# Emergence of putative energy parasites within *Clostridia* revealed by genome analysis of a novel endosymbiotic clade

**DOI:** 10.1101/2023.07.03.547156

**Authors:** Kazuki Takahashi, Hirokazu Kuwahara, Yutaro Horikawa, Kazuki Izawa, Daiki Kato, Tatsuya Inagaki, Masahiro Yuki, Moriya Ohkuma, Yuichi Hongoh

## Abstract

The *Clostridia* is a dominant bacterial class in the guts of various animals and are considered to nutritionally contribute to the animal host. Here, we discovered clostridial endosymbionts of cellulolytic protists in termite guts, which have never been reported with evidence. We obtained (near-)complete genome sequences of three endosymbiotic *Clostridia*, each associated with a different parabasalid protist species with various infection rates: *Trichonympha agilis*, *Pseudotrichonympha grassii*, and *Devescovina* sp. All these protists are previously known to harbor permanently-associated, mutualistic *Endomicrobia* or *Bacteroidales* that supplement nitrogenous compounds. The genomes of the endosymbiotic *Clostridia* were small in size (1.0–1.3 Mbp) and exhibited signatures of an obligately-intracellular parasite, such as an extremely limited capability to synthesize amino acids, cofactors, and nucleotides and a disrupted glycolytic pathway with no known net ATP-generating system. Instead, the genomes encoded ATP/ADP translocase and, interestingly, regulatory proteins that are unique to eukaryotes in general and are possibly used to interfere with host cellular processes. These three genomes formed a clade with metagenome-assembled genomes (MAGs) derived from the guts of other animals, including human and ruminants, and the MAGs shared the characteristics of parasites. Gene flux analysis suggested that the acquisition of the ATP/ADP translocase gene in a common ancestor was probably key to the emergence of this parasitic clade. Taken together, we provide novel insights into the multilayered symbiotic system in the termite gut by adding the presence of parasitism and present an example of the emergence of putative energy parasites from a dominant gut bacterial clade.

## Introduction

*Clostridia*, a class in the phylum *Bacillota* (*Firmicutes*), is one of the dominant bacterial taxa in the gastrointestinal tract of various animals and contributes to host nutrition via the degradation of food materials and homeostasis via the production of various compounds [1, 2]. For instance, two clusters of *Clostridia* corresponding to *Clostridium* clusters IV and XIVa [3] occupy a large part (10–40%) of human-gut bacterial microbiota, and their production of short-chain fatty acids and catecholamines contributes to immune homeostasis in the gut environment and gut-brain axis [2]. *Clostridia* are also dominant in the gut microbiota of various termite species [4–8]. Termites thrive exclusively on dead plant matter and play a key role in the terrestrial carbon cycle in temperate to tropical regions [9], while being notorious as pests of wooden buildings. The ability of termites to efficiently digest lignocellulose and survive on nitrogen-poor food is mostly attributable to the activities of their gut microbiota, comprising protists, bacteria, and archaea [10]. The protists play a central role in lignocellulose digestion in the gut of “lower”, or phylogenetically basal, termites, and *Clostridia* as well as other bacterial assemblages contribute in reductive acetogenesis [11], polysaccharide degradation [12, 13], nitrogen fixation [14], and nitrogen recycling via uric acid [15].

Certain members of *Clostridia* colonize the mucus layer of the human and murine guts [16], and in rumens, *Clostridia* attach to plant fibers and are involved in polysaccharide degradation [17]. In termite guts, previous 16S rRNA-based analyses revealed that *Clostridia* inhabit the gut wall or luminal fluid [18, 19]. In the gut of lower termites, the cellulolytic protists generally harbor phylogenetically diverse prokaryotic symbionts within the cytoplasm [20–24], nucleoplasm [25], and/or on the surface of the cell [26–32]. Many studies have reported the mutualistic functions of those endo- (i.e., intracellular) and ecto- (i.e., extracellularly attached) symbionts based on genome analyses. For example, “*Candidatus* Endomicrobium trichonymphae” (phylum *Elusimicrobiota*) is an obligate endosymbiont of the protist *Trichonympha agilis* and supplements various amino acids and cofactors [20]. “*Candidatus* Azobacteroides pseudotrichonymphae” (phylum *Bacteroidota*), an obligate endosymbiont of the protist *Pseudotrichonympha grassii*, fixes dinitrogen and recycles nitrogen via uric acid [21], and “*Candidatus* Armantifilum devescovinae” (*Bacteroidota*), an obligate ectosymbiont of the protist *Devescovina* spp., also fixes dinitrogen [32]. Although direct microscopic evidence for the presence of clostridial endosymbionts of termite-gut protists has never been reported, clostridial 16S rRNA gene sequences have occasionally been obtained from cell suspension of certain protist species [23, 33]. Apart from the termite gut microbiota, it has been reported that an endosymbiont of the aquatic ciliate *Metopus* sp. is closely related to “*Clostridium aminobutyricum*” (family “Anaerovoracaceae”) [34] and that an endosymbiont of the ciliate *Tryimyema compressum* belongs to the family *Syntrophomonadaceae* [35]. These two are only known examples of endosymbionts belonging to the class *Clostridia*, and no information on their functions is available.

Here, we report the discovery of endosymbiotic *Clostridia* within the cells of protists in termite guts and our examination of their ecology and evolution based on morphological, phylogenetic, and genomic characteristics. Our results indicated that the endosymbiotic *Clostridia* were obligately-intracellular parasites; this is the first report of parasites infecting cellulolytic protists in the termite gut with genome-based evidence. In addition, we found that these endosymbiotic *Clostridia* constituted a clade with metagenome-assembled genomes (MAGs) exhibiting similar parasitic traits from the gastrointestinal tracts of various animals, including humans and ruminants. Our results revealed another factor, i.e., parasitism, within the multilayered symbiotic system in the termite gut and extend the phylogenetic, ecological, and physiological diversity of *Clostridia*, providing novel insights into the evolutionary emergence of a parasitic lineage within a free-living bacterial clade.

## Materials and Methods

### Termites and protists

We collected three lower termite species: *Reticulitermes speratus* (family Rhinotermitidae) from Saitama Prefecture, Japan; *Coptotermes formosanus* (Rhinotermitidae) from Chiba, Kagoshima, and Okinawa Prefectures, Japan; and *Neotermes sugioi* (formerly *N. koshunensis*) (family Kalotermitidae) from Okinawa Prefecture, Japan. The entire gut of a worker termite was removed, and the gut contents were suspended in Trager’s solution U [36]. Single cells of the parabasalid protists *Trichonympha agilis*, *Pseudotrichonympha grassii*, and *Devescovina* sp. in the guts of *R. speratus*, *C. formosanus,* and *N. sugioi*, respectively, were physically isolated using a Leica AM6000 inverted micromanipulation microscope system.

### 16S rRNA gene sequencing and phylogenetic analysis

The isolated protist cells were subjected to whole-genome amplification (WGA) using the Illustra GenomiPhi V2 Kit (GE Healthcare) as described previously [37]. Near full-length 16S rRNA genes were amplified by PCR using Phusion Hi-Fidelity DNA Polymerase (New England BioLabs) and a bacteria-specific primer set, 27F-mix and 1492R or 1390R [29]. Purification of PCR products, cloning, and Sanger sequencing were performed as described previously [25].

Obtained clostridial sequences were aligned to reference sequences (Table S1) retrieved from the SILVA database r138 [38], the All-Species Living Tree Project database LTP_06_2022 [39], and the Type Strains Genome Database [40], using MAFFT v7.490 with the “linsi” option [41]. The 16S rRNA genes detected in MAGs (see Supplementary Methods) were also added to the dataset. Alignments were trimmed using TrimAL v1.4rev22 [42] with the “gappyout” option, and a maximum-likelihood tree was constructed using IQ-TREE with the GTR+I+G4 model and the ultrafast bootstrap and SH-aLRT tests with 1,000 replicates [43].

### Fluorescence in situ hybridization and transmission electron microscopy

Oligonucleotide probes specific to the respective clostridial 16S rRNA phylotypes were designed using ARB: RsTa-C01-76 (5′-GTGCCTTGCAAAACTCCG-3′), CfP3-15-656 (5′-CCGTTCGCCTCTACTTTAC-3′), and NkDv07-142 (5′-TCCATCAGCTATCTCCCAC-3′). Fluorescence in situ hybridization (FISH) was conducted as described previously [27, 44] with hybridization at 50°C for 1.5 h. Cells were observed under an Olympus BX51 epifluorescence microscope. Samples for transmission electron microscopy (TEM) were prepared as described previously [28, 45] and observed under a JEM-1400Plus transmission electron microscope (JEOL).

### DNA preparation for genome sequencing

Collected single protist cells were washed several times in solution U and then transferred to water droplets containing 0.01% Tween 20 (Nacalai Tesque, Kyoto, Japan). Bacterial cells that leaked from the ruptured host cell were collected with a glass capillary attached to the micromanipulation system and subjected to WGA using Thermo Scientific EquiPhi29 DNA Polymerase. The reaction was performed at 10-µL scale at 45℃ for 3 h (see Supplementary Methods for details). PCR was conducted using primer sets specific to the respective 16S rRNA phylotypes (Table S2), to check for the presence of their genomes. The WGA samples containing the target were re-amplified using EquiPhi29 DNA polymerase at 50-µL scale. The reaction was performed at 45℃ for 2 h.

### Genome sequencing

Genome sequencing was conducted on the MiSeq platform (Illumina) and the MinION platform (Oxford Nanopore Technologies). The libraries for MiSeq were prepared using the TruSeq DNA Sample Preparation Kit for low DNA input (Illumina). Sequencing was performed with the MiSeq Reagent Kit V3 (Illumina). Sequencing libraries for MinION were prepared using the SQK-LSK308 1D^2^ Sequencing Kit (Oxford Nanopore Technologies) for the NkDv07 sample and the SQK-LSK109 Ligation Sequencing Kit for RsTa-C01 and CfP3-15. The prepared libraries were loaded onto an R9.5 Spot-On Flow cell (FLO-MIN107) for NkDv07 and onto R10.3 Spot-On Flow cells (FLO-MIN109) for RsTa-C01 and CfP3-15. Detailed procedures for the MinION library preparation were described in Supplementary Methods.

### Quality filtering, assembly, and gene annotation

Adaptor removal and quality filtering of MiSeq reads were performed as described previously [26]. For MinION reads, base calling, adapter removal, and quality trimming, were performed using guppy v4.0.11, porechop v0.2.4 [46], and Nanofilt v2.7.1 [47], respectively. Quality-trimmed reads were assembled to construct contigs using SPAdes v3.14.1 [48] and Flye v2.8 [49] (Supplementary Methods). Automatic gene prediction and annotation were performed using DFAST [50], and the gene loci and annotations were manually curated based on the results of BLASTp searches of the NCBI non-redundant protein database. Metabolic pathways were reconstructed using the KEGG automatic annotation server (KAAS) [51] and KEGG Mapper [52]. The genomes were taxonomically classified using GTDB-tk v1.7.0 [53].

### Phylogenomics, COG analysis, and gene flux analysis

To construct a phylogenomic tree, we retrieved genome sequences of “Acutalibacteraceae” as references from GTDB r202 [54]. Also added were genome sequences assigned using GTDB-tk v1.7.0 to “Acutalibacteraceae” in the Unified Human Gastrointestinal Genome collection [55] and the Cow-rumen catalogue v1.0 (https://www.ebi.ac.uk/metagenomics/genome-catalogues/cow-rumen-v1-0) as well as MAGs detected in a recent large-scale analysis of the ruminant gut microbiome [56] and those from the termite gut microbiome [57]. These reference genome sequences were clustered based on the criterion of ≥95% average nucleotide identity (ANI), and representative genomes were selected using Galah (https://github.com/wwood/galah).

ANI and average amino acid identity (AAI) were calculated using ANI/AAI-Matrix [58]. Gene prediction was conducted using Prodigal v2.6.3 [59], and 31 single-copy gene markers used by Graham et al. (2018) [60] were extracted using the IdentifyHMM script (https://github.com/edgraham/PhylogenomicsWorkflow). The genome sequences with at least 16 marker genes were used to construct a phylogenomic tree (Table S3). Amino acid sequences were aligned using MAFFT v7.471 with default settings, concatenated, and trimmed using TrimAL v1.4rev22 with the “-automated1” option. A maximum-likelihood tree was constructed using IQ-TREE as above with a substitution model selected by its model finder.

The predicted genes and pseudogenes were classified into clusters of orthologous genes (COG) using RPS-BLAST v2.13.0+ with a cut-off e-value <10^−4^ [61], and each COG was classified into a COG functional category. For each genome, the relative abundance of the COG functional categories was calculated, and a centered log- ratio transformation was performed. The transformed data were used for principal component analysis (PCA), which was performed using a Scikit-learn python package [62]. All genes of “Acutalibacteraceae” genomes without COG annotation were subjected to de novo clustering using OrthoFinder v2.5.5 with default settings [63], and orthogroups were defined. Gene flux analysis was performed using Count (http://www.iro.umontreal.ca/~csuros/gene_content/count.html) [64] with Wagner parsimony, based on the matrix of COG and orthogroup and the phylogenetic tree obtained from the above analysis. The gene gain/loss penalty was set to 2.

### Phylogenetic analysis of protein-coding genes

For the phylogenetic analysis of ATP/ADP translocase genes, up to 100 reference sequences were identified by BLASTp searches of the UniProtKB database and were retrieved. Those used by Schmitz-Esser et al. (2004) [65] were added to the reference sequences (Table S4). After removing redundant sequences, the amino acid sequences were aligned using MAFFT v7.471 with “linsi” and trimmed with TrimAL v1.4rev22 using “gappyout”.

For eukaryotic-like genes, reference sequences were similarly retrieved (Table S5). For those genes in the RsTa-C01 genome, a tBLASTn search of the *T. agilis* draft genome sequence from an ongoing genome project was performed, and the aligned region of the top-hit sequence (LC770199–206) was added. For those in the CfP3-15 genome, BLASTp searches of a *P. grassii* transcriptome dataset [66] was performed, and the top-hit sequence was added. Alignments were carried out using T-Coffee [67] in the “expresso” mode and trimmed using TrimAL v1.4rev22 with “gappyout”. For genes for which alignment of the entire region was difficult, the most conserved region identified by the NCBI Conserved Domain Search was used for alignment. The sequences containing gaps in more than 50% of the aligned region were removed. Maximum-likelihood trees were constructed using IQ-TREE with settings as in the phylogenomic analysis.

### RNA extraction and RT-PCR

The gut contents of 30 *R. speratus* or *C. formosanus* workers were suspended in solution U. The suspensions were centrifuged, and the precipitates were subjected to total RNA extraction using the *Quick*-RNA Fungal/Bacterial Microprep Kit (Zymo Research). DNA removal, reverse transcription, and RT-PCR with specific primer sets for respective eukaryotic-like genes (Table S2), were conducted as described previously [68]. No specific amplifications were detected in RNA samples without reverse transcription. The RT-PCR products were cloned using the Invitrogen TOPO TA Cloning Kit and were sequenced as described above.

### Accession numbers

The 16S rRNA sequences of endosymbiotic *Clostridia* will appear under accession numbers LC760310–12. The genome sequence data obtained in this study have been deposited in DDBJ under the BioProject PRJDB15329.

## Results

### Localization, morphology, and phylogenetic positions of endosymbiotic *Clostridia*

During our 16S rRNA gene cloning analyses of bacterial microbiota associated with protist cells in termite guts, we identified three phylotypes affiliated with *Clostridia*: RsTa-C01 (LC760312) from *T. agilis* in an *R. speratus* gut, CfP3-15 (LC760310) from *P. grassii* in a *C. formosanus* gut, and NkDv07 (LC760311) from *Devescovina* sp. in an *N. sugioi* gut (see Supplementary Results for details). We performed FISH with specific probes and detected cocci exclusively within the cytoplasm of each host protist species (Figs. 1A–H). The rates of infection to the host protist species greatly varied, depending on the termite colony: 0–71% (RsTa-C01, surveyed termite colonies [n] = 7), 0–100% (CfP3-15, n = 10), and 0–18% (NkDv07, n = 14) (Table S6). A single host cell harbored ca. 40–100 cells of RsTa-C01, 10^2^–10^3^ cells of CfP3-15, and 10–100 cells of NkDv07. RsTa-C01 cells were mostly diplococci (Fig. 1E), while CfP3-15 were detected as single, diplo-, or triplococci (Fig. 1G), and NkDv07 as aggregates (Fig. 1H) or single cocci when its cell number was small (Figs. S1A, B). TEM images showed that both RsTa-C01 and NkDv07 frequently contained an endospore and that their cells or aggregates were surrounded by the host membrane (Figs. 1I–L). We also putatively identified CfP3-15 cells by TEM among the cells of “*Ca.* Azobacteroides pseudotrichonymphae” [21] within *P. grassii* cells (Figs. S1C–E).

**Figure 1.**
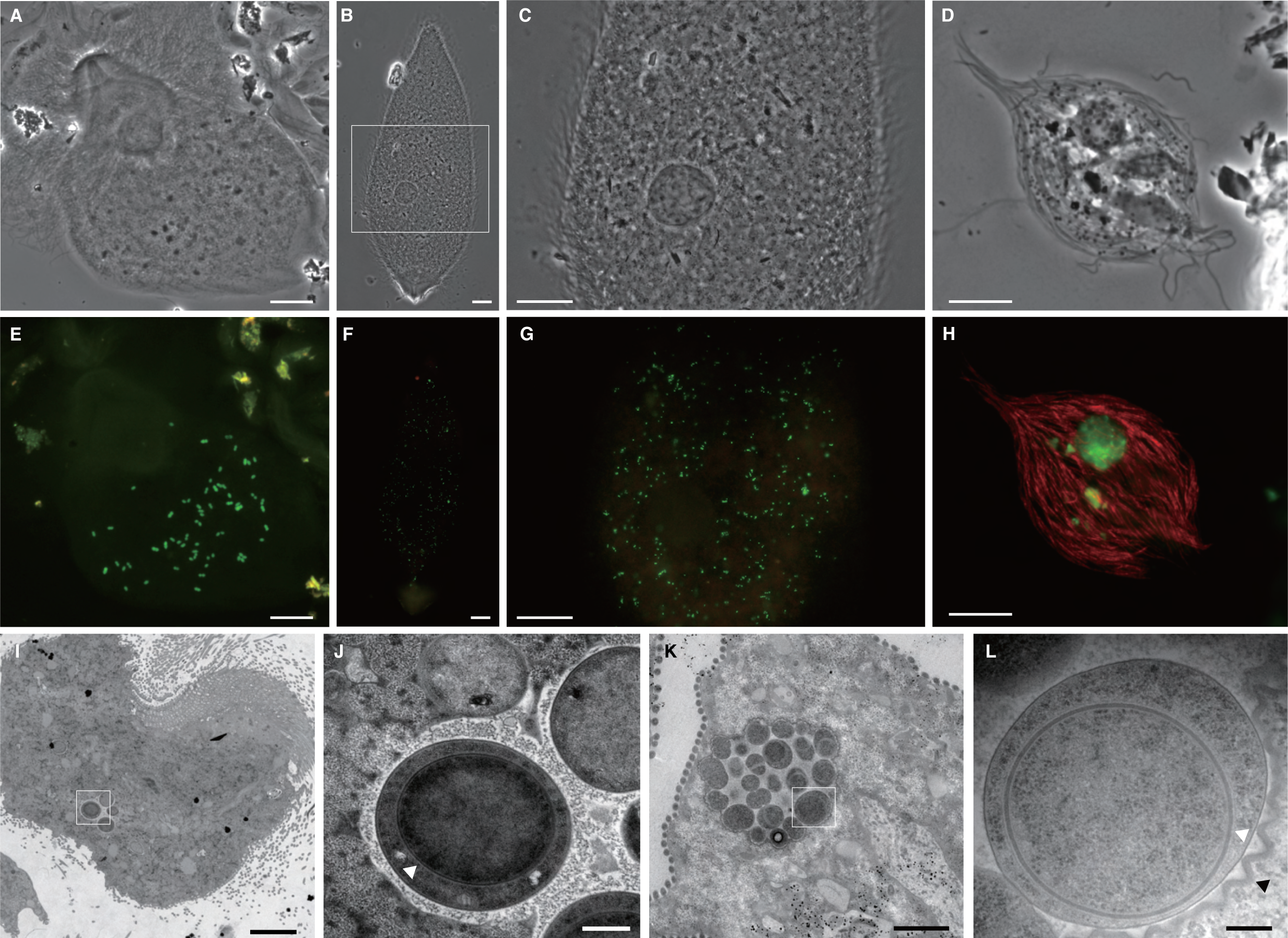
Fluorescence in situ hybridization (FISH) analysis of clostridial 16S rRNA phylotypes, RsTa-C01, CfP3-15, and NkDv07 (**A–H**) and transmission electron micrographs of endosymbionts of *Trichonympha agilis* and *Devescovina* sp. (**I–K**). *T. agilis* from the gut of *Reticulitermes speratus* (**A, E, I, J**), *Pseudotrichonympha grassii* from the gut of *Coptotermes formosanus* (**B, C, F, G**), and *Devescovina* sp. from the gut of *Neotermes sugioi* (**D, H, K, L**) were observed. Endosymbiotic clostridial cells were detected with oligonucleotide probes specific to each phylotype (6FAM-labelled, green), and “*Candidatus* Armantifilum” cells [28, 99] were detected with probe Bactd-937 (5′-CCACATGTTCCTCCGCTT-3′) (Texas red-labelled, red) [100]. (**C, G**) Magnified images of the area enclosed by a rectangle in panel B. (**J, L**) Magnified images of the area indicated in panels **I** and **K,** respectively. White arrowheads indicate an endospore, and black arrowhead indicates a membrane structure. Bars: (**A**–**C, E–G**), 20 μm; (**D, H**), 10 µm; (**I**), 5 µm; (**J**), 500 nm; (**K**), 2 µm; (**L**), 200 nm.

Phylogenetic analysis based on the 16S rRNA gene revealed that these endosymbionts belonged to *Clostridium* cluster IV (or the *Clostridium leptum* group) [3] within the family *Oscillospiraceae* (Figs. 2 and S2A). This cluster is one of the dominant groups in the termite gut [4–8] as well as in the human and cattle gastrointestinal tracts [69, 70]. The endosymbionts formed a monophyletic group with sequences derived from the gastrointestinal tracts of various animals (e.g., AB746684), wastewater reactors (GQ134270), denitrifying biofilm (KJ399537), and clones from *Trichonympha* cells in *Reticulitermes santonensis* guts [23] (Figs. 2 and S2B).

**Figure 2.**
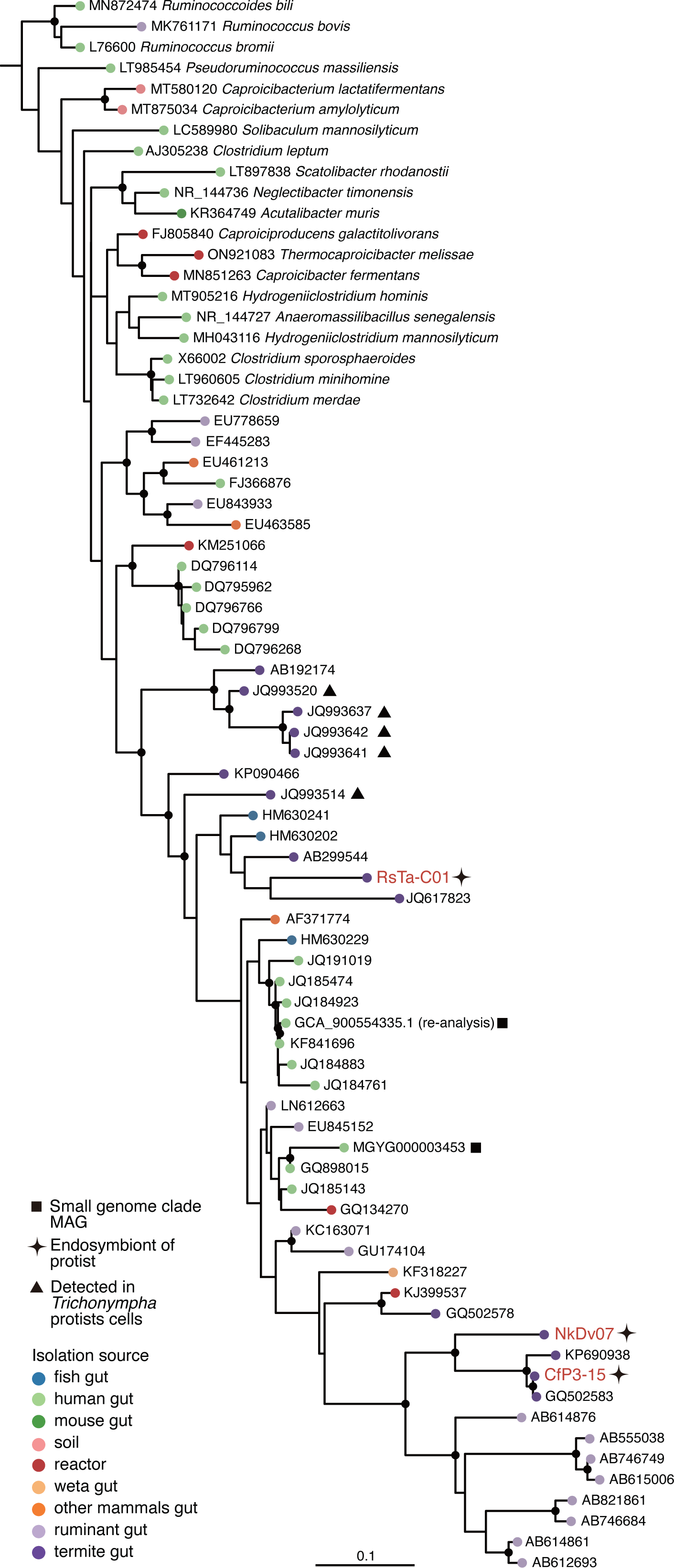
Phylogenetic positions of endosymbiotic *Clostridia* based on 16S rRNA gene sequences. Described species of “Acutalibacteraceae” in the GTDB taxonomy and uncultured clones closely related to endosymbiotic *Clostridia* were included in the tree. Maximum-likelihood tree was constructed with the GTR+I+G4 nucleotide substitution model and rooted with *Sporobacter termitidis* (Z49863) and *Oscillibacter ruminantium* (JF750939) as the outgroup. Highly supported nodes (SH-aLRT value ≥ 80% and ultrafast bootstrap support value ≥ 95%) are indicated with a closed circle.

### Overview and comparison of genome features

To predict the functions of these three endosymbiotic *Clostridia*, we obtained the complete genome sequences of RsTa-C01 and CfP3-15 and a draft genome sequence of NkDv07. To estimate the completeness of the NkDv07 genome consisting of four contigs, we modified the universal set of bacterial marker genes used by CheckM [71] according to the complete CfP3-15 genome, as shown in Table S7. The estimated completeness was 100% with no contaminating sequence detected, using the modified marker gene set. All MAGs simultaneously obtained from the WGA samples are listed in Table S8. The basic genomic features of the three endosymbiotic *Clostridia* are shown in Table 1 in comparison with *Clostridium leptum* DSM753 and *Ruminococcus bromii* ATCC27255, which are well-characterized gut bacteria among most closely related described species within *Clostridium* cluster IV (Figs. 2 and S2). The genome sizes of these three endosymbionts were small (1.0–1.3 Mbp), and CfP3-15 boasted the smallest genome (1,007,634 bp) among the known complete genome sequences of *Clostridia* (Fig. S3). Accordingly, the number of protein-coding sequences (CDSs) was much fewer (790–839) than in the free-living members of *Oscillospiraceae* (ca. 1,500–4,700). The GC content was also lower than those of the majority of *Clostridia* (Fig. S3). The GC skew profile of RsTa-C01 did not exhibit clear shifts that generally enable the identification of the replication origin and terminus site (Fig. S4).

**Table 1.**
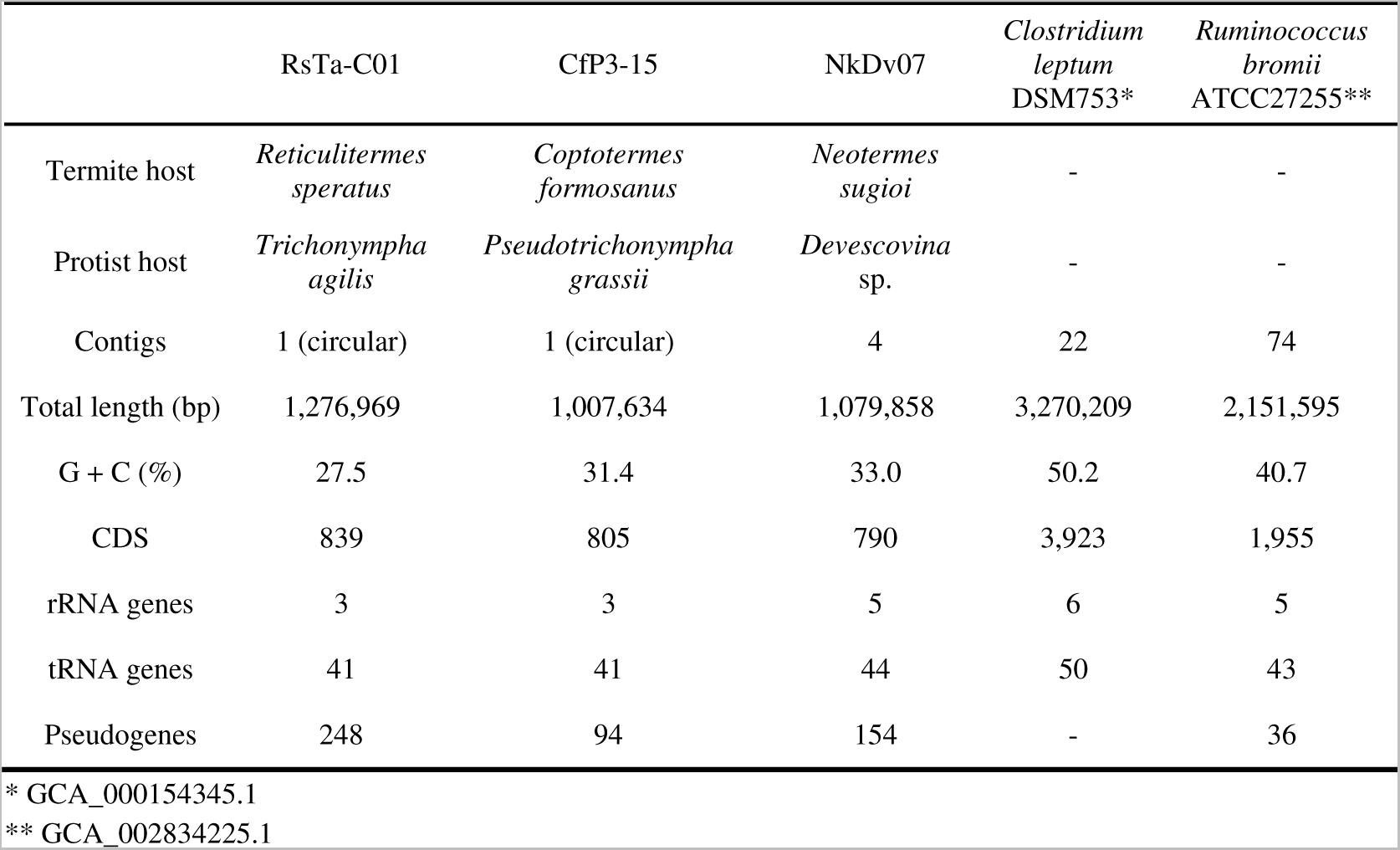
General genome features of endosymbiotic Clostridia obtained in this study and comparison with cultured relatives.

The predicted CDSs were classified into COG functional categories and compared (Fig. S5). The distribution patterns of CDSs were very similar among the three endosymbionts, and the number of genes in most categories were reduced to approximately a half to one-tenth of their free-living relatives (Fig. S5A). In particular, the reduction in categories E (amino acid transport and metabolism), G (carbohydrate transport and metabolism), and K (transcription), was prominent; the proportion of genes was reduced to 0.4–0.6 of the free-living relatives (Fig. S5B). Important genes involved in DNA replication were missing; CfP3-15 and NkDv07 lacked the *polA* gene for DNA polymerase I, and RsTa-C01 lacked *dnaA*, *rnhA,* and *recG*, encoding chromosomal replication initiator protein DnaA, ribonuclease HI, and ATP-dependent DNA helicase RecG, respectively. The three genomes contained 94–248 pseudogenes (Tables 1 and S9), and the pseudogenes were classified into mainly categories X (mobilome: prophages, transposons) and L (replication, recombination, and repair) (Fig. S6). No CRISPR-Cas system was found in these three genomes.

Phylogenomic analysis based on conserved single-copy genes inferred that the endosymbiotic *Clostridia* belonged to the family “Acutalibacteraceae” in the GTDB taxonomy, which corresponds to a portion of the validly approved family *Oscillospiraceae*. “Acutalibacteraceae” contain many cultured species (Figs. 2 and S2A) and MAGs from the gastrointestinal tracts of various animal species, and this is the first report of intracellular symbionts in this group.

### Predicted metabolism of endosymbiotic *Clostridia*

The predicted metabolic capacities based on the genome sequences of NkDv07 and CfP3-15 were very similar, while that of RsTa-C01 exhibited substantial differences from the other two (Fig. 3). Although all three genomes encoded a phosphotransferase system (PTS) that functions to import and phosphorylate glucose to glucose-6-phosphate (G6P), the glycolytic pathway is fragmented and cannot be used to generate ATP from glucose (Fig. 3). All three genomes encoded isomerase, which converts G6P to fructose-6-phosphate, a precursor for peptidoglycan biosynthesis. However, the RsTa-C01 genome lacked phosphofructokinase and fructose-bisphosphate aldolase; only CfP3-15 and NkDv07 can produce dihydroxyacetone phosphate (DHAP) from glucose, which is a precursor for glycerophospholipid biosynthesis. All three genomes lacked the gene for enolase, and therefore phosphoenolpyruvate (PEP), which is required for the activity of PTS, must be produced from pyruvate by the action of pyruvate-phosphate dikinase with ATP hydrolysis or from oxaloacetate by the action of phosphoenolpyruvate carboxykinase with GTP hydrolysis. Nevertheless, the genes for these enzymes were found only in the genomes of CfP3-15 and NkDv07 and were not identified in the RsTa-C01 genome. RsTa-C01 retained the pathway from 2-phosphoglycerate to DHAP (Fig. 3); thus, RsTa-C01 apparently needs to import phosphorylated glycerate to synthesize glycerophospholipids and needs to import PEP to activate PTS, although the associated transporters were not identified in its genome.

**Figure 3.**
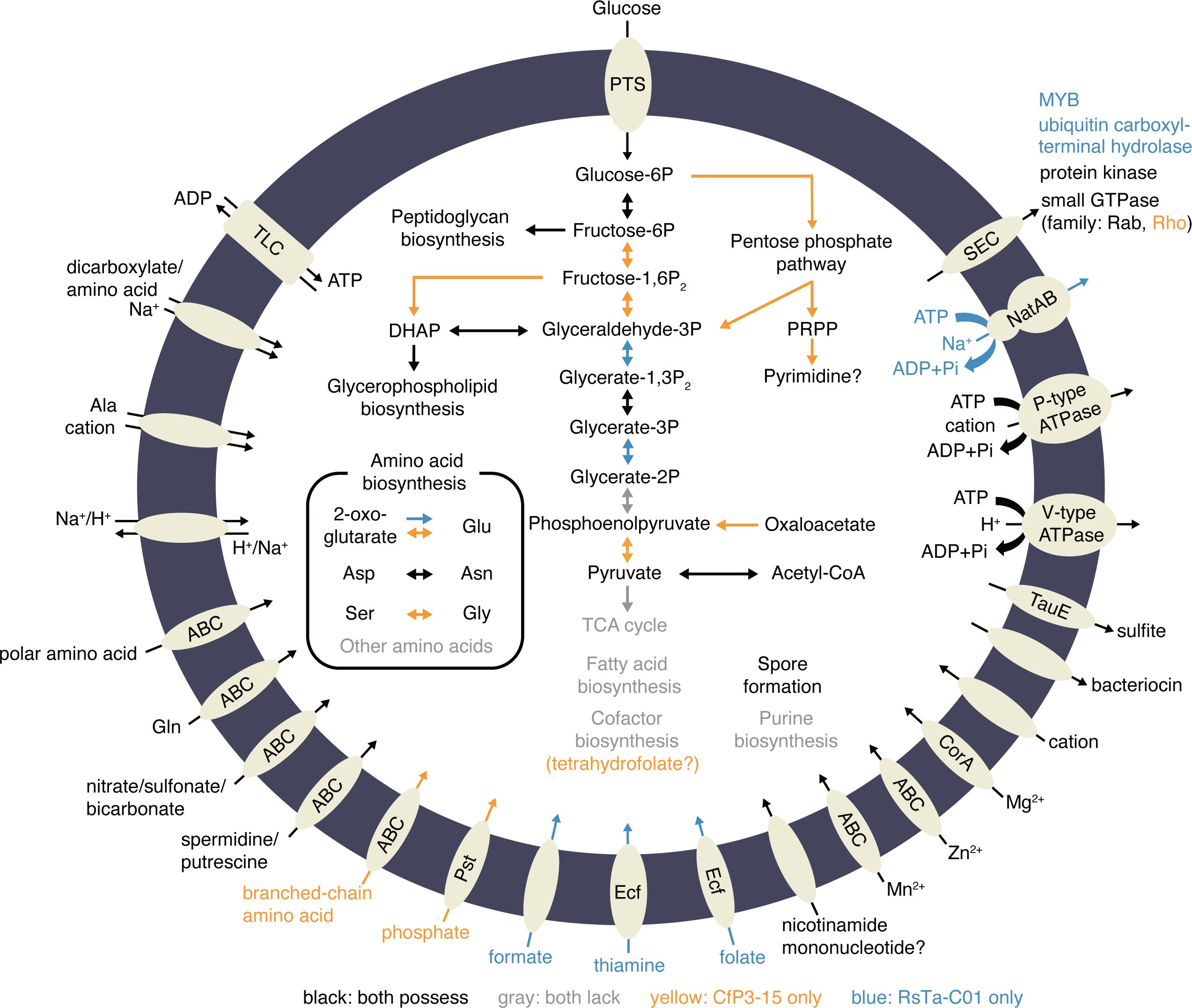
Predicted metabolic pathways of RsTa-C01 and CfP3-15. The metabolic capacity of NkDv07 is not shown because of its substantial similarity to that of CfP3-15. Pathways found only in the RsTa-C01 genome are highlighted with blue, and those only in the CfP3-15 genome are highlighted with yellow. Incomplete pathways are highlighted with grey.

Certain species of *Clostridia* possess the Rnf complex and/or energy-converting hydrogenase for ATP production [72]; however, the related genes were not identified in any of the three genomes. Alternative ATP-generating pathways, such as the Entner-Doudoroff pathway, Stickland reaction, and arginine dihydrolase pathway, were also not identified in these genomes. Instead, the gene encoding ATP/ADP translocase, a transporter that imports ATP and exports ADP and phosphate, was found in all three genomes. Important amino acid residues for nucleotide transport [73] were conserved in these homologues (Fig. S7). The apparent absence of net ATP-generating system and the presence of ATP/ADP translocase strongly suggest that the endosymbiotic *Clostridia* rely on ATP imported from the host. This transporter may be used also for other nucleotides [74]. The transcription of the ATP/ADP translocase genes of CfP3-15 and RsTa-C01 was confirmed by RT-PCR (Fig. S8). We did not attempt RT-PCR for NkDv07 because we lost the termite colony that contained this endosymbiont before the experiment.

CfP3-15 and NkDv07, but not RsTa-C01, retained the non-oxidative pentose phosphate pathway used to synthesize phosphoribosyl pyrophosphate (PRPP) (Fig. 3). However, CfP3-15 and NkDv07 did not possess the genes required for the de novo biosynthesis of purines and can only produce UMP/UDP/uridine for pyrimidine biosynthesis. Almost no biosynthetic pathways for amino acids or cofactors were present in the three genomes (Fig. 3). Each genome encoded several transporters for amino acids and cofactors. For example, the RsTa-C01 genome encoded transporters for formate and folate, which are used as substrates to synthesize 10-formyl-tetrahydrofolate. Although CfP3-15 and NkDv07 did not possess these transporters, their genomes encoded the biosynthetic pathway used for obtaining tetrahydrofolate from GTP. No known functions beneficial for the protist and/or termite hosts were identified in all three genomes, including nitrogen fixation/recycling [21, 22, 24, 31, 32, 75], reductive acetogenesis [22, 24], hydrogen removal [21, 22, 24, 26, 76, 77], polysaccharide degradation [31, 77, 78], and nutrient production [20–22, 24, 31, 75, 79].

### Eukaryotic-like protein-coding genes

Several genes unique to eukaryotes in general were identified in all three genomes (Table S10). For example, the CfP3-15 genome contained 18 genes for small GTPase, mainly of the Rab family, which are involved in intracellular trafficking, cell motility, and signal transduction in eukaryotic cells [80]. These proteins, such as small GTPases, Ser/Thr protein kinases, and Myb-like transcription factors, have been found almost exclusively in eukaryotes and several pathogenic bacterial clades, including *Legionella* [81] and *Chlamydiota* [82]. Among them, genes for Rab-GTPase have been found almost exclusively in *Legionella* outside of eukaryotes and never in the *Clostridia* class. The repertoires of these genes for eukaryotic-like proteins differed among the three genomes; the genes were acquired independently. Phylogenetic analyses indicated that many of the genes were closely related to those of protists, and at least 12 genes were most closely related to those of parabasalids (Figs. 4 and S9, Table S10). A Ser/Thr kinase of RsTa-C01 was phylogenetically closest to that of *Trichonympha agilis* (LC770206) (Fig. 4A), implying that this gene was acquired horizontally from the host protist. It was predicted that at least 20 of the 42 eukaryotic-like protein-coding genes contained signal peptides (Table S10); these are most probably secreted outside the cell through the Sec system.

**Figure 4.**
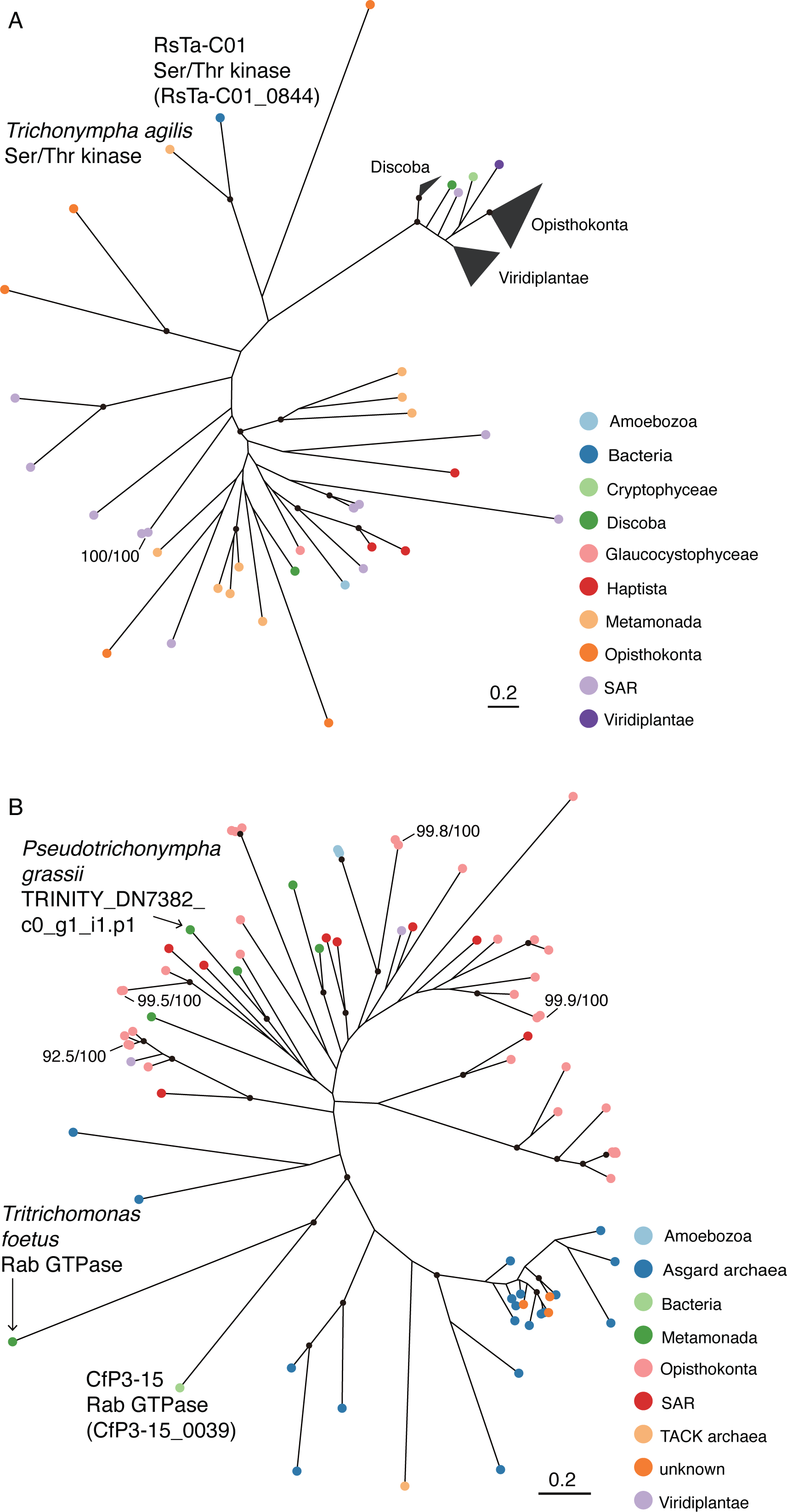
Phylogenetic positions of eukaryotic-like genes of RsTa-C01 and CfP3-15. (**A**) Phylogenetic relationships of Ser/Thr kinase of RsTa-C01 and its homologs. (**B**) Phylogenetic relationships of Rab-GTPase of CfP3-15 and its homologs. Maximum likelihood trees were generated using the LG+I+G4 amino acid substitution model. Highly supported nodes (SH-aLRT value ≥ 80% and ultrafast bootstrap support value ≥ 95%) are indicated with a closed circle.

Transcription of three out of ten eukaryotic-like genes tested was confirmed by RT-PCR and Sanger sequencing (Fig. S10, Table S10). The failure to detect the other transcripts may be due to intraspecific genetic variations because we had to use different termite colonies from different geographic locations for the RT-PCR experiments after genome sequencing analysis. Indeed, the sequence obtained for a small GTPase gene contained multiple single-nucleotide polymorphisms against the genome sequence (Fig. S11).

Phylogenetic and comparative genome analysis within the family “Acutalibacteraceae”

The phylogenetic positions of the three endosymbionts in the family “Acutalibacteraceae” were analyzed based on conserved single-copy genes (Fig. 5A, Table S3). The three genomes formed a monophyletic group with 23 MAGs from the gastrointestinal tracts of several animal species, including a termite, ruminants, and humans. This monophyletic group was divided into at least six genus-level clades based on GTDB r207 (Fig. 5A). Genomes of each genus-level clade shared at least 50% AAI, although ANI was occasionally below 70%, including the ANI among the three endosymbionts (Fig. S12). The predicted genome size and GC content of these MAGs were similar to those of the three endosymbionts: 0.86–1.4 Mbp (except for MGYG000004496: 1.9 Mb) and 30–39%, respectively (Fig. 5B, Table S11). This monophyletic group is designated hereafter the “small genome clade”, the genomic features and host information for which are summarized in Table S11.

**Figure 5.**
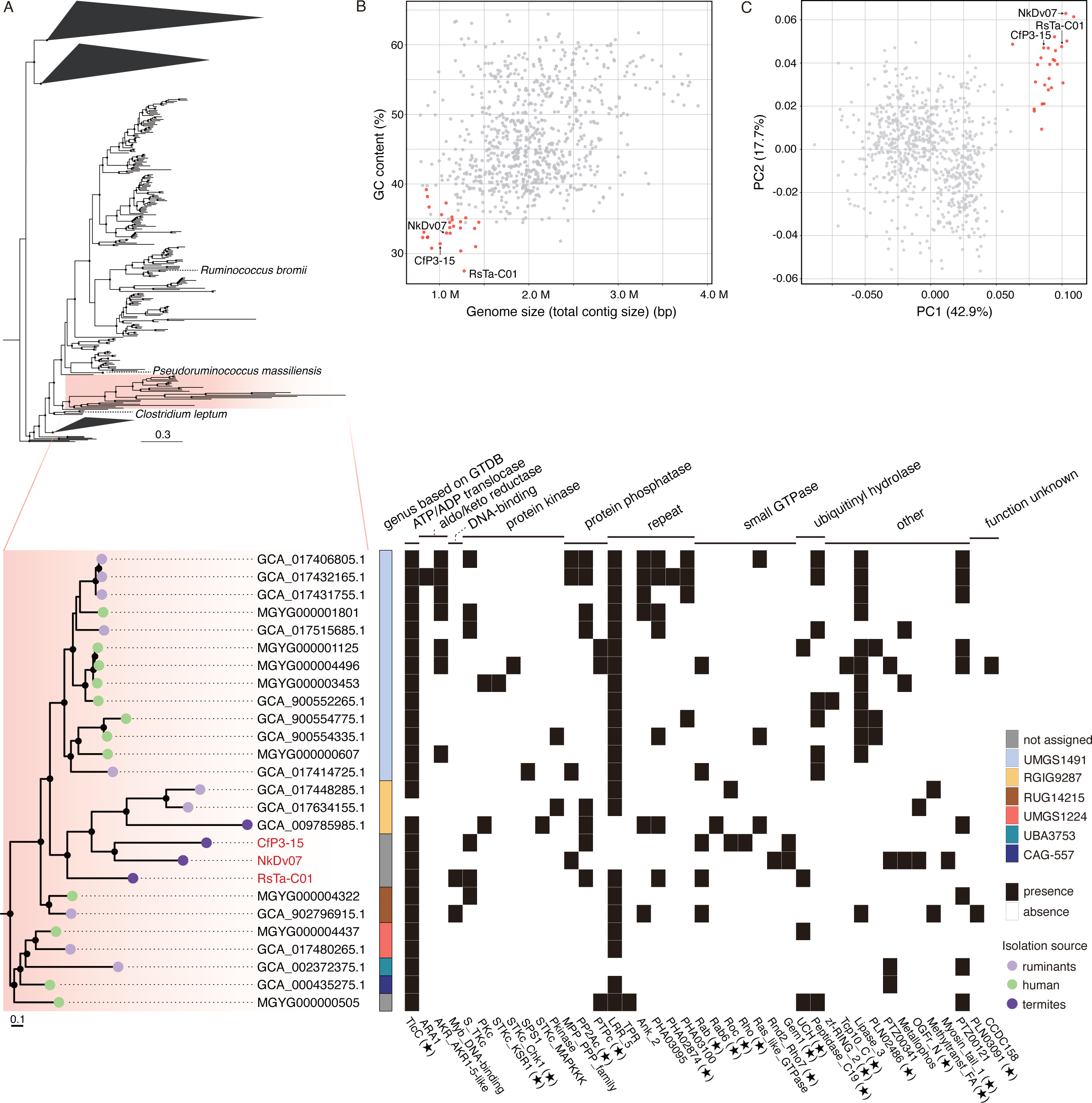
Genomic signatures of *Clostridia* belonging to the small genome clade. (**A**) Phylogenomic tree of bacteria belonging to “Acutalibacteraceae” based on concatenated amino acid sequences of single-copy marker genes. Maximum likelihood tree was constructed using the LG+F+I+G4 amino acid substitution model and rooted with *Syntrophomonas wolfei* (GCF_000014725.1) and *Thermosyntropha lipolytica* (GCF_900129805.1) as the outgroup. Highly supported nodes (SH-aLRT value ≥ 80% and ultrafast bootstrap support value ≥ 95%) are indicated with a closed circle. The small genome clade is magnified, and the genus-level clades based on GTDB r207 and distribution of ATP/ADP translocase (TlcC) and eukaryotic-like domains (Tables S10 and S12) are also depicted. The domains of protein-coding genes with a star were detected only in the small genome clade among “Acutalibacteraceae” genomes. (**B**) Relationships between genome size (or total contig lengths of draft genomes) and GC content. (**C**) Principal component analysis of genomes based on the distribution patterns of the COG functional categories. Genomes of the small genome clade are highlighted in red in **B** and **C**.

The COG distribution patterns were compared within “Acutalibacteraceae” (Fig. 5C). The genomes of the small genome clade clustered together, with a characteristic reduction in the proportion of genes related to metabolisms, especially G (carbohydrate transport and metabolism), and an increase in those related to J (translation, ribosomal structure, and biogenesis) (Fig. S13). Several genes containing eukaryotic-like domains were also identified from MAGs of the small genome clade. The identified domains included those of small GTPases, protein kinases, protein phosphatases, and ubiquitinyl hydrolases (Fig. 5A, Table S12).

### Gene flux analysis

To estimate the loss and gain events of gene families in the ancestors of the small genome clade, we conducted the gene flux analysis. The results suggested that the last common ancestor of the small genome clade gained the genes for ATP/ADP translocase (COG3202), 3-methyladenine DNA glycosylase Mpg (COG2094), serine transporter YbeC (COG0531), and others (Table S13). Among these, genes for ATP/ADP translocase and hypothetical proteins OG0003398, OG0004343, and OG0005234 were exclusively found in the small genome clade, whereas the others were also sparsely found in other clades in “Acutalibacteraceae” (Fig. S14). Of these, the ATP/ADP translocase gene was retained by all members of the small genome clade except for GCA_017634155.1 (Fig. 5A). The three hypothetical proteins were absent from 30–70% of genomes in the small genome clade, including the complete genomes of RsTa-C01 and CfP3-15 (Fig. S14).

The ATP/ADP translocase genes from the small genome clade formed two monophyletic clusters; one was a sister group of a clade mainly comprising those from *Alphaproteobacteria* and *Chlamydiae* (designated *Clostridia* ATP/ADP translocase I), and the other was in a large clade mainly consisting of those from “*Candidatus* Dependentiae” and *Chlamydiae* (*Clostridia* ATP/ADP translocase II) (Fig. 6). All genomes in the small genome clade, except GCA_017634155.1, encoded *Clostridia* ATP/ADP translocase I, and only a portion of them additionally possessed the other gene. Because the tree topology of *Clostridia* ATP/ADP translocase I was largely congruent with the species tree (Fig. S15A), the ATP/ADP translocase gene acquired by the last common ancestor of the small genome clade was most probably *Clostridia* ATP/ADP translocase I. *Clostridia* ATP/ADP translocase II was possibly acquired from a member of “*Ca.* Dependentiae” or *Chlamydiae* after the small genome clade had diversified (Figs.6 and S15B).

**Figure 6.**
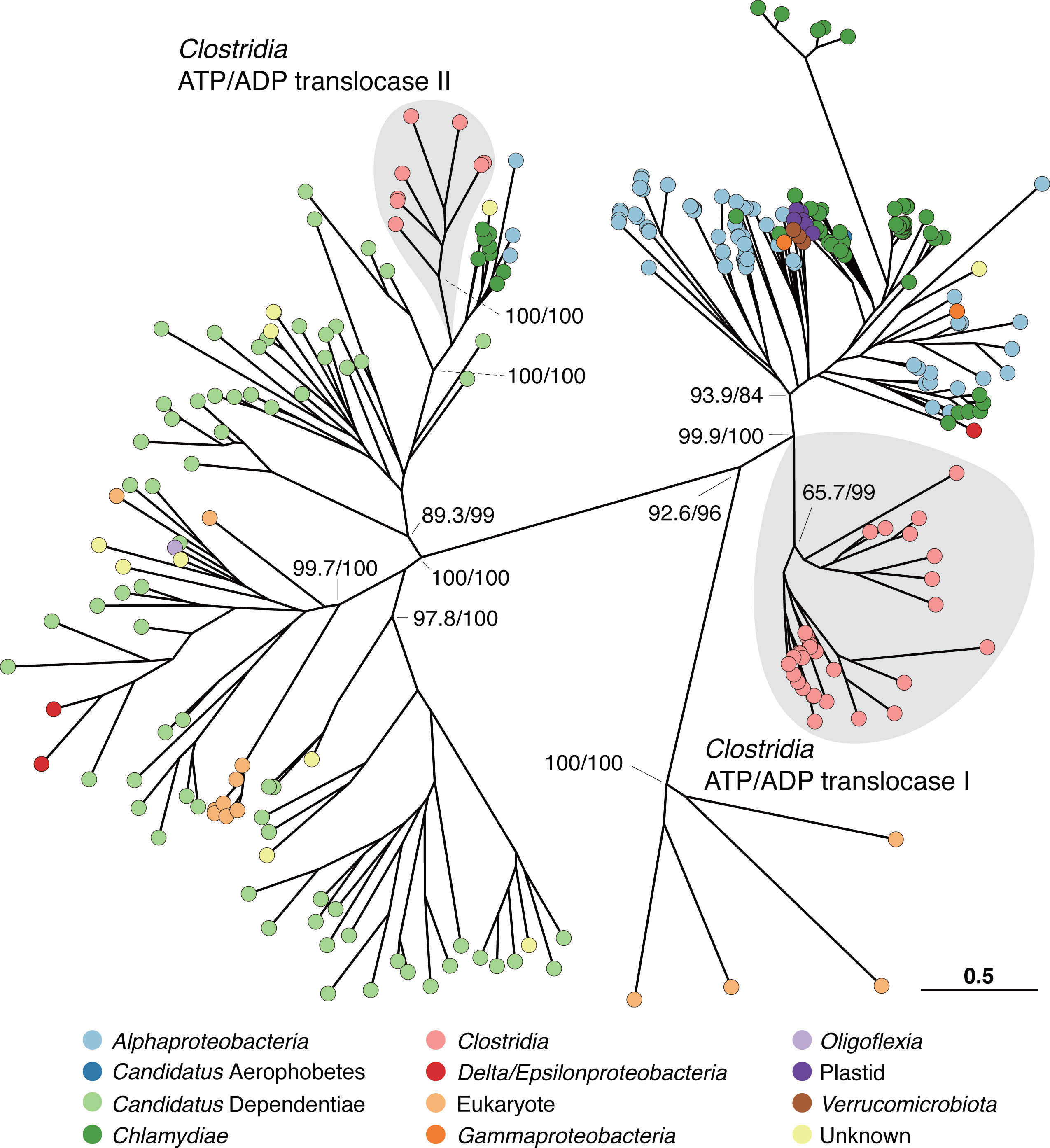
Phylogenetic positions of ATP/ADP translocase genes of the small genome clade. Maximum likelihood tree was constructed using the LG+F+G4 amino acid substitution model. Only support values (SH-aLRT/ultrafast bootstrap support values) for major clades are shown for clarity.

## Discussion

In this study, we discovered and analyzed the genomes of endosymbiotic *Clostridia* found exclusively within the cells of cellulolytic protists in termite guts. The total dependence of nutrition on their protist hosts, i.e., the lack of most biosynthetic pathways of nucleotides, amino acids, cofactors, and of systems to generate ATP, made it obvious that these *Clostridia* are obligately-intracellular parasites unable to reproduce outside the host cell, although the possibility that they possess certain beneficial functions not evident from the annotated genes cannot be excluded. To our knowledge, they are the most host-dependent *Clostridia* with the smallest genome sizes. The obligately parasitic life style of the endosymbiotic *Clostridia* contrasts with diverse endo- and ectosymbiotic bacteria of termite-gut protists studied thus far based on genome sequences. All these previously analyzed symbionts were suggested to be mutualists that critically contribute to the nutrition and metabolism of their protist hosts [20–22, 26, 31, 32, 75–79]. It seems likely that the endosymbiotic *Clostridia* have exploited these protist-bacteria mutualistic complexes as resources.

The habitat of this clostridial parasitic clade is not confined within the gut of lower termites; we found that the small genome clade is widely distributed in the gastrointestinal tracts of various animals, including “higher” (or phylogenetically apical) termites with no cellulolytic gut protists, humans throughout the world, and ruminants such as cattle, sheep, and deer (Figs. 5A and S16, Table S11). Although their localizations remain unknown in the gastrointestinal tract of these animals, higher termites can house a small number of protists in their gut, including *Nyctotherus* ciliates and *Trichomonas*-like parabasalids [83] and human guts can harbor commensal or pathogenic genera such as *Trichomonas*, *Entamoeba,* and *Giardia* [84]. In addition, clostridia belonging to the families *Oscillospiraceae* and *Lachnospiraceae* were reportedly enriched in ciliate fractions compared to protist-free bacterial fractions in the 16S rRNA gene amplicon sequencing analysis of cow ruminal contents [85]. Indeed, we confirmed the presence of 16S rRNA gene sequences related to those of the endosymbiotic *Clostridia* from ciliate fractions (Fig. S16), whereas no evidence was provided by FISH or TEM in a previous study [85]. Thus, these protists might be the hosts for the clostridia of the small genome clade. Previous reports focused on prey-predator relationships between eukaryotes and prokaryotes in the gastrointestinal tract of mammals [86, 87]; intracellular parasitism should also be considered.

The evolutionary emergence of this parasitic clade in a dominant gut bacterial group is likely attributable to the horizontal acquisition of ATP/ADP translocase, possibly from alphaproteobacteria, by the last common ancestor of the small genome clade, as suggested by our gene flux analysis (Fig. 6). Microsporidia, eukaryotic parasites, were previously suggested to have acquired the ATP/ADP translocase gene at an early stage in the transition to an intracellular lifestyle [88]. Because the acquisition of ATP/ADP translocase is also inferred in an early evolutionary stage of obligate host association of *Rickettsiales* [89], it is assumed that the acquisition of this gene was significant in establishing parasitism in organisms, irrespective of Gram-negative bacteria, Gram-positive bacteria, or eukaryotes. After intracellular parasitism emerged by the acquisition of ATP/ADP translocase, members of the small genome clade appear to have taken a common strategy: they independently acquired genes for eukaryotic-like regulatory proteins possibly from their protist hosts. These proteins may be used to adapt to their respective host cell environments by manipulating the host cellular processes. For example, the eukaryotic-like Ser/Thr protein kinase of diverse pathogenic bacteria reportedly disrupts various host cell functions, such as the innate immune system and the formation of actin cytoskeleton [90]. Although experimental evidence is absent, it has been hypothesized that Rab-GTPase proteins mimic and disrupt host cell functions [81]. Thus, it seems likely that convergent evolution has occurred in the small genome clade as seen in previously known, other intracellular parasites.

Evolutionary convergence was also observed in the DNA replication system of the small genome clade as endosymbionts; CfP3-15 and NkDv07 lack *polA*, and RsTa-C01 lacks *dnaA*, *rnhA,* and *recG*. The absence of *polA* has been reported in several mutualistic endosymbionts of insects [91] and the endosymbiotic cyanobacterium (chromatophore) of the photosynthetic protist *Paulinella* [92]. Possibly, slow-growing bacteria, including endosymbionts, can withstand the absence of PolA, as shown in an *Escherichia coli* strain lacking *polA* [93, 94]. The *dnaA* gene is also absent or pseudogenized in certain endosymbiotic bacteria of insects [95], water ferns [96], and a protist in the termite gut [20, 68]. The lack of this gene might be a consequence of streamlining adaptations along with the use of other replication mechanisms independent of DnaA [94]. The absence of *dnaA* is consistent with the atypical GC skew pattern of RsTa-C01 (Fig. S4), as seen in the *dnaA*-pseudogenized genome of *“Ca.* Endomicrobium trichonymphae” [20]. On the other hand, because double mutants of *rnhA* and *recG* are lethal in *E. coli* [97, 98], it is unclear how RsTa-C01 endures under the absence of these genes. Further studies of members of the small genome clade in environments other than termite guts will deepen our understanding of the ecology and evolution of this novel parasitic clade.

Here, we propose the novel genera and species names “*Candidatus* Improbicoccus pseudotrichonymphae”, “*Candidatus* Improbicoccus devescovinae”, and “*Candidatus* Paraimprobicoccus trichonymphae”, for CfP3-15, NkDv07, and RsTa-C01, respectively, based on their phylogeny, morphology, and genome sequences. The details are described in Supplementary Results.

## Acknowledgements

We are grateful to Gaku Tokuda, Kumiko Kihara, and Yukihiro Kinjo for cooperation in termite collection. We also thank Michiru Shimizu, Katsura Igai, and Tomoyuki Sato for assisting with experiments, and Osamu Kanazawa for checking the Latin grammar of scientific names. Sanger sequencing was done in the RIKEN BSI and in the Biomaterial Analysis Center in the Tokyo Institute of Technology. TEM was assisted by Keiko Ikeda in the latter facility. This study was financially supported by NEXT and KAKENHI grants from JSPS to YH (GS009, 22241046, 16H04840, 20H02897, 20H05584, 22K19342) and to MO (17H01447 and 19H05689), by JST-CREST (14532219) to YH, and also by JST SPRING (JPMJSP2106) to KT.

## Author contributions

YH and MO conceived research and provided equipment and reagents; YH and KT designed research; YH, TI, HK, and KT collected termites; KT, HK, and MY sequenced genomes; KT, YHori, and YH performed FISH and Sanger sequencing; KI, HK, and KT performed TEM; KT, YH, DK, and TI analyzed data; KT and YH wrote the paper with input from all authors.

## Compliance with ethical standards

