## Supplementary Materials for "Emergence of putative energy parasites within *Clostridia* revealed by genome analysis of a novel endosymbiotic clade"

**Supplementary Text**

**Supplementary Methods**

Whole genome amplification

The collected bacterial cells were transferred to 0.2-mL PCR tube containing 0.4-μL sterile double-distilled water and were subjected to cell lysis and denaturing double strand DNA using 0.5 μL of lysis buffer (0.4 M KOH, 100 mM dithiothreitol, 10 mM EDTA). After neutralization with 0.5 μL of buffer (400 mM HCl, 600 mM Tris-HCl, pH7.5), 10 μL reaction mixture containing 0.2 μL of EquiPhi29 DNA Polymerase, 1 μL of Reaction Buffer in the kit, 0.1 μL of 100 mM DTT, 0.5 μL of 500 μM exo-resistant random primer (Thermo Scientific) and 0.4 μL of 25 mM dNTP (Thermo Scientific) were added, and isothermal whole genome amplification (WGA) was performed at 45°C for 3 h. All plasticware and reagents except for the enzyme and random hexamer were UV-irradiated to decontaminate DNA.

Sequence library preparation for the MinION platform

Before preparing libraries for MinION, we conducted debranching and single-strand DNA digestion of WGA products using EquiPhi29 DNA polymerase and S1 nuclease (Takara Bio, Shiga, Japan), respectively [1]. After the treatment, DNA was purified by ethanol precipitation and fragmented using the Covaris g-TUBE for the RsTa-C01 and CfP3-15 samples. DNA fragments were separated by agarose gel electrophoresis, and those longer than 2 kbp for NkDv07 and 2–12 kbp for RsTa-C01 and CfP3-15 were excised and purified using the Zymoclean Large Fragment DNA Recovery Kit (Zymo Research). Sequencing libraries were then prepared and loaded onto flow cells as described in the main text.

Genome assembly

- RsTa-C01

MiSeq reads were trimmed and quality-filtered using the cutadapt [2] and prinseq [3] programs, respectively. Unpaired reads were removed using cmpfastq_pe (http://compbio.brc.iop.kcl.ac.uk/software/download/cmpfastq_pe). Quality-trimmed reads were then assembled using SPAdes v3.14.1 [4] with '--meta' option. The obtained contigs were designated “Contig-set 1”. Contig-set 1 was binned into respective bacterial assemblages using MyCC_2017.ova [5] with the “56mer” option. A bin consisted of 28 contigs and containing the 16S rRNA gene of RsTa-C01 was obtained. Among the contigs not included in the RsTa-C01 bin, we detected a high-coverage circular contig (36 kb) with high sequence similarity to certain contigs in the RsTa-C01 bin. Based on the gene repertories of this circular contig, we inferred that it was the complete genome sequence of a phage infecting RsTa-C01. We considered that the high sequence similarity between this phage genome and certain RsTa-C01 genome regions caused the low contiguity of the RsTa-C01 genome assembly. We therefore attempted re-assembling reads into contigs as above after excluding reads showing high sequence similarity to the phage genome. Obtained contigs were again binned as above, and the results were refined using Graphbin v1.2 [6]. We further detected and masked regions sharing identity with the phage genome by BLASTn searches with an e-value threshold of $10^{-4}$, using a script (https://github.com/tkhsgani/blast_masking.py). The resulting RsTa-C01 bin without the regions showing sequence similarities to the phage genome was designated “Contig-set 2”.

MinION reads after adapter trimming were filtered using Nanofilt v2.7.1 [7] with the “-q 13 -l 1000 --maxlength 20000 --headcrop 75” option, and chimeric reads were removed using yacrd v0.6.2 [8] with the “-c 4 -n 0.4 scrubb” option. Obtained reads ≥ 1 kb were mapped onto Contig-set 2 using Minimap2 v2.17 [9], and the mapped long reads were assembled into contigs using Flye v2.8 [10]. The resulting RsTa-C01 contig was circular, but certain regions were not covered with the long reads. To identify misassembled regions, we used Raven v1.6.0 [11] to create another assembly from the above mapped reads, and the results were compared using Nucmer v3.1 [12]. Regions showing discrepancy were replaced by contigs of the Raven assembly using a script (https://github.com/dkato2021/get-substituted-sequences), and we confirmed that the resulting circular contig was fully covered by the long reads. Furthermore, regions where discrepancy with Contig-set 1 was found were replaced using Nucmer v3.1 and the above script (https://github.com/dkato2021/get-substituted-sequences). Finally, the contig sequence was polished with the quality-trimmed MiSeq reads using Pilon v1.23 [13] with one iteration.

- CfP3-15

Assembly of MiSeq reads and binning were performed as in the genome of RsTa-C01. The MinION reads after adapter trimming were filtered using Nanofilt v2.7.1 with the "-q 13 -l 3000 --maxlength 20000 --headcrop 75" options, and palindromes introduced by whole-genome amplification were corrected with Pacasus (downloaded on 2020/09) [14] with the "--filter_factor=0.01 --query_coverage=0.01 --query_identity=0.01 --relative_score=0.01 --base_score=1.0" options. From the obtained reads ≥ 5 kb, chimeric reads were removed using yacrd v0.6.2 with the "-c 4 -n 0.4 scrubb" option. Obtained reads were assembled into contigs using Flye v2.8. From the contigs ≥ 10 kb, the CfP3-15 contigs were recovered by BLASTn searches using the above short read-derived CfP3-15 bin as queries. The resulting contigs were manually checked by BLASTn searches of the KEGG GENOME database and also by CAT v5.1.2 [15] annotation, and then sequences identified as non-*Clostridia* were excluded. The long reads were mapped to these CfP3-15 contigs using Minimap2 v2.17 and the mapped reads were recovered. This mapping revealed that this sample has a high degree of CfP3-15 genome-wide variation in read coverage, and therefore it was subsampled to read coverage of 200 using samNormalize.pl [16]. Subsampled reads were reassembled using Flye v2.8, and a circular CfP3-15 contig was obtained.

The quality-filtered MiSeq reads were mapped to this circular CfP3-15 contig, and mapped reads were recovered using Bowtie2 v2.4.1 [17]. The mapped reads were assembled using SPAdes v3.14.1 with the "--sc" and "--trusted-contigs" options, with the CfP3-15 circular contig being specified as the trusted contig. Contigs derived from this SPAdes assembly were used to replace regions of the circular CfP3-15 contig by using Nucmer v3.1 and the above script for substitution, where discrepancy was found. Finally, the contig was polished using Pilon v1.23 with one iteration.

- NkDv07

Assembly of MiSeq reads and binning were performed as in the genome of RsTa-C01, and the resulting NkDv07 bin was similarly designated "Contig-set 1". MinION reads were processed using Nanofilt v2.7.1 with "-q 7 -l 500 --maxlength 20000 --headcrop 75". The long reads were assembled into contigs using Flye v2.8, and NkDv07 contigs were recovered by BLASTn searches using Contig-set 1 as queries. Recovered contigs were manually checked by BLASTn searches of the KEGG GENOME database, and non-*Clostridia* sequences were excluded. The resulting contigs were cut at misassembled points using Tigmint v1.2.5 [18] with the "tigmint-make tigmint-long span=auto G=1.2e6 dist=auto" options and then scaffolded using LINKS v2.0.0 [19] with the "-d 1000, 2000, 3000, 4000" option, SLR [20] and LongStitch v1.0.1 [21]. The scaffolds were merged using CAMSA v1.3 [22], and gaps in the scaffolds were filled using TGS-GapCloser v1.1.1 [23]. The sequences were polished using Racon v1.4.16 [24] with two iterations and medaka v1.0.3 (https://github.com/nanoporetech/medaka). The resulting contigs were manually cut based on the results of mapping the long and short reads. Regions of the contigs were replaced by comparing them with Contig-set 1 using Nucmer v3.1 and the above script for substitution. The resulting contigs were then polished using Pilon v1.23 with three iterations with default settings and one additional iteration with the "--unpaired" option.

The resulting contig set contained a contig (~11 kb) that appeared to be chimera composed of several regions on other contigs, and indeed few MiSeq reads were mapped onto the contig. Therefore, the contig was discarded. Finally, regions covered with only a few reads were verified by PCR amplification and Sanger sequencing.

<u>Circular map of CfP3-15 and RsTa-C01</u>

Circular maps of the two complete genomes of the endosymbiotic *Clostridia* were depicted using Proksee (https://proksee.ca/). GC content and GC skew were calculated using a window size of 100,000 and a step size of 100. OriC and Ter sites were predicted using Ori-Finder 2022 [25].

<u>Detection of 16S rRNA gene in the MAG of GCA_90054335.1</u>

We were able to find 16S rRNA genes ≥ 500 bp only from two MAGs of the small genome clade, i.e., MGYG000003453 and GCA_900554335.1. Whereas the 16S rRNA gene of MGYG000003453 formed a monophyletic cluster with those of RsTa-C01, CfP3-15 and NkDv07 in consistence with the phylogenomic tree shown in Fig. 5A, the gene of GCA_900554335.1 did not. We suspected that the 16S rRNA gene of GCA_900554335.1 had possibly been misassembled in a previous study [26], and therefore, we reassembled the reads using SPAdes v3.14.1 with the "--meta" option, and resulting contigs were binned using MetaBat2 v2.12.1 [27]. BinSPreader [28] was used to refine the binning result, and the "--read" option was used to retrieve the read set of the bin corresponding to GCA_900554335.1. The read set was reassembled using SPAdes v3.14.1 with the "--only-assembler" option, and the 16S rRNA gene was detected in the obtained contigs using Barrnap v0.9 (https://github.com/tseemann/barrnap). The newly assembled and binned 16S rRNA gene formed a monophyletic cluster with the other four genes as shown in Fig. 2.

Re-binning of GCA_009785985.1

The MAG of GCA_009785985.1 was placed in the small genome clade by phylogenomic analysis, but its genome size was unusually large (ca. 2.7 Mbp) compared to the other MAGs in this clade. We suspected that the MAG probably contained contaminating contigs, and indeed, our CAT v.5.1.2-based contig classification showed that 239 of the 648 contigs were not assigned to the phylum *Bacillota*. We therefore re-binned the contigs using MyCC with the "56mer" option, and as a result, a 1.2-Mbp MAG consisting mostly of *Clostridia*-related contigs was obtained and used for the subsequent analyses.

Eukaryotic-like gene detection and signal peptide prediction

The proteins encoded by the small genome clade were used as queries of BLASTp searches of the NCBI non-redundant (nr) database from eukaryotes. The query proteins that showed significant sequence similarity with e-value of $10^{-4}$ were then used as queries of DIAMOND BLASTp v2.0.13 [29] searches of the NCBI nr database with the "--masking seg --ultra-sensitive" options. The protein sequences from members of the small genome clade present in the NCBI database (i.e., GCA_000435275.1, GCA_009785985.1, GCA_017406805.1, GCA_017414725.1, GCA_017431755.1, GCA_017432165.1, GCA_017448285.1, GCA_017480265.1, GCA_017515685.1, GCA_017634155.1) were ignored. When protein sequences showed the highest similarity to those from eukaryotes or when 90% of the detected database sequences were from eukaryotes, they were selected as candidates of eukaryotic-like proteins. These candidates were then used as queries for NCBI Conserved Domain Search (CD-search) in the web server (2022/09/19), and detected domains were selected as candidates of eukaryotic-like domains. Based on the NCBI and InterPro databases, the eukaryotic-like domain candidates were manually checked, and domains that are common also in bacteria and/or archaea were removed. The remaining domains were regarded as eukaryotic-like domains. Finally, local CD-search was performed using RPS-BLAST of Conserved Domain Database v3.20 for all proteins encoded by the small genome clade, and the proteins showing the highest similarity to eukaryotic-like domains were designated as eukaryotic-like proteins. Local CD-search was also performed for other genomes of "Acutalibacteraceae", and eukaryotic-like proteins were selected in the same way based on the eukaryotic-like domains specified above.

For the identified eukaryotic-like genes of CfP3-15, NkDv07, and RsTa-C01, signal peptides were predicted using SignalP 6.0 [30].

Phylogenetic analysis including short 16S rRNA gene amplicon sequences

The 16S rRNA gene amplicon sequences obtained by Levy and Jami (2018) [31] and Utami et al. (2018) [32] were placed onto the maximum likelihood tree shown in Fig. S2B, using EPA-NG v0.3.8 [33]. EPA-NG performs phylogenetic placement based on maximum likelihood, and each sequence can be assigned to multiple places of the tree with different likelihoods. We obtained a tree, where each amplicon sequence was represented on the position with the highest likelihood on the tree, using GAPPA v.0.8.3 [34] with the "graft --fully-resolve" option.

Tree visualization

Phylogenetic trees were visualized using the ggtree R package [35].

Reconstruction of other MAGs and statistics

The quality-trimmed reads of each sample were assembled using SPAdes v3.14.1 with the "--sc" option and binned using MyCC with the default mode. The completeness of reconstructed bins was estimated using CheckM [36] and also CheckM2 [37]. The MAGs with CheckM or CheckM2 completeness >50% were taxonomically classified using GTDB-tk v2.1.0 [38] with the r207 database. Reads alignment rates for each MAG were calculated using Bowtie2 v2.4.1.

#### Supplementary Results

##### 16S rRNA gene cloning analysis of bacterial microbiota associated with single protist cells

During the sequencing analysis of 16S rRNA genes amplified by PCR from a suspension of ten *Trichonympha agilis* cells in our previous study [39], six identical sequences among 46 clones were assigned to the class *Clostridia* and were designated as phylotype RsTa-C01. The remaining clones comprised 16 sequences of the dominant obligate endosymbiont "*Candidatus* Endomicrobium trichonymphae" which supplements nitrogenous compounds [40-42], 20 sequences of the dominant hydrogen-oxidizing obligate ectosymbiont "*Candidatus* Desulfovibrio trichonymphae" [39, 43], and four minor, facultative *Mycoplasmatales* symbionts [44].

16S rRNA phylotype CfP3-15 was identified during preliminary sequencing analyses of 16S rRNA genes amplified by PCR from single cells of *Pseudotrichonympha grassii* for the previous genome analysis of "*Candidatus* Azobacteroides trichonymphae" [45]. In a single-cell WGA sample of *P. grassii*, five of 93 clones were identical sequences belonging to the class *Clostridia* and were designated as phylotype CfP3-15. The remaining clones comprised the predominant, dinitrogen-fixing obligate endosymbiont "*Ca.* Azobacteroides trichonymphae" [45-47] and a clone related to "*Candidatus* Ancillula" [48, 49].

16S rRNA phylotype NkDv07 was identified during preliminary sequencing analyses of 16S rRNA genes amplified by PCR from single cells of the largest morphotype among multiple *Devescovina* species in the gut of *Neotermes sugioi* [50] for our unpublished genome analysis of "*Candidatus* Armantifilum devescovinae". Three identical sequences among 95 clones were assigned to the class *Clostridia* and were designated as phylotype NkDv07. The remaining clones comprised the predominant, dinitrogen-fixing ectosymbiont "*Candidatus* Armantifilum devescovinae" [50-53], 14 clones related to "*Candidatus* Nucleococcus" (phylum *Verrucomicrobiota*) [54], and three clones of the order *Holosporales.*

##### Other MAGs detected in WGA samples

During assembling and binning, several other MAGs in addition to the clostridial ones were recovered and listed in Table S8. Most of those MAGs were assigned to either of "*Ca*. Azobacteroides", *Endomicrobium* or *Spirochaetales* (*Termitinemataceae* and *Treponemataceae*). Among them, "*Ca.* Azobacteroides pseudotrichonymphae" and "*Ca.* Endomicrobium trichonymphae" are previously known mutualistic, permanently-associated endosymbionts of *Pseudotrichonympha grassii* and *Trichonympha agilis*, respectively [42, 45]. Any MAGs were not shared by different WGA samples, indicating these MAGs were not contaminants in reagents. Members of *Spirochaetales* have frequently been detected as ectosymbionts of termite gut protists [55-57].

**The functions of hypothetical proteins OG0003398, OG0004343, and OG0005234**

We attempted to predict the functions of the hypothetical proteins for which the gene flux analysis showed that the last common ancestor of the small genome clade gained. The functional prediction was conducted based on sequence homology using the NCBI Conserved Domain Search and structural homology using AlphaFold2 [58] and Dali server [59]. However, the NCBI Conserved Domain Search did not show significantly homologous domains, and AlphaFold2 also did not predict the structure with a high confidence score. Therefore, the functions of the hypothetical proteins were currently unknown.

**Description of '*Candidatus* Improbicoccus' gen. nov**

*Improbicoccus* [L. masc. adj. *improbus*, disloyal, wicked; M.L. m. n. *coccus*, from Gr. a berry, coccus; N.L. m. n. *Improbicoccus*, a parasitic coccus]. The bacteria specifically colonize the cytoplasm of protist cells in the termite guts. They are cocci of 1–2 μm and have a Gram-positive type cell wall. They are obligate intracellular parasites and have not been cultured. The type species is '*Candidatus* Improbicoccus pseudotrichonymphae', corresponding to CfP3-15, with its genome sequence as the type material.

**Description of '*Candidatus* Paraimprobicoccus' gen. nov**

*Paraimprobicoccus* [Gr. prep. para, alike, alongside of; N.L. m. n. *Improbicoccus*, taxonomic name of a bacterial genus; N.L. m. n. *Improbicoccus*, taxonomic name of a bacterial *Improbicoccus*-alike genus]. The bacteria have cell structure and metabolic capacity similar to *Candidatus* Improbicoccus. ANI and AAI between this genus and *Candidatus* Improbicoccus are below 70% and 50%, respectively. The type species is '*Candidatus* Paraimprobicoccus trichonymphae', corresponding to RsTa-C01, with its genome sequence as the type material.

**Description of '*Candidatus* Improbicoccus pseudotrichonymphae' sp. nov**

*Improbicoccus pseudotrichonymphae* (pseudo. tri.cho.nym'phae. N.L. n. *Pseudotrichonympha*, a genus of flagellated protists; N.L. gen. pseudotrichonymphae, of *Pseudotrichonympha*, referring to the host genus). The fundamental attributes of this species are the same as those for the genus. The bacterium colonizes the cytoplasm of the protist *P. grassii* in the gut of the termite *Coptotermes formosanus*. The cell dimensions are 0.5–1.0 μm by 0.5–1.0 μm. The assignment is based on the 16S rRNA gene, the complete genome (SAMD00586941), and hybridization with the 16S rRNA-targeted probe CfP3-15-656 (5′-CCGTTCGCCTCTACTTTAC-3′). This species corresponds to CfP3-15.

**Description of '*Candidatus* Improbicoccus devescovinae' sp. nov**

*Improbicoccus devescovinae* (de.ves.co.vi'nae. N.L. fem. n. *Devescovina*, a genus of flagellated protists; N.L. gen. fem. n. devescovinae, of *Devescovina*, referring to the host genus). The fundamental attributes of this species are the same as those for the genus. The bacterium colonizes the cytoplasm of the protist *Devescovina* sp. in the gut of the termite *Neotermes sugioi*. The cell dimensions are 1.0–2.0 μm by 1.0–2.0 μm and the cells usually form an aggregate. The assignment is based on the 16S rRNA gene, the draft genome (SAMD00586942), and hybridization with the 16S rRNA-targeted probe NkDv07-142 (5′-TCCATCAGCTATCTCCCAC-3′). This species corresponds to NkDv07.

**Description of '*Candidatus* Paraimprobicoccus trichonymphae' sp. nov**

*Paraimprobicoccus trichonymphae* (tri.cho.nym'phae. N.L. fem. n. *Trichonympha*, a genus of flagellated protists; N.L. gen. fem. n. Trichonymphae, of *Trichonympha*, referring to the host genus). The fundamental attributes of this species are the same as those for the genus. The bacterium colonizes the cytoplasm of the protist *Trichonympha agilis* in the gut of termite *Reticulitermes speratus*. The cell dimensions are 1.0–2.0 μm by 1.0–2.0 μm and form diplococci. The assignment is based on the 16S rRNA gene, the complete genome (SAMD00586943), and hybridization with the 16S rRNA-targeted probe RsTa-C01-76 (5′-GTGCCTTGCAAAACTCCG-3′). This species corresponds to RsTa-C01.

**Legends to Supplementary Figures**

**Figure S1.** Fluorescence in situ hybridization analysis of 16S rRNA phylotype NkDv07 (**A**, **B**) and transmission electron micrographs of intracellular bacterial symbionts of *Pseudotrichonympha grassii* (**C**–**E**). Phase contrast image of *Devescovina* sp. from the gut of *Neotermes sugioi* (**A**) and NkDv07 cells detected using 6FAM-labelled probe NkDv07-142 (green) as single cocci (**B**). Amorphous yellowish autofluorescence in **B** was emitted from wood particles ingested by *Devescovina* sp. Putative clostridial cell (C), and "*Candidatus* Azobacteroides pseudotrichonymphae" cells (D). In panel **E**, a putative clostridial cell apparently during fission was in the center and a "*Ca.* A. pseudotrichonymphae" cell in the lower right. Bars: **A** and **B**, 10 µm; **C**, 100 nm; **D**, 200 nm; **E**, 500 nm.

**Figure S2.** Phylogenetic positions of endosymbiotic *Clostridia* based on 16S rRNA gene sequences. (**A**) Maximum-likelihood tree of endosymbiotic *Clostridia* and other members of the family *Oscillospiraceae*. Described species of *Oscillospiraceae* were included in the tree. The tree was constructed with the GTR+I+G4 nucleotide substitution model and rooted with *Syntrophomonas wolfei* (CP000448) and *Thermosyntropha lipolytica* (FQWY01000001) as the outgroup. Clusters of *Clostridium* [60] and the clade corresponding to "Acutalibacteraceae" in the GTDB taxonomy are highlighted in gray and red, respectively. (**B**) Maximum-likelihood tree of endosymbiotic *Clostridia* and other members of the "Acutalibacteraceae". Uncultured clones closely related to described species and endosymbiotic *Clostridia* were included. The tree was constructed with the GTR+I+G4 nucleotide substitution model and rooted with *Sporobacter termitidis* (Z49863) and *Oscillibacter ruminantium* (JF750939) as the outgroup. Highly supported nodes (SH-aLRT value ≥ 80% and ultrafast bootstrap support value ≥ 95%) are highlighted with a closed circle.

**Figure S3.** Relationships between the size and GC content of the complete genomes in the class *Clostridia*.

**Figure S4.** Circular representation of the CfP3-15 chromosome (**A**) and the RsTa-C01 chromosome (**B**). The rings denote the following features (from inside): (i) Scale in mega base pairs of the chromosome (black); (ii) GC content (black); (iii) GC skew (green and purple); (iv) predicted genes present on the forward strand (+); (v) genes present on the reverse strand (-). OriC and Ter sites are indicated by arrows in **A**. For the RsTa-C01 genome, we could not detect these sites, using Ori-Finder 2022.

**Figure S5.** Comparison of the number (**A**) and ratio (**B**) of genes classified into clusters of orthologous genes (COG) functional categories. The genomes of CfP3-15, NkDv07, RsTa-C01, and their free-living relatives *Clostridium leptum* and *Ruminococcus bromii* were compared. The categories denote the following functions: (A) RNA processing and modification; (B) chromatin structure and dynamics; (C) energy production and conversion; (D) cell cycle control, cell division, chromosome partitioning; (E) amino acid transport and metabolism; (F) nucleotide transport and metabolism; (G) carbohydrate transport and metabolism; (H) coenzyme transport and metabolism; (I) lipid transport and metabolism; (J) translation, ribosomal structure and biogenesis; (K) transcription; (L) replication, recombination and repair; (M) cell wall/membrane/envelope biogenesis; (N) cell motility; (O) posttranslational modification, protein turnover, chaperones; (P) inorganic ion transport and metabolism; (Q) secondary metabolites biosynthesis, transport and catabolism; (R) general function prediction only; (S) function unknown; (T) signal transduction mechanisms; (U) intracellular trafficking, secretion, and vesicular transport; (V) defense mechanisms; (W) extracellular structures; (X) mobilome: prophages, transposons; (Y) nuclear structure; (Z) cytoskeleton.

**Figure S6.** Comparison of the number (**A**) and ratio (**B**) of pseudogenes classified into clusters of orthologous genes (COG) functional categories. See also the legend to Fig. S5.

**Figure S7.** Amino acid sequence alignment of ATP/ADP translocase and other nucleotide transport proteins from the endosymbiotic *Clostridia*, other parasitic bacteria, and plastids. Conserved residues important for substrate specificity, transport efficiency, and counter exchange of ATP/ADP translocase (CAA89201.2) of the plastid of *Arabidopsis thaliana* are indicated with arrows.

**Figure S8.** RT-PCR analysis of the ATP/ADP translocase gene. Agarose gel electrophoresis of RT-PCR products from total RNA of *R. speratus* guts (**A**) and *C. formosanus* guts (**B**). N, negative control (RNA samples without reverse transcription); P, products of RT-PCR using primer sets specific to the respective ATP/ADP translocase genes (see Table S2). These bands were excised from gels, and their sequences were verified by cloning and Sanger sequencing analysis.

**Figure S9.** Phylogenetic positions of eukaryotic-like genes contained in the genomes of CfP3-15, NkDv07, and RsTa-C01. The following genes were used as queries to identify homologues as reference sequences. (**A**) CfP3-15_0049. (**B**) CfP3-15_0165. (**C**) CfP3-15_0235. (**D**) CfP3-15_0282. (**E**) CfP3-15_0445. (**F**) CfP3-15_0456. (**G**) CfP3-15_0548. (**H**) CfP3-15_0552. (**I**) CfP3-15_0565. (**J**) CfP3-15_0581. (**K**) CfP3-15_0591. (**L**) CfP3-15_0627. (**M**) CfP3-15_0653. (**N**) CfP3-15_0780. (**O**) CfP3-15_0790. (**P**) CfP3-15_0852. (**Q**) CfP3-15_0856. (**R**) CfP3-15_0934. (**S**) NkDv07_0114. (**T**) NkDv07_0121. (**U**) NkDv07_0512. (**V**) NkDv07_0536. (**W**) NkDv07_0615. (**X**) RsTa-C01_0144. (**Y**) RsTa-C01_0199. (**Z**) RsTa-C01_0208. (**AA**) RsTa-C01_0431. (**AB**) RsTa-C01_0774. (**AC**) RsTa-C01_0775. (**AD**) RsTa-C01_0992. Highly supported nodes (SH-aLRT value ≥ 80% and ultrafast bootstrap support value ≥ 95%) are highlighted with closed circles. The sequence IDs highlighted by green, blue, and light blue correspond to the proteins of the endosymbiotic *Clostridia* used to find homologs, proteins of the protist hosts, and proteins from parabasalid protists, respectively.

**Figure S10.** RT-PCR analysis of eukaryotic-like protein-coding genes. Agarose gel electrophoresis of RT-PCR products from total RNA of *R. speratus* guts (**A**) and *C. formosanus* guts (**B**). N, negative control (RNA samples without reverse transcription); P, products of RT-PCR using primer sets specific to the respective eukaryotic-like protein-coding genes (see Table S2). The amplification products of eukaryotic-like protein-coding genes are indicated by arrows, which were confirmed by Sanger sequencing.

**Figure S11.** Comparison of a eukaryotic-like gene, CfP3-15_0552, in the CfP3-15 genome and that obtained by RT-PCR from *C. formosanus* guts. (**A**) Nucleotide sequences. (**B**) Amino acid sequences of **A**.

**Figure S12.** Pairwise ANI (**A**) and AAI (**B**) heatmap among the genomes in the small genome clade. The phylogenetic tree of the small genome clade is shown. In **B**, the genus-level clade based on GTDB r207 are indicated.

**Figure S13.** Principal component analysis of genomes of "Acutalibacteraceae" (Table S3) based on the relative abundance of the COG functional categories (**A**) and comparison of the number (**B**) and ratio (**C**) of the COG functional categories. Genomes belonging to the small genome clade are highlighted with red. Arrows indicate the loadings of each functional category. Several arrows are omitted to improve visibility. See also the legend to Fig. S5.

**Figure S14.** Distribution of genes acquired by the last common ancestor of the small genome clade, predicted in the gene flux analysis. The phylogenetic tree corresponds to that shown in Fig. 5A with the small genome clade being shaded in red.

**Figure S15.** Comparison between the phylogenomic tree of the small genome clade and the gene tree of their ATP/ADP translocase. (**A**) Comparison of the *Clostridia* ATP/ADP translocase I (left) with the phylogenomic tree (right). (**B**) Comparison of the *Clostridia* ATP/ADP translocase II (left) and the phylogenomic tree (right). Highly supported nodes (SH-aLRT value ≥ 80% and ultrafast bootstrap support value ≥ 95%) are indicated with a closed circle.

**Figure S16.** Phylogenetic positions of 16S rRNA gene amplicon sequences derived from the protistan fraction of a cattle rumen [31] and the guts of termites and cockroaches [32].

**References to Supplementary Materials**


1. Zhang K, Martiny AC, Reppas NB, Barry KW, Malek J, Chisholm SW, et al. Sequencing genomes from single cells by polymerase cloning. Nat Biotechnol. 2006;24:680-6.
2. Martin M. Cutadapt removes adapter sequences from high-throughput sequencing reads. EMBnet J. 2011;17:10.
3. Schmieder R, Edwards R. Quality control and preprocessing of metagenomic datasets. Bioinformatics. 2011;27:863-4.
4. Bankevich A, Nurk S, Antipov D, Gurevich AA, Dvorkin M, Kulikov AS, et al. SPAdes: a new genome assembly algorithm and its applications to single-cell sequencing. J Comput Biol. 2012;19:455-77.
5. Lin HH, Liao YC. Accurate binning of metagenomic contigs via automated clustering sequences using information of genomic signatures and marker genes. Sci Rep. 2016;6:e24175.
6. Mallawaarachchi V, Wickramarachchi A, Lin Y. GraphBin: refined binning of metagenomic contigs using assembly graphs. Bioinformatics. 2020;36:3307-13.
7. De Coster W, D'Hert S, Schultz DT, Cruts M, Van Broeckhoven C. NanoPack: visualizing and processing long-read sequencing data. Bioinformatics. 2018;34:2666-9.
8. Marijon P, Chikhi R, Varre JS. yacrd and fpa: upstream tools for long-read genome assembly. Bioinformatics. 2020;36:3894-6.
9. Li H. Minimap2: pairwise alignment for nucleotide sequences. Bioinformatics. 2018;34:3094-100.
10. Kolmogorov M, Yuan J, Lin Y, Pevzner PA. Assembly of long, error-prone reads using repeat graphs. Nat Biotechnol. 2019;37:540-6.
11. Vaser R, Šikić M. Time- and memory-efficient genome assembly with Raven. Nat Comput Sci. 2021;1:332-6.
12. Kurtz S, Phillippy A, Delcher AL, Smoot M, Shumway M, Antonescu C, et al. Versatile and open software for comparing large genomes. Genome Biol. 2004;5:R12.
13. Walker BJ, Abeel T, Shea T, Priest M, Abouelliel A, Sakthikumar S, et al. Pilon: an integrated tool for comprehensive microbial variant detection and genome assembly improvement. PLoS One. 2014;9:e112963.
14. Warris S, Schijlen E, van de Geest H, Vegesna R, Hesselink T, Te Lintel Hekkert

B, et al. Correcting palindromes in long reads after whole-genome amplification. BMC Genom. 2018;19:e798.
15. Von Meijenfeldt FAB, Arkhipova K, Cambuy DD, Coutinho FH, Dutilh BE. Robust taxonomic classification of uncharted microbial sequences and bins with CAT and BAT. Genome Biol. 2019;20:e217.
16. Eccles D. gringer/bioinfscripts: Tree Lab / Global River Release (1.3.0-TRL). 2019.
17. Langmead B, Salzberg SL. Fast gapped-read alignment with Bowtie 2. Nat Methods. 2012;9:357-9.
18. Jackman SD, Coombe L, Chu J, Warren RL, Vandervalk BP, Yeo S, et al. Tigmint: correcting assembly errors using linked reads from large molecules. BMC Bioinform. 2018;19:e393.
19. Warren RL, Yang C, Vandervalk BP, Behsaz B, Lagman A, Jones SJ, et al. LINKS: Scalable, alignment-free scaffolding of draft genomes with long reads. Gigascience. 2015;4:e35.
20. Luo J, Lyu M, Chen R, Zhang X, Luo H, Yan C. SLR: a scaffolding algorithm based on long reads and contig classification. BMC Bioinform. 2019;20:e539.
21. Coombe L, Li JX, Lo T, Wong J, Nikolic V, Warren RL, et al. LongStitch: high-quality genome assembly correction and scaffolding using long reads. BMC Bioinform. 2021;22:e534.
22. Aganezov SS, Alekseyev MA. CAMSA: a tool for comparative analysis and merging of scaffold assemblies. BMC Bioinform. 2017;18:e496.
23. Xu M, Guo L, Gu S, Wang O, Zhang R, Peters BA, et al. TGS-GapCloser: A fast and accurate gap closer for large genomes with low coverage of error-prone long reads. Gigascience. 2020;9:giaa094.
24. Vaser R, Sovic I, Nagarajan N, Sikic M. Fast and accurate de novo genome assembly from long uncorrected reads. Genome Res. 2017;27:737-46.
25. Dong MJ, Luo H, Gao F. Ori-Finder 2022: A comprehensive web server for prediction and analysis of bacterial replication origins. Genomics Proteomics Bioinformatics. 2022;20:1207-13.
26. Almeida A, Mitchell AL, Boland M, Forster SC, Gloor GB, Tarkowska A, et al. A new genomic blueprint of the human gut microbiota. Nature. 2019;568:499-504.
27. Kang DD, Li F, Kirton E, Thomas A, Egan R, An H, et al. MetaBAT 2: an adaptive binning algorithm for robust and efficient genome reconstruction from metagenome assemblies. PeerJ. 2019;7:e7359.
28. Tolstoganov I, Kamenev Y, Kruglikov R, Ochkalova S, Korobeynikov A.

BinSPreader: Refine binning results for fuller MAG reconstruction. iScience. 2022;25:e104770.
29. Buchfink B, Reuter K, Drost HG. Sensitive protein alignments at tree-of-life scale using DIAMOND. Nat Methods. 2021;18:366-8.
30. Teufel F, Almagro Armenteros JJ, Johansen AR, Gislason MH, Pihl SI, Tsirigos KD, et al. SignalP 6.0 predicts all five types of signal peptides using protein language models. Nat Biotechnol. 2022;40:1023-5.
31. Levy B, Jami E. Exploring the prokaryotic community associated with the rumen ciliate protozoa population. Front Microbiol. 2018;9:e2526.
32. Utami YD, Kuwahara H, Murakami T, Morikawa T, Sugaya K, Kihara K, et al. Phylogenetic diversity and single-cell genome analysis of "*Melainabacteria*", a non-photosynthetic cyanobacterial group, in the termite gut. Microbes Environ. 2018;33:50-7.
33. Barbera P, Kozlov AM, Czech L, Morel B, Darriba D, Flouri T, et al. EPA-ng: Massively parallel evolutionary placement of genetic sequences. Syst Biol. 2019;68:365-9.
34. Czech L, Barbera P, Stamatakis A. Genesis and Gappa: processing, analyzing and visualizing phylogenetic (placement) data. Bioinformatics. 2020;36:3263-5.
35. Yu G, Smith DK, Zhu H, Guan Y, Lam TTY, McInerny G. ggtree: an R package for visualization and annotation of phylogenetic trees with their covariates and other associated data. Methods Ecol Evol. 2016;8:28-36.
36. Parks DH, Imelfort M, Skennerton CT, Hugenholtz P, Tyson GW. CheckM: assessing the quality of microbial genomes recovered from isolates, single cells, and metagenomes. Genome Res. 2015;25:1043-55.
37. Chklovski A, Parks DH, Woodcroft BJ, Tyson GW. CheckM2: a rapid, scalable and accurate tool for assessing microbial genome quality using machine learning. bioRxiv. 2022.
38. Chaumeil PA, Mussig AJ, Hugenholtz P, Parks DH. GTDB-Tk: a toolkit to classify genomes with the Genome Taxonomy Database. Bioinformatics. 2019;36:1925-7.
39. Sato T, Hongoh Y, Noda S, Hattori S, Ui S, Ohkuma M. *Candidatus* Desulfovibrio trichonymphae, a novel intracellular symbiont of the flagellate *Trichonympha agilis* in termite gut. Environ Microbiol. 2009;11:1007-15.
40. Stingl U, Radek R, Yang H, Brune A. "*Endomicrobia*": Cytoplasmic symbionts of termite gut protozoa form a separate phylum of prokaryotes. Appl Environ Microbiol. 2005;71:1473-9.

### Supplementary Tables

**Table S1: 16S rRNA genes used in phylogenetic analysis (see Fig. 2 and S2).**

| accession numbers in SILVA or LTP used in Fig. 2 | |
|---|---|
| AB192174 | KP090466 |
| AB299544 | KP690938 |
| AB555038 | LN612663 |
| AB612693 | X66002 |
| AB614861 | MH043116 |
| AB614876 | MN851263 |
| AB615006 | MT875034 |
| AB746684 | FJ805840 |
| AB746749 | AJ305238 |
| AB821861 | KR364749 |
| AF371774 | L76600 |
| DQ795962 | MN872474 |
| DQ796114 | MK761171 |
| DQ796268 | LC589980 |
| DQ796766 | MT580120 |
| DQ796799 | MT905216 |
| EF445283 | NR_144727 |
| EU461213 | NR_144736 |
| EU463585 | LT985454 |
| EU778659 | LT897838 |
| EU843933 | ON921083 |
| EU845152 | LT732642 |
| FJ366876 | LT960605 |
| GQ134270 | Z49863* |
| GQ502578 | JF750939* |
| GQ502583 | |
| GQ898015 | |
| GU174104 | |
| HM630202 | |
| HM630229 | |
| HM630241 | |
| JQ184761 | |
| JQ184883 | |
| JQ184923 | |
| JQ185143 | |
| JQ185474 | |
| JQ191019 | |
| JQ617823 | |
| JQ993514 | |
| JQ993520 | |
| JQ993637 | |
| JQ993641 | |
| JQ993642 | |
| KC163071 | |
| KF318227 | |
| KF841696 | |
| KJ399537 | |
| KM251066 | |

| accession numbers in SILVA, LTP, or type strain genome database used in Fig. S2A | | | |
|---|---|---|---|
| EF529620 | EU474489 | AB011057 | JAOQKF000000000 |
| FR749897 | DQ015339 | ACXX02000011 | JAOQKG000000000 |
| X66002 | HM328208 | X71856 | GCM10018885 |
| MH043116 | KM650259 | MK935571 | LN866995 |
| MN851263 | KM650490 | AB267266 | NR_147398 |
| MT875034 | EU459548 | FR749966 | NR_179526 |
| FJ805840 | CCFG01000002 | FJ815191 | NR_147370 |
| AJ305238 | HQ769937 | AB793710 | NR_144722 |
| KR364749 | DQ794293 | HE862234 | NR_144727 |
| L76600 | DQ394630 | AEDB02000059 | NR_144736 |
| MN872474 | DQ796766 | X71846 | LN881596 |
| MK761171 | DQ796799 | AY959944 | LT631513 |
| LC589980 | KT337631 | AB186359 | LT576393 |
| L09177 | KT337629 | CP000568 | LT598578 |
| AY295777 | JQ993514 | AB125279 | LN870316 |
| AY487928 | JQ993641 | MK138666 | LT558850 |
| ACEC01000059 | EU828413 | LN868252 | LT725660 |
| L76596 | EU828406 | HG003571 | KY777734 |
| AB910742 | EU843382 | AJ575187 | LT985392 |
| AM915269 | EU463443 | AB910750 | LT934440 |
| CP002403 | EF096602 | GQ900631 | Y18180 |
| MH200617 | EF097212 | AJ310082 | LT985454 |
| AJ315980 | EF097423 | AB221372 | LT897838 |
| EU158190 | DQ796325 | KT630605 | GCM10018801 |
| LT576408 | EF097263 | LC522508 | GCM10018815 |
| LT854295 | EU469153 | LC522510 | OL617409 |
| KU999999 | HQ780956 | LC522513 | ON921083 |
| X98011 | KT337628 | MN055917 | LT732642 |
| X97852 | EU887962 | MN055919 | LT960605 |
| KM098109 | EU887966 | KX146426 | CP000448* |
| LN846908 | EU887963 | MT580120 | FQWY01000001* |
| AJ518869 | EU887967 | MZ310595 | |
| AJ413954 | DQ796114 | AJ270469 | |
| KX150462 | DQ796268 | DQ057476 | |
| ABCA03000037 | HQ789326 | LC670256 | |
| LC371917 | JQ993637 | MT905130 | |
| AY136666 | GQ132241 | MT905132 | |
| AGCK01000014 | EU843933 | MT905147 | |
| X81125 | KX672314 | MT905148 | |
| JF750939 | KX672322 | MT905149 | |
| AP012044 | KM251066 | MT905206 | |
| MG963288 | KM251154 | MT905207 | |
| AF167711 | DQ015324 | MT905208 | |
| Z49863 | JQ617823 | MT905216 | |
| KP889099 | JQ625023 | MT905210 | |
| L34618 | EU461213 | JAOQJZ000000000 | |
| KY978733 | HM218859 | JAOQKB000000000 | |
| CP001348 | DQ394629 | JAOQJE000000000 | |

| accession numbers in SILVA or LTP used in Fig. S2B | | | |
|---|---|---|---|
| AB034072 | EF686627 | KX672258 | JQ993642 |
| AB185796 | EU622658 | KX672556 | KC163071 |
| AB506382 | EU622736 | LAQZ01000049 | KF318227 |
| AB606299 | EU828413 | LC036223 | KF841696 |

| | | | |
|---|---|---|---|
| AB622826 | FJ880466 | LC036224 | KJ399537 |
| AB626936 | FPLM01005507 | LC194924 | KM251066 |
| AB627566 | FPLO01002175 | AB192174 | KP090466 |
| AB627625 | GQ402108 | AB299544 | KP690938 |
| AB627689 | GQ451304 | AB555038 | LN612663 |
| AB627717 | GQ999966 | AB612693 | X66002 |
| AB700411 | GU907824 | AB614861 | MH043116 |
| AB746644 | GU907825 | AB614876 | MN851263 |
| AB746645 | HM124146 | AB615006 | MT875034 |
| AB746650 | HM124266 | AB746684 | FJ805840 |
| AB746713 | HQ400121 | AB746749 | AJ305238 |
| AB746717 | HQ400259 | AB821861 | KR364749 |
| AB746718 | HQ746235 | AF371774 | L76600 |
| AB746727 | HQ780956 | DQ795962 | MN872474 |
| AB746748 | HQ789326 | DQ796114 | MK761171 |
| AB746764 | HQ805976 | DQ796268 | LC589980 |
| AB818589 | HQ806126 | DQ796766 | MT580120 |
| AB818596 | HQ810098 | DQ796799 | MT905216 |
| AB821695 | HQ814750 | EF445283 | NR_144727 |
| AB821707 | JF766393 | EU461213 | NR_144736 |
| AB824445 | JN653041 | EU463585 | LT985454 |
| AB930590 | JQ084431 | EU778659 | LT897838 |
| AB969538 | JQ085232 | EU843933 | ON921083 |
| AF332709 | JQ248107 | EU845152 | LT732642 |
| AF371771 | JQ607884 | FJ366876 | LT960605 |
| AJ295664 | JQ799169 | GQ134270 | Z49863 * |
| AM278898 | JQ996700 | GQ502578 | JF750939 * |
| AM697405 | JX575812 | GQ502583 | |
| AY949858 | JX575926 | GQ898015 | |
| AY949859 | KC163015 | GU174104 | |
| BCAA01000057 | KC163094 | HM630202 | |
| CCFG01000002 | KC555197 | HM630229 | |
| DQ394691 | KJ734862 | HM630241 | |
| DQ777904 | KM244841 | JQ184761 | |
| DQ905330 | KM244901 | JQ184883 | |
| DQ905407 | KM250930 | JQ184923 | |
| DQ905581 | KM454168 | JQ185143 | |
| DQ905824 | KM978279 | JQ185474 | |
| EF071387 | KT337628 | JQ191019 | |
| EF400741 | KT337630 | JQ617823 | |
| EF404262 | KT337631 | JQ993514 | |
| EF405002 | KT935669 | JQ993520 | |
| EF405280 | KT952843 | JQ993637 | |
| EF686593 | KX672228 | JQ993641 | |

* outgroup sequence

**Table S2: PCR primers designed in this study.**

| name | sequence (5' – 3') | target |
|---|---|---|
| RsTa-C01_Fwd | tagccggactgagaggttg | 16S rRNA gene of RsTa-C01 |
| RsTa-C01_Rev | catctcacgacacgagctg | 16S rRNA gene of RsTa-C01 |
| CfP3-15_Fwd | tgagtaaagtagaggcgaacg | 16S rRNA gene of CfP3-15 |
| CfP3-15_Rev | aggccatctttgagggattag | 16S rRNA gene of CfP3-15 |
| NkDv07_Fwd | aacacatgcaagtcgaacg | 16S rRNA gene of NkDv07 |
| NkDv07_Rev | ccttgcggttaggctatcg | 16S rRNA gene of NkDv07 |
| RsTa-C01_tlcA_189F | aatacgctgtgatgggagcc | *tlcA* of RsTa-C01 |
| RsTa-C01_tlcA_493R | gatgagactgcaatcgtacc | *tlcA* of RsTa-C01 |
| CfP3-15_tlcA-1_862F | atgcttccggtgtgttgtgg | *tlcA* (*Clostridia* ATP/ADP translocase 1) of CfP3-15 |
| CfP3-15_tlcA-1_1227R | acaatacgttgcccctttgc | *tlcA* (*Clostridia* ATP/ADP translocase 1) of CfP3-15 |
| CfP3-15_tlcA-2_558F | tccgttggtagttgccttgg | *tlcA* (*Clostridia* ATP/ADP translocase 2) of CfP3-15 |
| CfP3-15_tlcA-2_933R | ttcaacgcccgaatacctcg | *tlcA* (*Clostridia* ATP/ADP translocase 2) of CfP3-15 |
| CfP3-15_0049_210F | agtcagtgaaggcgaagacg | CfP3-15_0049 |
| CfP3-15_0049_569R | tctgcaccaacagtcggaac | CfP3-15_0049 |
| CfP3-15_0165_156F | tgtgggtgacgttcatggag | CfP3-15_0165 |
| CfP3-15_0165_474R | atgaatttccgcagcagcac | CfP3-15_0165 |
| CfP3-15_0552_289F | gagggtttctgccgatatgc | CfP3-15_0552 |
| CfP3-15_0552_541R | ttgcgtaatggccgtcatc | CfP3-15_0552 |
| CfP3-15_0627_105F | tgcattgttggcagtttcgg | CfP3-15_0627 |
| CfP3-15_0627_411R | gggtatagaaagcaacgccg | CfP3-15_0627 |
| CfP3-15_0790_107F | tcggtgatgaaggatcggg | CfP3-15_0790 |
| CfP3-15_0790_479R | acttcgtcccttagagtcgc | CfP3-15_0790 |
| CfP3-15_0852_325F | gttggtggtttgggtttggc | CfP3-15_0852 |
| CfP3-15_0852_648R | ttcttgtcccgtggtactcc | CfP3-15_0852 |
| RsTa-C01_0144_48F | gtttgcacaagtttcggctc | RsTa-C01_0144 |
| RsTa-C01_0144_392R | tcaggtgcaatttcacaagaagc | RsTa-C01_0144 |
| RsTa-C01_0431_85F | atgagcggtaacaagccaaatg | RsTa-C01_0431 |

| | | |
|---|---|---|
| RsTa-C01_0431_453R | ttccaagaaccaggctaaac | RsTa-C01_0431 |
| RsTa-C01_0774_124F | attgagacacatggcactgg | RsTa-C01_0774 |
| RsTa-C01_0774_468R | ttggtattcgtagcaaaggg | RsTa-C01_0774 |
| RsTa-C01_0844_930F | acaagtgttgccgcaaagtc | RsTa-C01_0844 |
| RsTa-C01_0844_1249R | agcggcatgctactatcatc | RsTa-C01_0844 |

**Table S3: MAGs used in phylogenetic and other comparative analyses (see Figs. 5, S13 and S14).**

| accession number or UHGG ID | accession number or UHGG ID | accession number or UHGG ID | accession number or UHGG ID |
|---|---|---|---|
| GCA_000432435.1 | GCA_017428625.1 | GCA_017631705.1 | MGYG000003581 |
| GCA_000432995.1 | GCA_017429345.1 | GCA_017631795.1 | MGYG000003591 |
| GCA_000433975.1 | GCA_017431505.1 | GCA_017631955.1 | MGYG000003628 |
| GCA_000434915.1 | GCA_017431755.1 | GCA_017634155.1 | MGYG000003629 |
| GCA_000435275.1 | GCA_017432165.1 | GCA_017648315.1 | MGYG000003641 |
| GCA_000435335.1 | GCA_017432485.1 | GCA_017650655.1 | MGYG000003752 |
| GCA_000577335.1 | GCA_017432865.1 | GCA_017935265.1 | MGYG000003768 |
| GCA_000752215.1 | GCA_017433225.1 | GCA_017935325.1 | MGYG000003809 |
| GCA_001261775.1 | GCA_017433905.1 | GCA_017936365.1 | MGYG000003841 |
| GCA_001512765.1 | GCA_017434895.1 | GCA_017936615.1 | MGYG000003842 |
| GCA_001513045.1 | GCA_017434985.1 | GCA_017938345.1 | MGYG000003863 |
| GCA_001513365.1 | GCA_017435305.1 | GCA_017938985.1 | MGYG000003872 |
| GCA_002069015.1 | GCA_017435385.1 | GCA_017939855.1 | MGYG000003889 |
| GCA_002069695.1 | GCA_017435665.1 | GCA_017939885.1 | MGYG000003947 |
| GCA_002119605.1 | GCA_017435765.1 | GCA_017942105.1 | MGYG000003955 |
| GCA_002160025.1 | GCA_017435865.1 | GCA_017942115.1 | MGYG000004027 |
| GCA_002201475.1 | GCA_017436085.1 | GCA_017959985.1 | MGYG000004032 |
| GCA_002293315.1 | GCA_017436225.1 | GCA_017963405.1 | MGYG000004049 |
| GCA_002297415.1 | GCA_017436255.1 | GCA_900048895.1 | MGYG000004075 |
| GCA_002305575.1 | GCA_017436285.1 | GCA_900100595.1 | MGYG000004076 |
| GCA_002308635.1 | GCA_017436375.1 | GCA_900101355.1 | MGYG000004106 |
| GCA_002309055.1 | GCA_017436725.1 | GCA_900176635.1 | MGYG000004115 |
| GCA_002309275.1 | GCA_017436795.1 | GCA_900184925.1 | MGYG000004116 |
| GCA_002314305.1 | GCA_017436915.1 | GCA_900197595.1 | MGYG000004129 |
| GCA_002314645.1 | GCA_017437005.1 | GCA_900240385.1 | MGYG000004131 |
| GCA_002315505.1 | GCA_017437165.1 | GCA_900291955.1 | MGYG000004146 |
| GCA_002315685.1 | GCA_017438225.1 | GCA_900314355.1 | MGYG000004159 |
| GCA_002315765.1 | GCA_017438505.1 | GCA_900314795.1 | MGYG000004229 |
| GCA_002315935.1 | GCA_017438585.1 | GCA_900315875.1 | MGYG000004265 |
| GCA_002316805.1 | GCA_017438845.1 | GCA_900315985.1 | MGYG000004279 |
| GCA_002320555.1 | GCA_017439545.1 | GCA_900316025.1 | MGYG000004292 |
| GCA_002329225.1 | GCA_017439585.1 | GCA_900316345.1 | MGYG000004322 |
| GCA_002329285.1 | GCA_017439765.1 | GCA_900316385.1 | MGYG000004334 |
| GCA_002329365.1 | GCA_017439845.1 | GCA_900316925.1 | MGYG000004413 |
| GCA_002338255.1 | GCA_017439875.1 | GCA_900317765.1 | MGYG000004437 |
| GCA_002350765.1 | GCA_017440755.1 | GCA_900317795.1 | MGYG000004470 |
| GCA_002359465.1 | GCA_017440765.1 | GCA_900318905.1 | MGYG000004496 |
| GCA_002361875.1 | GCA_017440825.1 | GCA_900320995.1 | MGYG000004543 |
| GCA_002361935.1 | GCA_017440965.1 | GCA_900321195.1 | MGYG000004555 |
| GCA_002363095.1 | GCA_017441385.1 | GCA_900321595.1 | MGYG000004560 |
| GCA_002371985.1 | GCA_017442225.1 | GCA_900539165.1 | MGYG000004616 |
| GCA_002372375.1 | GCA_017442265.1 | GCA_900539325.1 | MGYG000004643 |
| GCA_002378845.1 | GCA_017442345.1 | GCA_900539425.1 | MGYG000004723 |
| GCA_002391625.1 | GCA_017442445.1 | GCA_900540085.1 | MGYG000004727 |
| GCA_002397355.1 | GCA_017443805.1 | GCA_900540275.1 | MGYG000004807 |
| GCA_002399225.1 | GCA_017445905.1 | GCA_900540295.1 | MGYG000004815 |
| GCA_002399445.1 | GCA_017446095.1 | GCA_900541565.1 | MGYG000004850 |
| GCA_002407645.1 | GCA_017448285.1 | GCA_900542375.1 | MGYG000004892 |
| GCA_002407675.1 | GCA_017449935.1 | GCA_900542575.1 | MGYG000290008 |
| GCA_002409375.1 | GCA_017450005.1 | GCA_900542875.1 | MGYG000290033 |
| GCA_002409585.1 | GCA_017450135.1 | GCA_900543095.1 | MGYG000290055 |
| GCA_002409675.1 | GCA_017450185.1 | GCA_900543215.1 | MGYG000290071 |
| GCA_002410755.1 | GCA_017450645.1 | GCA_900544375.1 | MGYG000290075 |
| GCA_002411365.1 | GCA_017451425.1 | GCA_900545655.1 | MGYG000290084 |
| GCA_002424295.1 | GCA_017452785.1 | GCA_900546265.1 | MGYG000290120 |

| | | | |
|---|---|---|---|
| GCA_002437245.1 | GCA_017453685.1 | GCA_900546785.1 | MGYG000290125 |
| GCA_002437905.1 | GCA_017454485.1 | GCA_900547095.1 | MGYG000290153 |
| GCA_002438505.1 | GCA_017454815.1 | GCA_900547105.1 | MGYG000290169 |
| GCA_002438685.1 | GCA_017455295.1 | GCA_900547305.1 | MGYG000290178 |
| GCA_002449395.1 | GCA_017456015.1 | GCA_900547445.1 | MGYG000290186 |
| GCA_002451715.1 | GCA_017456445.1 | GCA_900547915.1 | MGYG000290211 |
| GCA_002451755.1 | GCA_017456845.1 | GCA_900548305.1 | MGYG000290227 |
| GCA_002451855.1 | GCA_017457155.1 | GCA_900549675.1 | MGYG000290295 |
| GCA_002490425.1 | GCA_017457955.1 | GCA_900549755.1 | MGYG000290335 |
| GCA_002491825.1 | GCA_017458105.1 | GCA_900549825.1 | MGYG000290439 |
| GCA_002493005.1 | GCA_017458845.1 | GCA_900551425.1 | MGYG000290441 |
| GCA_002493325.1 | GCA_017458945.1 | GCA_900552265.1 | MGYG000290485 |
| GCA_002493595.1 | GCA_017458975.1 | GCA_900553525.1 | MGYG000290552 |
| GCA_002493755.1 | GCA_017460565.1 | GCA_900554165.1 | MGYG000290559 |
| GCA_002494125.1 | GCA_017461005.1 | GCA_900554335.1 | MGYG000290567 |
| GCA_002834225.1 | GCA_017461235.1 | GCA_900554775.1 | MGYG000290582 |
| GCA_002834235.1 | GCA_017461625.1 | GCA_900555015.1 | MGYG000290690 |
| GCA_003268275.1 | GCA_017462405.1 | GCA_900555805.1 | MGYG000290832 |
| GCA_003438075.1 | GCA_017463345.1 | GCA_900754145.1 | MGYG000290859 |
| GCA_003499325.1 | GCA_017463985.1 | GCA_900761545.1 | MGYG000290913 |
| GCA_003505905.1 | GCA_017464405.1 | GCA_900764665.1 | MGYG000290943 |
| GCA_003514385.1 | GCA_017465085.1 | GCA_900766895.1 | MGYG000291000 |
| GCA_003517915.1 | GCA_017465285.1 | GCA_900767015.1 | MGYG000291022 |
| GCA_003520555.1 | GCA_017465465.1 | GCA_900768035.1 | MGYG000291055 |
| GCA_003521625.1 | GCA_017465505.1 | GCA_900769375.1 | MGYG000291062 |
| GCA_003538135.1 | GCA_017465825.1 | GCA_900769535.1 | MGYG000291101 |
| GCA_003612555.1 | GCA_017466205.1 | GCA_900772705.1 | MGYG000291106 |
| GCA_004103755.1 | GCA_017466365.1 | GCA_902761345.1 | MGYG000291180 |
| GCA_004555265.1 | GCA_017466905.1 | GCA_902761375.1 | MGYG000291183 |
| GCA_004555625.1 | GCA_017468425.1 | GCA_902762615.1 | MGYG000291203 |
| GCA_004560045.1 | GCA_017470025.1 | GCA_902763855.1 | MGYG000291207 |
| GCA_004560275.1 | GCA_017471345.1 | GCA_902765855.1 | MGYG000291209 |
| GCA_004768785.1 | GCA_017471605.1 | GCA_902766775.1 | MGYG000291284 |
| GCA_005601135.1 | GCA_017472575.1 | GCA_902766885.1 | MGYG000291342 |
| GCA_009746625.1 | GCA_017473165.1 | GCA_902767155.1 | MGYG000291369 |
| GCA_009774715.1 | GCA_017473435.1 | GCA_902767285.1 | MGYG000291396 |
| GCA_009774735.1 | GCA_017473555.1 | GCA_902767325.1 | MGYG000291400 |
| GCA_009777055.1 | GCA_017473775.1 | GCA_902768825.1 | MGYG000291443 |
| GCA_009777895.1 | GCA_017473925.1 | GCA_902771755.1 | MGYG000291527 |
| GCA_009779455.1 | GCA_017474045.1 | GCA_902773045.1 | MGYG000291606 |
| GCA_009779495.1 | GCA_017474125.1 | GCA_902773265.1 | MGYG000291625 |
| GCA_009783605.1 | GCA_017474205.1 | GCA_902773635.1 | MGYG000291638 |
| GCA_009784075.1 | GCA_017475765.1 | GCA_902774325.1 | MGYG000291646 |
| GCA_009785335.1 | GCA_017476375.1 | GCA_902777275.1 | MGYG000291751 |
| GCA_009785985.1 | GCA_017477885.1 | GCA_902785365.1 | MGYG000291753 |
| GCA_009786275.1 | GCA_017478005.1 | GCA_902785425.1 | MGYG000291756 |
| GCA_009786735.1 | GCA_017480115.1 | GCA_902785555.1 | MGYG000291761 |
| GCA_009787315.1 | GCA_017480265.1 | GCA_902786345.1 | MGYG000291777 |
| GCA_009917525.1 | GCA_017481085.1 | GCA_902787145.1 | MGYG000291785 |
| GCA_009930155.1 | GCA_017481115.1 | GCA_902790475.1 | MGYG000291791 |
| GCA_009936035.1 | GCA_017481395.1 | GCA_902790985.1 | MGYG000291793 |
| GCA_012511005.1 | GCA_017481575.1 | GCA_902796915.1 | MGYG000291887 |
| GCA_012511485.1 | GCA_017481865.1 | GCA_902798365.1 | MGYG000291916 |
| GCA_012516295.1 | GCA_017481885.1 | GCA_902798605.1 | MGYG000291946 |
| GCA_012517675.1 | GCA_017482035.1 | GCA_902798755.1 | MGYG000291949 |
| GCA_012519975.1 | GCA_017483265.1 | GCA_902800005.1 | MGYG000291967 |
| GCA_012520115.1 | GCA_017500965.1 | GCA_902800065.1 | MGYG000292005 |

| | | | |
|---|---|---|---|
| GCA_012520505.1 | GCA_017501865.1 | GCA_902801245.1 | MGYG000292015 |
| GCA_012522615.1 | GCA_017502005.1 | GCA_902802795.1 | MGYG000292022 |
| GCA_012523185.1 | GCA_017502085.1 | GCA_902802985.1 | MGYG000292086 |
| GCA_012524025.1 | GCA_017502405.1 | GCA_902803805.1 | MGYG000292095 |
| GCA_012728445.1 | GCA_017502425.1 | GCA_902809935.1 | MGYG000292159 |
| GCA_012839185.1 | GCA_017503135.1 | GCA_902810075.1 | MGYG000292186 |
| GCA_012839345.1 | GCA_017503625.1 | MGYG000000069 | MGYG000292239 |
| GCA_012839705.1 | GCA_017503705.1 | MGYG000000129 | MGYG000292281 |
| GCA_012840445.1 | GCA_017503785.1 | MGYG000000295 | MGYG000292320 |
| GCA_014384705.1 | GCA_017504645.1 | MGYG000000388 | MGYG000292352 |
| GCA_014465955.1 | GCA_017504765.1 | MGYG000000390 | MGYG000292489 |
| GCA_017379935.1 | GCA_017507805.1 | MGYG000000395 | MGYG000292574 |
| GCA_017380115.1 | GCA_017511525.1 | MGYG000000443 | MGYG000292617 |
| GCA_017380355.1 | GCA_017512465.1 | MGYG000000453 | MGYG000292623 |
| GCA_017380575.1 | GCA_017513165.1 | MGYG000000456 | MGYG000292633 |
| GCA_017380615.1 | GCA_017513525.1 | MGYG000000505 | MGYG000292754 |
| GCA_017381035.1 | GCA_017514345.1 | MGYG000000607 | MGYG000292779 |
| GCA_017381515.1 | GCA_017515685.1 | MGYG000000646 | MGYG000292782 |
| GCA_017382015.1 | GCA_017516315.1 | MGYG000000669 | MGYG000292814 |
| GCA_017382915.1 | GCA_017516445.1 | MGYG000000704 | MGYG000292850 |
| GCA_017383605.1 | GCA_017516485.1 | MGYG000000726 | MGYG000292854 |
| GCA_017383985.1 | GCA_017516965.1 | MGYG000000752 | MGYG000292917 |
| GCA_017384945.1 | GCA_017517205.1 | MGYG000000924 | MGYG000292997 |
| GCA_017385305.1 | GCA_017517525.1 | MGYG000000932 | MGYG000293109 |
| GCA_017386515.1 | GCA_017517665.1 | MGYG000000937 | MGYG000293123 |
| GCA_017387385.1 | GCA_017517865.1 | MGYG000001029 | MGYG000293161 |
| GCA_017387475.1 | GCA_017518415.1 | MGYG000001069 | MGYG000293167 |
| GCA_017389925.1 | GCA_017519685.1 | MGYG000001081 | MGYG000293196 |
| GCA_017390725.1 | GCA_017520635.1 | MGYG000001082 | MGYG000293206 |
| GCA_017390825.1 | GCA_017521125.1 | MGYG000001116 | MGYG000293234 |
| GCA_017391425.1 | GCA_017521925.1 | MGYG000001125 | MGYG000293289 |
| GCA_017391765.1 | GCA_017522205.1 | MGYG000001169 | MGYG000293318 |
| GCA_017391805.1 | GCA_017522225.1 | MGYG000001212 | MGYG000293344 |
| GCA_017391865.1 | GCA_017522455.1 | MGYG000001244 | MGYG000293370 |
| GCA_017393565.1 | GCA_017523865.1 | MGYG000001569 | MGYG000293383 |
| GCA_017396175.1 | GCA_017524165.1 | MGYG000001572 | MGYG000293642 |
| GCA_017396345.1 | GCA_017528095.1 | MGYG000001654 | MGYG000293699 |
| GCA_017397065.1 | GCA_017531385.1 | MGYG000001685 | MGYG000293767 |
| GCA_017397105.1 | GCA_017536525.1 | MGYG000001737 | MGYG000293769 |
| GCA_017397655.1 | GCA_017537245.1 | MGYG000001749 | MGYG000293853 |
| GCA_017397845.1 | GCA_017537365.1 | MGYG000001760 | MGYG000293887 |
| GCA_017398425.1 | GCA_017539305.1 | MGYG000001801 | MGYG000293910 |
| GCA_017398485.1 | GCA_017543285.1 | MGYG000001837 | MGYG000294093 |
| GCA_017399065.1 | GCA_017546535.1 | MGYG000001860 | MGYG000294096 |
| GCA_017399405.1 | GCA_017546685.1 | MGYG000001956 | MGYG000294115 |
| GCA_017400255.1 | GCA_017547465.1 | MGYG000002001 | MGYG000294117 |
| GCA_017401365.1 | GCA_017547725.1 | MGYG000002019 | MGYG000294154 |
| GCA_017402845.1 | GCA_017547865.1 | MGYG000002031 | MGYG000294286 |
| GCA_017403065.1 | GCA_017548025.1 | MGYG000002038 | MGYG000294288 |
| GCA_017403265.1 | GCA_017548045.1 | MGYG000002093 | MGYG000294362 |
| GCA_017403985.1 | GCA_017548305.1 | MGYG000002135 | MGYG000294373 |
| GCA_017404145.1 | GCA_017549105.1 | MGYG000002159 | MGYG000294398 |
| GCA_017404665.1 | GCA_017549285.1 | MGYG000002192 | MGYG000294482 |
| GCA_017405065.1 | GCA_017549565.1 | MGYG000002196 | MGYG000294524 |
| GCA_017406805.1 | GCA_017550105.1 | MGYG000002228 | MGYG000294526 |
| GCA_017407645.1 | GCA_017550625.1 | MGYG000002232 | MGYG000294547 |
| GCA_017407665.1 | GCA_017551295.1 | MGYG000002233 | MGYG000294561 |

| | | | |
|---|---|---|---|
| GCA_017407705.1 | GCA_017552225.1 | MGYG000002258 | MGYG000294654 |
| GCA_017407755.1 | GCA_017552785.1 | MGYG000002260 | MGYG000294693 |
| GCA_017407995.1 | GCA_017552845.1 | MGYG000002262 | MGYG000294701 |
| GCA_017408455.1 | GCA_017554125.1 | MGYG000002553 | MGYG000294705 |
| GCA_017408945.1 | GCA_017554405.1 | MGYG000002570 | MGYG000294754 |
| GCA_017409265.1 | GCA_017555765.1 | MGYG000002694 | MGYG000294801 |
| GCA_017409475.1 | GCA_017556785.1 | MGYG000002702 | MGYG000294827 |
| GCA_017410195.1 | GCA_017557825.1 | MGYG000002705 | MGYG000294848 |
| GCA_017410265.1 | GCA_017559305.1 | MGYG000002805 | MGYG000294892 |
| GCA_017410545.1 | GCA_017560255.1 | MGYG000002857 | MGYG000294994 |
| GCA_017410605.1 | GCA_017560725.1 | MGYG000002934 | MGYG000294995 |
| GCA_017411385.1 | GCA_017613735.1 | MGYG000002984 | MGYG000295003 |
| GCA_017411425.1 | GCA_017614835.1 | MGYG000003004 | MGYG000295013 |
| GCA_017411645.1 | GCA_017615405.1 | MGYG000003059 | MGYG000295030 |
| GCA_017411835.1 | GCA_017618135.1 | MGYG000003156 | MGYG000295033 |
| GCA_017414705.1 | GCA_017618535.1 | MGYG000003160 | MGYG000295123 |
| GCA_017414725.1 | GCA_017621885.1 | MGYG000003161 | MGYG000295151 |
| GCA_017414965.1 | GCA_017622415.1 | MGYG000003176 | MGYG000295160 |
| GCA_017415025.1 | GCA_017623315.1 | MGYG000003211 | MGYG000295212 |
| GCA_017415515.1 | GCA_017624355.1 | MGYG000003236 | MGYG000295220 |
| GCA_017416065.1 | GCA_017624675.1 | MGYG000003243 | MGYG000295236 |
| GCA_017416345.1 | GCA_017624735.1 | MGYG000003278 | MGYG000295277 |
| GCA_017416765.1 | GCA_017625095.1 | MGYG000003296 | MGYG000295278 |
| GCA_017418545.1 | GCA_017625435.1 | MGYG000003412 | MGYG000295283 |
| GCA_017419505.1 | GCA_017625835.1 | MGYG000003428 | MGYG000295359 |
| GCA_017422485.1 | GCA_017626295.1 | MGYG000003437 | MGYG000295423 |
| GCA_017423825.1 | GCA_017626835.1 | MGYG000003453 | MGYG000295437 |
| GCA_017425455.1 | GCA_017629015.1 | MGYG000003473 | MGYG000295480 |
| GCA_017425745.1 | GCA_017629275.1 | MGYG000003509 | MGYG000295483 |
| GCA_017427315.1 | GCA_017630985.1 | MGYG000003524 | MGYG000295535 |
| GCA_017428085.1 | GCA_017631355.1 | MGYG000003557 | MGYG000295546 |
| GCA_017428525.1 | GCA_017631515.1 | MGYG000003576 | GCA_000014725.1* |
| | | | GCA_900129805.1* |

* outgroup sequence

**Table S4. ATP/ADP translocase genes used in phylogenetic analysis (see Fig. 6).**

| UniProtKB ID | UniProtKB ID | UniProtKB ID | UniProtKB ID | UniProtKB ID | ID* |
|---|---|---|---|---|---|
| R6GHN2 | A0A2A4YG67 | A0A3B7D973 | A0A431IJ07 | A0A0G0BB48 | AJ582018 |
| B0RZB7 | F8L6N5 | H8K3Y2 | A0A2N2F6F9 | A0A7V1HE39 | AJ582019 |
| Q1MSG2 | A0A2H0VPK6 | A0A6H3RXP3 | A0A523TUZ0 | A0A431IIR0 | AY120885 |
| A0A0K8MFA8 | A0A6M2A234 | A0A126E8I3 | A0A0F9ZA13 | A0A0D2K661 | AP002546 |
| A0A4Q7DJM4 | A0A6M1YH83 | A0A0F3RDJ1 | A0A0G0BWV0 | A0A1F4QN25 | NC_002491 |
| A0A1F8JKC1 | A0A2H9SU81 | A0A0F3R7J0 | A0A1F4PIB3 | A0A345ZCK4 | Ntt2_Cab |
| A0A7W0FTB3 | A0A0C1C570 | H8K922 | A0A1F4QN61 | A0A1F4PQA9 | AJ010587 |
| A0A1M3BBW7 | A0A7W1RFY4 | A8GLZ6 | A0A523RJZ1 | A0A345ZCK5 | AE002346 |
| A0A0U5JD23 | D6YSH7 | A0A0C2QYA1 | A0A3D4XWN9 | A0A0G0V8C0 | AJ582023 |
| A0A2H0VTK1 | A0A2E3LYY9 | A0A7U0FXV9 | A0A1V5QV55 | A0A0G0V828 | AJ582020 |
| A0A3D2GUK3 | A0A7W1JM80 | A0A510G6D1 | A0A0G1B5V2 | V6DIH2 | AJ582022 |
| A0A077AUX9 | A0A090CYR0 | A8EXC0 | A0A355KQK0 | A0A7W1KD37 | AJ582021 |
| A0A7W1M910 | A0A1G3Z804 | A0A0F3QE15 | A0A0G1B3B1 | A0A0G0B9Z5 | NM_106679 |
| A0A1M3GVC4 | A0A3D2HI91 | A0A1C3EWJ7 | A0A355KRG2 | A0A0G0JXR7 | AC013453 |
| A0A358T439 | A0A1J4XF64 | A0A0F3QKS4 | A0A355KQN5 | A0A0G0I272 | Y10821 |
| A0A1W6N3J5 | A0A1Q3P7P4 | A0A261DBG2 | A0A349U2F8 | A0A0G0I2Y5 | AY098893 |
| A0A355VTN1 | A0A3S8T1B0 | A0A4Q3DK00 | A0A0G0IDH3 | A0A7T9BFC6 | AP003234 |
| A0A355VUV8 | A0A5B8XEK2 | A0A2A5B6J7 | A0A0G0J7G9 | A0A0F9Y0G0 | AJ251356 |
| A0A7W2B8S5 | A0A6J5JXQ0 | A0A3S0J7L7 | A0A7S1N3C9 | | AE008630 |
| A0A1M3IUX2 | A0A7V9ZM35 | Q842J1 | A0A0G0JZ48 | | EAA26488 |
| A0A1M3IQK7 | A0A258XDK4 | A0A0F4VNY1 | A0A2E3UXJ6 | | AJ507306 |
| A0A0W0YL02 | A0A1M3IT18 | A0A1V2N7Z3 | A0A7T8YHP0 | | AJ507305 |
| A0A7W1REL4 | A0A2M7T3X8 | A0A424FMM2 | A0A7T8YHT3 | | AJ235271_1 |
| A0A4S1MAI1 | A0A2D9KQJ9 | A0A2T4VXV6 | A0A7T9BEV6 | | AJ507307 |
| A0A1F8JIF2 | A0A2N6AEX9 | C6XHQ3 | A0A0F9XX69 | | AE008663 |
| A0A6M1ZTJ2 | A0A5C0UI02 | D0V1N0 | A0A0G0AQJ7 | | EAA26151 |
| A0A6M1YG67 | A0A2R8F168 | F7XV04 | A0A0G0AA79 | | AJ507312 |
| A0A212KR42 | A0A2U3RFE7 | A0A2A5BCR7 | A0A0D2I1H5 | | AJ235273 |
| A0A7H5P4T7 | A0A0F3NT69 | A0A0H3AUS9 | V6DI64 | | AJ507311 |
| A0A0H2X0K7 | A0A2U3QZ94 | A0A7W1MBQ0 | A0A1F4QNL7 | | AJ507313 |
| O84068 | A0A2U3REE8 | A0A258XCV7 | A0A2E6XWN8 | | AJ507309 |
| A0A0H3MGN2 | A0A2U3RPD8 | A0A258X8X1 | A0A2E6XVP0 | | AJ235272 |
| A0A069ZTL9 | A5CCY4 | A0A258XDT1 | A0A345ZCC7 | | AJ507310 |
| S7KMH9 | A0A5P2GM32 | A0A2H0VVI2 | A0A2A4WV73 | | AJ507308 |
| A0A3B0PRX3 | A0A2U3R2Y2 | A0A3B8U8W6 | A0A2A4WU23 | | AE008626 |
| Q823H5 | A0A0F3RK25 | A0A6M1ZTP4 | A0A3D4XV95 | | EAA25317 |
| A0A656IUS2 | A0A0F3MIA5 | K1Y0X6 | A0A0F9XZ47 | | AE008614 |
| S7IUE0 | D5AVX6 | A0A6M2AEZ9 | A0A0G0IC44 | | EAA25449 |
| A0A0E2IZV9 | Q68XS7 | A0A0H5DUE5 | A0A0G0J7L7 | | AJ507303 |
| Q5L657 | Q83W30 | A0A090D030 | A0A0G0DCP8 | | AJ507302 |
| A0A5C1MW90 | C4YY86 | A0A1F8JLG4 | A0A7S4MBI5 | | AJ235271 |
| A0A173DZV0 | A0A161QL04 | A0A0G0Y769 | A0A7S0IL69 | | AJ507304 |
| W8JLD3 | A0A0H3AVQ4 | A0A0G0V565 | A0A7S4B8E2 | | AJ582017 |
| Q254E3 | C4K173 | A0A0G0Y710 | A0A7S4EWG1 | | NC_003235 |
| V8TP30 | H6PVW1 | A0A523U2E7 | A0A7S4B8I1 | | NC_003236_2 |
| A0A0F7WSG7 | Q7PAI8 | V6DFH5 | A0A7S3TPR1 | | NC_003236_3 |
| A0A0F7X0C7 | A0A0F3QLL0 | A0A7W1KD24 | A0A7S3XCL4 | | NC_003236_1 |
| A0A2R8FAW5 | Q92JI6 | A0A7W1KD99 | A0A7S0LVZ6 | | |
| A0A1A9HYH4 | H8LP27 | A0A0D2JKG0 | A0A7T8YHP5 | | |
| A0A2A4WVV1 | C3PM91 | A0A0F9YPF6 | A0A0G0KIK2 | | |
| A0A3B8U7H6 | H8KBF8 | A0A7V1MPS5 | A0A0G0GNG3 | | |
| A0A6G4ZQ23 | Q83W40 | A0A7V1HEP3 | A0A0F9XZR0 | | |
| A0A6M1YYA5 | A8F0G0 | A0A0F9YC01 | A0A0G0KKH5 | | |
| A0A317JCP9 | H6QKS7 | K1XSV0 | A0A7T9BE65 | | |
| A0A486XJX4 | A0A0S2CAX6 | A0A7T9H7B6 | A0A0F9YAB8 | | |

* Sequence IDs of the Schmitz-Esser et al. (2004) dataset

**Table S5. Eukaryotic-/Eukaryotic-like genes used in phylogenetic analyses (see Figs. 4 and S9).**

| UniProtKB ID used in Fig. 4A | UniProtKB ID used in Fig. 4B * | UniProtKB ID used in Fig. S7A * | UniProtKB ID used in Fig. S7B * | UniProtKB ID used in Fig. S7C * |
|---|---|---|---|---|
| A2EJW2 | A0A0F9MGS0 | A0A061H4X5 | A0A0F7FJ36 | A0A1D2NJL0 |
| A0A6U1Z744 | X0S206 | A0A8H7JWK5 | A0A429GSS9 | A0A813DRP3 |
| A0A6U2ELB9 | A0A7K4GPA7 | A0A8H7IUB6 | A0A3R9RI95 | A0A812IE99 |
| C1E7N3 | A0A524EX74 | A0A163GEM8 | A1RX13 | A0A812JE06 |
| A0A6U0NFR7 | A0A5C9E0T7 | A0A6A5RGB5 | A0A1J4JF35 | A0A812TX60 |
| A0A6U3GEW5 | A0A5C9ERY9 | A0A6A6ZV80 | A0A1J4KDX1 | A0A812K2K7 |
| A0A699YPG0 | A0A8H4D775 | A0A8K0R389 | A0A1J4K7N4 | A0A813AUC2 |
| A0A6U1H2R9 | A0A2H0ZHW0 | A0A6A5QU68 | TRINITY_DN8548_c0_g1_i1.p1 | A0A1Q9CR86 |
| A0A6V1YRM0 | A0A0L0NZ33 | A0A178AMZ5 | A0A1J4KD15 | A0A7S1LKL5 |
| A0A6U2EHJ1 | A0A510P9E2 | A0A4Q4S4B3 | A0A1J4KRZ2 | A0A7S2IRI7 |
| A0A0D2LTI7 | A0A890D766 | A0A6A5JYC4 | A2DXF5 | A0A7J6L878 |
| A8J4U2 | A0A8H7GXT7 | M2U9S4 | A0A1J4K1A2 | A0A7J6LHD6 |
| D8UDK2 | Q6BZD6 | M2SNR4 | A0A1J4KUG9 | C5L3R7 |
| A0A0K9RC48 | C4YEB1 | A0A8H7BFT7 | A0A1Y1S6A6 | Q3SDK0 |
| A0A446LWG6 | A0A2N1J7A6 | A0A6A5UZE4 | L2GPN7 | Q3SDM6 |
| A0A446LWE5 | G0R2Q5 | A0A2V1DYD2 | I3EPT9 | Q3SDJ6 |
| A0A446KLF2 | J9EG22 | A0A6A5U2U5 | A0A177EMV5 | I7MHF3 |
| A0A446KLA1 | A0A1I9G785 | A0A6A6UYI8 | Q22Z33 | J9FS83 |
| A0A5J9TXC9 | A0A1I8FUI2 | A0A6A6W8B4 | M7X1F0 | A0A1J5WYR8 |
| A0A2G5E2A4 | A0A1I8I979 | A0A0D2AI17 | B0EHJ8 | A0A7S2XY36 |
| A0A2G5E294 | A0A336KHI3 | A0A3N4LBX1 | K2GBZ6 | A0A7M5WXS5 |
| A0A2G5E279 | A0A7K5SL79 | A0A3N4JA11 | A0A0N4ZVF7 | A0A0G4FIG9 |
| A0A5D3DBB3 | A0A7K7MQT5 | A0A5J5FBU2 | A0A0N5BX46 | A0A8J2F5M1 |
| A0A5A7UIP3 | A0A0L0FS10 | A0A3N4IQI6 | A0A0K0F9K4 | A0A8J1ZLN5 |
| A0A0K9P1Q3 | A0A7S3PVA0 | A0A8H3ENN8 | A0A090LGS6 | Q3SDK3 |
| A0A067GRF5 | E1B4V3 | A0A8H3I3Z6 | A0A8D8RFF8 | Q3SDN0 |
| A0A6N2LJI4 | A0A0R3WE28 | A0A8H6FY27 | A0A8D9BZJ7 | Q3SD33 |
| A0A6N2LJG2 | A0A7S2FKN1 | W9CLP2 | A0A8D8RJQ9 | Q3SCZ8 |
| A0A6N2LLJ9 | A0A5B8MUA6 | A0A8H2ZN92 | A0A8D9DSE8 | G0QZL5 |
| A0A371GFU8 | A0A177EFK8 | A0A061HJ54 | A0A8D9C1K6 | Q231B4 |
| A0A0L9TV98 | I3EH04 | N1JDN0 | A0A1B6JRI2 | A0A8B6XT86 |
| A0A7J9CP80 | K2GS41 | A0A8H7X6D0 | A0A1B6LQE9 | A0A7S4IGM2 |
| A0A5J5AT40 | Q4QT07 | A0A2V1CHG3 | A0A0A9XGR4 | A0A4D9C553 |
| A5B7L8 | B0EQW5 | A0A370TYT6 | A0A146M8W9 | A0A819J260 |
| A0A7N0VJ53 | Q5EGP4 | A0A559M7G0 | A0A023FBL8 | A0A816VTI4 |
| G3QAS5 | Q3SDK3 | A0A8H8S098 | A0A812B110 | A0A819MPL4 |
| A0A4Z2AY33 | Q3SDN0 | A0A4Q4SX16 | A0A812AZZ3 | A0A816PWM8 |
| A0A7J8JP11 | A0A8J2F5M1 | A0A4Q4VKJ9 | A0A1W0WS83 | A0A819BVD7 |
| A0A7J8JP19 | A0A1J4JVC5 | A0A4V1XFC0 | A0A8B7YH20 | A0A818N9U5 |
| A0A151P2S4 | TRINITY_DN7382_c0_g1_i1.p1 | A0A4V1XEQ1 | A0A8B8CKK3 | A0A821P7P9 |
| C3YDA6 | A0A852KM46 | A0A4Q4ZRQ1 | A0A8B8CGL8 | A0A815S408 |
| A0A1S3HH50 | A0A7K7LJJ0 | A0A4Q4WVV0 | A0A6J8AYA3 | A0A818TY56 |
| A0A6J8EBK4 | A0A7K9USC6 | A0A4Q4ZPE7 | A0A8B6F6T3 | A0A821C8F7 |
| A0A6J8E7S2 | A0A8C4Q5Z8 | A0A1Y2WCP9 | A0A076FH86 | A0A815TV65 |
| R7V8P8 | A0A1J4J649 | W3X4Y5 | K0KXS0 | A0A814XKZ4 |
| A0A6F9DUG7 | A2ESK3 | A0A1Y2XGL1 | A0A1E3P3J0 | A0A814QLZ4 |
| A4S845 | A0A7S3CVP5 | A0A1Y2TZ92 | A0A061AVJ6 | A0A815JDL7 |
| L1IXS2 | T1JME3 | A0A1Y2EE74 | A0A830I1H1 | A0A820TZQ3 |
| A0A6U4DLJ2 | E9G2W8 | A0A1W2TMU9 | Q5D6W2 | A0A7S3YE35 |
| D2VZY0 | A0A8J2WG29 | A0A4Z0Z3W2 | A2DJK6 | A0A7S3YE77 |
| A0A2V2V2B7 | J9I1W7 | E3Q2P6 | A0A1J4J4M1 | A0A3S1BD86 |
| V5BIE1 | S7W8I7 | A0A066XFF5 | A2EN26 | L7K0I4 |
| A0A7J6Y812 | A2FIJ0 | A0A1S1VTF1 | A2DHX2 | L2GVC0 |
| A0A2V2VCT5 | A0A820ZVC4 | A0A8H6JPY0 | A0A1S3D0W5 | A0A671LEH6 |
| A0A2V2X2H9 | A0A820Y630 | A0A4U6XBJ2 | A0A7S0HB51 | A0A672RYS7 |
| A0A3R7PIC2 | A0A818ZAY2 | A0A5Q4BX13 | Q98RX2 | A0A6P3WCN0 |
| A0A061IZA1 | A0A814III5 | N4VJD2 | A0A654GLI6 | A0A3B4DW63 |
| A0A422NP63 | A0A814KCR9 | A0A2T2ZT58 | A0A7M3QA85 | A0A5N5KZE5 |
| A0A1X0NNW0 | A0A251U4W4 | A0A5N5MTR3 | A0A196SLS2 | A0A4W4EQZ8 |

Q385X0
A0A1G4ID09
A0A3L6KRT6
D0A6U2
G0V079
G0UAY8
S9TWK5
S9VC96
A0A7G2CKC4
L0AV63
A0A131XVM0
A0A6P6XR97
I7MKZ2
A0A1R2C869
Q22TX5
A0D4B0
A0D283
A0A1J4KTU1
A0A7E4ZT81
A0CGY4
A2FJN1
A2DVU2
A2FCN9
A0A146K3G2
B3RJY9
A0A6T2AG77
A0A151ZH60
A0A6T6W9C0
A0A6V5V221
A0A6U8BAU6
A0A6T1BLD3
A0A6V1QD79
A0A6V3D4S8
A0A6U4IVP2
T0PWX7
A0A6T6JJ43
A0A6P6RQX7
A0A1J4KIC4
A2DY61

A0A2H4SKE0
X6MGW6
A0A7K4GX73
A0A850M276
A0A1J4K2N7
A0A5B9DE64
A0A5B9DFX9
A0A497QHU6
A0A842QXE0
A0A524F9Y6
A0A7K4GLD5
A0A524D9Y9
A0A5C9DTX2
A0A5C9ERY4
A0A842T0B9
A0A524C8J4
A0A524FK55
X0ZEA4
A0A5C9DWI3
A0A7K4H0C6
A0A532TI34
A0A523W0B6

A0A5N5NSZ1
A0A086T1H5
A0A0F7ZQD3
A0A167A5G7
A0A7U3Q103
A0A8K0J7J9
M1W7M1
A0A063CBP1
A0A179HZ11
A0A367L240
A0A2C5Z555
A0A544ZV83
A0A0D2DR91
A0A0D1YEH3
A0A177FHB8
N1Q0R6
A0A6A6C2K9
G8C275
A0A0C7N6G5
A0A1G4JTJ0
Q6CTC6
A0A5P2U6P3
A0A0A8LB19
W0THH7
A0A8F2RLY9
D2V9Q0
TRINITY_DN11080_c4_g1_i1.p1
I4Y7M5
A0A316UT68
V5F1D1
I2G6S2
A0A1K0FYU1

D8LW26
A2FFL0
J9JAC8
A2EAD2
A2EUE8
A0A1J4KZL1
A0A1J4KD55
A0A7S3DNW8
A0A1Z5K7K3
A0A8J7XL72
A0A843BTD3
A0A7C1CCU2
A0A522XYA9
A0A557SVK8
A0A838D347
A0A654MAD2
A0A550GFH8
A0A550GJA5
A0A7C5RKA0
A0A832HX82
A0A7J3Z5U8
A0A6B2C8A9
A0A7C5TWK9
E1QT99
F0QSH4
A0A497GTR0
A0A7J4AYN1
A0A2K2V4B8
A0A2H5UVI2
A0A662PPR2
A0A7C4CNQ4
A0A7C4WGB0
A0A7J2HY37
A0A256Y5X4
A0A7J3NL68
A0A832BXS7

A0A673IPV9
A0A7K8ER36
A0A8C3D6V0
A0A532TY91
A0A524ELR9
A0A850MQ89
V4C1F0
A2EJI6
A0A816MK02
A0A820CEI0
A0A820L5M8
A0A8C5LU88
A0A8C5LUB8
A0A834ITL8
A0A8H7GXT7
A0A2V1B0Y8
A0A2V1AFA3
A0A8J5QGJ0
A5E1E8
C4YM78
A0A1D8PT80
A0A8H6F3V0
A0A1L0BQ30
M7WFW0
TRINITY_DN9732_c0_g2_i1.p1
A2G2G1
A2E2M2
A0A437AKV0
A0A7S3CVP5
A0A8J4V5F7

| **UniProtKB ID used in Fig. S7D** * | **UniProtKB ID used in Fig. S7E** * | **UniProtKB ID used in Fig. S7F** * | **UniProtKB ID used in Fig. S7G** * | **UniProtKB ID used in Fig. S7H** * |
|---|---|---|---|---|
| A0A0C2I723 | A0A812IE99 | G0QQ18 | A0A812IE99 | A0A821TKA4 |
| A0A1D2MB91 | A9CRJ3 | A0A0A8LBP3 | A9CRJ3 | A0A818Z5J4 |
| A0A1D2MC16 | A0A1Y1S664 | Q6CJ90 | A0A1Y1S664 | A0A821JVW3 |
| A0A835CSA2 | A0A0R0M7T3 | W0T9B2 | A0A0R0M7T3 | A0A1B7SPZ4 |
| A0A835CQY0 | S7W8I7 | G8ZY21 | S7W8I7 | W1Q751 |
| A2DCC5 | A0A7S3CVP5 | A0A7H9HWT1 | A0A7S3CVP5 | A0A2G5B9W1 |
| A0A4Q9KY41 | A0A8J4V5F7 | A0A7G3ZNG9 | A0A8J4V5F7 | A0A0K8S640 |
| R0MLC0 | Q6BZD6 | H2ANL1 | Q6BZD6 | A0A6H5GSU7 |
| A0A6A5BNC4 | A0A2V1B0Y8 | A0A8H2ZGY6 | A0A2V1B0Y8 | A0A0K8S654 |
| D2VNZ8 | TRINITY_DN14133_c1_g1_i1.p1 | C7GQX9 | TRINITY_DN14133_c1_g1_i1.p1 | A0A0A9XIZ1 |
| B0EK11 | A0A812DZM0 | B3RHE4 | A0A812DZM0 | A0A6A4JR53 |
| K2GDQ4 | A0A369SI02 | G2WJ86 | C4YM78 | T1IAQ6 |
| A2EXV3 | C4YM78 | A0A8B8UW66 | A0A1D8PT80 | A0A0N7Z8L5 |
| Q3SD70 | A0A1D8PT80 | J6ENU0 | A0A8H6F3V0 | A0A224XWC7 |
| A0A8S1NU57 | A0A8H6F3V0 | A0A0L8RFE5 | B9WM58 | A0A069DQN9 |
| A0A8J5QY30 | B9WM58 | J8LKE2 | C5M3H7 | A0A8I6RWP9 |
| A0A0C9QHS2 | C5M3H7 | G8BUE5 | A0A8J5QGJ0 | A0A6J0YCN0 |
| E2ATU0 | A0A8J5QGJ0 | A0A0W0DYK8 | A0A8H8DBP2 | A0A6P3TNI1 |
| A0A836EZX9 | A0A8H8DBP2 | Q6FMR0 | A0A421JIW5 | A0A8C8JIG8 |
| A0A833RY36 | A0A421JIW5 | J7S7W3 | A5E1E8 | A2DMY4 |
| A0A154PCY4 | A5E1E8 | G0VE40 | G3B8G4 | N9T9B7 |
| A0A7M7GVQ5 | G3B8G4 | A0A1Q3ABC0 | G3B8G3 | A0A2G5UH10 |
| A0A6S7GTF6 | G3B8G3 | A0A1Q2ZX36 | A0A1Y2CT12 | B6IHN6 |
| TRINITY_DN6741_c0_g1_i1.p1 | A0A1Y2CT12 | A0A8J2T4F1 | A0A1Y3NQ15 | E4YE67 |
| A0A1E3P0Y1 | A0A1Y3NQ15 | A0A1S7HVI1 | A0A1Y1XCZ7 | A0A7D9CZM6 |
| A0A1A0H717 | A0A1Y1XCZ7 | A0A7H9B7A8 | A0A1Y2D0F9 | A2EQ49 |
| A0A512UI12 | A0A1Y2D0F9 | A7TQ91 | A0A2N1J7A6 | A0A1J4K8L6 |
| A0A1L0BSK3 | A0A2N1J7A6 | G8JR72 | A0A1E3PR62 | A0A1J4KG41 |
| Q6BYB0 | A0A1E3PR62 | R9XEI8 | A0A850MMV7 | A0A1J4KRH2 |
| Q5NT15 | A0A850MMV7 | A0A1G4MCU1 | V4C1F0 | A0A1J4K6W7 |
| K2GWH3 | V4C1F0 | C5DEM2 | T1FYQ5 | A0A6A7G7T8 |
| K2H1L3 | T1FYQ5 | A0A1G4IXA3 | A0A369SI02 | A0A6A7GAK5 |
| B0E8D8 | Q3SCY3 | A0A1E3NXK2 | Q3SCY3 | A0A1R2CEW5 |
| A0A510P9E2 | Q3SDN0 | A0A1D2VNR0 | Q3SDN0 | A0A7S3G5U1 |
| A0A2H0ZHW0 | Q3SDK3 | G3B8G4 | Q3SDK3 | A0A7S3G9Z6 |
| A0A0L0NZ33 | Q3SCZ8 | G3B8G3 | Q3SCZ8 | A0A1J4KBR9 |
| G8YEB7 | A0A8J2F5M1 | M3K269 | A0A8J2F5M1 | A0A1J4K4P4 |
| G8YBW7 | A0A8J1ZLN5 | A0A0C2MUI2 | A0A8J1ZLN5 | A0A1J4L2Z7 |
| A0A1B6D225 | A0A1R2B4K9 | G0R2Q5 | A0A1R2B4K9 | TRINITY_DN10941_c1_g1_i3.p1 |
| X6MGW6 | D2VB98 | A0A5J9VNJ2 | F0ZUW8 | D3YVQ6 |
| A0A0R0M7T3 | A0A6P8J1G3 | A0A1Y3NQ15 | D2VB98 | A0A8C6I502 |
| A0A6P7S593 | F0ZUW8 | A2FIJ0 | A0A6P8J1G3 | Q0PD39 |
| A0A8B8E6D2 | L7K0I4 | A2G974 | L7K0I4 | A0A8C6I458 |
| S6B8L6 | A0A0F9ZEV9 | A0A1J4KTB4 | A0A0F9ZEV9 | A0A6P5PI80 |
| L0B2B3 | A0A7M7NPW1 | A0A1J4K7M7 | A0A7M7NPW1 | A0A6P5PP20 |
| J4CDX6 | A0A7M7RC83 | A0A1J4JG74 | A0A7M7RC83 | B2RZ46 |
| A0A813B3T2 | Q3SDK0 | A0A1J4JW69 | Q3SDK0 | A0A218WJE0 |
| A0A8J8T7F8 | Q3SDM6 | A0A0V0RA50 | Q3SDM6 | A0A1J4KXA3 |
| A0A3R7W9T9 | Q3SDM2 | Q3SDI3 | Q3SDM2 | A0A524DV28 |
| A0A6U9WAJ3 | Q3SD42 | A0A804KB13 | Q3SD42 | A0A7M5UI19 |
| A0A1E7EU26 | A0A672L021 | A0A077Z5W6 | A0A672L021 | T2MAC0 |
| A0A6A7FZ22 | A0A672L036 | A0A131ZVZ2 | A0A673JC67 | A0A183ICB6 |
| A0A0D9YVS9 | A0A671LSV7 | W6L7P9 | A0A6P6R6Z5 | S7XTQ9 |
| A0A7J7CM12 | A0A673JC67 | A0A7S3CVP5 | A0A3Q3VRG5 | Q3SDI2 |
| | A0A6P6R6Z5 | A0A1J4KG86 | A0A3Q3VU11 | A0CE74 |
| | A0A7J6B401 | A0A7C8VL15 | A0A667YXN5 | Q3SCZ7 |
| | W5UKR5 | A0A7C8PHK9 | A0A8C4A543 | A9UYR4 |

| | | | |
|---|---|---|---|
| A0A3Q3VRG5 | A0A6G1M4C8 | A0A672L036 | A0A8B7DFA2 |
| A0A3Q3VU11 | A0A7C8NBB5 | A0A671LSV7 | L8GIC5 |
| A0A667YXN5 | A0A369SI02 | A0A7J6B401 | L8HAR7 |
| A0A8C4A543 | A0A497QHU6 | W5UKR5 | A0A1C7NPA6 |
| A0A3B4DB85 | A0A5B9DE64 | A0A3B1JUY8 | A0A8H7RBX5 |
| M4ANU7 | A0A1V0SEB6 | A0A5N5Q526 | A0A8H7BVQ6 |
| A0A3B5MRM0 | I3ER36 | M4ANU7 | V4A2G3 |
| A0A3Q4AL16 | A2DRV6 | A0A3B5MRM0 | A0A6J8BEV0 |
| A0A8C2WLV0 | A0A0A1U9N5 | A0A3Q4AL16 | A0A8B6D857 |
| A0A6J2W5V0 | A0A7S1XW42 | A0A8C2WLV0 | A0A2K6F864 |
| A0A5N5Q526 | A0A7J6EUN9 | A0A6J2W5V0 | A0A2K6F866 |
| A0A3B1JUY8 | R0MLC0 | A0A3B4DB85 | A0A833R3N8 |
| A0A8C9SBL6 | TRINITY_DN16833_c1_g1_i5.p1 | A0A8C9SBL6 | A0A328CY65 |
| A0A2U3W9G9 | A0A890D766 | A0A2U3W9G9 | A0A1J5WW56 |
| M3Y931 | A0A2H0ZHW0 | A0A3Q7SCR8 | A0A813S638 |
| A0A2Y9IDN2 | A0A510P9E2 | F1PTE3 | A0A815Q636 |
| A0A2U3W4P2 | A0A0L0NZ33 | A0A6J2ELM6 | A0A819MUY6 |
| A0A6J2ELM6 | A3GFW1 | A0A2U3W4P2 | A0A815LWT6 |
| A0A452SH85 | Q6BZD6 | A0A452SH85 | A0A819MLH3 |
| A0A3Q7SCR8 | A0A812DZM0 | M3Y931 | A0A814PMR1 |
| F1PTE3 | A0A146K9N6 | A0A2Y9IDN2 | A0A819M2X1 |
| A0A8C5WCM5 | I4Y7M5 | A0A8C5WCM5 | A0A813XYH8 |
| A0A1I8GTC3 | A0A4V4ML05 | A0A1I8GTC3 | A0A814M3X0 |
| A0A7S1EZY4 | A0A4T0QER5 | A0A7S1EZY4 | A0A814B7Z5 |
| A0A7S2IRI7 | A0A4T0FQE2 | A0A7S2IRI7 | A0A814LNX5 |
| A0A813DRP3 | A0A2G5B6K2 | A0A813DRP3 | A0A815WYL7 |
| A0A812JE06 | A0A1J4L1H1 | A0A812JE06 | A0A815FY79 |
| A0A812TX60 | A0A1J4K4P4 | A0A812TX60 | A0A819USI6 |
| A0A813AUC2 | A0A1J4KKP2 | A0A812K2K7 | A0A816Z0F8 |
| A0A1Q9CR86 | Q23W08 | A0A813AUC2 | A0A815HVY4 |
| A0A812K2K7 | | A0A1Q9CR86 | A0A818FR11 |
| | | | A0A818HMD5 |
| | | | A0A817DM21 |
| | | | A0A818JL34 |
| | | | A0A818MM80 |
| | | | A0A820Q620 |
| | | | A0A821BPC6 |
| | | | A0A818SBF8 |
| | | | A0A813QBB6 |
| | | | A0A816CFM2 |

| UniProtKB ID used in Fig. S7I * | UniProtKB ID used in Fig. S7J * | UniProtKB ID used in Fig. S7K * | UniProtKB ID used in Fig. S7L * | UniProtKB ID used in Fig. S7M * |
|---|---|---|---|---|
| A0A1E4RHJ0 | A0A3M1UVP2 | A3GFW1 | I1JKY2 | A0A1W0E6K4 |
| A0A8B9J6F6 | L7K0I4 | B0ESL8 | A0A0B2SHG0 | A0A8H7GXT7 |
| Q6FGX3 | L2GVC0 | K2HM75 | K7K3L6 | A0A8C7HQG5 |
| G5E2Z7 | S7XTQ9 | M7WHL4 | I7MDK8 | A0A8C7ML11 |
| A0A8M1FTS7 | T0MER4 | A0A1J4KV97 | G0QU16 | A0A4W5RCX9 |
| A0A662YRE0 | A0A059F2S3 | A2FIJ0 | I7LY19 | A0A1S3SZL2 |
| A0A3Q2NP45 | A0A1R2BEZ6 | A0A0V0RA50 | A0BU91 | A0A1S3SZM7 |
| A0A8C7FF29 | A0A7S4P836 | A0A7S3K8N0 | A0A5J4VWM7 | A0A1S3SZL4 |
| A0A0S7H0U4 | C5LKH4 | A0A078ABN3 | TRINITY_DN10457_c0_g1_i1.p1 | A0A1S3SZK7 |
| A0A2U9B8I3 | A0A812Q5K2 | A0A0V0QFZ1 | B0E8E1 | A0A1S3SZL6 |
| A0A4W5LAI3 | A0A8J2B4L0 | A0A8J1ZLN5 | M7VW78 | A0A1S3SZL0 |
| A0A8C7IGL2 | A0A5J5F566 | A0A8B7YSX1 | A0A7J6YYW5 | A0A8C7HRP8 |
| A0A8C7IDH7 | A0A0J9X8R7 | S7W8I7 | T1EJ40 | A0A8C8D7G3 |
| A0A8C8LLP2 | A0A5A9MZE3 | A9CRJ3 | A0BTF9 | A0A8C7Q6K0 |
| A0A384DFD5 | H9H629 | N9URX2 | A0A1J4K080 | A0A674C9N3 |
| A0A803J357 | A0A7S3D6C9 | M7WXJ3 | A0A7M5TRI7 | A0A674C9A5 |
| A0A803K6A7 | A0A4D9DI20 | K2GIL3 | A0A6A5PI19 | A0A8C7Q6D8 |
| A0A8C1H6E7 | A0A7S4GI31 | B0EGQ1 | A0A6A4R8Y0 | A0A4W5RJY3 |
| Q6P304 | Q39861 | G0R2Q5 | A0A1J7GDT5 | A0A8C8CZY7 |
| A0A669BTR5 | W4ITV4 | A0A1J4KPE5 | A0A3P8YVY0 | A0A3P8Z3G0 |
| A0A668VST9 | A0A2P9GG46 | A2DMZ1 | A0A8B9LQG6 | A0A8C7K7P4 |
| A0A3Q0S8J9 | A0A2P9CKT7 | A0A1J4JQ83 | A0A433PM26 | A0A8C8HWB6 |
| A0A667WH53 | A0A2P9BSY1 | A0A1J4KKP2 | A0DHR0 | A0A8C7TU69 |
| H2SGX0 | Q94659 | A0A1J4KCU8 | A0A817MCN2 | A0A8C7TPK2 |
| A0A3B5A1Y3 | Q9UB85 | TRINITY_DN12480_c0_g1_i1.p1 | A0A433DE42 | A0A060VW08 |
| A0A671WCX9 | Q9U466 | A0A146K690 | A0A4P9YVV2 | A0A4W5LTD9 |
| A0A3P8SFA4 | A0A024VVH2 | B0ER43 | A0A4P9XJF4 | A0A671TEJ5 |
| A0A3Q1F295 | A0A060RR02 | A0A1X0QHI2 | A0A1Y2F8X0 | A0A3P8W3Y6 |
| A0A8C6LZA1 | A0A2P9D242 | S7XUF5 | A0A8C4NAN0 | A0A3B3ZJE7 |
| H2MNT7 | A0A2P9B9K4 | A0A842R0Y3 | A0A1E3PFL4 | A0A3Q3GML6 |
| A0A3P9KVA3 | K6UKL1 | A0A842R6F5 | A0A2U1L0V4 | A0A6J2QWT6 |
| A0A3B3CV27 | W7ARZ9 | A0A0F9DFN3 | V7B6H7 | A0A6J2QYY2 |
| A0A671NG68 | D2VB98 | A2EV06 | A0A0S3RTU3 | A0A8C2WVL9 |
| A0A673HFT2 | A0A6A5ATY7 | A0A1Y1ZC38 | V7B2F7 | A0A3Q3GTA3 |
| A0A673HEV8 | A0A0N5B7E3 | A0A1Y1Z3T9 | A0A4D6NGP5 | A0A2I4CHV6 |
| A0A671LVR5 | A0A0K0EV93 | A0A4P9XMJ8 | A0A445J787 | A0A672GPV2 |
| A0A671M211 | A0A0N4Z015 | A0A397SAP9 | A0A151T9Y3 | A0A672GR95 |
| A0A8C1I4S5 | A0A090LNS9 | A0A372QVP2 | A0A2Z6MZY7 | A0A6G1PQP7 |
| A0A8C1FT99 | A0A0K0E4U5 | A0A2Z6RC45 | G7KX35 | A0A3Q3JV56 |
| A0A671LXY7 | A0A4U8URS0 | A0A2P4QMJ6 | B7FKS6 | A0A3B5AQ46 |
| A0A6P6R3B2 | A0A6P4YM79 | A0A2N1ND68 | I3SXF6 | A0A3P8SRD2 |
| A0A8N1YWU9 | A0A4Y2AL00 | A0A1D1Z9A9 | G7KX34 | A0A2D0SW35 |
| A0A672STW9 | A0A1I8I5X4 | A0A1D1Y4X4 | A0A2K3N539 | A0A8J4X8C3 |
| A0A672SSR4 | A0A267GBT0 | A0A8H4EP33 | A0A6P5NKL3 | A0A8C1RCY6 |
| A0A672SU05 | A0A1I8HBZ8 | A0A397UBB2 | A0A6J1DXN1 | A0A8C1P9W6 |
| A0A672SU16 | E4X453 | A0A397J5N9 | A0A6J1F3N4 | A0A6P3WCN8 |
| A0A8C1ZXG6 | E4YFK5 | A0A507FP66 | A0A6J1J588 | A0A6P8G119 |
| A0A8C1FUW3 | F1LAQ0 | F4RPA3 | A0A6J1I1J9 | A0A6J2VNP7 |
| A0A6P6JR13 | A0A0N4V9P2 | A0A180GLG9 | A0A6J1GBU9 | V4C1F0 |
| A0A7J6DCV7 | A0A2A6CCI0 | A0A2N5T6G8 | A0A2C9VBG7 | A0A0V1HW41 |
| A0A8C2DXU4 | R7TCP3 | A0A4T0FQE2 | A0A218W3F2 | A0A7N4V6I7 |
| A0A8C1L5F7 | A0A7M5WZ22 | M5E8W1 | A0A6I9RAS3 | F7EMD6 |
| A0A671NFZ2 | A0A023GIH7 | E6ZSW7 | A0A6I9RRU4 | A0A8C0HHI1 |
| A0A672N163 | A0A8C3EUA6 | A0A2N8U8K1 | A0A5B7AKH8 | A0A8C3FV55 |
| A0A673HCY8 | A0A8D0G4V8 | A0A5C3FH31 | D7SR02 | A0A8C3XUY1 |
| A0A4W4H640 | A0A8D1XRF6 | A0A8H7PPI3 | A0A2P5BS67 | A0A8C4UJ25 |
| A0A3B4BVR2 | A0A8B9BYZ8 | A0A1X2HBN1 | A0A8K0H8R8 | A0A8C5MSH7 |
| A0A5N5M6K9 | A0A3P8WB11 | A0A2T9YQM0 | A0A6J5XNJ7 | A0A8J1KXT4 |

A0A8C4BDM6
A0A2D0RVW6
A0A3B1K8N3
V9KRJ0
H3DHF1
A0A8D2JKY5
A0A670K8G6
A0A803SWP1
A0A8C9AWC4
A0A8B7PP98
D3PGL3
A0A482W300
A0A023FTB6
A0A8J9Z090
A0A814DN77
A0A814DVR9
A0A813WYI6
R7TE99
A0A7E6FI07
A0A7I8W2J6
S2JFB7
A0A8H4BS43
A0A0C9LVN5
A0A8H7R068
C4QZ12
Q55ET3
A0A814H4B2
A0A6P7V5T8
TRINITY_DN21322_c0_g1_i1.p1
A0A8K1CRQ3
A0A421JIW5
A0A8J5QGJ0
G3AJT2
H8X559
G8BFM2
A0A1L0BQ30

A0A669PIE6
A0A8C9SBL6
A0A816QNG3
A0A7S3DFN8
A0A7S2J5E9
D2V1N7
Q3SDM2
Q3SDJ6
Q3SD54
Q3SDM6
A0A0C2M9K0
A0A812IE99
A0A812K2K7
G0QZL5
Q3SCY3
A0A814ZZ42
A0A3G4ZV93
A2EXV3
V4C1F0
A0A3Q0KGU5
A0A8F2RLY9
A0A5Q0HU94
C4Y1Y5
TRINITY_DN15383_c0_g1_i6.p1
A0A7S3GPX1
A0A0F9RE60

C5M895
A0A367YIS9
A0A367XW76
A0A0H5CAD7
A0A1E4S882
I2JWX4
A0A7D9CYJ2
A0A448YNW1
A0A8K0J7J9
M1W7M1
F7W162
A0A0B0DJ31
U7PXU8
A0A094GEK6
A0A428NCL8
A0A428P6F5
A0A063CBP1
A0A545UPL6
D3B840
A0A890D766
A0A510P9E2
A0A2H0ZHW0
Q6BZD6
C4YEB1
A0A8H6F1C2

A0A6J5V7B8
A0A5E4EKJ3
A0A7J9E626
A0A7J9BWN7
A0A0D2S189
A0A5D2TN29
A0A5D2BFA5
A0A1U8J4P7
A0A7J8ZF24
A0A7J9N594
A0A7J9IZH6
A0A2U7S651
A0A5D2D7H0
A0A5D3A178
A0A0B0PHB8
A0A7J9BNG4
A0A6P6A8Y4
A0A8B8JSV2
A0A8B8JT80
V7AM67
A0A4D6MQ08
A0A0S3SXJ8
A0A371I7U0

A0A8C8VPJ8
A0A6P8SN84
A0A8M1N0P6
A0A5A9NHF9
A0A4X2JNH9
A0A3Q2PPV4
Q5ZIT5
G3NVY1
A0A7N4NL08
A0A811JV18
A0A1I7S8W3
A7S5Z2
A0A1I8I5X4
A0A267GBT0
G0QZL5
A0A3G4ZJY4
A0A817ZCR9
A0A817JHV4
A0A815JFP7
A0A814C117
A0A814X8Z0
A0A815KTP7
A0A818N9U5
A0A815S408
A0A821P7P9
A0A815BI71
A0A814Y729
TRINITY_DN16011_c1_g1_i13.p1
A0A1J4KQ65
A9CRJ3

| **UniProtKB ID used in Fig. S7N** * | **UniProtKB ID used in Fig. S7O** * | **UniProtKB ID used in Fig. S7P** * | **UniProtKB ID used in Fig. S7Q** * | **UniProtKB ID used in Fig. S7R** * |
|---|---|---|---|---|
| A0A8J2F5M1 | Q4G292 | C5M895 | V4C1F0 | A0A0L0NEF2 |
| A0A3G4ZT94 | A0A146K6A4 | A0A367YIS9 | A0A7S3GPX1 | A0A2K3QHR0 |
| A0A0S8JIQ7 | A0A2T6ZM98 | A0A367XW76 | B9WMJ7 | A0A8K0J7J9 |
| A0A0A1U4S6 | A0A3N4J5V2 | S7W8I7 | Q59X89 | A0A8K0WQ63 |
| A0A7S4ISA4 | A0A292PLE6 | A0A437AKV0 | C5M895 | W3X4Y5 |
| Q4G2B3 | Q9HDY0 | A9CRJ3 | A0A2P7YV86 | A0A1V1SRE9 |
| A0A1J4L0X4 | Q4G2D0 | A0A1W0E6K4 | A0A6C1E7V3 | A0A7C8MNX7 |
| A0A1J4L2Z7 | A0A0V0G7Q0 | A0A814XKZ4 | G8YBW7 | A0A4Z0Z3W2 |
| C5M895 | A0A6B2LN26 | A0A814QLZ4 | A0A0V1PV62 | A0A1Y2XGL1 |
| A0A367YIS9 | A0A0L0D2G5 | A0A815JDL7 | Q6BZD6 | A0A447CM14 |
| A0A367XW76 | A0A7S1YM28 | A0A820TZQ3 | TRINITY_DN12381_c1_g4_i1.p1 | B2AXF0 |
| A0A2P4QMJ6 | A2DSR0 | A0A816WMC3 | A0A842R0Y3 | G2QDJ0 |
| A0A2N1ND68 | A0A1B7SPZ4 | A0A818TY56 | A2EGH0 | F8MKM8 |
| A0A372QVP2 | W1Q751 | A0A821C8F7 | A0A8K0EKR2 | A0A0B0DJ31 |
| A0A1D1Y4X4 | A0A6S9AIZ9 | A0A815TV65 | A0A7D9E2I1 | F7W162 |
| TRINITY_DN11656_c0_g1_i2.p1 | A0A6U5J0A2 | A0A818N9U5 | A0A3M7QKZ9 | S3DHW5 |
| R4UVI8 | B8CFX8 | A0A821P7P9 | A0A7R8H7D1 | A0A2T3B6K3 |
| A0A1E3JPZ7 | A0A1J4KE73 | A0A815S408 | A0A0N5B7E3 | A0A8H6FY27 |
| A0A1E3HFS5 | A0A1J4JFK0 | A0A8B7DFA2 | A0A8S1XYM0 | A0A2L2U0T5 |
| A0A7S4GJ87 | Q4G2A0 | A0A5S6QR92 | Q3SD54 | A0A0B7JT57 |
| A0A1J4KAE0 | A0A5N6L5I0 | A0A7M5WXS5 | A0A8S1S9H3 | A0A167A5G7 |
| A0A8B7DFA2 | A0A821ZHC2 | A0A822FMM7 | A0A8S1U6V2 | A0A0A1UZN6 |
| A0A2G8LBI1 | A0A818DT02 | A0A8L2Q1F9 | A0A8S1UWG0 | A0A0B4HZP5 |
| A0A369RU39 | A0A821MGB8 | A0A1U7QV97 | A0A8S1WE94 | A0A7D5Z9U8 |
| A7RL89 | A0A817VQV9 | A0A3B3QXD3 | A0A8S1NHV9 | A0A0B4FV86 |
| V4C1F0 | A0A813V7X3 | A0A6S7GTF6 | A0A8S1NCU8 | A0A395N7P0 |
| A0A3M7SVY6 | A0A818EWQ6 | A0A2G8LBI1 | Q3SDM2 | E0S8K8 |
| A2E2M2 | A0A820XAV1 | A0A7M5X1D9 | A0A8S1Q3Y0 | I6UQ87 |
| A2G473 | A0A4W4EY14 | A0A1I8I446 | A0A8S1PM58 | I6ZJR5 |
| T1JME3 | A0A6P3WB05 | A0A1I8I5X4 | A0A8S1YEW8 | M1JID1 |
| D2W0R0 | A0A3B1JCF8 | A0A267GBT0 | I2H316 | A0A0B2UDK7 |
| A0A0F9ZEV9 | A0A3B4BP68 | A0A0F9ZEV9 | A0A0C2I723 | A0A8B8C0K3 |
| A0A182P478 | A0A7S4E9E6 | A0A3G4ZV93 | A0A7S3DFN8 | A0A3S1H8P1 |
| A0A6E8VSE3 | A0A7S3ZYW7 | A0A3G4ZT94 | A0A2R6XGH9 | A0A8C5MQN1 |
| A0A182VEW5 | F0YR38 | A0A1R2BEZ6 | A0A8S1RK63 | A7S215 |
| A0A182I2G0 | A0A397KZ26 | A7RL89 | A0A8S1UFZ0 | A0A293M7B3 |
| A0A182U3T0 | A0A8D9GR65 | A0A1J4JW69 | A0A8S1LD78 | G3AWI7 |
| A0A7S1EZY4 | A0A3P5ZWS8 | A2ESK3 | A0A8S1MMC2 | A0A0R3T291 |
| A0A7S3MGG9 | A0A6J0N9A9 | A0A1J4JBD1 | Q3SDN0 | TRINITY_DN6159_c0_g1_i1.p1 |
| A0A7L2NHC4 | V4MA47 | A2G2G1 | A0A8S1UJG9 | Q3SDI3 |
| A0A7L3M763 | A0A087GL06 | A2E8M0 | Q3SDK3 | Q4G295 |
| A0A7L3QA71 | A0A178V1R4 | A2GBJ0 | W7K002 | M7W6P6 |
| A0A7K5W7Z0 | A0A7G2F456 | A0A1J4JSU3 | A5K9N1 | A0A6V7TNB2 |
| A0A852EWV5 | A0A2Z4HJG9 | A0A1J4KCA8 | A0A1J1H3I3 | A0A6V7VDS8 |
| A0A7L0YXX8 | R9S9A9 | A0A146KAX7 | A0A8C9H9P9 | A0A1Y1ZC38 |
| A0A852G279 | A0A2Z4HJH7 | A2G412 | A0A6V7T708 | V5F1D1 |
| A0A7K5SL79 | A0A6D2IHV2 | A0A1J4KAE0 | A0A6G3MIX9 | A0A5C3PZW7 |
| A0A7K7MQT5 | G7J3R3 | TRINITY_DN13871_c2_g2_i1.p1 | L8H896 | A0A1Y1YUI7 |
| A0A7K4ULZ9 | A0A7J9C4M2 | A0A1J4JTG9 | A0A7R9XIL7 | A0A1Y1Z117 |
| A0A8C4A543 | A0A5C7HVA3 | A0A1J4JC27 | A0A3S0ZT37 | A0A8H3QPA1 |
| A0A2I4CSF6 | A0A6A3BGW3 | A0A1J4L0E6 | A0A8B7YSX1 | A0A2I1EWS0 |
| A0A3Q3EKQ8 | A0A5B6ZN47 | A2FI73 | | A0A8H3XAU5 |
| A0A3B4DB85 | L1JL95 | Q3SD49 | | A0A397TZM0 |
| A0A5N5Q526 | A0A135SN23 | Q3SD15 | | M2N699 |
| A0A673JC67 | A0A1Y1XHS8 | B9X1R2 | | A0A3N4JGL0 |
| A0A7J6B401 | A0A167NJP4 | A0A2H6K891 | | A0A1Y1I5H5 |
| W5UKR5 | A0A0G4IZJ1 | A0A061D9C8 | | A0A7J7M8Q8 |

A0A672L021
A0A8C2WLV0
A0A4W4EH74
A0A672QSA5
A0A8C9SBL6
V9LBN5
A0A452GH23
A0A8C4Y8Q3
A0A4D9DN93
A0A8C8RFF0
A0A6P7WJQ3
A0A6P7WQX3
A0A6P8P719
A0A8C5RET6
A0A098LYT8
A0A8C5WCM5
A0A821Q2A1
Q561P0
Q08AW6
I7MHF3
A0A4Z2DEI3
A0A7S3LSK1
A0A7S0VY16
D2VB98
A0A7S3D6C9
A0A7S3D4T3
A0A1J1HA99
A0A0D9QK24
A0A1Y3DQI3
K6UKL1
A0A1B1E0W5
A0A7S0GVJ0
A0A8J1ZLN5

F0WQ13
A0A1D2VNR0
A0A1E3QZZ2
A0A448YGF3
A0A875SBY1
A0A1E4SZ00
A0A1E4TMP5
C4R716
A7TQ91
A0A0N5AEB6
A0A0V0QP51
G0R2Q5
G0R2Z6
G0QZW6
Q23FM5
A0EC83
Q3SDR7
E9I3R6
G0QSP4
E1CB06
Q24CD0
A0A0V0QP25
G0QN99
B0ESL8
TRINITY_DN14379_c0_g1_i1.p1
A0A1J4L2Z7
Q4FID5
Q4G2B3
A0A1J4JDR8
A0A0V0QMI9
A2EI68
A0A146K818
K2HZF0
B0EJW3

J7S7W3
A0A7H9B7A8
A0A437AN12
A0A1J4KAP9
A0A1J4KF25
A2EV06
A0A6J8BC13
A0A2V1B0Y8
A0A2V1AFA3
A0A8F2RLY9
A0A8H7GXT7
C4YEB1
A0A0V1PV62
A0A812DZM0
A0A7R9M6F0
A0A836INF4
A0A448YEI3
A0A7L0DYT5
A0A091NRN7
A0A093J334
A0A7K6SFJ9
A0A850WD84
A0A7L0CYS0
A0A7L1JGW0
A0A7L1I795
A0A3B4B770
A0A8C6TDV0
A0A2P4QMJ6
A0A2N1ND68
A0A2Z6RC45
A0A372QVP2
A0A397SAP9
A0A1D1Y4X4
A0A1Y1ZC38
A0A8H7PPI3
A0A0B7N846

A0A0E9NEX6
I2JWX4
A0A7D9CYJ2
A0A875S066
A0A1E4S882
H0H091
J5RIJ4
A0A376B2F0
A0A0P1KTR4
C5DME6
A0A367YIS9
A0A367XW76
C5M895
A0A8J5BMJ0
A0A8J5BLS6
A0A3N4LLT7
A0A3N4IQI6
A0A6J3MGU2
A0A6A6FC18
A0A2I0RXR3
N1QIF5
A0A2G5ICM3
A0A6A7BSU4
A0A2K1QUJ9
A0A6A6JFC6
A0A063CBP1
A0A151GXK7
A0A0F7ZQD3
A0A2C5Z555
A0A8H4PTS8
T5A660
A0A179HZ11
A0A2S4LAA2

| UniProtKB ID used in Fig. S7S |
|---|
| H3ZPY0 |
| A0A7K9WWI1 |
| A0A7L2AY31 |
| A0A7L1SUJ0 |
| A0A672UB01 |
| A0A851Y506 |
| A0A7K4WN53 |
| A0A851F6L6 |
| A0A7K6AAI7 |
| A0A7K5K657 |
| A0A7L1YUW8 |
| A0A7L0J6I4 |
| A0A7K8YZE4 |
| A0A7K7HMY4 |
| A0A093PX51 |
| A0A7R5KUX4 |
| A0A7K6DW71 |
| A0A7K9ZLN4 |
| A0A7L1TJ62 |
| A0A852NNU4 |
| A0A091MQW9 |
| A0A7L0P0G0 |
| A0A091P6R5 |
| A0A7L4A6K9 |
| A0A663E6D2 |
| A0A8B9MXT3 |
| A0A7L0C2Q5 |
| A0A7K6SER2 |
| A0A7K6WP83 |
| A0A1V4JGU5 |
| A0A7K4S2M6 |
| A0A674KCU8 |
| A0A8C3HIH7 |
| M7BLR1 |
| A0A8C3XTW6 |
| A0A7K9M987 |
| A0A7K8PRT3 |
| A0A3L8SQL4 |
| W9XVC4 |
| I1HI21 |
| A0A2R5G1A2 |
| A0A665SW59 |
| A0A3B4XVT7 |
| A0A7C3SYP7 |
| A0A271M1F8 |
| A0A8H7VJ05 |
| A0A3Q4HSR0 |
| A0A8C4BIY7 |
| A0A7K7W9V8 |
| A0A851T1Y0 |
| B0WDG7 |
| G2WCA2 |
| A0A6A6A6U5 |
| Q9QZ84 |
| A0A2R8QAS3 |
| A0A2A3EN12 |

| UniProtKB ID used in Fig. S7T |
|---|
| A0A067CQT3 |
| A0A0A1TYQ4 |
| A0A0A1TZW0 |
| E3N8T3 |
| A0A0B2VJL4 |
| A0A183UYI0 |
| A0A3P7G8C6 |
| F1KY49 |
| A0A183E4B6 |
| A0A238C3C9 |
| A0A0K0F094 |
| A0A0N5B5P7 |
| A0A183UPP3 |
| A0A183VFL5 |
| A0A0F7FJ36 |
| A0A3G1A6P0 |
| A0A6G3LK14 |
| S6A5D1 |
| A0A7C1CET5 |
| A0A7C3RFL1 |
| A0A7C4NJF3 |
| A0A132NZ21 |
| C6LN26 |
| A0A1I5ZVK2 |
| A0A5P9FG66 |
| A0A7Z8LFA9 |
| A0A1Z5JLA5 |
| A0A1Z5K7K3 |
| A0A497KTA2 |
| A0A497N359 |
| A0A497NL17 |
| A0A523VQX2 |
| A0A550GFH8 |
| A0A550GJA5 |
| A0A7C3U715 |
| A0A832GST8 |
| A0A4P2VDB4 |
| A0A7J2VYQ3 |
| A0A7C4BDQ5 |
| A0A7J3Z5U8 |
| A0A7M1UPU6 |
| A0A7J3SKM9 |
| A0A7J2ZGM5 |
| A0A7L4RFL7 |
| A0A1Q9MXI9 |
| A0A4E0PVR7 |
| A0A7D5INX6 |
| A2G4Y4 |
| A0A1J4J6C0 |
| A0A1J4JHP7 |
| A0A1J4K2U0 |
| A0A1J4L0B3 |
| A0A0D8XU78 |
| A0A183BZJ7 |
| A0A4U5MK84 |
| A0A6P8JNP9 |
| B4P8D7 |

| UniProtKB ID used in Fig. S7U |
|---|
| A0A0A1U419 |
| A0A0J7NLU2 |
| A0A151WVR9 |
| A0A195FX94 |
| A0A836G5Y4 |
| A0A836GA01 |
| A0A2A3EPC7 |
| A0A7M7FY08 |
| A0A6J3L2F3 |
| A0A6P3UV69 |
| A0A6P3UDD3 |
| A0A6I9W1B6 |
| E2AVV6 |
| A0A146KMR0 |
| A0A1J4KS07 |
| A0D4Q1 |
| A0A5J5BR95 |

| UniProtKB ID used in Fig. S7V |
|---|
| A0A061IXG7 |
| A0A081B323 |
| W3A4R2 |
| W2RI07 |
| W2XW22 |
| W2JU56 |
| V9G0E8 |
| W2HPW2 |
| W2LZU9 |
| A0A0W8CHQ3 |
| A0A0W8D7D9 |
| A0A329RS92 |
| A0A225WRX5 |
| A0A6A4AKH4 |
| A0A6G0SI28 |
| G4ZAV4 |
| A0A485L1L2 |
| W4HCB8 |
| A0A1E7ERA3 |
| A0A448Z563 |
| A0A1Z5JT78 |
| A0A1Z5KGE1 |
| A0A8J9X7B7 |
| A0A6U6EFP9 |
| A0A7S4IG93 |
| K0TD91 |
| A0A6H5KS84 |
| A0A6V1NP81 |
| A0A196SCC1 |
| A0A196SHI8 |
| A0A0L0DGH7 |
| A0A0L0DSS8 |
| A0A0S4IYE4 |
| A0A7S1KZ37 |
| A0A7S1J1G0 |
| A0A7S4GF14 |
| A0A0B7A4Z5 |
| A0A7I8VYK0 |
| A0A132AE08 |
| A0A834R0H4 |
| A0A6F9DH37 |
| A0A1D1VY25 |
| A0A7M7JD61 |
| A0A7M7JHA0 |
| A0A7M7JDD3 |
| A0A0A1TZW0 |
| A0A644F932 |
| A8B5G7 |
| E1F524 |
| A0A6G3LK14 |
| S6A5D1 |
| A2FGA9 |
| A0A1J4J0K8 |
| A0A226ED05 |
| A0A087HKP6 |
| A0A087HKP7 |
| A0A178WIX8 |

| UniProtKB ID used in Fig. S7W |
|---|
| A0A0C3DNR7 |
| A0A139GX96 |
| A0A139HDY9 |
| A0A139HE38 |
| A0A8H6R9V0 |
| M3B473 |
| N1PPB3 |
| A0A1Y6LCR3 |
| F9X5K1 |
| A0A2H1G483 |
| A0A1C5XZZ2 |
| A0A2D0TBD0 |
| A0A3B4BG89 |
| A0A2S7QNH0 |
| W9CRX6 |
| A0A292QA45 |
| A0A2T7A7X9 |
| A0A8H6CJU6 |

B4I7X4
B4QGM3
Q7KVM5
O77294
Q8SXA5
A0A7S0AJV3
A0A1J4K7E9
A0A132NU74
E1F400
A8B747
A0A4Z1SM88
A0A132NVA3
C6LUS8
A8B748
V6TNS6
E1F401
A0A0R3WSU4
A0A5D2UNJ7
A0A0V0Q868
A0A1D3CZ57
A0A6P6RYW6
A0A212EII7
A0A2H1X0P5
A0A6J1X8S4
E4X4C7
E4YI80
Q5C227
A0A7S1VIC8
A0A7J7IEY4
A0A7S4CJD1
A0A7S1TPF1
A0A7S2RK65
B8BY29
T0RUL9

A0A5S9SDH6
A0A7G2DQD3
A0A1P8AQ82
A0A1J4MQL9
B6AEM6
A0A0N4UH75
A0A6A5DL12
A0A8C1JXR9
A0A834CN59
A0A409WJC1
A0A5N6DP27
A0A6A6JG40
A0A448WL55
A0A6C0ECQ0
A0A3P7LUJ7
A0A6G0YTM2
A0A1E5R165
A0A2P2I617
A0A6A7G8M1
A0A6Q2XRZ8
A0A4Z2DR72
A0A7S1SFW7
A0A812SLM4
A0A812Y9I1
A0A812SUV1
A0A813D987
A0A813G871
A0A813E6D2
A0A813ES76
A0A813ILK3
A0A813L8A9
A0A813JKW0
A0A813ENB4
A0A813G2P3
A0A7S2QNK6
A0A7S4DZM0
A0A7S1M792
A0A1G4I817
Q583K6
A0A3L6LAG3

| UniProtKB ID used in Fig. S7X | UniProtKB ID used in Fig. S7Y | UniProtKB ID used in Fig. S7Z | UniProtKB ID used in Fig. S7AA | UniProtKB ID used in Fig. S7AB |
|---|---|---|---|---|
| A0A1J4J8K4 | A0A1R2B755 | A0A7S4MA53 | A0A381WSD4 | A0A0F9WHB2 |
| I7LTW8 | A0A6V7RZD1 | A0A6A5PKL5 | A2EJI5 | A0A0V0R280 |
| G0QLE1 | A0A6V7TAY8 | A0A5N5MCT4 | A0A1J4KK47 | A0A1J4J277 |
| A0A1J4KI54 | W7AKE4 | A0A4D9BTN4 | A0A1J4J5X7 | A0A1J4JS52 |
| A2G473 | A0A6V7SJJ4 | A0A445AB95 | A0A1J4JM42 | A0A1J4JKF4 |
| B0ESL8 | A0A449BX12 | A0A444ZRU0 | A0A1J4J1W8 | A2DDL1 |
| A2E2M2 | V7PWG1 | A0A445AAU9 | A2DPH2 | A2DR69 |
| A0A3Q4HJF3 | A0A078KAX5 | A0A444ZRR5 | A0A1J4JF19 | A2DLG2 |
| A0A668UPV9 | A0A1C6YJ52 | A0A5N5P234+A500 | A0A1J4L0F6 | A2DXR8 |
| A0A3Q3KP92 | A0A1C6YPX8 | A0A2P5XWQ2 | A0A1J4K3E3 | A2EGJ2 |
| A0A6A5ED41 | A0A2C9JIF3 | A0A1D6PJC2 | A2DYG6 | A2FY69 |
| A0A484D204 | A0A210R7P5 | A0A818JZY3 | A0A1J4K1G9 | A2DWP7 |
| A0A8C9ZDF8 | A0A816GMA8 | A0A8J5FVM1 | A0A1J4KPC6 | A2DL53 |
| A0A8C9Z5M5 | A0A814NQP4 | A0A445FSN0 | A0A1J4K5B7 | A2FUI1 |
| A0A3Q4BUT6 | A0A814LCP9 | A0A445DP98 | A0A1J4KEE6 | A2E117 |
| A0A8C4Z511 | A0A819UDZ7 | I7MER0 | A2FXH1 | A2FID7 |
| A0A8C5ADE2 | A0A6F9DMZ3 | A0A7J9JW60 | A0A1J4JPI8 | A2G0M4 |
| A0A8C5AGK9 | A0A0V0QUS7 | A0A7J9HIX1 | A2FK80 | A2E9J2 |
| A0A8C4Z911 | G8YJ57 | A0A7J9M609 | A0A1J4KY41 | A2FD09 |
| W5NJL9 | A0A067NBL4 | A0A7J9CF27 | A2FZQ5 | A2GJC5 |
| W5NJL8 | B0I556 | A0A7J8VF00 | A2DAL7 | A0A1J4K281 |
| A0A8C4AG32 | V5N917 | A0A7J6G4Z6 | A0A1J4JQ46 | A0A1J4K7T5 |
| A0A3N0XD70 | A0A804JWC8 | A0A8T2XKP9 | A0A1J4JLD7 | A2EBM3 |
| E9QB21 | A0A2P5C898 | A0A2N9GR34 | A0A1J4JN56 | A0A1J4KDC4 |
| A0A3M7Q9T4 | A0A2P5E5Y7 | A0A5J4Y1E9 | A0A1J4KLH2 | A0A1J4KI15 |
| V6LYP9 | A0A6D2JGI6 | A0A8C8CMB4 | A0A1J4J164 | A0A1J4K8I1 |
| A0A7R9M589 | A0A803ME03 | A0A8J2ASF7 | M7WIA0 | A2D981 |
| A0A5A9NME9 | A0A6A4QAE7 | B4R2C4 | B0E6U0 | A2DF81 |
| A0A7J5ZS88 | A0A422NME5 | S7W5E9 | K2H7X2 | A2EW46 |
| A0A3Q2ZTK7 | A0A163J3W0 | A2DEX3 | A0A0A1TV77 | A2FIY3 |
| A0A3P8T5G6 | T1EJ40 | A0A1J4JPT4 | A0A1R2AUF7 | A0A1J4J7M9 |
| A0A3P8T5E3 | A0A8S1L2W1 | A0A1J4KJ20 | G0QKQ4 | A0A1J4K7D8 |
| A0A2I4B0H1 | A0A8S1W812 | B0EGV2 | A0A1R2CZW2 | A2F8D7 |
| M3ZM01 | A0A8S1S0L3 | F2TYG9 | G0QN96 | A2G1U5 |
| A0A6J2QQH5 | A0A0A1U0D2 | A0A7S2WM62 | A0C3W5 | A0A1J4JE48 |
| A0A3Q3M723 | A0A838DXL1 | A0A1F5S4L6 | A0BF11 | A0A1J4JJL8 |
| A0A3B3ZC47 | A0A7C5I534 | A0A8J7XL72 | A0A1J4JTG1 | A0A1J4K5N1 |
| A0A3Q3JP80 | T2B550 | A0A1J4K4Q3 | A0A1J4KXC9 | A0A1J4KNE3 |
| A0A3Q2YB76 | M1CE33 | A0A1J4KAS0 | A0A1J4J3Z3 | A2DCQ9 |
| A0A669F9J7 | A0A4D9C2P2 | A0A1J4K230 | A0A2T6ZXE1 | A2G5L9 |
| A0A3Q0T011 | A0A0S4J575 | A0A1J4JBY4 | A0A5J5ERG7 | A0A1J4KXJ9 |
| A0A4W4FBX1 | A0A077Z461 | A0A1J4KBM0 | A0A3N4K202 | A0A1J4L356 |
| A0A803XWF9 | A0A085MFE0 | A0A1J4KMK6 | A0A292PVS3 | A0A1J4J312 |
| A0A6J2IEX0 | A0A5S6QQ87 | A0A1J4KGG0 | A0A317SEH9 | A0A1J4JCF0 |
| A0A218UNS0 | A0A443SWH3 | A0A448ZIF0 | D5G9K3 | A0A1J4L119 |
| A0A7K5PIN4 | A0A074YCB5 | A0A7S3LE49 | A0A3N4LVM4 | A0A1J4KHE9 |
| A0A1J4KQB1 | A0A074W7A7 | A0A7S0CDE3 | A0A0D6EJH7 | A0A1J4JK68 |
| A0A4U5PI34 | A0A074X5N7 | A0A072VBW5 | A0A2S5B088 | A2DBC8 |
| A0A6N1NUY3 | A0A074XB43 | A2Q450 | A0A120E829 | A2DCX2 |
| A0A6N1NW88 | A0A4T0A872 | A0A072VCK4 | A0A4U0W4G1 | A2ED22 |
| R0MLC0 | A0A4V6TCA5 | A0A2J6K9H7 | A0A0V0QPL4 | A2G6V3 |
| M1K5Y5 | M3IHX8 | A0A6S7MHE8 | A0A814EWS9 | A2FMR7 |
| Q8SW02 | G3AUZ8 | A0A6J1FXY8 | A2DFV7 | A0A1J4JRT3 |
| A0A0B2UM37 | A0A8J5Q0S6 | A0A6J1JF26 | A0A6B2KXK6 | A0A1J4KLZ2 |
| A0A0V0QP62 | A3LSM4 | A0A437APR0 | A0A6B2L0G7 | A0A1J4JBK5 |
| A0A0C3LPW8 | A0A0V1PW92 | A0A1J4KCS6 | A0A3P8XE13 | A0A1J4KMA6 |
| A0A0C2XQX9 | Q6BTM5 | A0A0A1U6Y8 | A0A0V0QQU1 | A0A1J4KEA6 |
| G4TJQ4 | G8JUE0 | A0A197K3Z8 | J9IDQ8 | A0A1J4K5M4 |

A0A8H3TZN1
V6U403
V6TMU8
Q9GU79
E1F2X5
A0A484E0L2
I7LSW1
Q231B4
A0A8H7S4J3
A0A0C2I723
D3BMH1
A0A7J7JHF2
M7WFW0
A0A1J4JAM7
A0A875SBY1
A0A1R2BA89
A0A1R2B1V3
A0A6A5BCH3
A0A8B8D3Y5
Q23W08
A0A1E4RM66
A0A1Z8JJK0
A0A1V2LQ27
A0A099P792
A0A1J4K3I2
A2DJT9
A0A1J4J8B6

A0A1R2BZJ2

A0A1Y1VGQ9
A0A1Y1VWN3
A0A1Y2ECC1
A0A0B1T0A4
A0A2X0LIH5
A0A2X0M815
A0A1U7LN28
A0A1R1Y2Z4
A0A815BAF5
A0A7L2X781
A0A0R3RIG7
A0A0D8Y5W0
A0A811LNT4
A0A6P9ECR0
A0A6P5NDZ8
A0A8J1ZKR6
A0A0D9QGL2
A0A1D3LGY4
A0A4V0K4K9
A0A1C6XER6
W7AM38
A0A6V7TEC2
A0A6V7RWG2
A0A6V7RXG6
A0A081IBW3
A0A509AI23
Q7RLG9
A0A077Y4Y0
V7PR35
A0A1A8YM92

J9ID37
A0A7W0JEE4
A0A4V4NF50
A0A0D9W5P5
A0A8J5WY44
A0A8J5W3R3
A0A6A6KYJ4
A0A811MS40
A0A061R1F1
A0A7S1T3L6
A0A4T0S531
A0A4T0UDE2
R9ARQ6
A0A4T0HL96
A0A4T0ILR3
A0A5B8MQ60
A0A7S3C6J7
A0A5J5AP43
A0A0K9RIK7
A0A6V7QVD3
A0A830CG88
A0A2G9G885
B6T5R0
A0A7J6CIS0
A0A1J1J6K2
A0A1I7SA30
A0A6V7HSW2
A0A132A5A7
A0A834VEN9
A0A8J2ED74
A0A3G4ZN36
A0A3G4ZZP1
A0A6C0J6C9
A0A6C0HVX1
A0A6C0H7D4
A0A6C0DB04
A0A381RMA6

A0A1J4KJQ5
A2G7D0
A2FHT0
A2FKU7
A0A1J4JRP0
A2ELG8
A0A1J4J998
A0A1J4K0I5
A2DB42
A2E5X3
A0A1J4JAI1
A0A1J4L3X5
A0A6G8MVV9
A0A5K1VLT1
B0ETD3
A2DWP6
A2F3W9
A0A1R2CTL5
A0A397IU75
A0A7S0ZLB8
A0A7S0ZLF1
A0A8S4GIZ7
A0A8S4GPI8
A0A1E5W9P7
A0A2U1L2J1
A0A2I0VIP8
A0A8T3A9R6
A0A371H1L5
A0A438CWV7
A0A438GQK5
E0CUH5
A0A8K0I3U2
A0A6J0JNJ0
A0A2G9HU10
J9HWH8
R0MQ89

| UniProtKB ID used in Fig. S7AC | UniProtKB ID used in Fig. S7AD |
|---|---|
| A0A438FEI1 | R7UH49 |
| A0A1J4K5M4 | A2DY61 |
| A0A1J4JLD9 | F6V890 |
| A0A1J4KPE8 | A0A6F9DLG8 |
| A0A1J4JSY7 | A0A8J4XSX2 |
| A2DK36 | B3RJY9 |
| A2DAT0 | T1FQZ0 |
| A2G2A8 | A0A7I8W6Z3 |
| A2FID7 | A0A6S7GDS7 |
| A2E117 | A0A7J7A2M3 |
| A2FBW2 | A0CGY4 |
| A2DIR4 | A2EIC5 |
| A2G1U7 | A7RH03 |
| A2EDQ7 | A0A8B8D6Q7 |
| A0A1J4JSA9 | A0A158QB13 |
| A2DF81 | A0A0N5CYC8 |
| A0A1J4K5N1 | E0VVH1 |
| A0A1J4KA14 | A0A8J8SVZ6 |
| A0A1J4JF52 | A0A834XPU3 |
| A0A1J4KHJ5 | A0A813PEX5 |
| A0A1J4JE48 | A0A1R2C949 |
| A0A1J4KA25 | A0A1R2AP99 |
| A0A1J4JJL8 | A0A1R2B613 |
| A0A1J4KRD7 | F0GWI9 |
| A2E5X3 | A0A133KEI5 |
| A2G2L1 | A0A1M5QV77 |
| A0A1J4L3X5 | R5IVQ1 |
| A0A1J4J3H2 | A0A358M8C2 |
| A0A0V0R280 | A0A3B9I2F9 |
| A0A0V0QS39 | A0A1M6DI04 |
| A0A2U1L2J1 | A0A1U7M316 |
| G0WAQ3 | A0A7V6XKZ4 |
| A0A1R2C6E8 | A0A285BKP8 |
| A0A1R2C3M4 | B7R9Z4 |
| A0A1R2BL46 | A0A7Y2L5X5 |
| A0A1R2C1E0 | A0A7C6RD96 |
| A0A1R2C1D5 | A0A1G7U587 |
| A0A1R2D3G5 | A0A1M4UKH6 |
| A0A1R2BD78 | I9KVI9 |
| A0A7I8VN78 | M8CSL9 |
| A0A8J8NGV7 | A0A0B3BPJ8 |
| A0A1R2BH76 | A0A5D8QG83 |
| A0A1R2CIQ1 | A0A269XM42 |
| A0A1R2BD27 | A0A8J7S2V1 |
| F4Q2J2 | A0A660Q0G1 |
| A0A151ZFK7 | A0A3M1C7C4 |
| A0A7J7J830 | A0A660Z1I5 |
| A0A7N4PR21 | A0A523NCT1 |
| A0A0V0R1R4 | A0A7X8FA76 |
| A0A5N5JUD3 | A0A832LCD7 |
| A0A0M0JLC1 | A0A3M1N4D4 |
| A0A7S4B0W8 | G8BTZ4 |
| J9D5Z6 | G0WCG8 |
| R0KXU5 | G0VAK7 |
| R0MQ89 | A0A8H2ZI03 |
| A0A6G0XI09 | A0A1X7R7E2 |
| A0A8E4LYS0 | A0A7G3ZJE4 |
| A0A8E4LYR5 | A0A7H9HV14 |

| | |
|---|---|
| A0A4Y7LEK7 | A0A1G4KAP7 |
| A0A4Y7L3K0 | F2QLM7 |
| O65366 | C4QXS1 |
| K9LWN6 | A0A1E4S1A3 |
| A0A6P5YTN1 | A0A0H5CIT2 |
| A0A6P5Z1E2 | A0A1E3NZD5 |
| A0A6J1CPT5 | A0A1E3QYY1 |
| A0A7N2L062 | R4XNV4 |
| A0A835F539 | A0A1Y2ES38 |
| A0A251UI60 | A0A1Y2DDV0 |
| A0A834T595 | A0A1Y1VN91 |
| A0A314LBJ9 | A0A1Y1X5H4 |
| A0A836DDK2 | R7QSN9 |
| A0A3P6A873 | A2EAS8 |
| R0GVT9 | G0QUM5 |
| A0A1S2XDH5 | F0Y2N0 |
| A0A6D2IRL1 | A0A8J4V512 |
| A0A1J3H1Z5 | V4B415 |
| A0A654G3C9 | A0A1J4L2Q9 |
| A0A178UCW3 | A0A183KEX8 |
| A0A2Z7AIN2 | A0A183NTI0 |
| A0A3N6SDE0 | W7WYR6 |
| A0A816QU65 | A0A7S2U6B1 |
| A0A0D3D1R9 | A0DG76 |
| A0A816ZHN7 | A0A7F5R7G2 |
| M4F200 | A0A7F5R7F1 |
| A0A067KW19 | A0A443QQT7 |
| Q8L5N8 | A0A8H3X323 |
| A0A200PW04 | A0A835YZ60 |
| A0A2Z0DYH0 | A0DJR9 |
| A0A2U1NB55 | A0A1R2AUS4 |
| W9R8B8 | J9IEM3 |
| A0A4Y7KVZ2 | A0BIS4 |
| A0A371ERQ2 | A0A8J4PVQ4 |
| A0A1S3TIW5 | A0A1J4K477 |
| A0A314KYX7 | A0A1R2B8U0 |
| A0A2C9UV00 | A0A1R2C538 |
| A0A438BQK6 | |
| A0A438GGA2 | |

* IDs starting with 'TRINITY' are sequences from Nishimura et al. (2020)

**Table S6. Infection rates of clostridial 16S rRNA phylotypes to cells of respective protist hosts.**

| 16S rRNA phylotype | termite colony ID | infection rate (%) |
|---|---|---|
| RsTa-C01 | A | 0 |
| (host protist: | B | 0 |
| *Trichonymph aagilis* ) | C | 0 |
| | D | 9.8 |
| | E | 54.7 |
| | F | 0.9 |
| | G | 71.4 |
| CfP3-15 | A * | 0 |
| (host protist: | B * | 0 |
| *Pseudotrichonympha* | C * | 0 |
| grassii) | D * | 0 |
| | E * | 0 |
| | F ** | 74.1 |
| | G ** | 100.0 |
| | H ** | 21.0 |
| | I *** | 4.0 |
| | J *** | 0.9 |
| NkDv07 | A | 0 |
| (host protist: | B | 0 |
| *Devescovina* sp.) | C | 0 |
| | D | 0 |
| | E | 0 |
| | F | 0 |
| | G | 0 |
| | H | 0 |
| | I | 0 |
| | J | 0 |
| | K | 18.3 |
| | L | 0 |
| | M | 0 |
| | N | 7.2 |

* From Kagoshima Prefecture.

** From Okinawa Prefecture.

*** From Chiba Prefecture.

**Table S7. Single-copy marker genes modified based on the CfP3-15 genome.**

| domain ID | gene annotation |
|---|---|
| PF00623 | RNA polymerase Rpb1, domain 2 |
| PF00889 | Elongation factor TS |
| PF00886 | Ribosomal protein S16 |
| PF11987 | Translation-initiation factor 2 |
| TIGR03723 | tRNA threonylcarbamoyl adenosine protein |
| PF01746 | tRNA (Guanine-1)-methyltransferase |
| PF00318 | Ribosomal protein S2 |
| PF01765 | Ribosome recycling factor |
| PF00162 | Phosphoglycerate kinase |
| PF00177 | Ribosomal protein S7p/S5e |
| PF03947 | Ribosomal Proteins L2, C-terminal domain |
| PF00673 | ribosomal L5P family C-terminus |
| PF00203 | Ribosomal protein S19 |
| TIGR03263 | Guanylate kinase |
| PF01195 | Peptidyl-tRNA hydrolase |
| PF06421 | GTP-binding protein LepA C-terminus |
| PF00380 | Ribosomal protein S9/S16 |
| PF04998 | RNA polymerase Rpb1, domain 5 |
| PF00252 | Ribosomal protein L16p/L10e |
| PF00453 | Ribosomal protein L20 |
| TIGR02432 | tRNA(Ile)-lysidine synthetase |
| PF00333 | Ribosomal protein S5, N-terminal domain |
| PF01245 | Ribosomal protein L19 |
| PF00312 | Ribosomal protein S15 |
| PF00189 | Ribosomal protein S3, C-terminal domain |
| PF01250 | Ribosomal protein S6 |
| PF00562 | RNA polymerase Rpb2, domain 6 |
| PF01281 | Ribosomal protein L9, N-terminal domain |
| PF01000 | RNA polymerase Rpb3/RpoA insert domain |
| PF02130 | Uncharacterized protein family UPF0054 |
| PF03948 | Ribosomal protein L9, C-terminal domain |
| PF00281 | Ribosomal protein L5 |
| PF00828 | Ribosomal protein L18e/L15 |
| PF13184 | NusA-like KH domain |
| PF02978 | Signal peptide binding domain |
| PF01795 | MraW methylase family |
| TIGR00755 | rRNA small subunit methyltransferase A |
| PF01016 | Ribosomal L27 protein |
| PF01196 | Ribosomal protein L17 |
| TIGR01079 | Ribosomal protein L24 |
| TIGR00855 | Ribosomal protein L7/L12 |
| PF02033 | Ribosome-binding factor A |
| PF01668 | SmpB protein |

| | |
|---|---|
| TIGR03594 | Ribosome-associated GTPase EngA |
| PF00572 | Ribosomal protein L13 |
| PF04983 | RNA polymerase Rpb1, domain 3 |
| PF00466 | Ribosomal protein L10 |
| PF06071 | Protein of unknown function (DUF933) |
| PF00416 | Ribosomal protein S13/S18 |
| PF00347 | Ribosomal protein L6 |
| PF00831 | Ribosomal L29 protein |
| PF00338 | Ribosomal protein S10p/S20e |
| TIGR00967 | Preprotein translocase, SecY subunit |
| PF04565 | RNA polymerase Rpb2, domain 3 |
| PF03719 | Ribosomal protein S5, C-terminal domain |
| TIGR00250 | RNAse H domain protein, YqgF family |
| PF00410 | Ribosomal protein S8 |
| PF04997 | RNA polymerase Rpb1, domain 1 |
| PF03484 | tRNA synthetase B5 domain |
| PF00297 | Ribosomal protein L3 |
| PF05697 | Bacterial trigger factor protein (TF) |
| PF00181 | Ribosomal Proteins L2, RNA binding domain |
| TIGR00344 | Alanine--tRNA ligase |
| PF01632 | Ribosomal protein L35 |
| PF05000 | RNA polymerase Rpb1, domain 4 |
| PF00573 | Ribosomal protein L4/L1 family |
| TIGR00810 | Preprotein translocase, SecG subunit |
| TIGR02075 | UMP kinase |
| PF00298 | Ribosomal protein L11, RNA binding domain |
| TIGR00459 | Aspartate--tRNA ligase |
| PF00411 | Ribosomal protein S11 |
| PF00164 | Ribosomal protein S12/S23 |
| PF04561 | RNA polymerase Rpb2, domain 2 |
| TIGR00392 | Isoleucine--tRNA ligase |
| PF00238 | Ribosomal protein L14p/L23e |
| TIGR00460 | Methionyl-tRNA formyltransferase |
| PF00861 | Ribosomal L18p/L5e family |
| TIGR00019 | Peptide chain release factor 1 |
| PF01193 | RNA polymerase Rpb3/Rpb11 domain |
| PF10385 | RNA polymerase beta subunit external 1 domain |
| PF00276 | Ribosomal protein L23 |
| PF08529 | NusA N-terminal domain |
| TIGR00329 | Metallohydrolase, glycoprotease/Kae1 family |
| PF00829 | Ribosomal prokaryotic L21 protein |
| TIGR00922 | Transcription termination factor NusG |
| PF03946 | Ribosomal protein L11, N-terminal domain |
| PF00237 | Ribosomal protein L22p/L17e |
| PF00366 | Ribosomal protein S17 |

| | |
|---|---|
| PF04563 | RNA polymerase beta subunit |
| PF01409 | tRNA synthetases class II core domain (F) |
| PF04560 | RNA polymerase Rpb2, domain 7 |
| PF00687 | Ribosomal protein L1p/L10e family |

**Table S8: MAGs (>50% CheckM or CheckM2 completeness) detected in WGA samples.**

| WGA sample | bin ID | classification* | total length (bp) | G+C (%) | CheckM** completeness (%) | CheckM2*** completeness (%) | alignment rate (%) |
|---|---|---|---|---|---|---|---|
| *P. grassii* | A | *Termitinemataceae* | 2,383,868 | 45.2 | 78.5 | 80.7 | 9.3 |
| | B | "*Ca* . Azobacteroides pseudotrichonymphae" | 1,290,540 | 32.8 | 100 | 100.0 | 57.1 |
| | C | *Treponemataceae* | 2,245,192 | 51.1 | 48 | 51.3 | 5.0 |
| | CfP3-15# | "Acutalibacteraceae" | 1,007,634 | 31.4 | 80.2 | 86.8 | 29.6 |
| *T. agilis* | A | *Treponemataceae* | 3,261,309 | 48.4 | 89.9 | 81.7 | 1.8 |
| | B | *Termitinemataceae* | 3,602,372 | 40.5 | 87.1 | 95.5 | 5.3 |
| | C | "*Ca.* Endomicrobium trichonymphae" | 1,131,688 | 35.2 | 96.6 | 100.0 | 51.4 |
| | D | *Rickettsiales* | 1,843,874 | 28.3 | 82.3 | 74.8 | 1.4 |
| | E | *Mycoplasmoidaceae* | 1,376,671 | 29.6 | 76.4 | 80.3 | 1.0 |
| | F | *Termitinemataceae* | 2,345,787 | 39.5 | 81.2 | 82.0 | 2.9 |
| | G | *Termitinemataceae* | 2,599,803 | 42.0 | 83.2 | 87.6 | 3.4 |
| | RsTa-C01# | "Acutalibacteraceae" | 1,276,969 | 27.5 | 85.3 | 93.9 | 14.0 |
| *Devescovina* | A | *Endomicrobium* | 885,870 | 31.7 | 86.2 | 90.4 | 13.7 |
| | NkDv07# | "Acutalibacteraceae" | 1,079,858 | 33.0 | 85.3 | 90.6 | 74.6 |

* Based on GTDB r207.

** Estimated completeness, using the universal set of bacterial marker genes.

*** Estimated completeness, using "General Model".

### Bins were after individually processed (see Supplementary Methods); therefore, CfP3-15 and RsTaC01 were complete geonomes.

**Table S9. Pseudogenes of endosymbiotic *Clostridia*.**

| ID [*] | gene annotation based on COG |
|---|---|
| CfP3-15_0011 | Beta-lactamase class A |
| CfP3-15_0035 | Transposase |
| CfP3-15_0259 | Transposase |
| CfP3-15_0279 | Transposase |
| CfP3-15_0414 | Transposase |
| CfP3-15_0447 | Transposase |
| CfP3-15_0547 | Transposase |
| CfP3-15_0567 | Transposase |
| CfP3-15_0066 | Phage-related protein |
| CfP3-15_0068 | Phage-related protein |
| CfP3-15_0082 | Transposase and inactivated derivatives, IS1 family |
| CfP3-15_0271 | Transposase and inactivated derivatives, IS1 family |
| CfP3-15_0458 | Transposase and inactivated derivatives, IS1 family |
| CfP3-15_0487 | Transposase and inactivated derivatives, IS1 family |
| CfP3-15_0594 | Transposase and inactivated derivatives, IS1 family |
| CfP3-15_0595 | Transposase and inactivated derivatives, IS1 family |
| CfP3-15_0645 | Transposase and inactivated derivatives, IS1 family |
| CfP3-15_0767 | Transposase and inactivated derivatives, IS1 family |
| CfP3-15_0925 | Transposase and inactivated derivatives, IS1 family |
| CfP3-15_0270 | ATP-dependent Lon protease, bacterial type |
| CfP3-15_0286 | DNA uptake channel protein ComEC, N-terminal domain |
| CfP3-15_0296 | DNA-binding transcriptional regulator, PucR/PutR family |
| CfP3-15_0306 | Signal recognition particle GTPase |
| CfP3-15_0312 | DNA repair protein RadA/Sms, contains AAA+ ATPase domain |
| CfP3-15_0658 | DNA repair protein RadA/Sms, contains AAA+ ATPase domain |
| CfP3-15_0388 | Transposase InsA |
| CfP3-15_0417 | DNA replication protein DnaC |
| CfP3-15_0517 | DNA replication protein DnaC |
| CfP3-15_0774 | DNA replication protein DnaC |
| CfP3-15_0623 | ATP-dependent DNA ligase |
| CfP3-15_0773 | ATP-dependent DNA ligase |
| CfP3-15_0626 | Holliday junction resolvase RusA (prophage-encoded endonuclease) |
| CfP3-15_0639 | PASTA domain, binds beta-lactams |
| CfP3-15_0687 | GTPase SAR1 family domain |
| NkDv07_0149 | 16S rRNA G527 N7-methylase RsmG (former glucose-inhibited division protein B) |
| NkDv07_0017 | Predicted transcriptional regulator, contains HTH domain |

| | |
|---|---|
| NkDv07_0247 | Fatty acid-binding protein DegV (function unknown) |
| NkDv07_0344 | DNA repair protein RadA/Sms, contains AAA+ ATPase domain |
| NkDv07_0369 | DNA repair protein RadA/Sms, contains AAA+ ATPase domain |
| NkDv07_0390 | Transposase |
| NkDv07_0414 | Transposase |
| NkDv07_0415 | Transposase |
| NkDv07_0439 | Transposase |
| NkDv07_0455 | Transposase |
| NkDv07_0480 | Transposase |
| NkDv07_0500 | Transposase |
| NkDv07_0545 | Transposase |
| NkDv07_0584 | Transposase |
| NkDv07_0611 | Transposase |
| NkDv07_0786 | Transposase |
| NkDv07_0795 | Transposase |
| NkDv07_0798 | Transposase |
| NkDv07_0799 | Transposase |
| NkDv07_0824 | Transposase |
| NkDv07_0967 | Transposase |
| NkDv07_0982 | Transposase |
| NkDv07_0418 | DNA replication protein DnaC |
| NkDv07_0451 | DNA replication protein DnaC |
| NkDv07_0423 | Uncharacterized protein domain, C-terminal to COG1479 DNAse/DNA nickase |
| NkDv07_0431 | 2-polyprenyl-3-methyl-5-hydroxy-6-metoxy-1,4-benzoquinol methylase |
| NkDv07_0435 | Transposase |
| NkDv07_0457 | Holliday junction resolvase RusA (prophage-encoded endonuclease) |
| NkDv07_0463 | 5-methylcytosine-specific restriction endonuclease McrA |
| NkDv07_0789 | 5-methylcytosine-specific restriction endonuclease McrA |
| NkDv07_0465 | ATP-dependent DNA ligase |
| NkDv07_0791 | ATP-dependent DNA ligase |
| NkDv07_0467 | Lipopolysaccharide biosynthesis protein, LPS:glycosyltransferase |
| NkDv07_0470 | Phage-related protein |
| NkDv07_0474 | Site-specific DNA recombinase SpoIVCA/DNA invertase PinE |
| NkDv07_0493 | Site-specific DNA recombinase SpoIVCA/DNA invertase PinE |
| NkDv07_0531 | Site-specific DNA recombinase SpoIVCA/DNA invertase PinE |
| NkDv07_0475 | DNA-directed RNA polymerase specialized sigma subunit, sigma24 family |
| NkDv07_0591 | DNA-directed RNA polymerase specialized sigma subunit, sigma24 family |
| NkDv07_0476 | DNA modification methylase |
| NkDv07_0589 | DNA modification methylase |

| | |
|---|---|
| NkDv07_0496 | Phage antirepressor protein YoqD, KilAC domain |
| NkDv07_0528 | Phage antirepressor protein YoqD, KilAC domain |
| NkDv07_0498 | Lyzozyme M1 (1,4-beta-N-acetylmuramidase), GH25 family |
| NkDv07_0526 | Lyzozyme M1 (1,4-beta-N-acetylmuramidase), GH25 family |
| NkDv07_0577 | Transposase and inactivated derivatives, IS1 family |
| NkDv07_0612 | Transposase (or an inactivated derivative) |
| NkDv07_0775 | Transposase (or an inactivated derivative) |
| NkDv07_0793 | Phage portal protein BeeE |
| NkDv07_0794 | Predicted phage phi-C31 gp36 major capsid-like protein |
| NkDv07_0952 | Signal transduction histidine kinase |
| NkDv07_0968 | DNA-binding transcriptional regulator, MerR family |
| NkDv07_0979 | Prophage antirepressor |
| NkDv07_0984 | Membrane protein TolA involved in colicin uptake |
| RsTa-C01_0020 | ATP-dependent helicase/DNAse subunit B |
| RsTa-C01_0029 | Ribonuclease HII |
| RsTa-C01_0043 | REP element-mobilizing transposase RayT |
| RsTa-C01_0084 | REP element-mobilizing transposase RayT |
| RsTa-C01_0216 | REP element-mobilizing transposase RayT |
| RsTa-C01_0234 | REP element-mobilizing transposase RayT |
| RsTa-C01_0260 | REP element-mobilizing transposase RayT |
| RsTa-C01_0298 | REP element-mobilizing transposase RayT |
| RsTa-C01_0403 | REP element-mobilizing transposase RayT |
| RsTa-C01_0525 | REP element-mobilizing transposase RayT |
| RsTa-C01_0783 | REP element-mobilizing transposase RayT |
| RsTa-C01_1099 | REP element-mobilizing transposase RayT |
| RsTa-C01_0044 | Retron-type reverse transcriptase |
| RsTa-C01_0052 | Retron-type reverse transcriptase |
| RsTa-C01_0055 | Retron-type reverse transcriptase |
| RsTa-C01_0204 | Retron-type reverse transcriptase |
| RsTa-C01_0274 | Retron-type reverse transcriptase |
| RsTa-C01_0349 | Retron-type reverse transcriptase |
| RsTa-C01_0371 | Retron-type reverse transcriptase |
| RsTa-C01_0388 | Retron-type reverse transcriptase |
| RsTa-C01_0402 | Retron-type reverse transcriptase |
| RsTa-C01_0712 | Retron-type reverse transcriptase |
| RsTa-C01_0975 | Retron-type reverse transcriptase |
| RsTa-C01_0990 | Retron-type reverse transcriptase |
| RsTa-C01_0058 | Transposase |
| RsTa-C01_0076 | Transposase |

| | |
|---|---|
| RsTa-C01_0146 | Transposase |
| RsTa-C01_0245 | Transposase |
| RsTa-C01_0278 | Transposase |
| RsTa-C01_0290 | Transposase |
| RsTa-C01_0424 | Transposase |
| RsTa-C01_1090 | Transposase |
| RsTa-C01_1098 | Transposase |
| RsTa-C01_0089 | Transposase |
| RsTa-C01_0593 | Transposase |
| RsTa-C01_0611 | Transposase |
| RsTa-C01_0713 | Transposase |
| RsTa-C01_0765 | Transposase |
| RsTa-C01_0778 | Transposase |
| RsTa-C01_1086 | Transposase |
| RsTa-C01_0122 | ABC-type polar amino acid transport system, ATPase component |
| RsTa-C01_0138 | Adenine specific DNA methylase Mod |
| RsTa-C01_0383 | Adenine specific DNA methylase Mod |
| RsTa-C01_0139 | Superfamily II DNA or RNA helicase |
| RsTa-C01_0140 | Transposase |
| RsTa-C01_0188 | Transposase |
| RsTa-C01_1124 | Transposase |
| RsTa-C01_0141 | Magnesium-transporting ATPase (P-type) |
| RsTa-C01_0150 | Superfamily II DNA or RNA helicase, SNF2 family |
| RsTa-C01_0155 | Type I restriction-modification system, DNA methylase subunit |
| RsTa-C01_0186 | Transposase and inactivated derivatives, IS5 family |
| RsTa-C01_0230 | Lantibiotic modifying enzyme |
| RsTa-C01_0322 | Small-conductance mechanosensitive channel |
| RsTa-C01_0348 | Alanyl-tRNA synthetase |
| RsTa-C01_0377 | Primase-polymerase (Primpol) domain protein |
| RsTa-C01_0391 | Phage-related protein YomH |
| RsTa-C01_0916 | Phage-related protein YomH |
| RsTa-C01_0399 | Lysophospholipase, alpha-beta hydrolase superfamily |
| RsTa-C01_0400 | Phage portal protein BeeE |
| RsTa-C01_0410 | 5-methylcytosine-specific restriction endonuclease McrA |
| RsTa-C01_0460 | ATP-dependent Clp protease, ATP-binding subunit ClpA |
| RsTa-C01_0529 | Predicted Mg-chelatase, contains ChlI-like and ATPase domains, YifB family |
| RsTa-C01_0551 | DNA uptake channel protein ComEC, N-terminal domain |
| RsTa-C01_0583 | DNA uptake channel protein ComEC C-terminal domain, metallo-beta-lactamase superfamily |
| RsTa-C01_0643 | CRISPR-associated protein Csa3, CARF domain |

| | |
|---|---|
| RsTa-C01_0848 | Aspartate kinase |
| RsTa-C01_0849 | Translation elongation factor EF-4, membrane-bound GTPase |
| RsTa-C01_0863 | Serine kinase of the HPr protein, regulates carbohydrate metabolism |
| RsTa-C01_0870 | L,D-peptidoglycan transpeptidase YkuD, ErfK/YbiS/YcfS/YnhG family |
| RsTa-C01_0871 | Periplasmic beta-glucosidase and related glycosidases |
| RsTa-C01_0883 | mRNA-degrading endonuclease MazF, toxin component of the MazEF toxin-antitoxin module |
| RsTa-C01_0889 | Diaminopimelate epimerase |
| RsTa-C01_0999 | Hydroxymethylpyrimidine/phosphomethylpyrimidine kinase |
| RsTa-C01_1000 | Hydroxyethylthiazole kinase, sugar kinase family |
| RsTa-C01_1125 | Uncharacterized conserved protein, DUF917 family |
| RsTa-C01_1126 | N-methylhydantoinase A/oxoprolinase/acetone carboxylase, beta subunit |

* Pseudogenes not assigned to any COG are not listed.

**Table S10: Eukaryotic-like genes detected in the genomes of endosymbiotic *Clostridia*.**

| ID | domain[*] | RT-PCR[**] | most closely related organism | signal peptide prediction |
|---|---|---|---|---|
| CfP3-15_0039 | Rab | n.t. | *Tritrichomonas foetus* | Sec signal peptide |
| CfP3-15_0049 | Rho | not detected | *Naegleria gruberi* | Sec signal peptide |
| CfP3-15_0165 | PP2Ac | not detected | *Amphiprora paludosa* | not detected |
| CfP3-15_0235 | Rab | n.t. | *Trichomonas vaginalis* | not detected |
| CfP3-15_0282 | Rab | n.t. | *Trichomonas vaginalis* | not detected |
| CfP3-15_0445 | Rab | n.t. | *Trichoplax* sp. H2 | Sec signal peptide |
| CfP3-15_0456 | Rab | n.t. | *Pseudotrichonympha grassii* | not detected |
| CfP3-15_0548 | Rab | n.t. | *Candidatus* Helarchaeota archaeon | Sec signal peptide |
| CfP3-15_0552 | Roc | detected | *Tritrichomonas foetus* | Sec signal peptide |
| CfP3-15_0565 | Rab | n.t. | *Brachionus calyciflorus* | not detected |
| CfP3-15_0581 | Rab | n.t. | *Pseudotrichonympha grassii* | not detected |
| CfP3-15_0591 | Rab | n.t. | *Trepomonas* sp. PC1 | Sec signal peptide |
| CfP3-15_0627 | S_TKc | not detected | *Protomyces lactucae-debilis* | not detected |
| CfP3-15_0653 | Rab | n.t. | *Metschnikowia persimmonesis* | Sec signal peptide |
| CfP3-15_0780 | Rab | n.t. | *Cryptococcus amylolentus* | Sec signal peptide |
| CfP3-15_0790 | Gem1 | not detected | *Trepomonas* sp. PC1 | not detected |
| CfP3-15_0852 | Rab | not detected | *Pseudotrichonympha grassii* | Sec signal peptide |
| CfP3-15_0856 | Rab | n.t. | *Trichomonas vaginalis* | not detected |
| CfP3-15_0929 | Rab | n.t. | n.t. (no significant blast hits) | Sec signal peptide |
| CfP3-15_0934 | Rab | n.t. | *Paramecium tetraurelia* | not detected |
| RsTa-C01_0144 | Rab | not detected | *Ichthyophthirius multifiliis* | Sec signal peptide |
| RsTa-C01_0199 | S_TKc | n.t. | candidate division WOR-3 bcterium | not detected |
| RsTa-C01_0208 | PP2Ac | n.t. | *Candidatus* Falkowbacteria bacterium | not detected |
| RsTa-C01_0269 | LRR_5 | n.t. | n.t. (repeat protein) | not detected |
| RsTa-C01_0431 | UCH | detected | *Candida inconspicua* | not detected |
| RsTa-C01_0505 | LRR_5 | n.t. | n.t. (repeat protein) | not detected |
| RsTa-C01_0710 | LRR_5 | n.t. | n.t. (repeat protein) | not detected |
| RsTa-C01_0762 | Myb_DNA-binding | n.t. | n.t. (too short alignment) | not detected |

| | | | | |
|---|---|---|---|---|
| RsTa-C01_0774 | Myb_DNA-binding | not detected | *Trichonympha agilis* | not detected |
| RsTa-C01_0775 | Myb_DNA-binding | n.t. | *Trichonympha agilis* | Lipoprotein signal peptide |
| RsTa-C01_0844 | S_TKc | detected | *Trichonympha agilis* | Sec signal peptide |
| RsTa-C01_0845 | LRR_5 | n.t. | n.t. (repeat protein) | Sec signal peptide |
| RsTa-C01_0983 | PHA03095 | n.t. | n.t. (repeat protein) | not detected |
| RsTa-C01_0984 | PHA03095 | n.t. | n.t. (repeat protein) | not detected |
| RsTa-C01_0992 | S_TKc | n.t. | *Stentor coeruleus* | Sec signal peptide |
| NkDv07_0114 | Myosin_tail_1 | n.t. | *Apis cerana cerana* | Sec signal peptide |
| NkDv07_0121 | Metallophos | n.t. | Rhodobacteraceae bacterium | Sec signal peptide |
| NkDv07_0395 | Gem1 | n.t. | n.t. (no significant blast hits) | Sec signal peptide |
| NkDv07_0512 | Rnd2_Rho7 | n.t. | *Nyssa sinensis* | Sec signal peptide |
| NkDv07_0536 | MPP_PPP_family | n.t. | *Trichomonas vaginalis* | not detected |
| NkDv07_0615 | OGFr_N | n.t. | *Letharia lupina* | not detected |
| NkDv07_0673 | PTZ00341 | n.t. | n.t. (too short alignment) | Sec signal peptide |

* Rab, Ras-related in brain (Rab) family of small guanosine triphosphatases (GTPases); Rho, Ras homology family (Rho) of small guanosine triphosphatases (GTPases); PP2Ac, Protein phosphatase 2A homologues, catalytic domain; Roc, Ras of Complex, Roc, domain of DAPkinase; S_TKc, Serine/Threonine protein kinases, catalytic domain; Gem1, GTPase SAR1 family domain; LRR_5, Leucine rich repeats (6 copies); UCH, Ubiquitin carboxyl-terminal hydrolase; Myb_DNA-binding, Myb-like DNA-binding domain; PHA03095, ankyrin-like protein; Myosin_tail_1, Myosin tail; Metallophos, Calcineurin-like phosphoesterase; Rnd2_Rho7, Rnd2/Rho7 GTPases; MPP_PPP_family, phosphoprotein phosphatases of the metallophosphatase superfamily, metallophosphatase domain; OGFr_N, Opioid growth factor receptor (OGFr) conserved region; PTZ00341, Ring-infected erythrocyte surface antigen.

** n.t., not tested.

**Table S11: General features of MAGs in the small genome clade.**

| accession number | host | geographic range | total length (bp) | number of contigs | G+C (%) | completeness (%)* | predicted genome size (Mbp) |
|---|---|---|---|---|---|---|---|
| GCA_902796915.1 | cattle | Europe | 1404720 | 145 | 31.0 | 100.0 | 1.40 |
| GCA_017406805.1 | goat | Asia | 1157853 | 53 | 33.9 | 97.8 | 1.18 |
| GCA_017414725.1 | goat | Asia | 864431 | 150 | 32.3 | 92.4 | 0.94 |
| GCA_017431755.1 | goat | Asia | 812295 | 116 | 32.3 | 94.6 | 0.86 |
| GCA_017432165.1 | goat | Asia | 1234861 | 103 | 33.7 | 98.9 | 1.25 |
| GCA_017448285.1 | goat | Asia | 869701 | 78 | 32.4 | 97.8 | 0.89 |
| GCA_000435275.1 | human | North America, Europe, Asia, Oceania | 1114344 | 132 | 34.5 | 96.7 | 1.15 |
| GCA_900552265.1 | human | Europe, Asia, Oceania | 1119698 | 17 | 32.9 | 100.0 | 1.12 |
| GCA_900554335.1 | human | South America | 1073904 | 14 | 37.3 | 98.9 | 1.09 |
| GCA_900554775.1 | human | North America, Europe, Asia, Oceania | 864877 | 127 | 38.2 | 91.3 | 0.95 |
| MGYG000000505 | human | Oceania, Africa | 1142930 | 172 | 34.9 | 84.8 | 1.35 |
| MGYG000000607 | human | Africa | 1023795 | 31 | 35.6 | 96.7 | 1.06 |
| MGYG000001125 | human | Asia | 1115602 | 240 | 33.7 | 83.7 | 1.33 |
| MGYG000001801 | human | Europe | 1294689 | 229 | 35.1 | 90.2 | 1.44 |
| MGYG000003453 | human | Oceania | 825696 | 257 | 33.1 | 75.0 | 1.10 |
| MGYG000004322 | human | Europe, Asia | 1235249 | 21 | 34.6 | 100.0 | 1.24 |
| MGYG000004437 | human | North America, Europe, Asia, Oceania, South America, Africa | 1139688 | 15 | 35.3 | 100.0 | 1.14 |
| MGYG000004496*** | human | Asia | 1393460 | 182 | 33.6 | 73.9 | 1.89 |
| GCA_017480265.1 | roe deer | Asia | 883795 | 85 | 36.7 | 98.9 | 0.89 |
| GCA_002372375.1 | sheep | Oceania | 1440018 | 55 | 34.5 | 100.0 | 1.44 |

| | | | | | | | |
|---|---|---|---|---|---|---|---|
| GCA_017515685.1 | sheep | Asia | 853097 | 306 | 39.2 | 72.8 | 1.17 |
| GCA_009785985.1 | soil-feeding termite (*Isognathotermes ugandensis* ) | Africa | 1238722 (2692270)** | 277 (648)** | 30.4 (30.5)** | 90.2 (92.4)** | 1.4 (2.9)** |
| GCA_017634155.1 | water deer | Asia | 912766 | 64 | 30.8 | 97.8 | 0.93 |

* Caluculated based on modified single-copy marker gene sets (Table S7).

** Before re-binning.

*** Detected single-copy marker gene set was considerably different from other members of the small genome clade;

**Table S12: Descriptions of the eukaryotic-like domains detected in the small genome clade.**

| domain | description |
|---|---|
| STKc_MAP | Aldo/keto reductase, related to diketogulonate reductase |
| AKR_AKR1-5-like | AKR1/2/3/4/5 family of aldo-keto reductase (AKR) and similar proteins |
| Myb_DNA-binding | Myb-like DNA-binding domain |
| S_TKc | Serine/Threonine protein kinases, catalytic domain |
| PKc | Catalytic domain of Protein Kinases |
| STKc_KSR1 | Catalytic domain of the Serine/Threonine Kinase, Kinase Suppressor of Ras 1 |
| STKc_Chk1 | Catalytic domain of the Serine/Threonine kinase, Checkpoint kinase 1 |
| SPS1 | Serine/threonine protein kinase |
| STKc_MAPKKK | Catalytic domain of the Serine/Threonine Kinase, Mitogen-Activated Protein Kinase Kinase Kinase |
| Pkinase | Protein kinase domain |
| MPP_PPP_family | Phosphoprotein phosphatases of the metallophosphatase superfamily, metallophosphatase domain |
| PP2Ac | Protein phosphatase 2A homologues, catalytic domain |
| PTPc | Catalytic domain of protein tyrosine phosphatases |
| LRR_5 | Leucine rich repeats (6 copies) |
| TPR | Tetratricopeptide repeat |
| Ank_2 | Ankyrin repeats (3 copies) |
| PHA03095 | Ankyrin-like protein |
| PHA02874 | Ankyrin repeat protein |
| PHA03100 | Ankyrin repeat protein |
| Rab | Ras-related in brain (Rab) family of small guanosine triphosphatases (GTPases) |
| Rab6 | Rab GTPase family 6 |
| Roc | Ras of Complex |

| | |
|---|---|
| Rho | Ras homology family (Rho) of small guanosine triphosphatases (GTPases) |
| Ras_like_GTPase | Rat sarcoma (Ras)-like superfamily of small guanosine triphosphatases (GTPases) |
| Rnd2_Rho7 | Rnd2/Rho7 GTPases |
| Gem1 | GTPase SAR1 family domain |
| UCH | Ubiquitin carboxyl-terminal hydrolase |
| Peptidase_C19 | Peptidase C19 |
| zf-RING_2 | Ring finger domain |
| Tcp10_C | T-complex protein 10 C-terminus |
| Lipase_3 | Lipase (class 3) |
| PLN02486 | Aminoacyl-tRNA ligase |
| PTZ00341 | Ring-infected erythrocyte surface antigen |
| Metallophos | Calcineurin-like phosphoesterase |
| OGFr_N | Opioid growth factor receptor (OGFr) conserved region |
| Methyltransf_FA | Farnesoic acid 0-methyl transferase |
| Myosin_tail_1 | Myosin tail |
| PTZ00121 | Merozoite apical erythrocyte binding ligand |
| PLN03091 | Hypothetical protein |
| CCDC158 | Coiled-coil domain-containing protein 158. |

**Table S13: Genes predicted to have been acquired in the last common ancestor of the small genome clade.**

| COG/orthogroup | description |
|---|---|
| COG3202 | ATP/ADP translocase |
| COG2094 | 3-methyladenine DNA glycosylase Mpg |
| COG0531 | serine transporter YbeC |
| COG3740 | phage head maturation protease |
| OG0000732 | NHLM bacteriocin system secretion protein |
| OG0000787 | multidrug and toxic compound extrusion-like protein |
| OG0001223 | leucine rich repeats containing protein |
| OG0002080 | head-tail connector protein gp6 |
| OG0002469 | phage tail protein |
| OG0000920 | hypothetical protein |
| OG0001554 | hypothetical protein |
| OG0003398 | hypothetical protein |
| OG0004343 | hypothetical protein |
| OG0005233 | hypothetical protein |
| OG0005234 | hypothetical protein |

#### Supplementary Figures

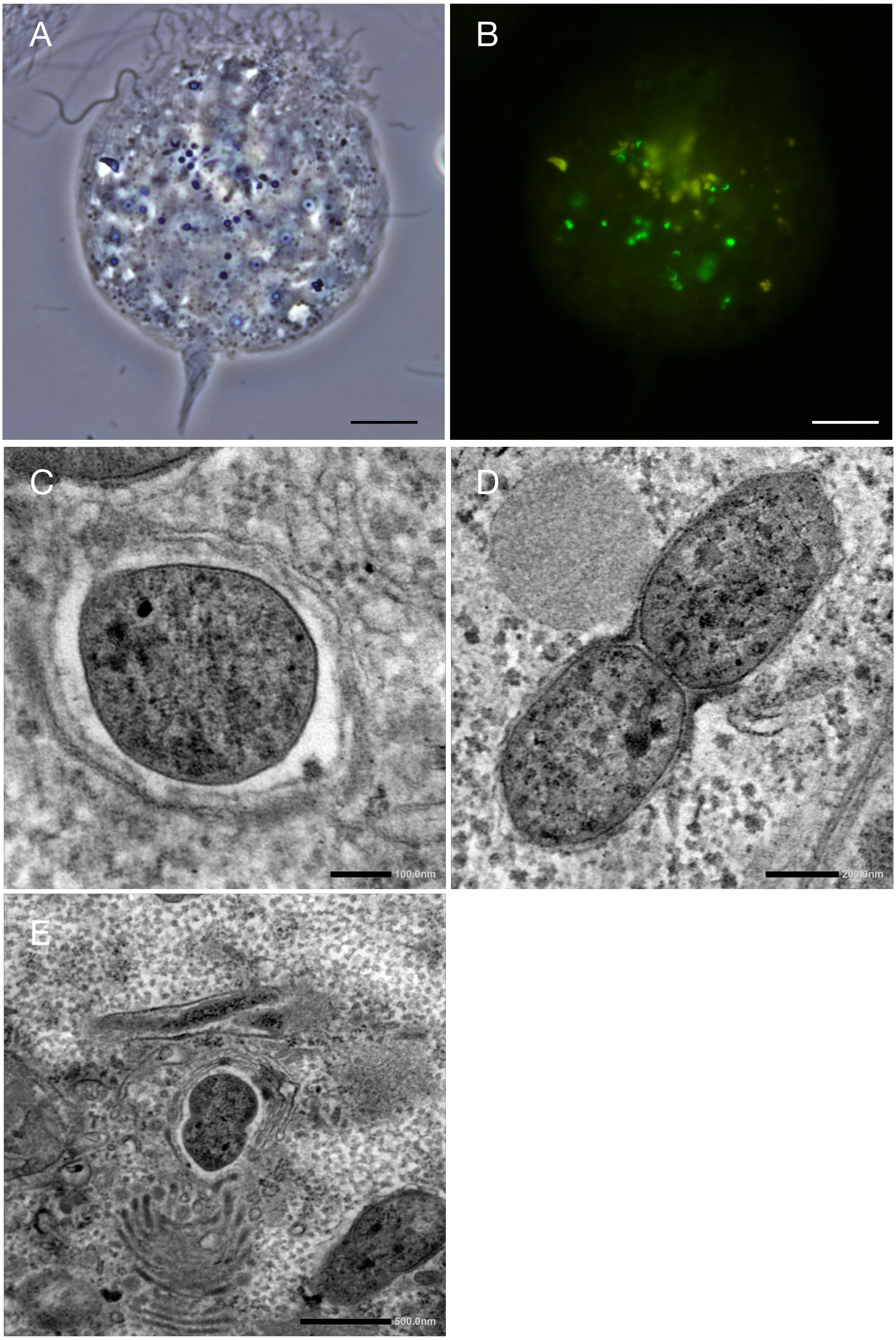


**Figure S1.** Fluorescence in situ hybridization analysis of 16S rRNA phylotype NkDv07 (**A, B**) and transmission electron micrographs of intracellular bacterial symbionts of *Pseudotrichonympha grassii* (**C–E**). Phase contrast image of *Devescovina* sp. from the gut of *Neotermes sugioi* (A) and NkDv07 cells detected using 6FAM-labelled probe NkDv07-142 (green) as single cocci (**B**). Amorphous yellowish autofluorescence in B was emitted from wood particles ingested by *Devescovina* sp. Putative clostridial cell (**C**), and "*Candidatus* Azobacteroides pseudotrichonymphae" cells (**D**). In panel E, a putative clostridial cell apparently during fission was in the center and a "*Ca.* A. pseudotrichonymphae" cell in the lower right. Bars: **A** and **B**, 10 µm; **C**, 100 nm; **D**, 200 nm; **E**, 500 nm.

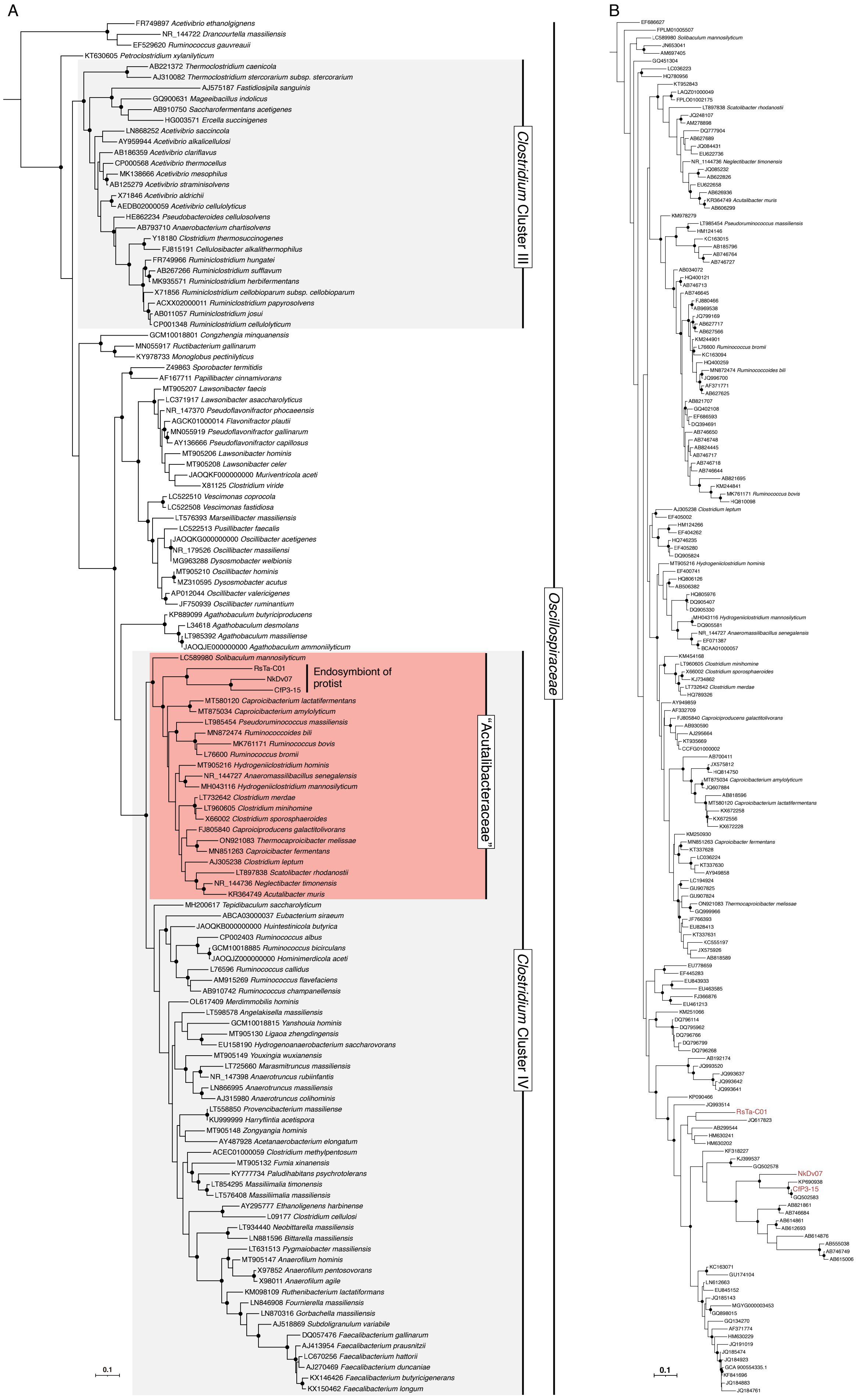


**Figure S2.** Phylogenetic positions of endosymbiotic *Clostridia* based on 16S rRNA gene sequences. (**A**) Maximum-likelihood tree of endosymbiotic *Clostridia* and other members of the family *Oscillospiraceae*. Described species of *Oscillospiraceae* were included in the tree. The tree was constructed with the GTR+I+G4 nucleotide substitution model and rooted with *Syntrophomonas wolfei* (CP000448) and *Thermosyntropha lipolytica* (FQWY01000001) as the outgroup. Clusters of *Clostridium* [60] and the clade corresponding to "Acutalibacteraceae" in the GTDB taxonomy are highlighted in gray and red, respectively. (**B**) Maximum-likelihood tree of endosymbiotic *Clostridia* and other members of the "Acutalibacteraceae". Uncultured clones closely related to described species and endosymbiotic *Clostridia* were included. The tree was constructed with the GTR+G4 nucleotide substitution model and rooted with *Sporobacter termitidis* (Z49863) and *Oscillibacter ruminantium* (JF750939) as the outgroup. Highly supported nodes (SH-aLRT value ≥ 80% and ultrafast bootstrap support value ≥ 95%) are highlighted with a closed circle.

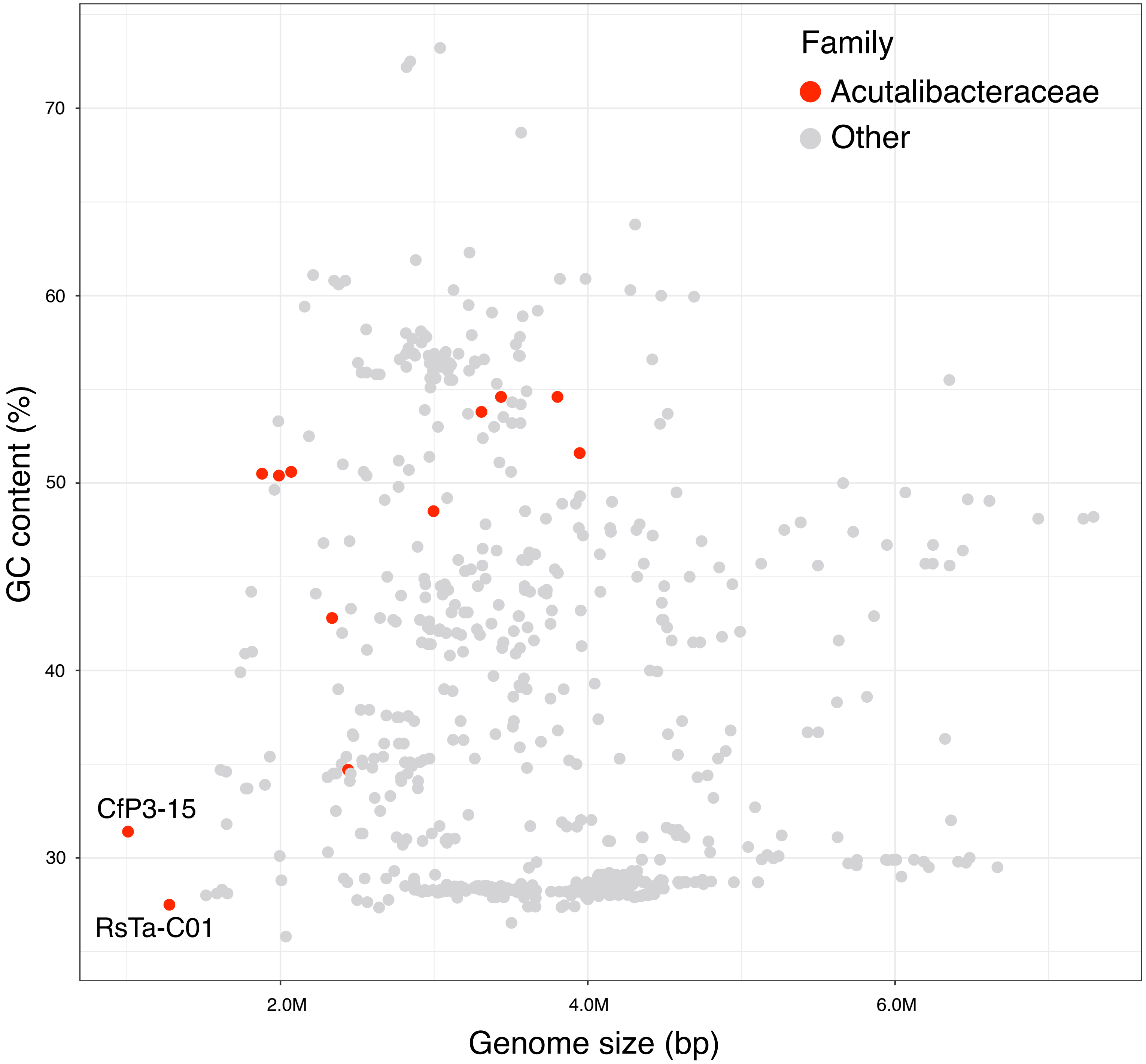


**Figure S3.** Relationships between the size and GC content of the complete genomes in the class *Clostridia*.

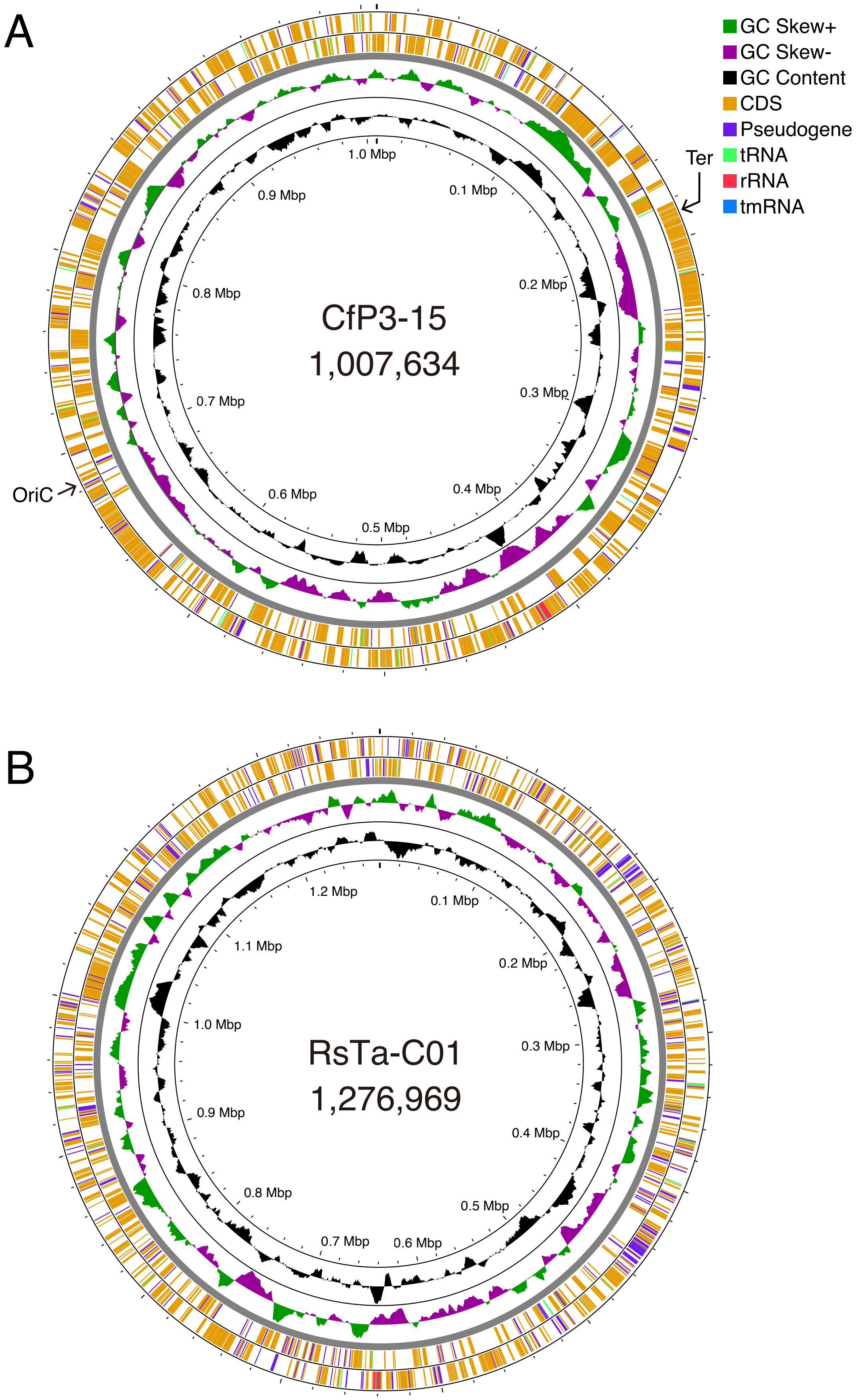


**Figure S4.** Circular representation of the CfP3-15 chromosome (**A**) and the RsTa-C01 chromosome (**B**). The rings denote the following features (from inside): (i) Scale in mega base pairs of the chromosome (black); (ii) GC content (black); (iii) GC skew (green and purple); (iv) predicted genes present on the forward strand (+); (v) genes present on the reverse strand (-). OriC and Ter sites are indicated by arrows in **A**. For the RsTa-C01 genome, we could not detect these sites, using Ori-Finder 2022.

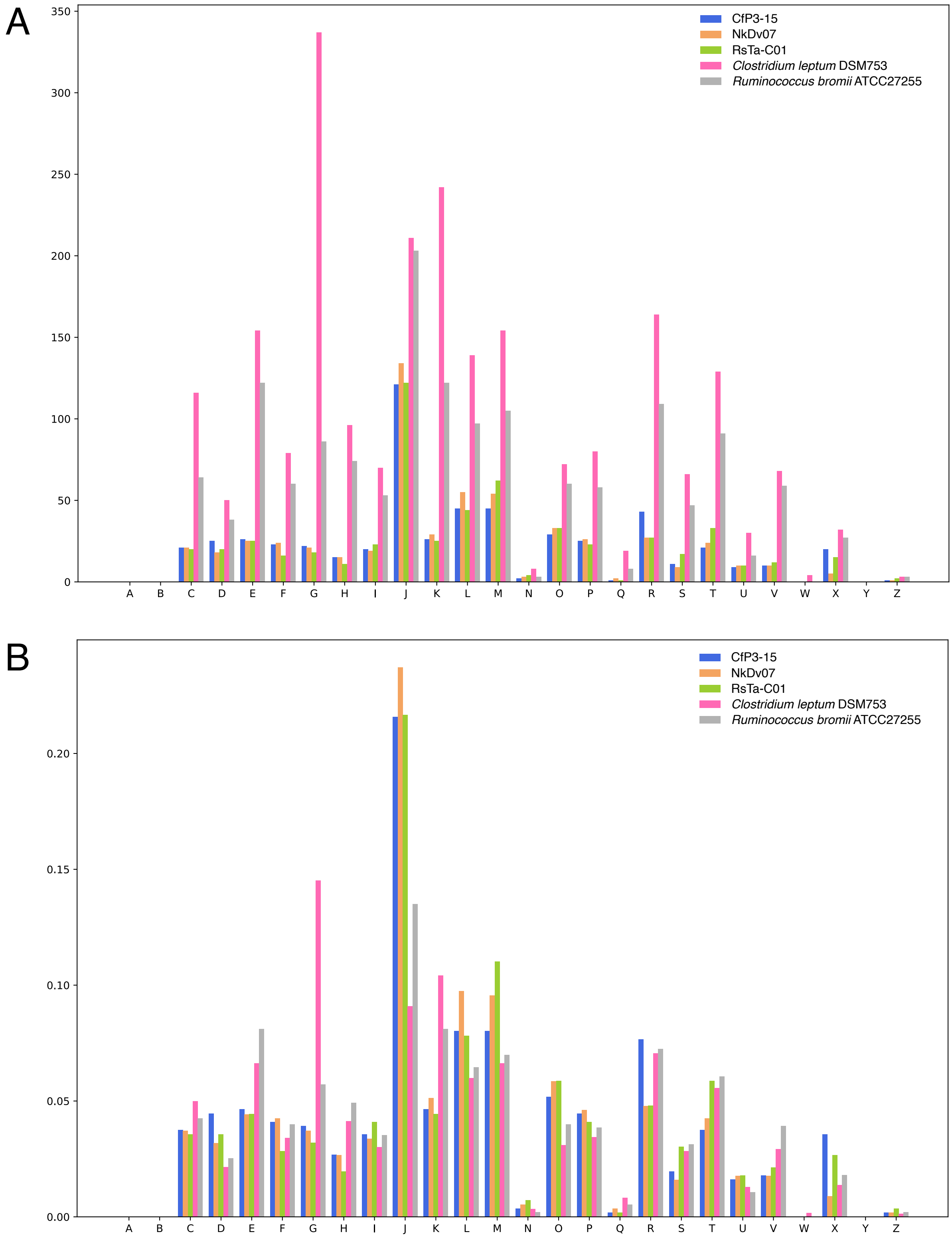


**Figure S5.** Comparison of the number (**A**) and ratio (**B**) of genes classified into clusters of orthologous genes (COG) functional categories. The genomes of CfP3-15, NkDv07, RsTa-C01, and their free-living relatives *Clostridium leptum* and *Ruminococcus bromii* were compared. The categories denote the following functions: (A) RNA processing and modification; (B) chromatin structure and dynamics; (C) energy production and conversion; (D) cell cycle control, cell division, chromosome partitioning; (E) amino acid transport and metabolism; (F) nucleotide transport and metabolism; (G) carbohydrate transport and metabolism; (H) coenzyme transport and metabolism; (I) lipid transport and metabolism; (J) translation, ribosomal structure and biogenesis; (K) transcription; (L) replication, recombination and repair; (M) cell wall/membrane/envelope biogenesis; (N) cell motility; (O) posttranslational modification, protein turnover, chaperones; (P) inorganic ion transport and metabolism; (Q) secondary metabolites biosynthesis, transport and catabolism; (R) general function prediction only; (S) function unknown; (T) signal transduction mechanisms; (U) intracellular trafficking, secretion, and vesicular transport; (V) defense mechanisms; (W) extracellular structures; (X) mobilome: prophages, transposons; (Y) nuclear structure; (Z) cytoskeleton.

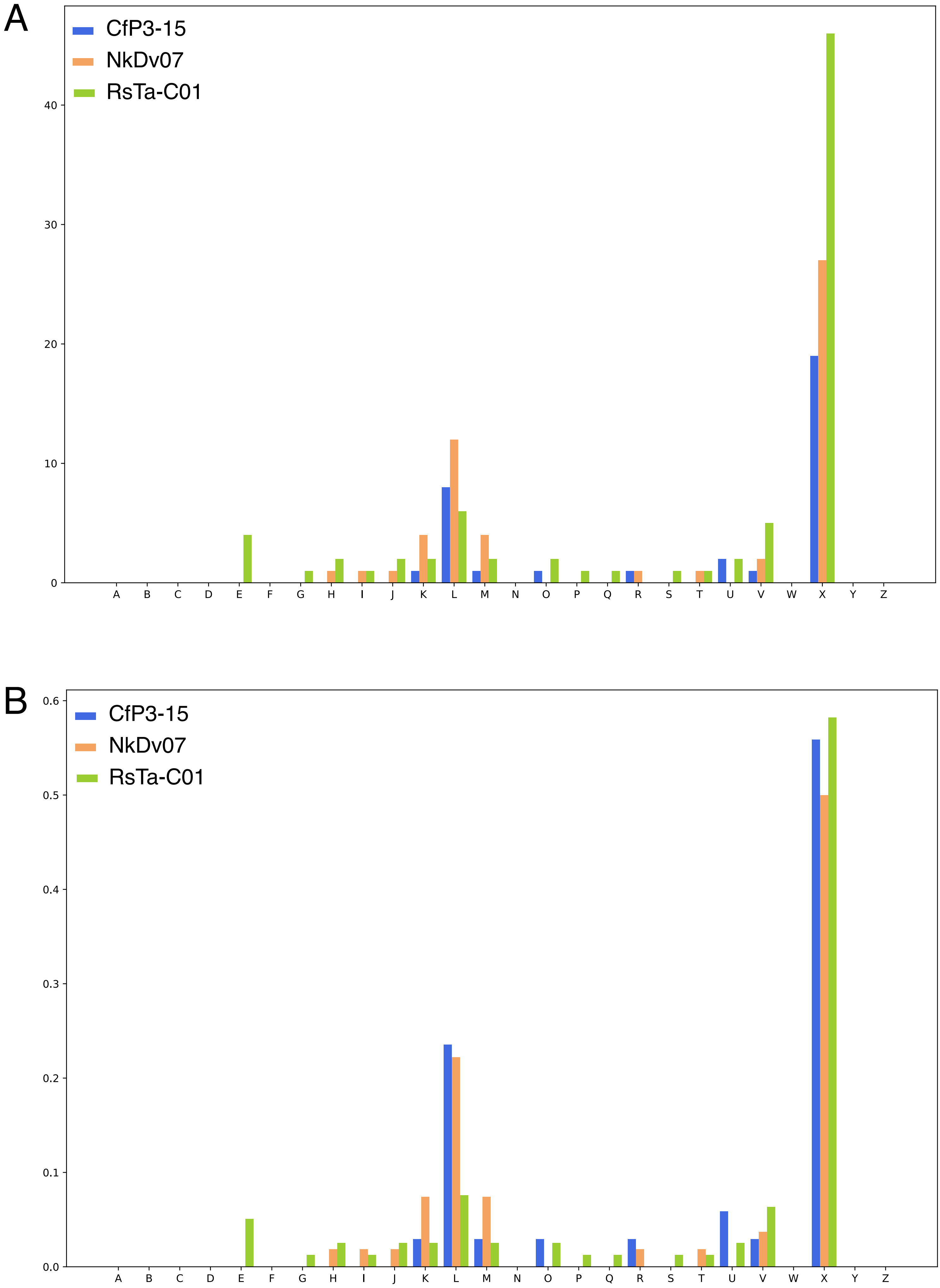


**Figure S6.** Comparison of the number (**A**) and ratio (**B**) of pseudogenes classified into clusters of orthologous genes (COG) functional categories. See also the legend to Fig. S5.

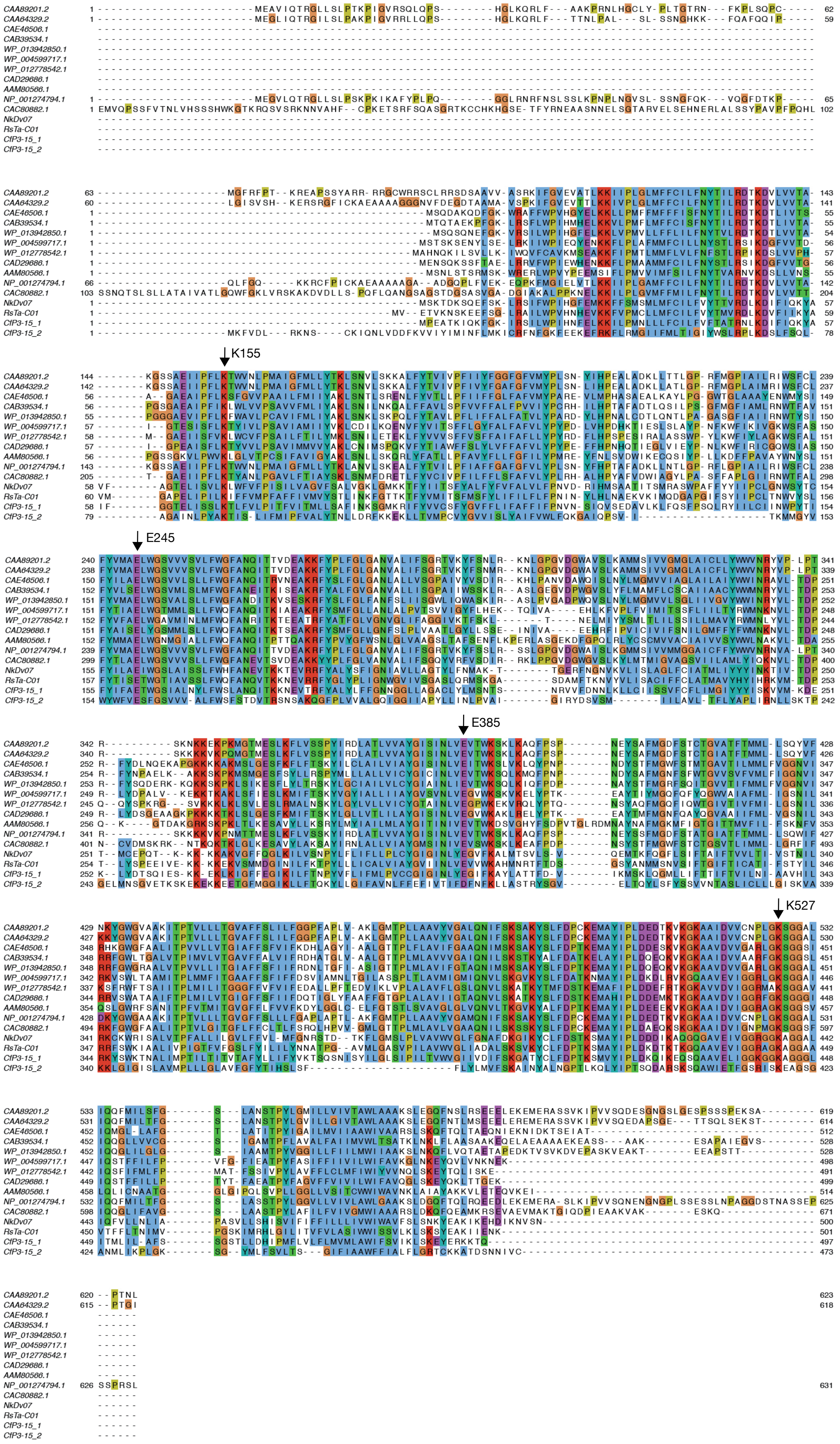


**Figure S7.** Amino acid sequence alignment of ATP/ADP translocase and other nucleotide transport proteins from the endosymbiotic *Clostridia*, other parasitic bacteria, and plastids. Conserved residues important for substrate specificity, transport efficiency, and counter exchange of ATP/ADP translocase (CAA89201.2) of the plastid of *Arabidopsis thaliana* are indicated with arrows.

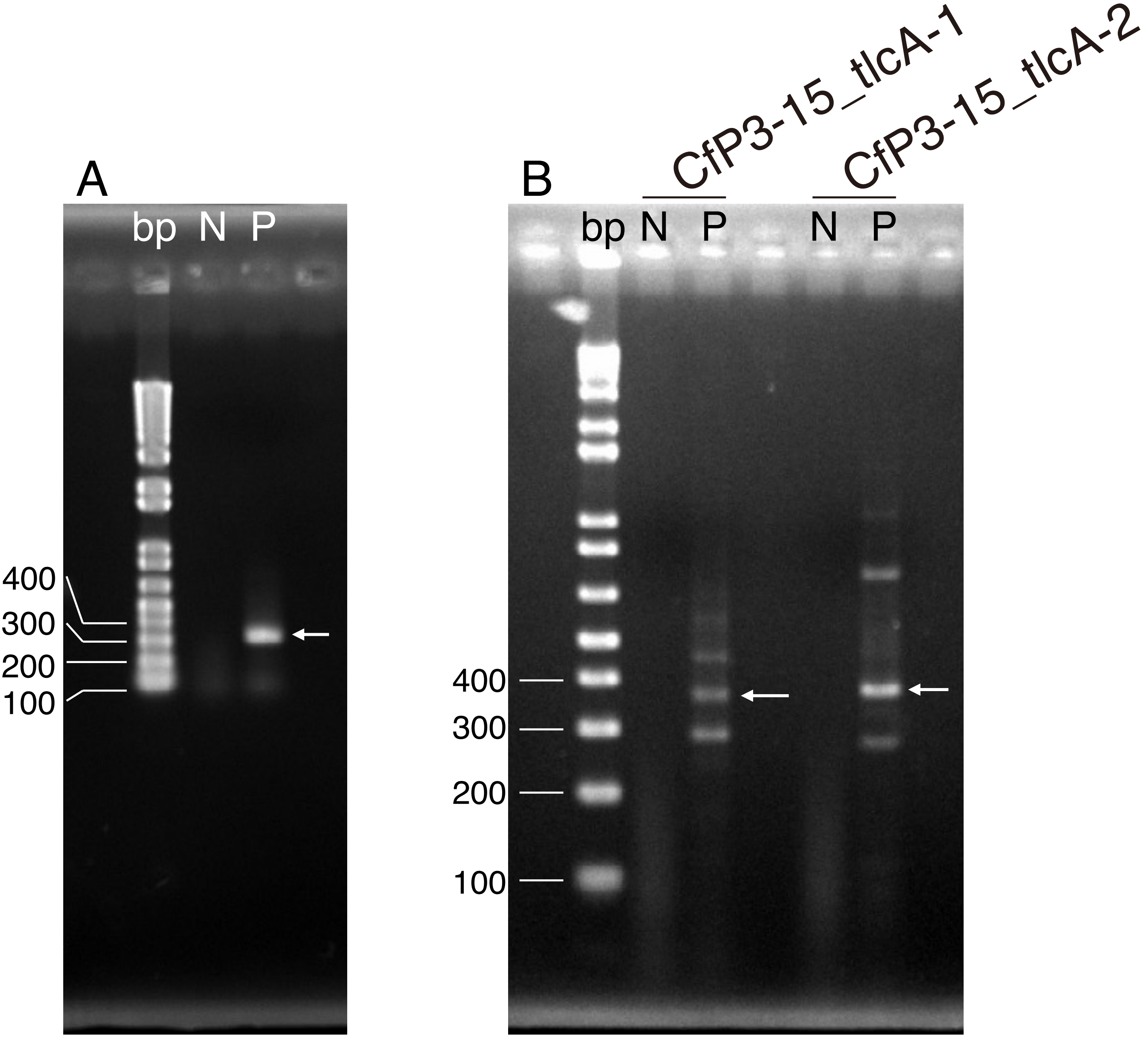


**Figure S8.** RT-PCR analysis of the ATP/ADP translocase gene. Agarose gel electrophoresis of RT-PCR products from total RNA of *R. speratus* guts (**A**) and *C. formosanus* guts (**B**). N, negative control (RNA samples without reverse transcription); P, products of RT-PCR using primer sets specific to the respective ATP/ADP translocase genes (see Table S2). These bands were excised from gels, and their sequences were verified by cloning and Sanger sequencing analysis.

A

B

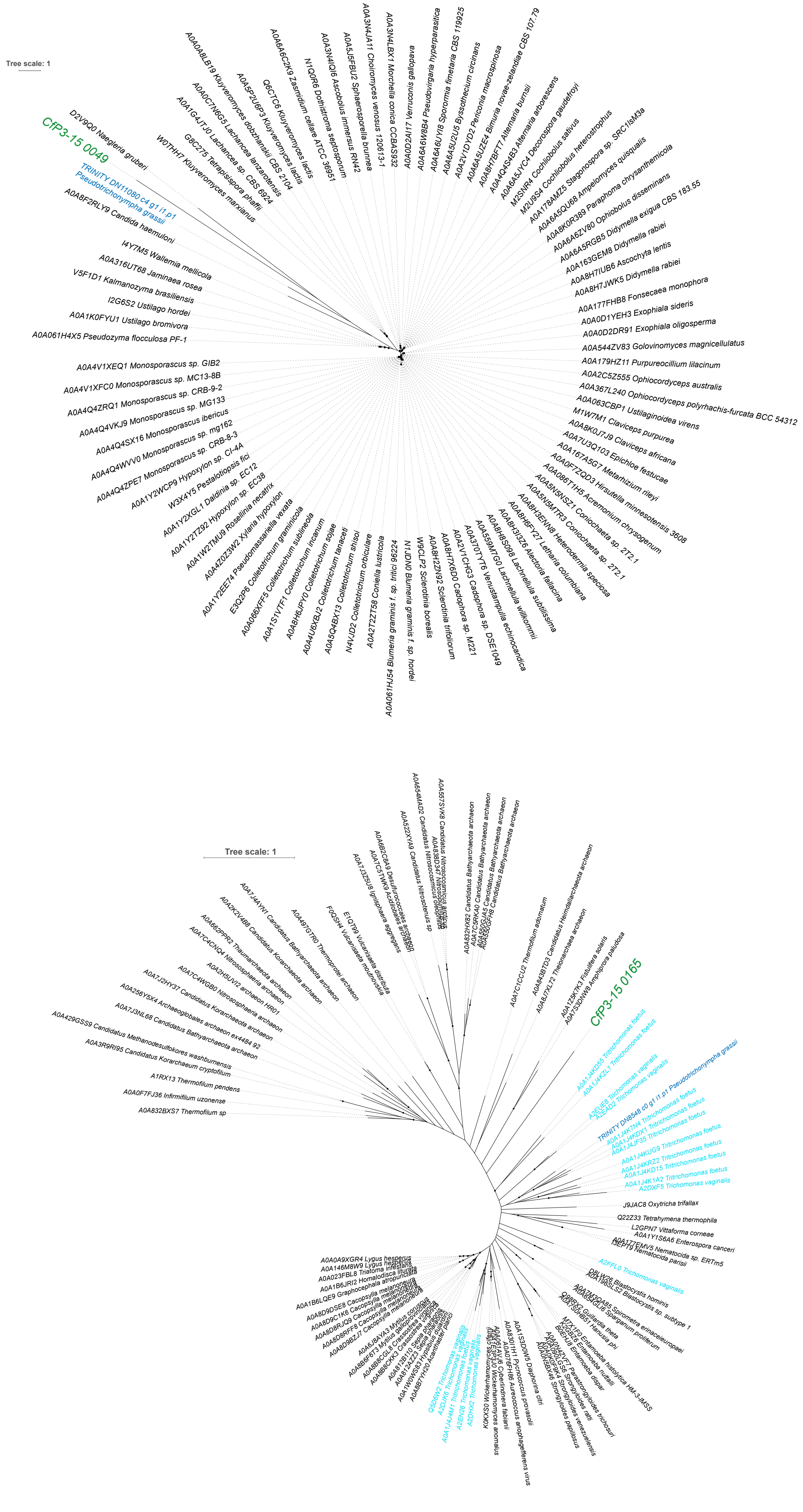
Tree scale: 1
CfP3-15 0049
TRINITY DN11080 c4 g1 i1.p1 Pseudotrichonympha grassii
Tree scale: 1
CfP3-15 0165

C

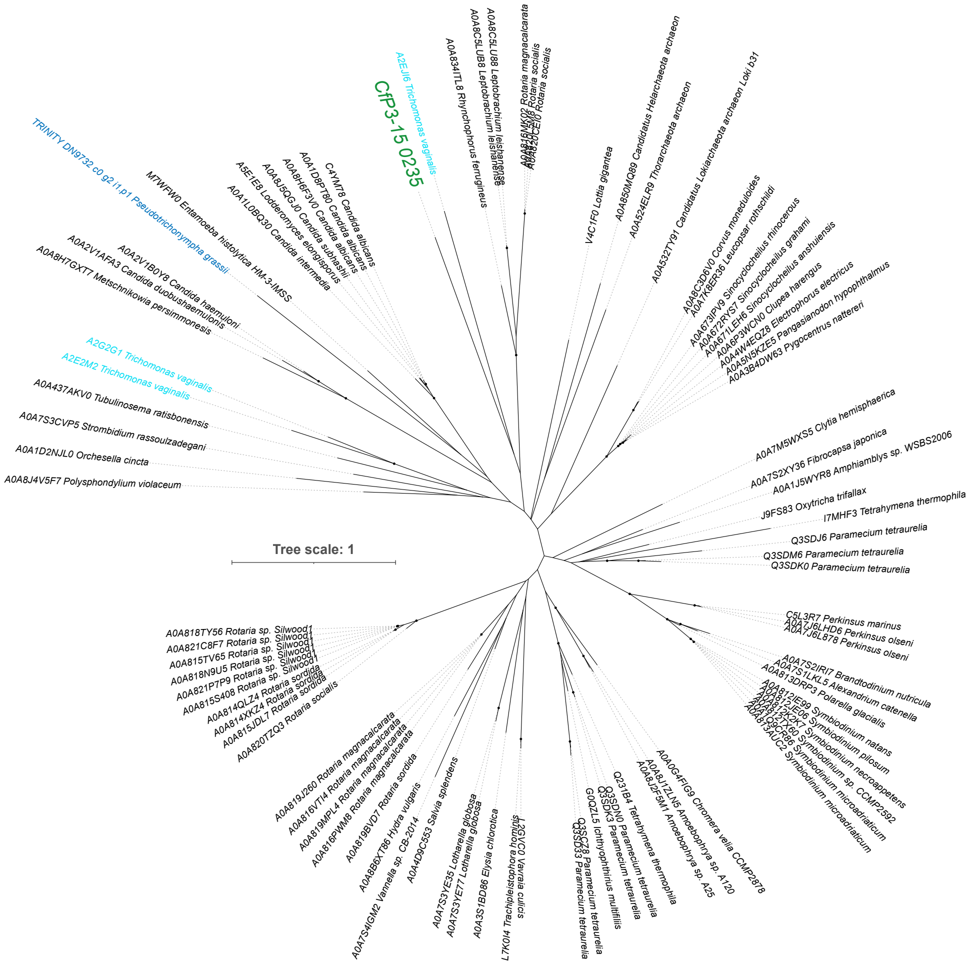

CfP3-15 0235
A2EJI6 Trichomonas vaginalis
TRINITY DN9732 c0 g2 i1.p1 Pseudotrichonympha grassii
M7WFW0 Entamoeba histolytica HM-3:IMSS
A2G2G1 Trichomonas vaginalis
A2E2M2 Trichomonas vaginalis
A0A437AKV0 Tubulinosema ratisbonensis
A0A7S3CVP5 Strombidium rassoulzadegani
A0A1D2NJL0 Orchesella cincta
A0A8J4V5F7 Polysphondylium violaceum
Tree scale: 1
V4C1F0 Lottia gigantea
A0A7M5WXS5 Clytia hemisphaerica
A0A7S2XY36 Fibrocapsa japonica
J9FS83 Oxytricha trifallax
I7MHF3 Tetrahymena thermophila
Q3SDJ6 Paramecium tetraurelia
Q3SDM6 Paramecium tetraurelia
Q3SDK0 Paramecium tetraurelia
C5L3R7 Perkinsus marinus
A0A0G4FIG9 Chromera velia CCMP2878
L7K0I4 Trachipleistophora hominis
A0A3S1BD86 Elysia chlorotica
A0A4D9C553 Salvia splendens


D

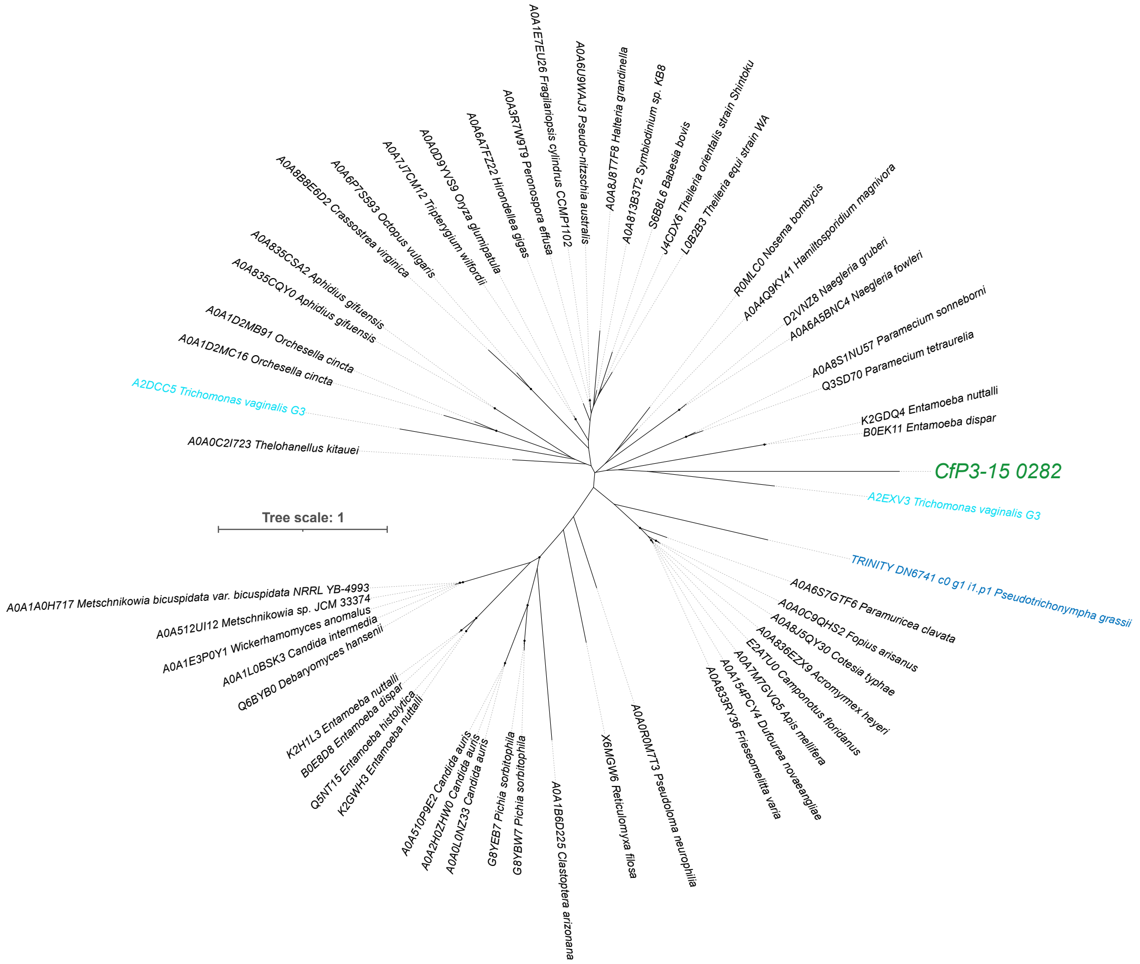

A0A1E7EU26 Fragilariopsis cylindrus CCMP1102
A0A6U9WAJ3 Pseudo-nitzschia australis
A0A8J8T7F8 Halteria grandinella
A0A813B3T2 Symbiodinium sp. KB8
S6BBL6 Babesia bovis
J4CDX6 Theileria orientalis strain Shintoku
L0B2B3 Theileria equi strain WA
A0A3R7W9T9 Peronospora effusa
A0A6A7FZ22 Hirondellea gigas
A0A0D9VVS9 Oryza glumipatula
A0A7J7CM12 Tripterygium wilfordii
A0A6P7S593 Octopus vulgaris
A0A8B8E6D2 Crassostrea virginica
A0A835CSA2 Aphidius gifuensis
A0A835CQY0 Aphidius gifuensis
A0A1D2MB91 Orchesella cincta
A0A1D2MC16 Orchesella cincta
A2DCC5 Trichomonas vaginalis G3
A0A0C2I723 Thelohanellus kitauei
R0MLC0 Nosema bombycis
A0A4Q9KY41 Hamiltosporidium magnivora
D2VNZ8 Naegleria gruberi
A0A6A5BNC4 Naegleria fowleri
A0A8S1NU57 Paramecium sonneborni
Q3SD70 Paramecium tetraurelia
K2GDQ4 Entamoeba nuttalli
B0EK11 Entamoeba dispar
CfP3-15 0282
A2EXV3 Trichomonas vaginalis G3
TRINITY DN6741 c0 g1 i1.p1 Pseudotrichonympha grassii
Tree scale: 1
A0A1A0H717 Metschnikowia bicuspidata var. bicuspidata NRRL YB-4993
A0A512UI12 Metschnikowia sp. JCM 33374
A0A1E3P0Y1 Wickerhamomyces anomalus
A0A1L0BSK3 Candida intermedia
Q6BYB0 Debaryomyces hansenii
K2H1L3 Entamoeba nuttalli
B0E8D8 Entamoeba dispar
Q5NT15 Entamoeba histolytica
K2GWH3 Entamoeba nuttalli
A0A510P9E2 Candida auris
A0A2H0ZHW0 Candida auris
A0A0L0NZ33 Candida auris
G8YEB7 Pichia sorbitophila
G8YBW7 Pichia sorbitophila
A0A1B6D225 Clastoptera arizonana
X6MGW6 Reticulomyxa filosa
A0A0R0M7T3 Pseudoloma neurophilia
A0A6S7GTF6 Paramuricea clavata
A0A0C9QHS2 Fopius arisanus
A0A8J5QY30 Cotesia typhae
A0A836EZX9 Acromyrmex heyeri
E2ATU0 Camponotus floridanus
A0A7M7GVQ5 Apis mellifera
A0A154PCY4 Dufourea novaeangliae
A0A833RY36 Friesomelitta varia

E

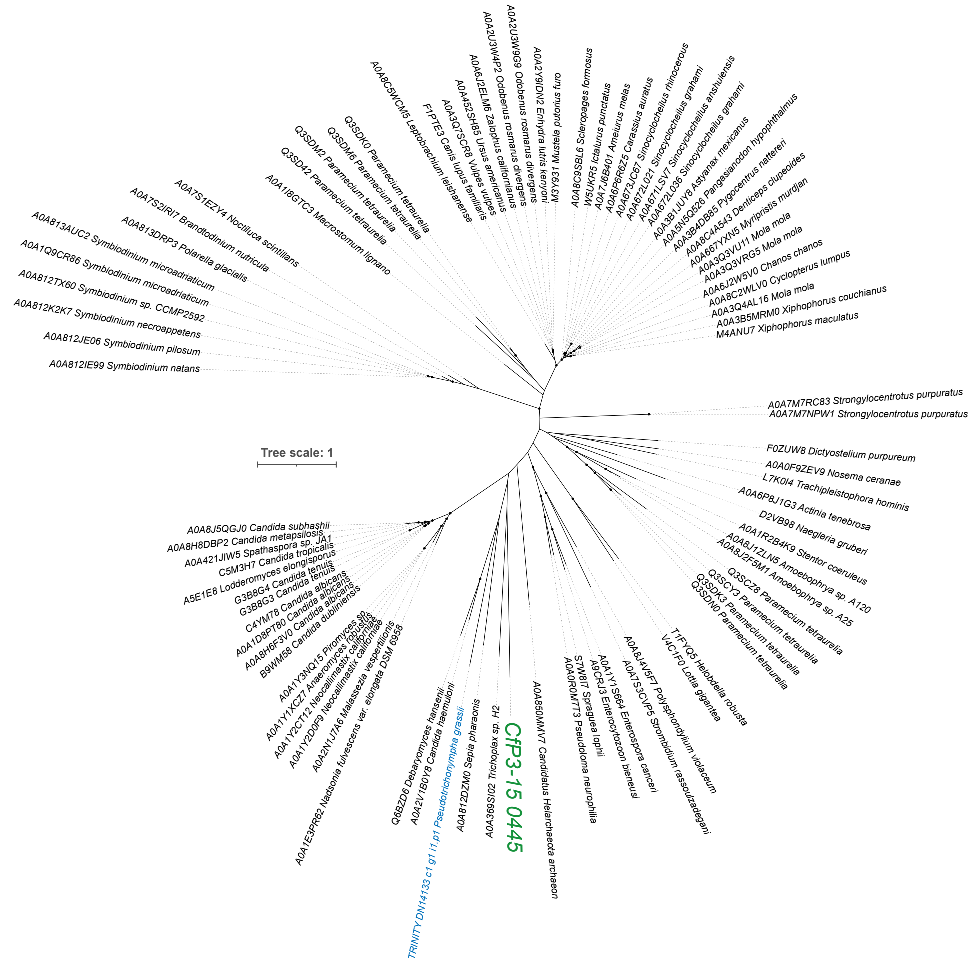
Tree scale: 1
A0A812IE99 Symbiodinium natans
A0A812JE06 Symbiodinium pilosum
A0A812K2K7 Symbiodinium necroappetens
A0A812TX60 Symbiodinium sp. CCMP2592
A0A1Q9CR86 Symbiodinium microadriaticum
A0A813AUC2 Symbiodinium microadriaticum
A0A813DRP3 Polarella glacialis
A0A7S2IRI7 Brandtodinium nutricula
A0A7S1EZY4 Noctiluca scintillans
A0A1I8GTC3 Macrostomum lignano
Q3SD42 Paramecium tetraurelia
Q3SDM2 Paramecium tetraurelia
Q3SDM6 Paramecium tetraurelia
Q3SDK0 Paramecium tetraurelia
A0A8C5WCM5 Leptobrachium leishanense
F1PTE3 Canis lupus familiaris
A0A3Q7SCR8 Vulpes vulpes
A0A452SHB5 Ursus americanus
A0A6J2ELM6 Zalophus californianus
A0A2U3W4P2 Odobenus rosmarus divergens
A0A2U3WG59 Odobenus rosmarus divergens
A0A2Y9IDN2 Enhydra lutris kenyoni
M3YXA31 Mustela putorius furo
A0A8C9SBL6 Scleropages formosus
W5UKR5 Ictalurus punctatus
A0A7J6B401 Ameiurus melas
A0A6P6R6Z5 Carassius auratus
A0A673JC67 Sinocyclocheilus rhinocerous
A0A672LSV7 Sinocyclocheilus grahami
A0A672L036 Sinocyclocheilus grahami
A0A3B1JUY8 Astyanax mexicanus
A0A5N5Q526 Pangasianodon hypophthalmus
A0A3B4DB85 Pygocentrus nattereri
A0A8C4A543 Denticeps clupeoides
A0A667YXN5 Myripristis murdjan
A0A3Q3VU11 Mola mola
A0A3Q3VRG5 Mola mola
A0A2J5W5V0 Chanos chanos
A0A8C2WLV0 Cyclopterus lumpus
A0A3Q4AL16 Mola mola
A0A3B5MRM0 Xiphophorus couchianus
M4ANU7 Xiphophorus maculatus
A0A7M7RC83 Strongylocentrotus purpuratus
A0A7M7NPW1 Strongylocentrotus purpuratus
F0ZUW8 Dictyostelium purpureum
A0A0F9ZEV9 Nosema ceranae
L7K0I4 Trachipleistophora hominis
A0A6P8J1G3 Actinia tenebrosa
D2VB98 Naegleria gruberi
A0A1R2B4K9 Stentor coeruleus
A0A8J1ZLN5 Amoebophrya sp. A120
A0A8J2F5M1 Amoebophrya sp. A25
Q3SCZ8 Paramecium tetraurelia
Q3SCY3 Paramecium tetraurelia
Q3SDK3 Paramecium tetraurelia
Q3SDN0 Paramecium tetraurelia
T1FYQ5 Helobdella robusta
V4C1F0 Lottia gigantea
A0A8J4V5F7 Polysphondylium violaceum
A0A7S3CVP5 Strombidium rassoulzadegani
A0A1Y1S664 Enterospora canceri
A9CRJ3 Enterocytozoon bieneusi
S7W8I7 Spraguea lophii
A0A0R0M7T3 Pseudoloma neurophilia
A0A850MMV7 Candidatus Helarchaeota archaeon
CfP3-15 0445
A0A369SI02 Trichoplax sp. H2
A0A812DZM0 Sepia pharaonis
TRINITY DN14133 c1 g1 i1.p1 Pseudotrichonympha grassii
A0A2V1B0Y8 Candida haemuloni
Q6BZD6 Debaryomyces hansenii
A0A1E3PR62 Nadsonia fulvescens var. elongata DSM 6958
A0A2N1J7A6 Malassezia vespertilionis
A0A1Y2D0F9 Neocallimastix californiae
A0A1Y2CT12 Neocallimastix californiae
A0A1Y1XCZ7 Anaeromyces robustus
A0A1Y3NQ15 Piromyces sp.
B9WM58 Candida dubliniensis
A0A8H6F3V0 Candida albicans
A0A1D8PT80 Candida albicans
C4YM78 Candida albicans
G3B8G3 Candida tenuis
G3B8G4 Candida tenuis
A5E1E8 Lodderomyces elongisporus
C5M3H7 Candida tropicalis
A0A421JIW5 Spathaspora sp. JA1
A0A8H8DBP2 Candida metapsilosis
A0A8J5QGJ0 Candida subhashii

F

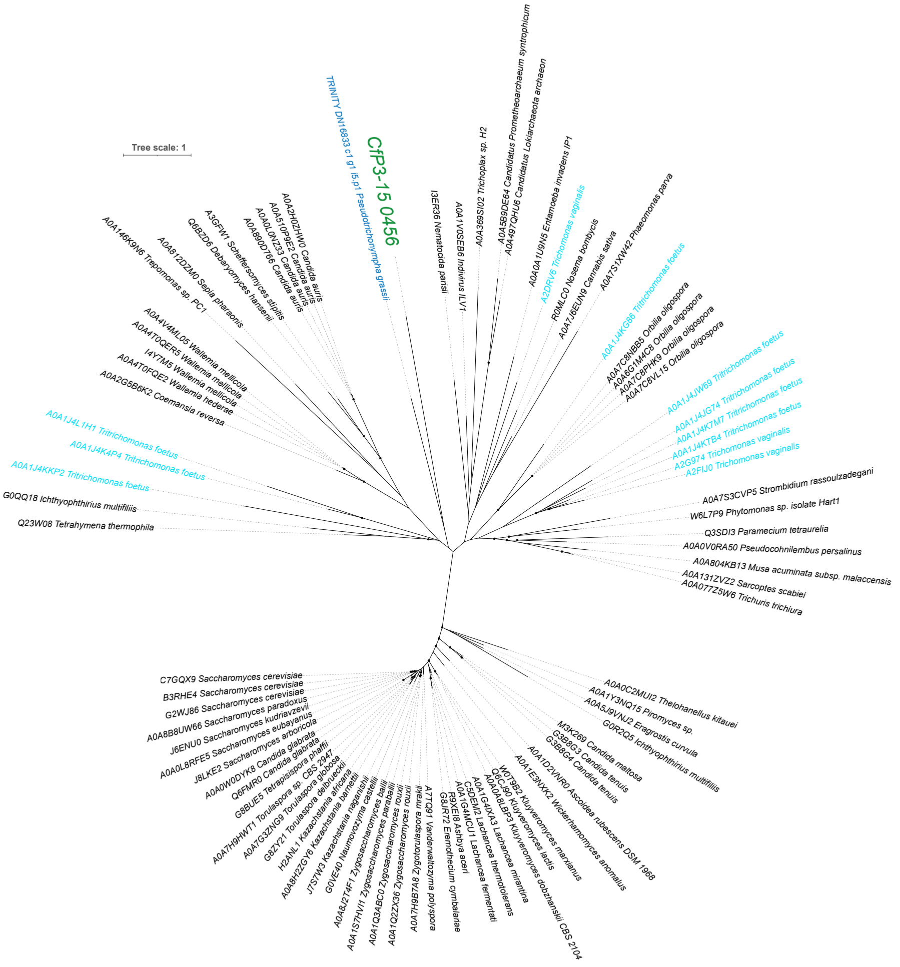
Tree scale: 1
TRINITY DN16833 c1 g1 i5.p1 Pseudotrichonympha grassii
CfP3-15 0456
A0A2H0ZHW0 Candida auris
A0A510P9E2 Candida auris
A0A0L0NZ33 Candida auris
A0A8Q0D766 Candida auris
A3GFW1 Scheffersomyces stipitis
Q6BZD6 Debaryomyces hansenii
A0A812DZM0 Sepia pharaonis
A0A146K8N6 Trepomonas sp. PC1
A0A4V4ML05 Wallemia mellicola
A0A4T0QER5 Wallemia mellicola
I4Y7M5 Wallemia mellicola
A0A4T0FQE2 Wallemia hederae
A0A2G5B6K2 Coemansia reversa
A0A1J4L1H1 Tritrichomonas foetus
A0A1J4K4P4 Tritrichomonas foetus
A0A1J4KKP2 Tritrichomonas foetus
G0QQ18 Ichthyophthirius multifiliis
Q23W08 Tetrahymena thermophila
I3ER36 Nematocida parisii
A0A1V0SEB6 Indivirus ILV1
A0A369SI02 Trichoplax sp. H2
A0A5B9DE64 Candidatus Prometheoarchaeum syntrophicum
A0A497QHU6 Candidatus Lokiarchaeota archaeon
A0A0A1U9N5 Entamoeba invadens IP1
A2DRV6 Trichomonas vaginalis
R0MLC0 Nosema bombycis
A0A7J6EUN9 Cannabis sativa
A0A7S1XW42 Phaeomonas parva
A0A1J4KG36 Tritrichomonas foetus
A0A7C8NBB5 Orbilia oligospora
A0A6G1M4C8 Orbilia oligospora
A0A7C8PHK9 Orbilia oligospora
A0A7C8VL15 Orbilia oligospora
A0A1J4JW69 Tritrichomonas foetus
A0A1J4JG74 Tritrichomonas foetus
A0A1J4K7M7 Tritrichomonas foetus
A0A1J4KTB4 Tritrichomonas foetus
A2G974 Trichomonas vaginalis
A2FIJ0 Trichomonas vaginalis
A0A7S3CVP5 Strombidium rassoulzadegani
W6L7P9 Phytomonas sp. isolate Hart1
Q3SDI3 Paramecium tetraurelia
A0A0V0RA50 Pseudocohnilembus persalinus
A0A804KB13 Musa acuminata subsp. malaccensis
A0A131ZVZ2 Sarcoptes scabiei
A0A077Z5W6 Trichuris trichiura
A0A0C2MUI2 Thelohanellus kitauei
A0A1Y3NQ15 Piromyces sp.
A0A5J9VNJ2 Eragrostis curvula
G0R2Q5 Ichthyophthirius multifiliis
M3K269 Candida maltosa
G3B8G3 Candida tenuis
G3B8G4 Candida tenuis
A0A1D2VNR0 Ascoidea rubescens DSM 1968
A0A1E3NXK2 Wickerhamomyces anomalus
W0TB82 Kluyveromyces marxianus
Q6CJ90 Kluyveromyces lactis
A0A0A8LBP3 Kluyveromyces dobzhanskii CBS 2104
A0A1G4JXA3 Lachancea mirantina
C5DM32 Lachancea thermotolerans
A0A1G4MCU1 Lachancea fermentati
R9XE18 Ashbya aceri
G8JR72 Eremothecium cymbalariae
A7TQ91 Vanderwaltozyma polyspora
A0A7H9B7A8 Zygotorulaspora mrakii
A0A1Q2ZX36 Zygosaccharomyces rouxii
A0A1Q3ABC0 Zygosaccharomyces parabailii
A0A1S7HVI1 Zygosaccharomyces bailii
A0A8J2T4F1 Zygosaccharomyces bailii
G0VE40 Naumovozyma castellii
J7S7W3 Kazachstania naganishii
A0A8H2ZGY6 Kazachstania africana
H2ANL1 Kazachstania africana
G8ZY21 Torulaspora delbrueckii
A0A7G3ZNG9 Torulaspora globosa
A0A7H9HWT1 Torulaspora sp. CBS 2947
G8BUE5 Tetrapisispora phaffii
Q6FMR0 Candida glabrata
A0A0W0DYK8 Candida glabrata
J8LKE2 Saccharomyces arboricola
A0A0L8RFE5 Saccharomyces eubayanus
J6ENU0 Saccharomyces kudriavzevii
A0A8B8UW66 Saccharomyces paradoxus
G2WJ86 Saccharomyces cerevisiae
B3RHE4 Saccharomyces cerevisiae
C7GQX9 Saccharomyces cerevisiae

G

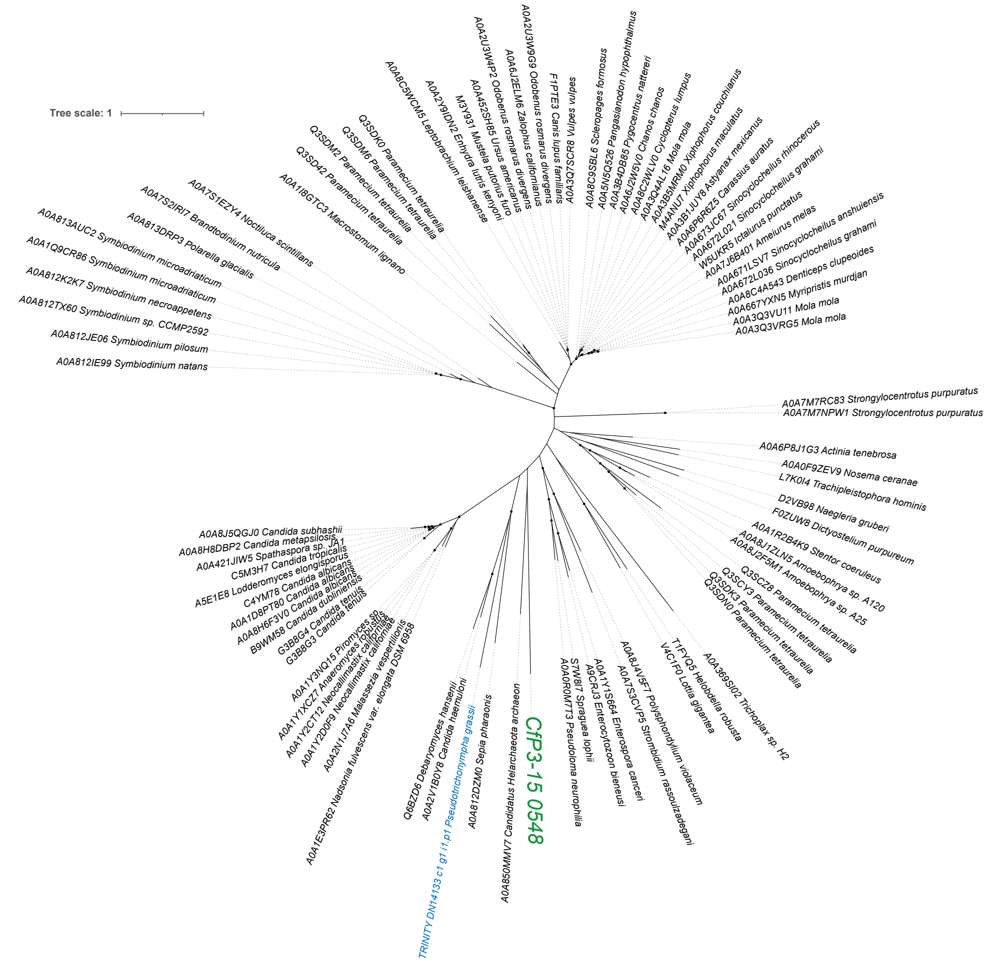
Tree scale: 1
CfP3-15 0548
TRINITY DN14133 c1 g1 i1.p1 Pseudotrichonympha grassii
A0A7M7RC83 Strongylocentrotus purpuratus
A0A7M7NPW1 Strongylocentrotus purpuratus
A0A812IE99 Symbiodinium natans
A0A812JE06 Symbiodinium pilosum
A0A3Q3VRG5 Mola mola
A0A3Q3VU11 Mola mola

H

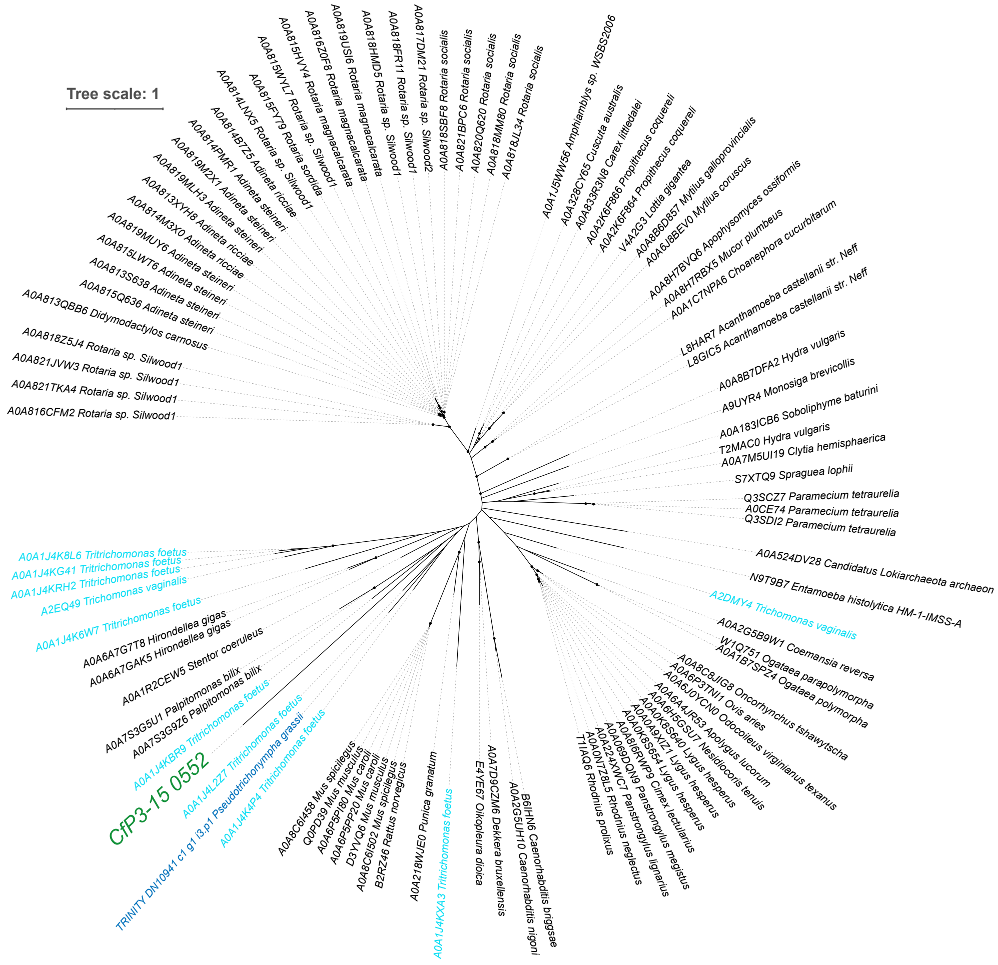
Tree scale: 1
CfP3-15 0552
TRINITY DN10941 c1 g1 i3.p1 Pseudotrichonympha grassii
A0A816CFM2 Rotaria sp. Silwood1
A0A821TKA4 Rotaria sp. Silwood1
A0A821JVW3 Rotaria sp. Silwood1
A0A818Z5J4 Rotaria sp. Silwood1
A0A524DV28 Candidatus Lokiarchaeota archaeon
N9T9B7 Entamoeba histolytica HM-1-IMSS-A
A2DMY4 Trichomonas vaginalis
S7XTQ9 Spraguea lophii
Q3SCZ7 Paramecium tetraurelia
A0CE74 Paramecium tetraurelia
Q3SDI2 Paramecium tetraurelia

I

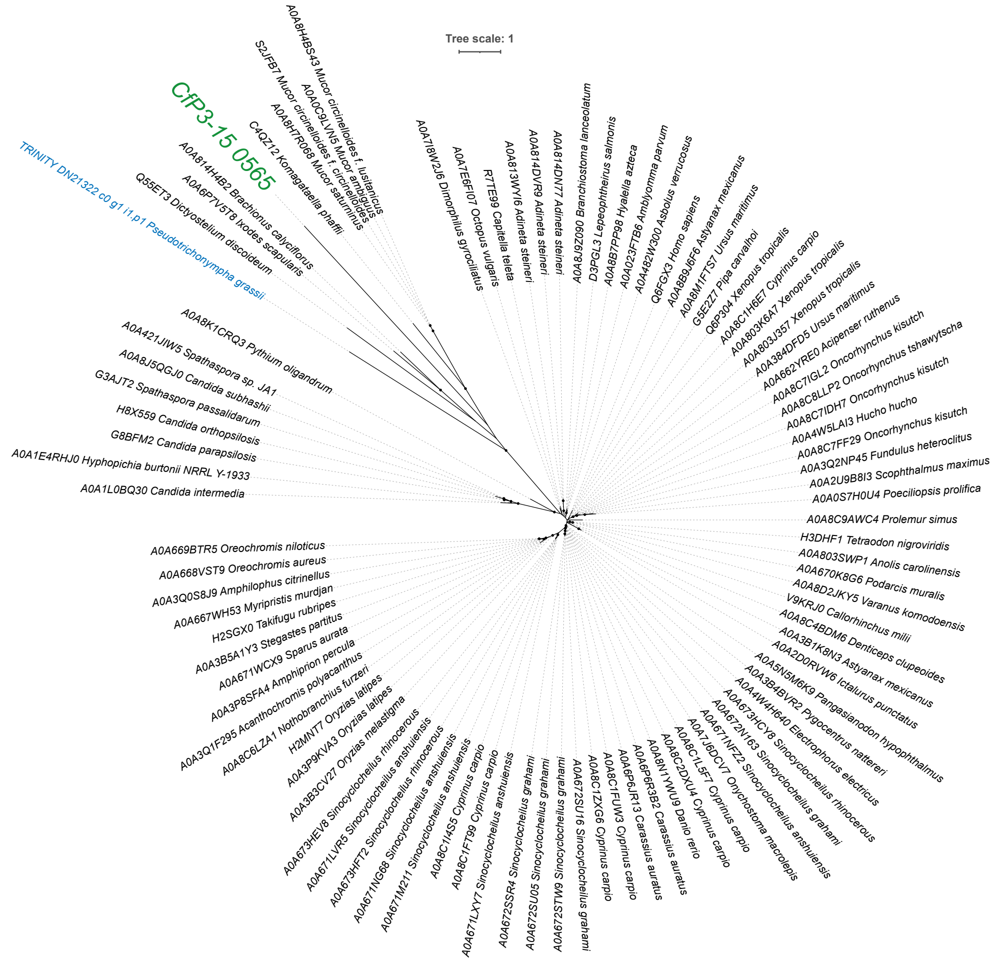
Tree scale: 1
CfP3-15 0565
TRINITY DN21322 c0 g1 i1.p1 Pseudotrichonympha grassii
A0A8H4BS43 Mucor circinelloides f. lusitanicus
A0A0C9LVN5 Mucor ambiguus
S2JFB7 Mucor circinelloides f. circinelloides
A0A8H7R068 Mucor saturninus
C4QZ12 Komagataella phaffii
A0A814H4B2 Brachionus calyciflorus
A0A6P7V5T8 Ixodes scapularis
Q55ET3 Dictyostelium discoideum
A0A7I8W2J6 Dimorphilus gyrociliatus
A0A7E6FI07 Octopus vulgaris
R7TE99 Capitella teleta
A0A813WYI6 Adineta steineri
A0A814DVR9 Adineta steineri
A0A814DN77 Adineta steineri
A0A8J9Z090 Branchiostoma lanceolatum
D3PGL3 Lepeophtheirus salmonis
A0A8B7PP98 Hyalella azteca
A0A023FTB6 Amblyomma parvum
A0A482W300 Asbolus verrucosus
Q6FGX3 Homo sapiens
A0A8B9J6F6 Astyanax mexicanus
A0A8M1FTS7 Ursus maritimus
G5E227 Pipa carvalhoi
Q6P304 Xenopus tropicalis
A0A8C1H6E7 Cyprinus carpio
A0A803K6A7 Xenopus tropicalis
A0A803J357 Xenopus tropicalis
A0A384DFD5 Ursus maritimus
A0A662YRE0 Acipenser ruthenus
A0A8C7IGL2 Oncorhynchus kisutch
A0A8C8LLP2 Oncorhynchus tshawytscha
A0A8C7IDH7 Oncorhynchus kisutch
A0A4W5LAI3 Hucho hucho
A0A8C7FF29 Oncorhynchus kisutch
A0A3Q2NP45 Fundulus heteroclitus
A0A2U9B8I3 Scophthalmus maximus
A0A0S7H0U4 Poeciliopsis prolifica
A0A8C9AWC4 Prolemur simus
H3DHF1 Tetraodon nigroviridis
A0A803SWP1 Anolis carolinensis
A0A670K8G6 Podarcis muralis
A0A8D2JKY5 Varanus komodoensis
V9KRJ0 Callorhinchus milii
A0A8C4BDM6 Denticeps clupeoides
A0A3B1K8N3 Astyanax mexicanus
A0A2D0RVW6 Ictalurus punctatus
A0A5N5M6K9 Pangasianodon hypophthalmus
A0A3B4BVR2 Pygocentrus nattereri
A0A4W4H640 Electrophorus electricus
A0A673HCY8 Sinocyclocheilus rhinocerous
A0A672N163 Sinocyclocheilus grahami
A0A671NFZ2 Sinocyclocheilus anshuiensis
A0A7J6DCV7 Onychostoma macrolepis
A0A8C1L5F7 Cyprinus carpio
A0A8C2DXU4 Cyprinus carpio
A0A8N1YWU9 Danio rerio
A0A6P6R3B2 Carassius auratus
A0A6P6JR13 Carassius auratus
A0A8C1FUW3 Cyprinus carpio
A0A8C1ZXG6 Cyprinus carpio
A0A672SU16 Sinocyclocheilus grahami
A0A672STM9 Sinocyclocheilus grahami
A0A672SU05 Sinocyclocheilus grahami
A0A672SSR4 Sinocyclocheilus grahami
A0A671LXY7 Sinocyclocheilus anshuiensis
A0A8C1FT99 Cyprinus carpio
A0A8C1I4S5 Cyprinus carpio
A0A671M211 Sinocyclocheilus anshuiensis
A0A671NG68 Sinocyclocheilus anshuiensis
A0A673HFT2 Sinocyclocheilus rhinocerous
A0A671LVR5 Sinocyclocheilus anshuiensis
A0A673HEV8 Sinocyclocheilus rhinocerous
A0A3B3CV27 Oryzias melastigma
A0A3P9KVA3 Oryzias latipes
H2MNT7 Oryzias latipes
A0A8C6LZA1 Nothobranchius furzeri
A0A3Q1F295 Acanthochromis polyacanthus
A0A3P8SFA4 Amphiprion percula
A0A671WCX9 Sparus aurata
A0A3B5A1Y3 Stegastes partitus
H2SGX0 Takifugu rubripes
A0A667WH53 Myripristis murdjan
A0A3Q0S8J9 Amphilophus citrinellus
A0A668VST9 Oreochromis aureus
A0A669BTR5 Oreochromis niloticus
A0A8K1CRQ3 Pythium oligandrum
A0A421JIW5 Spathaspora sp. JA1
A0A8J5QGJ0 Candida subhashii
G3AJT2 Spathaspora passalidarum
H8X559 Candida orthopsilosis
G8BFM2 Candida parapsilosis
A0A1E4RHJ0 Hyphopichia burtonii NRRL Y-1933
A0A1L0BQ30 Candida intermedia

J

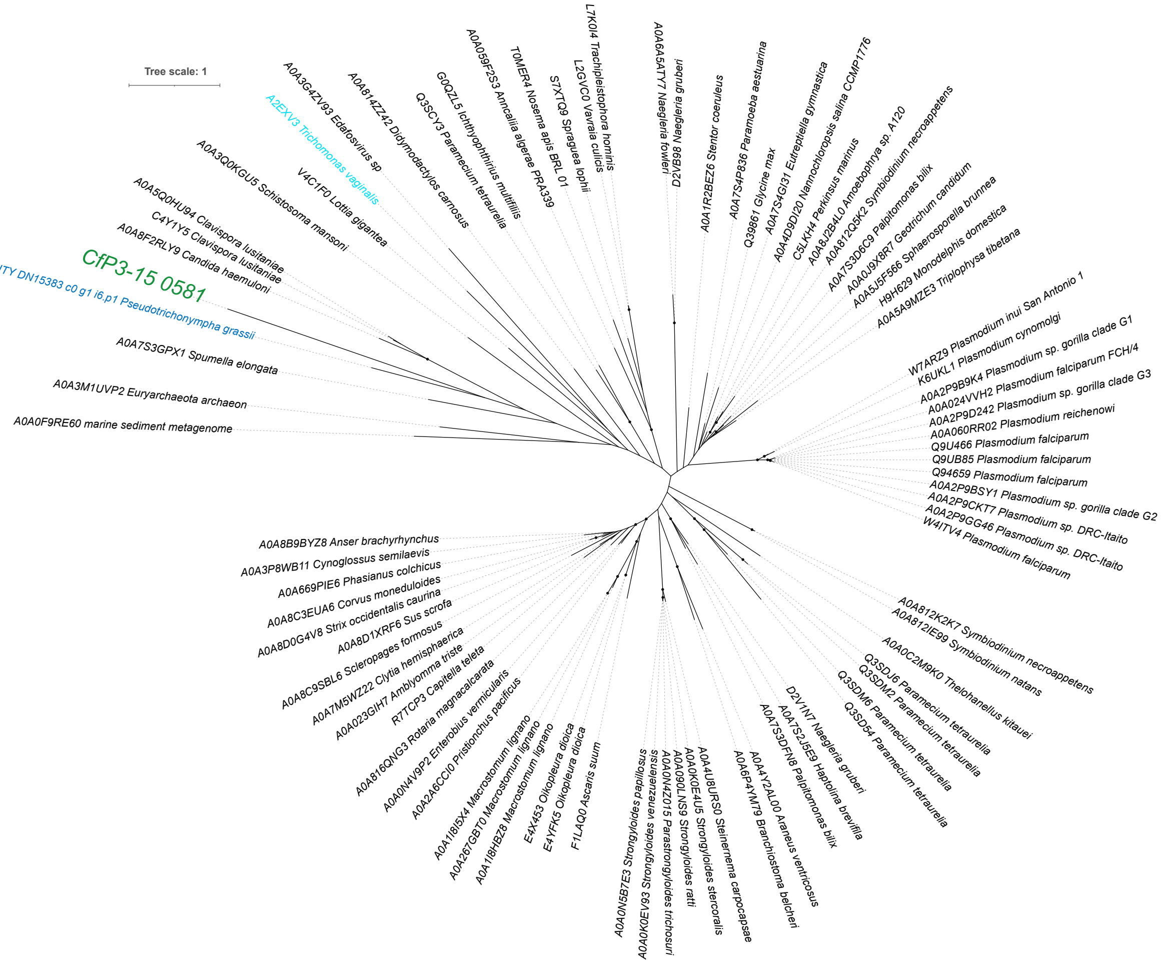
Tree scale: 1
CfP3-15 0581
TRINITY DN15383 c0 g1 i6.p1 Pseudotrichonympha grassii
A2EXV3 Trichomonas vaginalis
A0A3G4ZV93 Edafosvirus sp
A0A814ZZ42 Didymodactylos carnosus
Q3SCY3 Paramecium tetraurelia
G0QZL5 Ichthyophthirius multifiliis
A0A059F2S3 Anncaliia algerae PRA339
T0MER4 Nosema apis BRL 01
S7XTQ9 Spraguea lophii
L2GVC0 Vavraia culicis
L7K0I4 Trachipleistophora hominis
A0A6A5ATY7 Naegleria fowleri
D2VB98 Naegleria gruberi
A0A1R2BEZ6 Stentor coeruleus
A0A7S4P836 Paramoeba aestuarina
Q39861 Glycine max
A0A7S4GI31 Eutreptiella gymnastica
A0A4D9DI20 Nannochloropsis salina CCMP1776
C5LKH4 Perkinsus marinus
A0A8J2B4L0 Amoebophrya sp. A120
A0A812Q5K2 Symbiodinium necroappetens
A0A7S3D6C9 Palpitomonas bilix
A0A0J9X8R7 Geotrichum candidum
A0A5J5F566 Sphaerosporella brunnea
H9H629 Monodelphis domestica
A0A5A9MZE3 Triplophysa tibetana
W7ARZ9 Plasmodium inui San Antonio 1
K6UKL1 Plasmodium cynomolgi
A0A2P9B9K4 Plasmodium sp. gorilla clade G1
A0A024VVH2 Plasmodium falciparum FCH/4
A0A2P9D242 Plasmodium sp. gorilla clade G3
A0A060RR02 Plasmodium reichenowi
Q9U466 Plasmodium falciparum
Q9UB85 Plasmodium falciparum
Q94659 Plasmodium falciparum
A0A2P9BSY1 Plasmodium sp. gorilla clade G2
A0A2P9CKT7 Plasmodium sp. DRC-Itaito
A0A2P9GG46 Plasmodium sp. DRC-Itaito
W4ITV4 Plasmodium falciparum
A0A812K2K7 Symbiodinium necroappetens
A0A812IE99 Symbiodinium natans
A0A0C2M9K0 Thelohanellus kitauei
Q3SDJ6 Paramecium tetraurelia
Q3SDM2 Paramecium tetraurelia
Q3SDM6 Paramecium tetraurelia
Q3SD54 Paramecium tetraurelia
D2V1N7 Naegleria gruberi
A0A7S2JSE9 Haptolina brevifila
A0A7S3DFN8 Palpitomonas bilix
A0A4Y2AL00 Araneus ventricosus
A0A6P4YM79 Branchiostoma belcheri
A0A4U8UR50 Steinernema carpocapsae
A0A0K0E4U5 Strongyloides stercoralis
A0A090LNS9 Strongyloides ratti
A0A0N4Z015 Parastrongyloides trichosuri
A0A0K0EV93 Strongyloides venezuelensis
A0A0N5B7E3 Strongyloides papillosus
F1LAQ0 Ascaris suum
E4YFK5 Oikopleura dioica
E4X453 Oikopleura dioica
A0A1I8HBZ8 Macrostomum lignano
A0A267GBT0 Macrostomum lignano
A0A1I8I5X4 Macrostomum lignano
A0A2A6CCI0 Pristionchus pacificus
A0A0N4V9P2 Enterobius vermicularis
A0A816QNG3 Rotaria magnacalcarata
R7TCP3 Capitella teleta
A0A023GIH7 Amblyomma triste
A0A7M5WZ22 Clytia hemisphaerica
A0A8C9SBL6 Scleropages formosus
A0A8D1XRF6 Sus scrofa
A0A8D0G4V8 Strix occidentalis caurina
A0A8C3EUA6 Corvus moneduloides
A0A669PIE6 Phasianus colchicus
A0A3P8WB11 Cynoglossus semilaevis
A0A8B9BYZ8 Anser brachyrhynchus
A0A0F9RE60 marine sediment metagenome
A0A3M1UVP2 Euryarchaeota archaeon
A0A7S3GPX1 Spumella elongata
A0A8F2RLY9 Candida haemuloni
C4Y1Y5 Clavispora lusitaniae
A0A5Q0HU94 Clavispora lusitaniae
A0A3Q0KGU5 Schistosoma mansoni
V4C1F0 Lottia gigantea

K

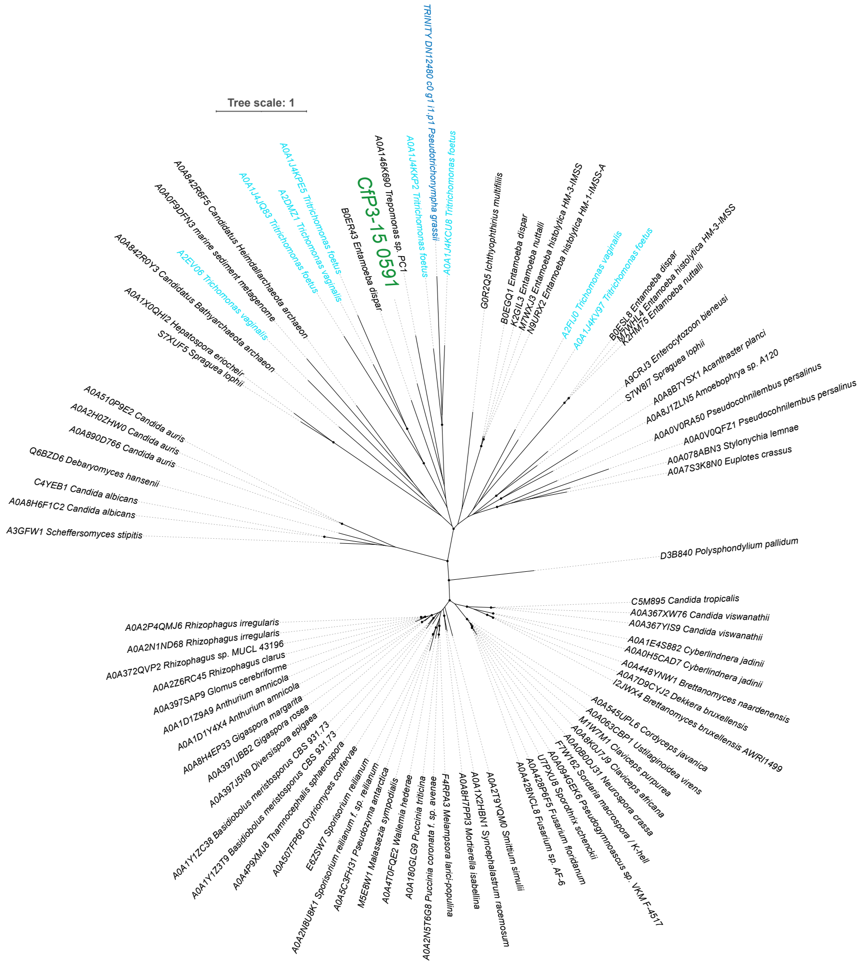


L

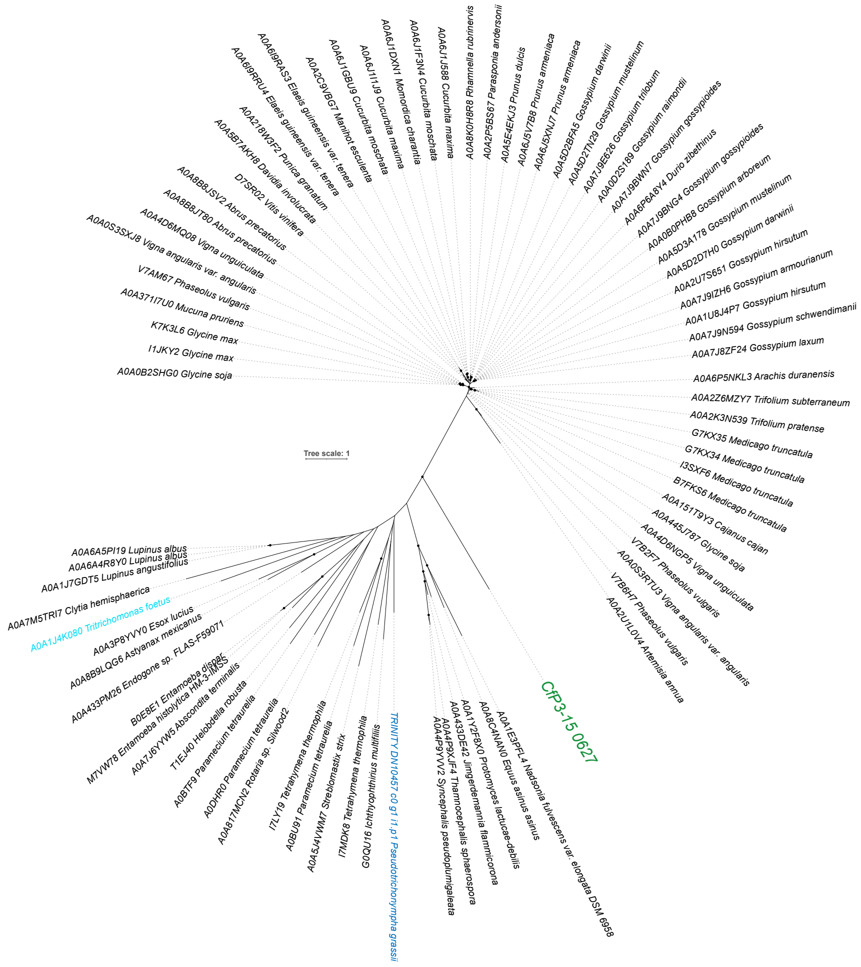

M

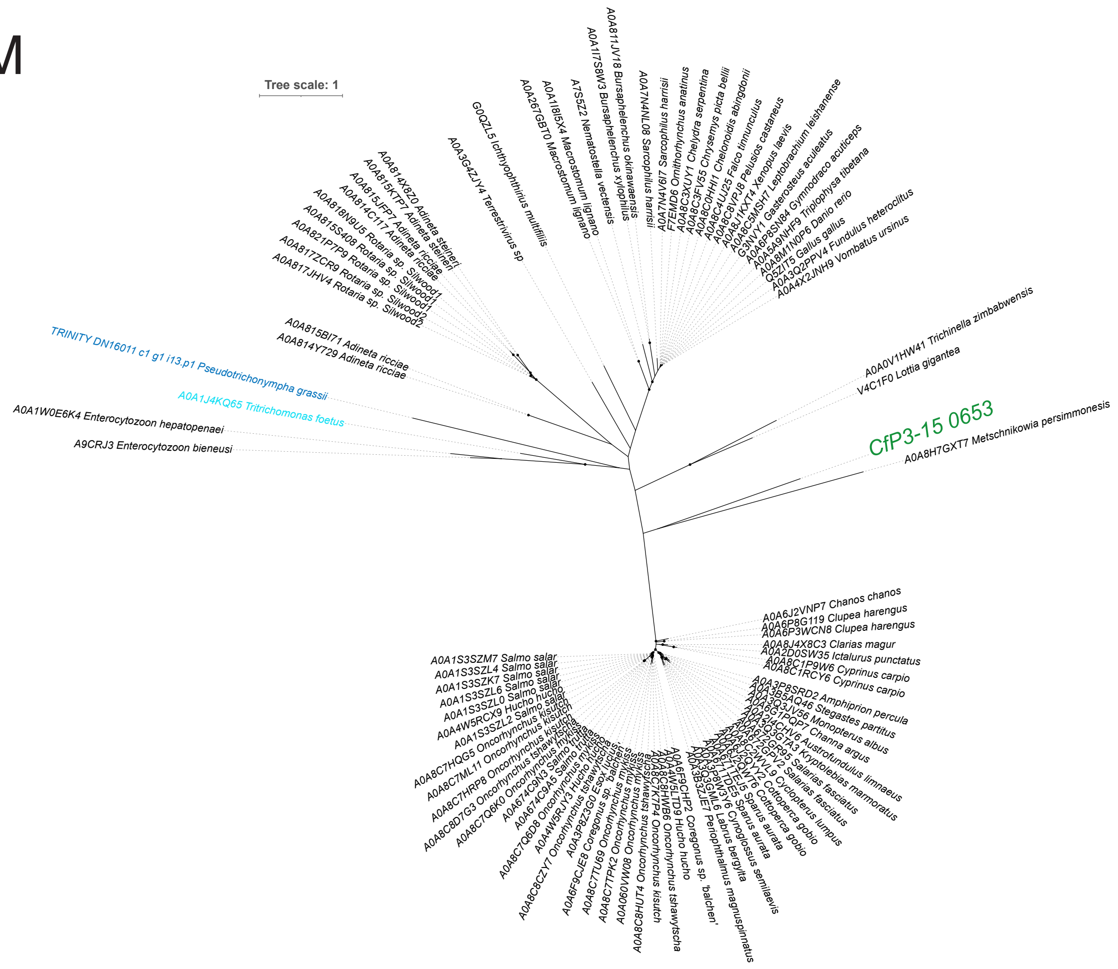
Tree scale: 1
A0A3G4ZJY4 Terrestrivirus sp
G0QZL5 Ichthyophthirius multifiliis
A0A814X8Z0 Adineta steineri
A0A815KTP7 Adineta steineri
A0A814C117 Adineta ricciae
A0A818N9U5 Rotaria sp. Silwood1
A0A815S408 Rotaria sp. Silwood1
A0A821P7P9 Rotaria sp. Silwood2
A0A817ZCR9 Rotaria sp. Silwood1
A0A817JHV4 Rotaria sp. Silwood2
A0A815BI71 Adineta ricciae
A0A814Y729 Adineta ricciae
TRINITY DN16011 c1 g1 i13.p1 Pseudotrichonympha grassii
A0A1J4KQ65 Tritrichomonas foetus
A0A1W0E6K4 Enterocytozoon hepatopenaei
A9CRJ3 Enterocytozoon bieneusi
A0A257GBT0 Macrostomum lignano
A0A1I8I5X4 Macrostomum lignano
A7S5Z2 Nematostella vectensis
A0A1I7S8W3 Bursaphelenchus xylophilus
A0A811JV18 Bursaphelenchus okinawaensis
A0A7N4NL08 Sarcophilus harrisii
A0A7N4V6I7 Sarcophilus harrisii
F7EMD6 Ornithorhynchus anatinus
A0A8C3XUY1 Chelydra serpentina
A0A8C3F655 Chrysemys picta bellii
A0A8C0HH11 Chelonoidis abingdonii
A0A8C4U2J5 Falco tinnunculus
A0A8J1KXT4 Xenopus laevis
A0A8C6VP38 Pelusios castaneus
A0A8C5MSH7 Leptobrachium leishanense
Gasterosteus aculeatus
Gymnodraco acuticeps
Triplophysa tibetana
Danio rerio
Q5ZIT5 Gallus gallus
A0A3Q2PPV4 Fundulus heteroclitus
A0A4X2JNH9 Vombatus ursinus
A0A0V1HW41 Trichinella zimbabwensis
V4C1F0 Lottia gigantea
CfP3-15 0653
A0A8H7GXT7 Metschnikowia persimmonesis
A0A6J2VNP7 Chanos chanos
A0A6P8G119 Clupea harengus
A0A6P3WCN8 Clupea harengus
A0A8J4X8C3 Clarias magur
A0A2D0SW35 Ictalurus punctatus
A0A8C1P9W6 Cyprinus carpio
A0A8C1RCY6 Cyprinus carpio
A0A1S3SZM7 Salmo salar
A0A1S3SZL4 Salmo salar
A0A1S3SZK7 Salmo salar
A0A1S3SZL6 Salmo salar
A0A1S3SZL0 Salmo salar
A0A4W5RCX9 Hucho hucho
A0A1S3SZL2 Salmo salar
A0A8C7HQG5 Oncorhynchus kisutch
A0A8C7ML11 Oncorhynchus kisutch
Amphiprion percula
Stegastes partitus
Monopterus albus
Channa argus
Austrofundulus limnaeus
Kryptolebias marmoratus
Salarias fasciatus
Cyclopterus lumpus
Cottoperca gobio
Sparus aurata
Cynoglossus semilaevis
Labrus bergylta
Periophthalmus magnuspinnatus
Coregonus sp. 'balchen'
Hucho hucho
Oncorhynchus tshawytscha
Oncorhynchus kisutch

N

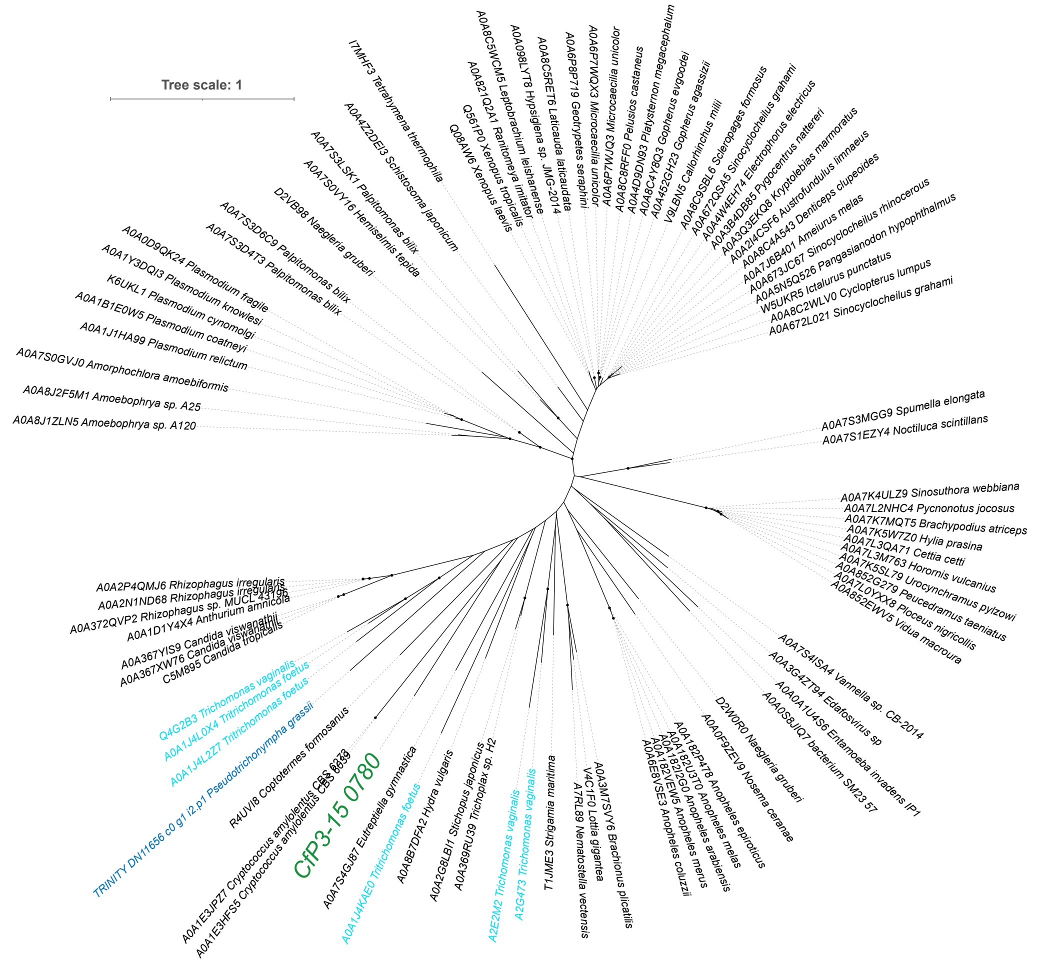
Tree scale: 1
I7MHF3 Tetrahymena thermophila
A0A4Z2DEI3 Schistosoma japonicum
A0A7S3LSK1 Palpitomonas bilix
A0A7S0VY16 Hemiselmis tepida
D2VB98 Naegleria gruberi
A0A7S3D6C9 Palpitomonas bilix
A0A7S3D4T3 Palpitomonas bilix
A0A0D9QK24 Plasmodium fragile
A0A1Y3DQI3 Plasmodium knowlesi
K6UKL1 Plasmodium cynomolgi
A0A1B1E0W5 Plasmodium coatneyi
A0A1J1HA99 Plasmodium relictum
A0A7S0GVJ0 Amorphochlora amoebiformis
A0A8J2F5M1 Amoebophrya sp. A25
A0A8J1ZLN5 Amoebophrya sp. A120
A0A8C5WCM5 Leptobrachium leishanense
A0A098LYT8 Hypsiglena sp. JMG-2014
A0A8C5RET6 Laticauda laticaudata
A0A6P8P719 Geotrypetes seraphini
A0A6P7WQX3 Microcaecilia unicolor
A0A6P7WJQ3 Microcaecilia unicolor
A0A8C8RFF0 Pelusios castaneus
A0A4D9DN93 Platysternon megacephalum
A0A4C4Y6Q3 Gopherus evgoodei
A0A452GH23 Gopherus agassizii
V9LBN5 Callorhinchus milii
A0A8C9SBL6 Scleropages formosus
A0A672QSA5 Sinocyclocheilus grahami
A0A4W4EH74 Electrophorus electricus
A0A3B4DB65 Pygocentrus nattereri
A0A3Q3EKQ8 Kryptolebias marmoratus
A0A2I4CSF6 Austrofundulus limnaeus
A0A8C4A543 Denticeps clupeoides
A0A7J6B401 Ameiurus melas
A0A673JC67 Sinocyclocheilus rhinocerous
A0A5N5Q526 Pangasianodon hypophthalmus
W5UKR5 Ictalurus punctatus
A0A8C2WLV0 Cyclopterus lumpus
A0A672L021 Sinocyclocheilus grahami
A0A821Q2A1 Ranitomeya imitator
Q561P0 Xenopus tropicalis
Q08AW6 Xenopus laevis
A0A7S3MGG9 Spumella elongata
A0A7S1EZY4 Noctiluca scintillans
A0A7K4ULZ9 Sinosuthora webbiana
A0A7L2NHC4 Pycnonotus jocosus
A0A7K7MQT5 Brachypodius atriceps
A0A7K5W7Z0 Hylia prasina
A0A7L3QA71 Cettia cetti
A0A7L3M763 Horornis vulcanius
A0A7K5SL79 Urocynchramus pylzowi
A0A852G279 Peucedramus taeniatus
A0A7L0YXX8 Ploceus nigricollis
A0A852EWV5 Vidua macroura
A0A7S4ISA4 Vannella sp. CB-2014
A0A3G4ZT94 Edafosvirus sp
A0A0A1U4S6 Entamoeba invadens IP1
A0A0S8JIQ7 bacterium SM23 57
D2W0R0 Naegleria gruberi
A0A0F9ZEV9 Nosema ceranae
A0A182P478 Anopheles epiroticus
A0A182U3T0 Anopheles melas
A0A182GNY0 Anopheles arabiensis
A0A182VEW5 Anopheles merus
A0A6E8W5E3 Anopheles coluzzii
A0A3M7SVY6 Brachionus plicatilis
V4C1F0 Lottia gigantea
A7RL89 Nematostella vectensis
T1JME3 Strigamia maritima
A2G473 Trichomonas vaginalis
A2E2M2 Trichomonas vaginalis
A0A369RU39 Trichoplax sp. H2
A0A2G8LBI1 Stichopus japonicus
A0A8B7DFA2 Hydra vulgaris
A0A7S4GJ87 Eutreptiella gymnastica
A0A1J4KAE0 Tritrichomonas foetus
CfP3-15 0780
A0A1E3JPZ7 Cryptococcus amylolentus CBS 6273
A0A1E3HFS5 Cryptococcus amylolentus CBS 6039
R4UVI8 Coptotermes formosanus
TRINITY DN11656 c0 g1 i2.p1 Pseudotrichonympha grassii
A0A1J4L2Z7 Tritrichomonas foetus
A0A1J4L0X4 Tritrichomonas foetus
Q4G2B3 Trichomonas vaginalis
A0A367YIS9 Candida viswanathii
A0A367XW76 Candida viswanathii
C5M895 Candida tropicalis
A0A1D1Y4X4 Anthurium amnicola
A0A372QVP2 Rhizophagus sp. MUCL 43196
A0A2N1ND68 Rhizophagus irregularis
A0A2P4QMJ6 Rhizophagus irregularis

O

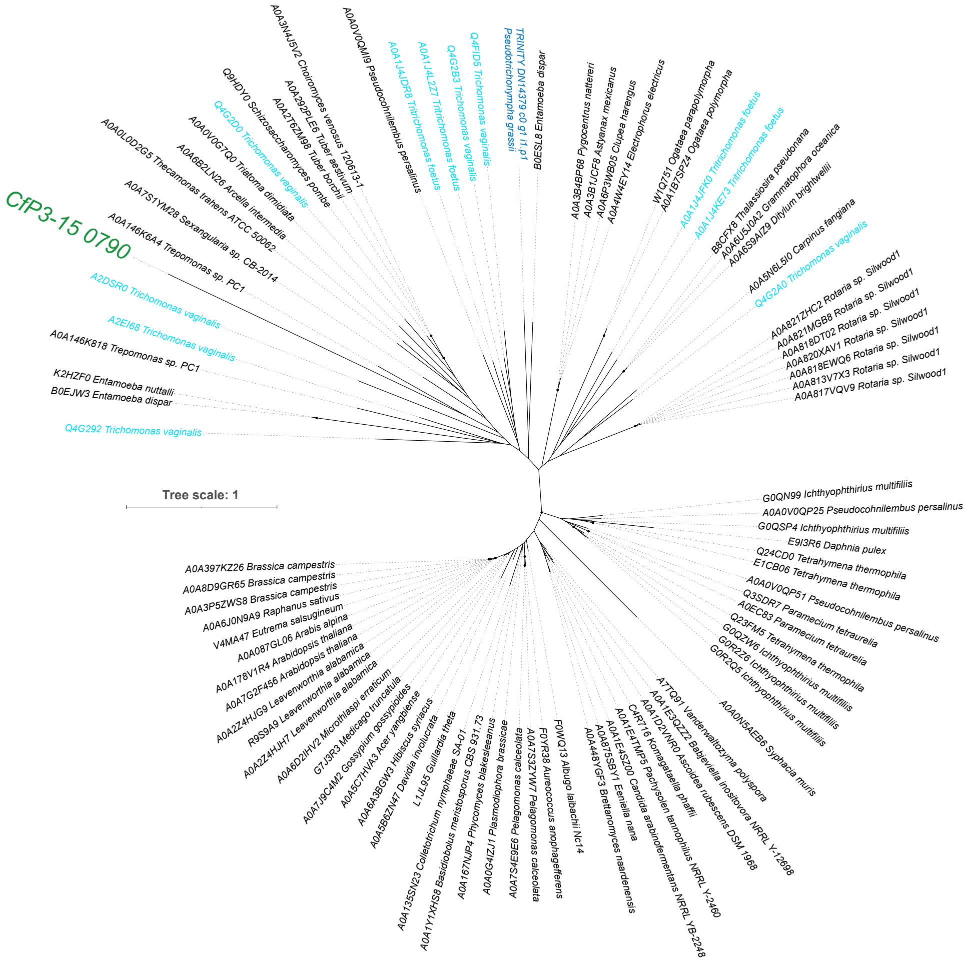

CfP3-15 0790
A0A3N4J5V2 Choiromyces venosus 120613-1
A0A0V0QM19 Pseudocohnilembus persalinus
Q9HDV0 Schizosaccharomyces pombe
A0A292PLE6 Tuber aestivum
A0A2T6ZM98 Tuber borchii
Q4G2D0 Trichomonas vaginalis
A0A0V0G7Q0 Triatoma dimidiata
A0A6B2LN26 Arcella intermedia
A0A0L0D2G5 Thecamonas trahens ATCC 50062
A0A7S1YM28 Sexangularia sp. CB-2014
A0A146K6A4 Trepomonas sp. PC1
A2DSR0 Trichomonas vaginalis
A2EI66 Trichomonas vaginalis
A0A146K818 Trepomonas sp. PC1
K2HZF0 Entamoeba nuttalli
B0EJW3 Entamoeba dispar
Q4G292 Trichomonas vaginalis
A0A1J4JDR8 Tritrichomonas foetus
A0A1J4L2Z7 Tritrichomonas foetus
Q4G2B3 Trichomonas vaginalis
Q4FID5 Trichomonas vaginalis
TRINITY DN14379 c0 g1 i1.p1
Pseudotrichonympha grassii
B0ESL8 Entamoeba dispar
A0A3B4BP68 Pygocentrus nattereri
A0A3B1JCF8 Astyanax mexicanus
A0A6P3WB05 Clupea harengus
A0A4W4EY14 Electrophorus electricus
W1Q751 Ogataea parapolymorpha
A0A1B7SPZ4 Ogataea polymorpha
A0A1J4JPK0 Tritrichomonas foetus
A0A1J4KE73 Tritrichomonas foetus
B8CFX8 Thalassiosira pseudonana
A0A6U5J0A2 Grammatophora oceanica
A0A6S9AI29 Ditylum brightwellii
A0A5N6L5I0 Carpinus fangiana
Q4G2A0 Trichomonas vaginalis
A0A821ZHC2 Rotaria sp. Silwood1
A0A821MGB8 Rotaria sp. Silwood1
A0A818DT02 Rotaria sp. Silwood1
A0A820XAV1 Rotaria sp. Silwood1
A0A818EWQ6 Rotaria sp. Silwood1
A0A813V7X3 Rotaria sp. Silwood1
A0A817VQV9 Rotaria sp. Silwood1
Tree scale: 1
G0QN99 Ichthyophthirius multifiliis
A0A0V0QP25 Pseudocohnilembus persalinus
G0QSP4 Ichthyophthirius multifiliis
E9I3R6 Daphnia pulex
Q24CD0 Tetrahymena thermophila
E1CB06 Tetrahymena thermophila
A0A0V0QP51 Pseudocohnilembus persalinus
Q3SDR7 Paramecium tetraurelia
A0EC83 Paramecium tetraurelia
Q23FM5 Tetrahymena thermophila
G0QZW6 Ichthyophthirius multifiliis
G0R226 Ichthyophthirius multifiliis
G0R2Q5 Ichthyophthirius multifiliis
A0A0N5AEB6 Syphacia muris
A7TQ91 Vanderwaltozyma polyspora
A0A1E3QZZ2 Babjeviella inositovora NRRL Y-12698
A0A1D2VNR0 Ascoidea rubescens DSM 1968
C4R716 Komagataella phaffii
A0A1E4TMP5 Pachysolen tannophilus NRRL Y-2460
A0A1E4SZ00 Candida arabinofermentans NRRL YB-2248
A0A875SBY1 Eeniella nana
A0A448YGF3 Brettanomyces naardenensis
F0WQ13 Albugo laibachii Nc14
F0YR38 Aureococcus anophagefferens
A0A7S3ZYW7 Pelagomonas calceolata
A0A7S4E9E6 Pelagomonas calceolata
A0A0G4IZJ1 Plasmodiophora brassicae
A0A167NJP4 Phycomyces blakesleeanus
A0A1Y1XHS8 Basidiobolus meristosporus CBS 931.73
A0A135SN23 Colletotrichum nymphaeae SA-01
L1J9L96 Guillardia theta
A0A5B6ZN47 Davidia involucrata
A0A6A3BGW3 Hibiscus syriacus
A0A5C7HVA3 Acer yangbiense
A0A7J9C4M2 Gossypium gossypioides
G7J3R3 Medicago truncatula
A0A6D2IHV2 Microthlaspi erraticum
A0A2Z4HJH7 Leavenworthia alabamica
R9S9A9 Leavenworthia alabamica
A0A2Z4HJG9 Leavenworthia alabamica
A0A7G2F456 Arabidopsis thaliana
A0A178V1R4 Arabidopsis thaliana
A0A087GL06 Arabis alpina
V4MA47 Eutrema salsugineum
A0A6J0N9A9 Raphanus sativus
A0A3P5ZWS8 Brassica campestris
A0A8D9GR65 Brassica campestris
A0A397KZ26 Brassica campestris


P

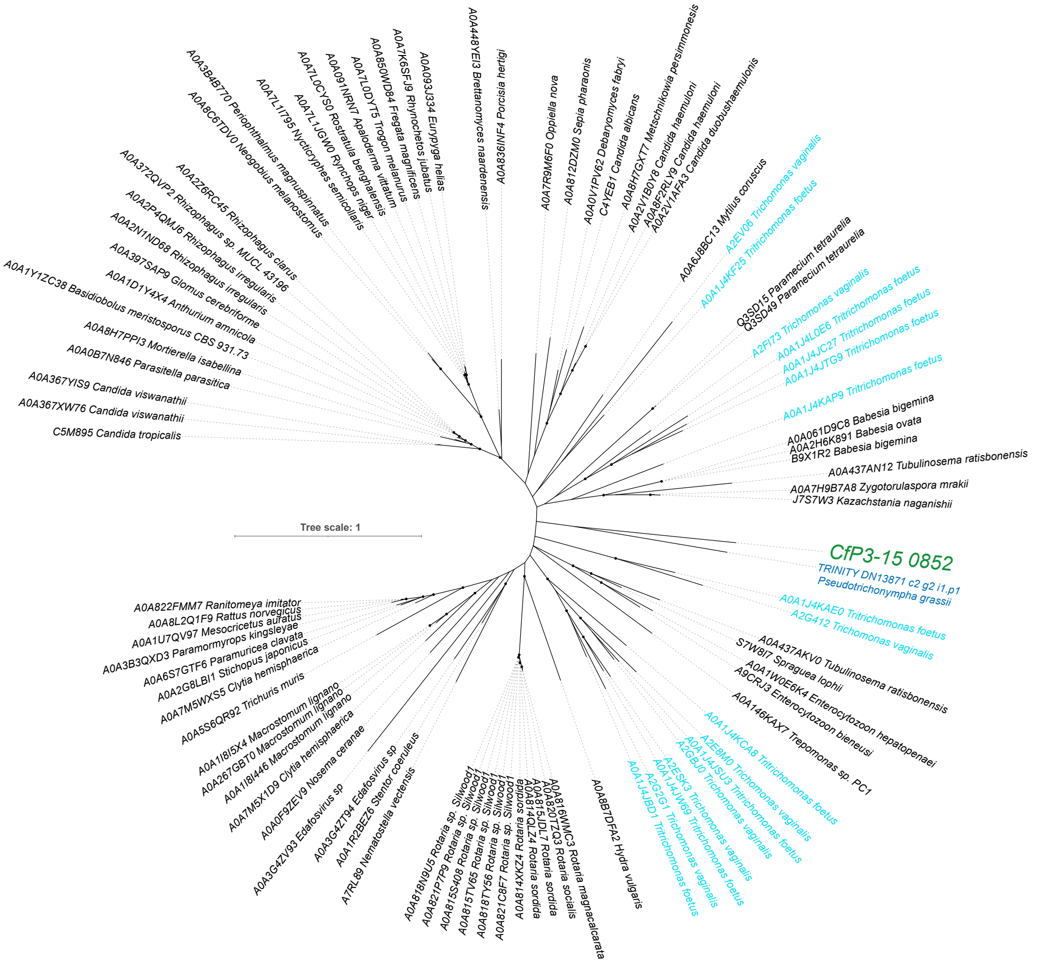

A0A448YEI3 Brettanomyces naardenensis
A0A836INF4 Porcisia hertigi
A0A093J334 Eurypyga helias
A0A7K6SFJ9 Rhynochetos jubatus
A0A850WD84 Fregata magnificens
A0A7L0DYT5 Trogon melanurus
A0A091NRN7 Apaloderma vittatum
A0A7L0CYS0 Rostratula benghalensis
A0A7L1JGW0 Rynchops niger
A0A7L1I795 Nyctibius semicollaris
A0A3B4B770 Periophthalmus magnuspinnatus
A0A8C6TDV0 Neogobius melanostomus
A0A372QVP2 Rhizophagus sp. MUCL 43196
A0A2Z6RC45 Rhizophagus clarus
A0A2P4QMJ6 Rhizophagus irregularis
A0A2N1ND68 Rhizophagus irregularis
A0A397SAP9 Glomus cerebriforme
A0A1D1Y4X4 Anthurium amnicola
A0A1Y1ZC38 Basidiobolus meristosporus CBS 931.73
A0A8H7PPI3 Mortierella isabellina
A0A0B7N846 Parasitella parasitica
A0A367YIS9 Candida viswanathii
A0A367XW76 Candida viswanathii
C5M895 Candida tropicalis
A0A7R9M6F0 Oppiella nova
A0A812DZM0 Sepia pharaonis
A0A0V1PV62 Debaryomyces fabryi
C4YEB1 Candida albicans
A0A8H7GXT7 Metschnikowia persimmonesis
A0A2V1B0Y8 Candida haemuloni
A0A8F2RLY9 Candida haemuloni
A0A2V1AFA3 Candida duobushaemulonis
A0A6J8BC13 Mytilus coruscus
A2EV08 Trichomonas vaginalis
A0A1J4KF25 Tritrichomonas foetus
Q3SD15 Paramecium tetraurelia
Q3SD49 Paramecium tetraurelia
A2FI73 Trichomonas vaginalis
A0A1J4L0E6 Tritrichomonas foetus
A0A1J4JC27 Tritrichomonas foetus
A0A1J4JTG9 Tritrichomonas foetus
A0A1J4KAP9 Tritrichomonas foetus
A0A061D9C8 Babesia bigemina
A0A2H6K891 Babesia ovata
B9X1R2 Babesia bigemina
A0A437AN12 Tubulinosema ratisbonensis
A0A7H9B7A8 Zygotorulaspora mrakii
J7S7W3 Kazachstania naganishii
Tree scale: 1
CfP3-15 0852
TRINITY DN13871 c2 g2 i1.p1
Pseudotrichonympha grassii
A0A1J4KAE0 Tritrichomonas foetus
A2G412 Trichomonas vaginalis
A0A437AKV0 Tubulinosema ratisbonensis
S7W8I7 Spraguea lophii
A0A1W0E6K4 Enterocytozoon hepatopenaei
A9CRJ3 Enterocytozoon bieneusi
A0A146KAX7 Trepomonas sp. PC1
A0A1J4KCA8 Tritrichomonas foetus
A2E8M0 Trichomonas vaginalis
A0A1J4JSU3 Tritrichomonas foetus
A2GBJ0 Trichomonas vaginalis
A2ESK3 Trichomonas vaginalis
A0A1J4JW69 Tritrichomonas foetus
A2G2G1 Trichomonas vaginalis
A0A1J4JBD1 Tritrichomonas foetus
A0A8B7DFA2 Hydra vulgaris
A0A816WNC3 Rotaria magnacalcarata
A0A820TZQ3 Rotaria socialis
A0A815UDL7 Rotaria sordida
A0A814QLZ4 Rotaria sordida
A0A814XKZ4 Rotaria sordida
A0A821C8F7 Rotaria sp. Silwood1
A0A818TY56 Rotaria sp. Silwood1
A0A815TV65 Rotaria sp. Silwood1
A0A815S408 Rotaria sp. Silwood1
A0A821P7P9 Rotaria sp. Silwood1
A0A818N9U5 Rotaria sp. Silwood1
A7RL89 Nematostella vectensis
A0A1R2BEZ6 Stentor coeruleus
A0A3G4ZT94 Edafosvirus sp
A0A3G4ZV93 Edafosvirus sp
A0A0F9ZEV9 Nosema ceranae
A0A7M5X1D9 Clytia hemisphaerica
A0A1I8I446 Macrostomum lignano
A0A267GBT0 Macrostomum lignano
A0A1I8I5X4 Macrostomum lignano
A0A5S6QR92 Trichuris muris
A0A7M5WXS5 Clytia hemisphaerica
A0A2G8LBI1 Stichopus japonicus
A0A6S7GTF6 Paramuricea clavata
A0A3B3QXD3 Paramormyrops kingsleyae
A0A1U7QV97 Mesocricetus auratus
A0A8L2Q1F9 Rattus norvegicus
A0A822FMM7 Ranitomeya imitator

Q

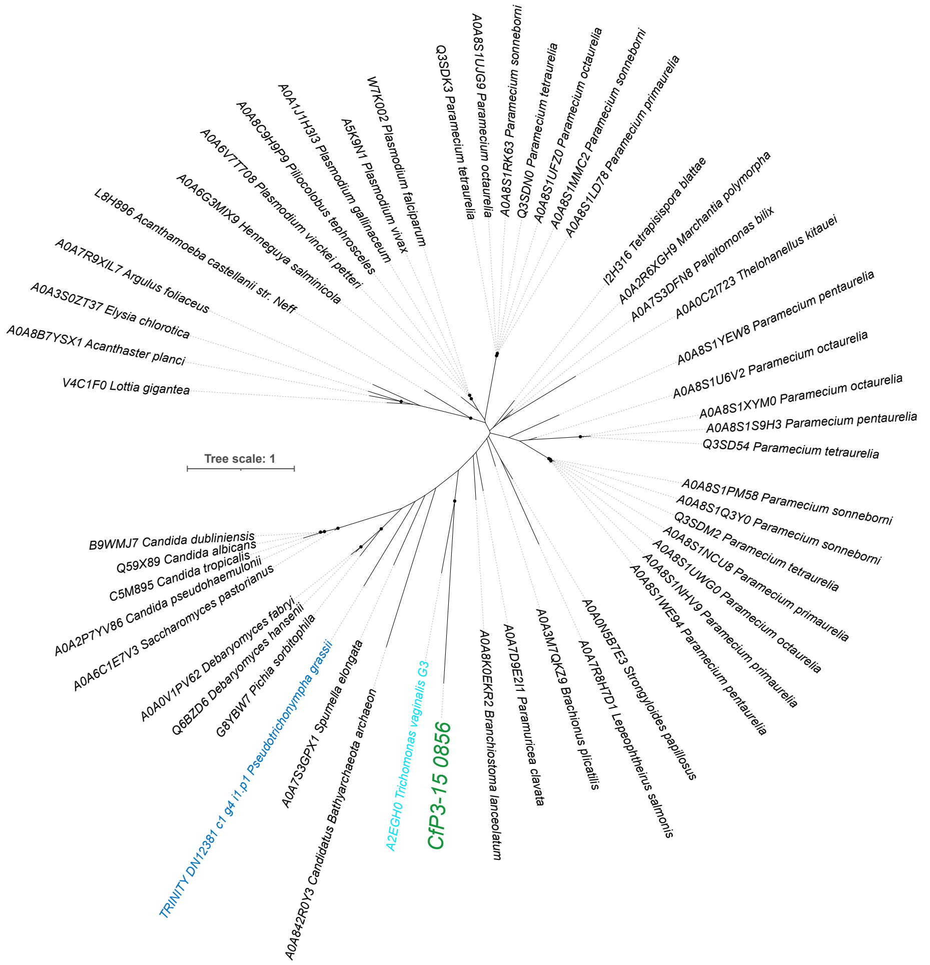

Tree scale: 1
Q3SDK3 Paramecium tetraurelia
A0A8S1UJG9 Paramecium octaurelia
A0A8S1RK63 Paramecium sonneborni
Q3SDN0 Paramecium tetraurelia
A0A8S1UFZ0 Paramecium octaurelia
A0A8S1MMC2 Paramecium sonneborni
A0A8S1LD78 Paramecium primaurelia
I2H316 Tetrapisispora blattae
A0A2R6XGH9 Marchantia polymorpha
A0A7S3DFN8 Palpitomonas bilix
A0A0C2I723 Thelohanellus kitauei
A0A8S1YEW8 Paramecium pentaurelia
A0A8S1U6V2 Paramecium octaurelia
A0A8S1XYM0 Paramecium octaurelia
A0A8S1S9H3 Paramecium pentaurelia
Q3SD54 Paramecium tetraurelia
A0A8S1PM58 Paramecium sonneborni
A0A8S1Q3Y0 Paramecium sonneborni
Q3SDM2 Paramecium tetraurelia
A0A8S1NCU8 Paramecium primaurelia
A0A8S1UWG0 Paramecium octaurelia
A0A8S1NHV9 Paramecium primaurelia
A0A8S1WE94 Paramecium pentaurelia
A0A0N5B7E3 Strongyloides papillosus
A0A7R8H7D1 Lepeophtheirus salmonis
A0A3M7QKZ9 Brachionus plicatilis
A0A7D9E2I1 Paramuricea clavata
A0A8K0EKR2 Branchiostoma lanceolatum
CfP3-15 0856
A2EGH0 Trichomonas vaginalis G3
A0A842R0Y3 Candidatus Bathyarchaeota archaeon
A0A7S3GPX1 Spumella elongata
TRINITY DN12381 c1 g4 i1.p1 Pseudotrichonympha grassii
G8YBW7 Pichia sorbitophila
Q6BZD6 Debaryomyces hansenii
A0A0V1PV62 Debaryomyces fabryi
A0A6C1E7V3 Saccharomyces pastorianus
A0A2P7YV86 Candida pseudohaemulonii
C5M895 Candida tropicalis
Q59X89 Candida albicans
B9WMJ7 Candida dubliniensis
V4C1F0 Lottia gigantea
A0A8B7YSX1 Acanthaster planci
A0A3S0ZT37 Elysia chlorotica
A0A7R9XIL7 Argulus foliaceus
L8H896 Acanthamoeba castellanii str. Neff
A0A6G3MIX9 Henneguya salminicola
A0A6V7T708 Plasmodium vinckei petteri
A0A8C9H9P9 Piliocolobus tephrosceles
A0A1J1H3I3 Plasmodium gallinaceum
A5K9N1 Plasmodium vivax
W7K002 Plasmodium falciparum


R

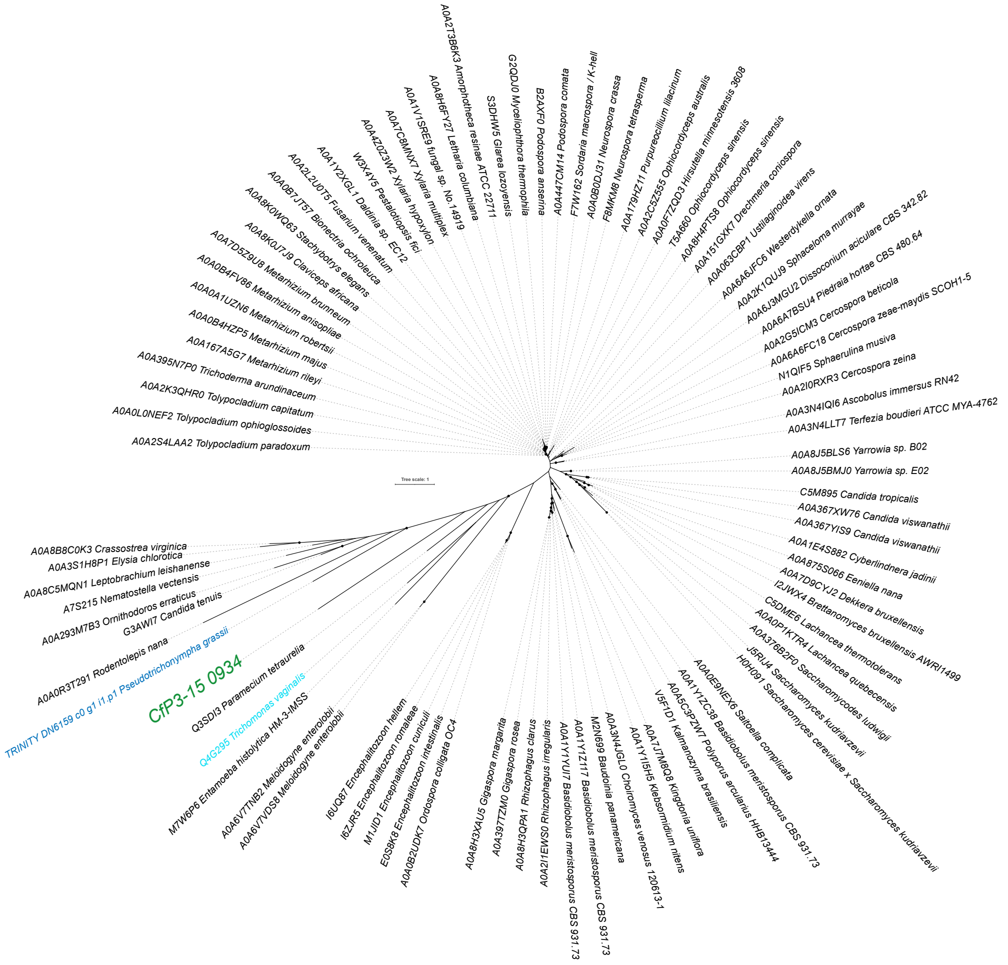

Tree scale: 1
A0A2T3B8K3 Amorphotheca resinae ATCC 22711
A0A8H6FY27 Letharia columbiana
A0A1V1SRE9 fungal sp. No. 11419
A0A7C8MNX7 Xylaria multiplex
A0A4Z0Z3W2 Xylaria hypoxylon
W3X4Y5 Pestalotiopsis fici
A0A1Y2XGL1 Daldinia sp. EC12
A0A2L2U0T5 Fusarium venenatum
A0A0B7JT57 Bionectria ochroleuca
A0A8K0WQ63 Stachybotrys elegans
A0A8K0U7J9 Claviceps africana
A0A7D5Z9U8 Metarhizium brunneum
A0A0B4FV86 Metarhizium anisopliae
A0A0A1UZN6 Metarhizium robertsii
A0A0B4HZP5 Metarhizium majus
A0A167A5G7 Metarhizium rileyi
A0A395N7P0 Trichoderma arundinaceum
A0A2K3QHR0 Tolypocladium capitatum
A0A0L0NEF2 Tolypocladium ophioglossoides
A0A2S4LAA2 Tolypocladium paradoxum
S3DHW5 Glarea lozoyensis
G2QDJ0 Myceliophthora thermophila
B2AXF0 Podospora anserina
A0A447CM14 Podospora comata
F7W162 Sordaria macrospora k-hell
A0A0B0DJ31 Neurospora crassa
F8MKM8 Neurospora tetrasperma
A0A179HZ11 Purpureocillium lilacinum
A0A2C5Z555 Ophiocordyceps australis
A0A0F7ZQ03 Hirsutella minnesotensis 3608
T5A660 Ophiocordyceps sinensis
A0A8H4PTS8 Ophiocordyceps sinensis
A0A151GXK7 Drechmeria coniospora
A0A063C8P1 Ustilaginoidea virens
A0A6A6JFC6 Westerdykella ornata
A0A2K1QUJ9 Sphaceloma murrayae
A0A6J3MGU2 Dissoconium aciculare CBS 342.82
A0A6A7BSU4 Piedraia hortae CBS 480.64
A0A2G5ICM3 Cercospora beticola
A0A6A6FC18 Cercospora zeae-maydis SCOH1-5
N1QIF5 Sphaerulina musiva
A0A2I0RXR3 Cercospora zeina
A0A3N4IQI6 Ascobolus immersus RN42
A0A3N4LLT7 Terfezia boudieri ATCC MYA-4762
A0A8J5BLS6 Yarrowia sp. B02
A0A8J5BMJ0 Yarrowia sp. E02
C5M895 Candida tropicalis
A0A367XW76 Candida viswanathii
A0A367YIS9 Candida viswanathii
A0A1E4S882 Cyberlindnera jadinii
A0A875S066 Eeniella nana
A0A7D9CYJ2 Dekkera bruxellensis
I2JWX4 Brettanomyces bruxellensis AWRI1499
C5DME6 Lachancea thermotolerans
A0A0P1KTR4 Lachancea quebecensis
A0A376B2F0 Saccharomycodes ludwigii
J5RIJ4 Saccharomyces kudriavzevii
H0H091 Saccharomyces cerevisiae x Saccharomyces kudriavzevii
A0A0E9NEX6 Saitoella complicata
A0A1Y1ZC38 Basidiobolus meristosporus CBS 931.73
A0A5C3P2W7 Polyporus arcularius HHB13444
V5F1D1 Kalmanozyma brasiliensis
A0A1J7M8Q8 Kingdonia uniflora
A0A1Y1I5H5 Klebsormidium nitens
A0A3N4JGL0 Choiromyces venosus 120613-1
M2N599 Baudoinia panamericana
A0A1Y1Z1T7 Basidiobolus meristosporus CBS 931.73
A0A1Y1YU7 Basidiobolus meristosporus CBS 931.73
A0A2I1EWS0 Rhizophagus irregularis
A0A8H3QPA1 Rhizophagus clarus
A0A397TZM0 Gigaspora rosea
A0A8H3XAU5 Gigaspora margarita
A0A0B2UQK7 Ordospora colligata OC4
E0S8K8 Encephalitozoon intestinalis
M1JID1 Encephalitozoon cuniculi
I6ZJR5 Encephalitozoon romaleae
I6UQ87 Encephalitozoon hellem
A0A6V7VDS8 Meloidogyne enterolobii
A0A6V7TNB2 Meloidogyne enterolobii
M7W6P6 Entamoeba histolytica HM-3:IMSS
Q4G295 Trichomonas vaginalis
Q3SDI3 Paramecium tetraurelia
CfP3-15 0934
TRINITY DN6159 c0 g1 i1.p1 Pseudotrichonympha grassii
A0A0R3T291 Rodentolepis nana
G3AWI7 Candida tenuis
A0A293M7B3 Ornithodoros erraticus
A7S215 Nematostella vectensis
A0A8C5MQN1 Leptobrachium leishanense
A0A3S1H8P1 Elysia chlorotica
A0A8B8C0K3 Crassostrea virginica

S

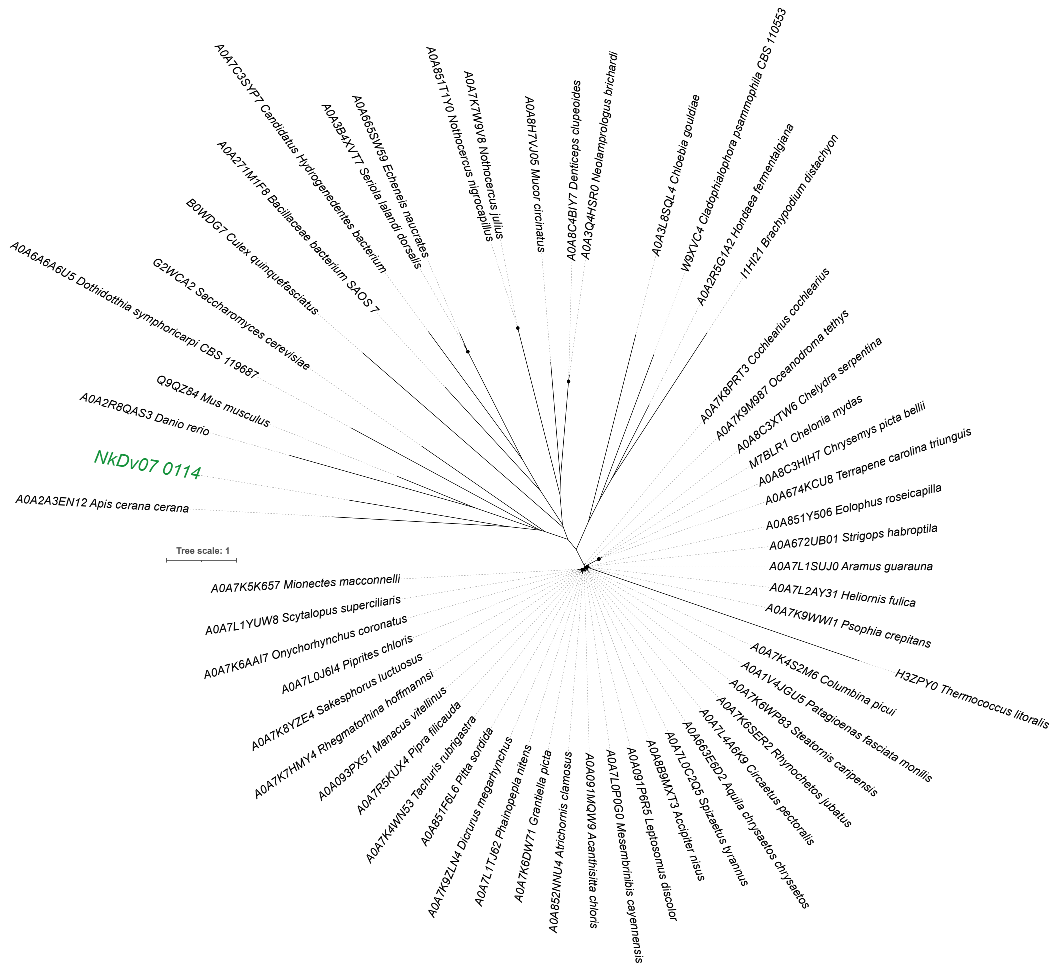
A0A7C3SYP7 Candidatus Hydrogenedentes bacterium
A0A665SW59 Echeneis naucrates
A0A3B4XVT7 Seriola lalandi dorsalis
A0A851T1Y0 Nothocercus nigrocapillus
A0A7K7W9V8 Nothocercus julius
A0A8H7VJ05 Mucor circinatus
A0A8C4BIY7 Denticeps clupeoides
A0A3Q4HSR0 Neolamprologus brichardi
A0A3L8SQL4 Chloebia gouldiae
W9XVC4 Cladophialophora psammophila CBS 110553
A0A2R5G1A2 Hondaea fermentalgiana
I1HI21 Brachypodium distachyon
A0A271M1F8 Bacillaceae bacterium SAOS 7
B0WDG7 Culex quinquefasciatus
G2WCA2 Saccharomyces cerevisiae
A0A6A6A6U5 Dothidotthia symphoricarpi CBS 119687
Q9QZ84 Mus musculus
A0A2R8QAS3 Danio rerio
NkDv07 0114
A0A2A3EN12 Apis cerana cerana
Tree scale: 1
A0A7K8PRT3 Cochlearius cochlearius
A0A7K9M987 Oceanodroma tethys
A0A8C3XTW6 Chelydra serpentina
M7BLR1 Chelonia mydas
A0A8C3HIH7 Chrysemys picta bellii
A0A674KCU8 Terrapene carolina triunguis
A0A851Y506 Eolophus roseicapilla
A0A672UB01 Strigops habroptila
A0A7L1SUJ0 Aramus guarauna
A0A7L2AY31 Heliornis fulica
A0A7K9WWI1 Psophia crepitans
H3ZPY0 Thermococcus litoralis
A0A7K5K657 Mionectes macconnelli
A0A7L1YUW8 Scytalopus superciliaris
A0A7K6AAI7 Onychorhynchus coronatus
A0A7L0J6I4 Piprites chloris
A0A7K6YZE4 Sakesphorus luctuosus
A0A7K7HMY4 Rhegmatorhina hoffmannsi
A0A093PX51 Manacus vitellinus
A0A7R5KUX4 Pipra filicauda
A0A7K4WN53 Tachuris rubrigastra
A0A851F6L6 Pitta sordida
A0A7K9ZLN4 Dicrurus megarhynchus
A0A7L1TJ62 Phainopepla nitens
A0A7K6DW71 Grantiella picta
A0A852NNU4 Atrichornis clamosus
A0A091MQW9 Acanthisitta chloris
A0A7L0P0G0 Mesembrinibis cayennensis
A0A091P6R5 Leptosomus discolor
A0A8B9MXT3 Accipiter nisus
A0A7L0C2Q5 Spizaetus tyrannus
A0A663E6D2 Aquila chrysaetos chrysaetos
A0A7L4A6K9 Circaetus pectoralis
A0A7K6SER2 Rhynochetos jubatus
A0A7K6WP83 Steatornis caripensis
A0A1V4JGU5 Patagioenas fasciata monilis
A0A7K4S2M6 Columbina picui

T

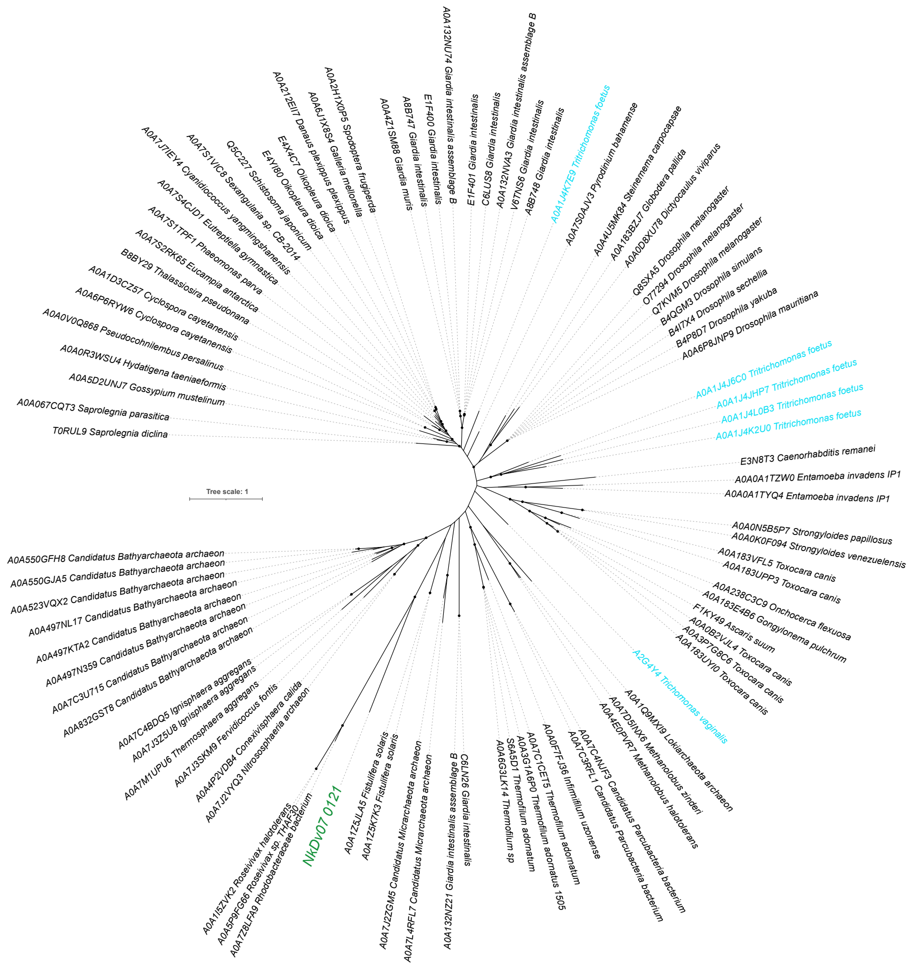
A0A132NU74 Giardia intestinalis assemblage B
A0A2H1X0P5 Spodoptera frugiperda
A0A6J1X8S4 Galleria mellonella
A0A212EII7 Danaus plexippus plexippus
E4X4C7 Oikopleura dioica
E4YI80 Oikopleura dioica
Q5C227 Schistosoma japonicum
A0A7S1VIC8 Sexangularia sp. CB-2014
A0A7J7IEY4 Cyanidiococcus yangmingshanensis
A0A7S4CJD1 Eutreptiella gymnastica
A0A7S1TPF1 Phaeomonas parva
A0A7S2RK65 Eucampia antarctica
B8BY29 Thalassiosira pseudonana
A0A1D3CZ57 Cyclospora cayetanensis
A0A6P6RYW6 Cyclospora cayetanensis
A0A0V0Q868 Pseudocohnilembus persalinus
A0A0R3WSU4 Hydatigena taeniaeformis
A0A5D2UNJ7 Gossypium mustelinum
A0A067CQT3 Saprolegnia parasitica
T0RUL9 Saprolegnia diclina
A0A4Z1SM88 Giardia muris
A8B747 Giardia intestinalis
E1F400 Giardia intestinalis
E1F401 Giardia intestinalis
C6LUS8 Giardia intestinalis
A0A132NVA3 Giardia intestinalis assemblage B
V6TNS6 Giardia intestinalis
A8B748 Giardia intestinalis
A0A1J4K7E9 Tritrichomonas foetus
A0A7S0AJV3 Pyrodinium bahamense
A0A4U5MK84 Steinernema carpocapsae
A0A183BZJ7 Globodera pallida
A0A0D8XU78 Dictyocaulus viviparus
Q8SXA5 Drosophila melanogaster
O77294 Drosophila melanogaster
Q7KVM5 Drosophila melanogaster
B4QGM3 Drosophila simulans
B4I7X4 Drosophila sechellia
B4P8D7 Drosophila yakuba
A0A6P8JNP9 Drosophila mauritiana
A0A1J4J6C0 Tritrichomonas foetus
A0A1J4JHP7 Tritrichomonas foetus
A0A1J4L0B3 Tritrichomonas foetus
A0A1J4K2U0 Tritrichomonas foetus
E3N8T3 Caenorhabditis remanei
A0A0A1TZW0 Entamoeba invadens IP1
A0A0A1TYQ4 Entamoeba invadens IP1
Tree scale: 1
A0A0N5B5P7 Strongyloides papillosus
A0A0K0F094 Strongyloides venezuelensis
A0A183VFL5 Toxocara canis
A0A183UPP3 Toxocara canis
A0A238C3C9 Onchocerca flexuosa
A0A183E4B6 Gongylonema pulchrum
F1KY49 Ascaris suum
A0A0B2VJL4 Toxocara canis
A0A3P7G8C6 Toxocara canis
A0A183UYI0 Toxocara canis
A2G4Y4 Trichomonas vaginalis
A0A550GFH8 Candidatus Bathyarchaeota archaeon
A0A550GJA5 Candidatus Bathyarchaeota archaeon
A0A523VQX2 Candidatus Bathyarchaeota archaeon
A0A497NL17 Candidatus Bathyarchaeota archaeon
A0A497KTA2 Candidatus Bathyarchaeota archaeon
A0A497N359 Candidatus Bathyarchaeota archaeon
A0A7C3U715 Candidatus Bathyarchaeota archaeon
A0A832GST8 Candidatus Bathyarchaeota archaeon
A0A7C4BDQ5 Ignisphaera aggregans
A0A7J3Z5U8 Ignisphaera aggregans
A0A7M1UPU6 Thermosphaera aggregans
A0A7J3SKM9 Fervidicoccus fontis
A0A4P2VDB4 Conexivisphaera calida
A0A7J2VYQ3 Nitrososphaeria archaeon
A0A1I5ZVK2 Roseivivax halotolerans
A0A5P9FG66 Roseivivax sp. THAF30
A0A7Z8LFA9 Rhodobacteraceae bacterium
NkDv07 0121
A0A1Z5JLA5 Fistulifera solaris
A0A1Z5K7K3 Fistulifera solaris
A0A7J2ZGM5 Candidatus Micrarchaeota archaeon
A0A7L4RFL7 Candidatus Micrarchaeota archaeon
A0A132NZ21 Giardia intestinalis assemblage B
C6LN26 Giardia intestinalis
A0A6G3LK14 Thermofilum sp
S6A5D1 Thermofilum adornatum
A0A3G1A6P0 Thermofilum adornatus 1505
A0A7C1CET5 Thermofilum adornatum
A0A0F7FJ36 Infirmifilum uzonense
A0A7C3RFL1 Candidatus Parcubacteria bacterium
A0A7C4NJF3 Candidatus Parcubacteria bacterium
A0A4E0PVR7 Methanolobus halotolerans
A0A7D5INX6 Methanolobus zinderi
A0A1Q9MXI9 Lokiarchaeota archaeon

U

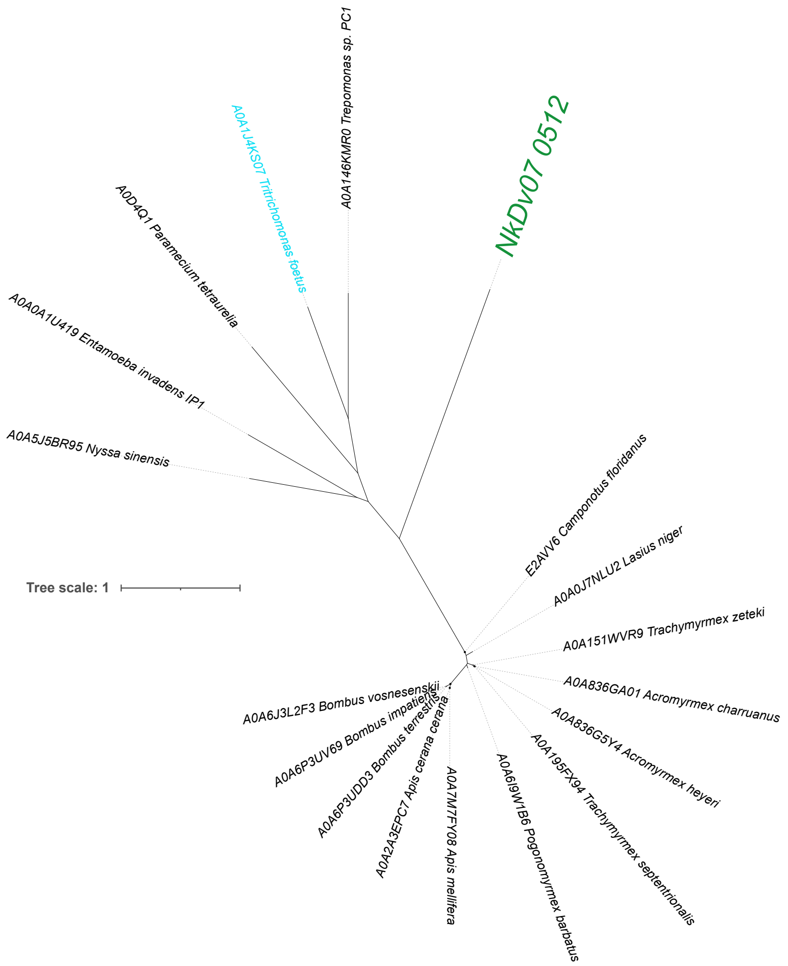

A0A1J4KS07 Tritrichomonas foetus
A0A146KMR0 Trepomonas sp. PC1
NkDv07 0512
A0D4Q1 Paramecium tetraurelia
A0A0A1U419 Entamoeba invadens IP1
A0A5J5BR95 Nyssa sinensis
Tree scale: 1
E2AVV6 Camponotus floridanus
A0A0J7NLU2 Lasius niger
A0A151WVR9 Trachymyrmex zeteki
A0A836GA01 Acromyrmex charruanus
A0A836G5Y4 Acromyrmex heyeri
A0A195FX94 Trachymyrmex septentrionalis
A0A6J9W1B6 Pogonomyrmex barbatus
A0A7M7FY08 Apis mellifera
A0A2A3EPC7 Apis cerana cerana
A0A6P3UDD3 Bombus terrestris
A0A6P3UV69 Bombus impatiens
A0A6J3L2F3 Bombus vosnesenskii


V

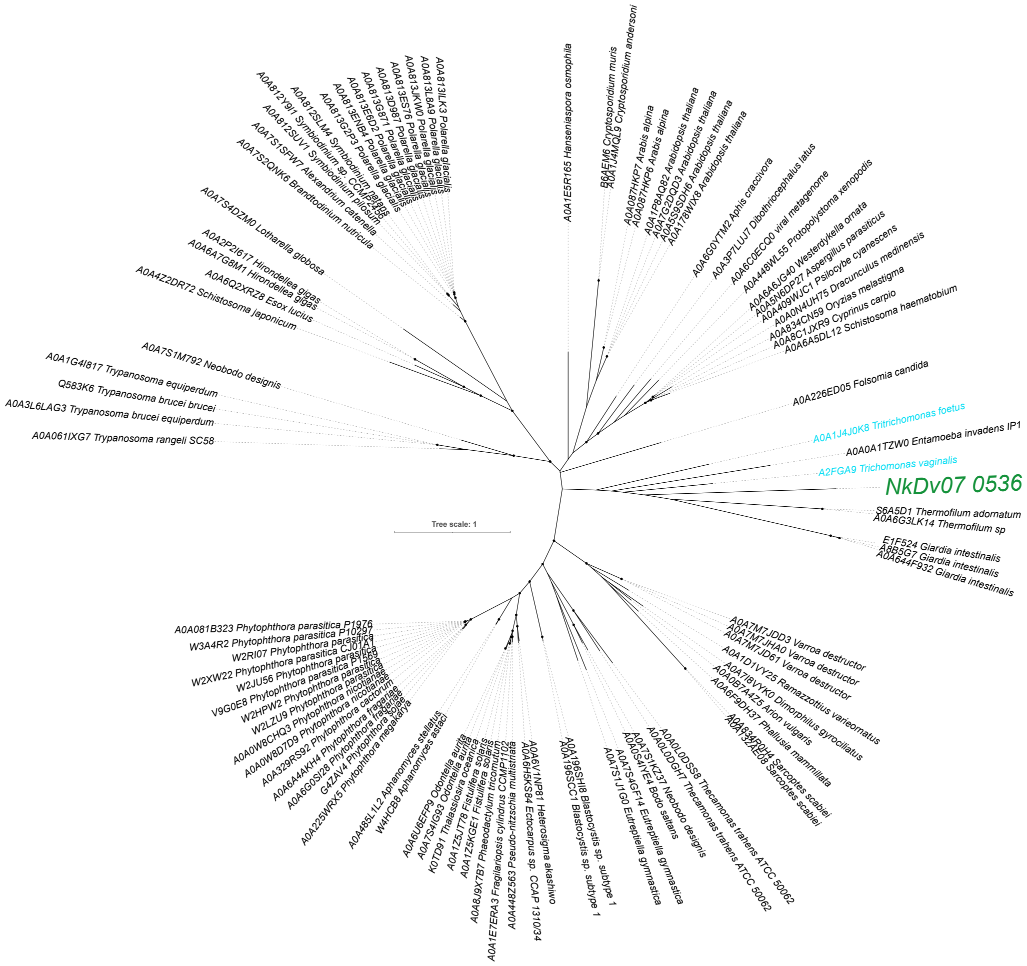

A0A1E5R165 Hanseniaspora osmophila
A0A7S4DZM0 Lotharella globosa
A0A2P2I617 Hirondellea gigas
A0A6A7G8M1 Hirondellea gigas
A0A6Q2XRZ8 Esox lucius
A0A4Z2DR72 Schistosoma japonicum
A0A7S1M792 Neobodo designis
A0A1G4I817 Trypanosoma equiperdum
Q583K6 Trypanosoma brucei brucei
A0A3L6LAG3 Trypanosoma brucei equiperdum
A0A061IXG7 Trypanosoma rangeli SC58
A0A226ED05 Folsomia candida
A0A1J4J0K8 Tritrichomonas foetus
A0A0A1TZW0 Entamoeba invadens IP1
A2FGA9 Trichomonas vaginalis
NkDv07 0536
S6A5D1 Thermofilum adornatum
A0A6G3LK14 Thermofilum sp
E1F524 Giardia intestinalis
A8B5G7 Giardia intestinalis
A0A644F932 Giardia intestinalis
Tree scale: 1
A0A081B323 Phytophthora parasitica P1976
W3A4R2 Phytophthora parasitica P10297
A0A7M7JDD3 Varroa destructor
A0A1D1VY25 Ramazzottius varieornatus

W

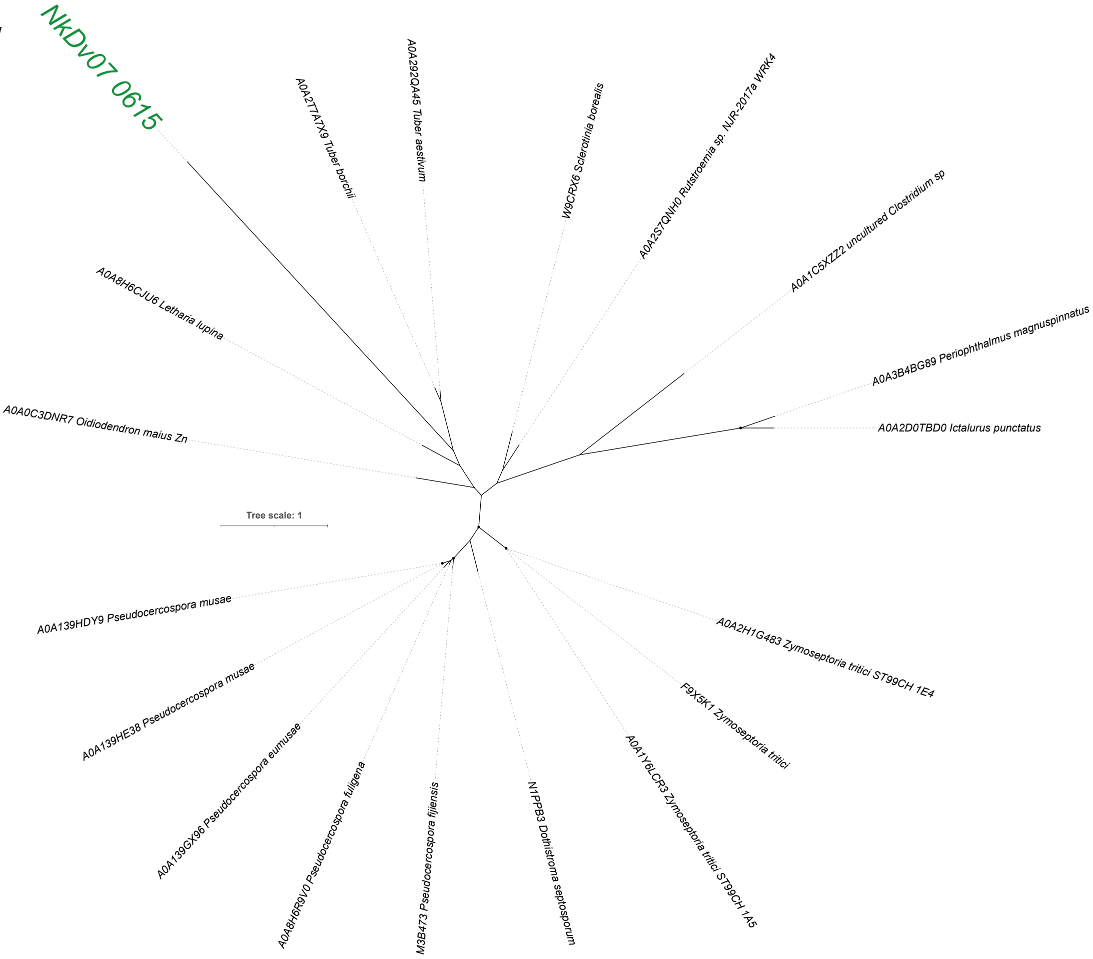
NkDv07 0615
A0A2T7A7X9 Tuber borchii
A0A292Q445 Tuber aestivum
W9CRX6 Sclerotinia borealis
A0A2S7QNH0 Rutstroemia sp. NJR-2017a WRK4
A0A1C5XZZ2 uncultured Clostridium sp
A0A3B4BG89 Periophthalmus magnuspinnatus
A0A2D0TBD0 Ictalurus punctatus
A0A8H6CJU6 Letharia lupina
A0A0C3DNR7 Oidiodendron maius Zn
Tree scale: 1
A0A139HDY9 Pseudocercospora musae
A0A139HE38 Pseudocercospora musae
A0A139GX96 Pseudocercospora eumusae
A0A8H6R9V0 Pseudocercospora fuligena
M3B473 Pseudocercospora fijiensis
N1PPB3 Dothistroma septosporum
A0A1Y6LCR3 Zymoseptoria tritici ST99CH 1A5
F9X5K1 Zymoseptoria tritici
A0A2H1G483 Zymoseptoria tritici ST99CH 1E4

X

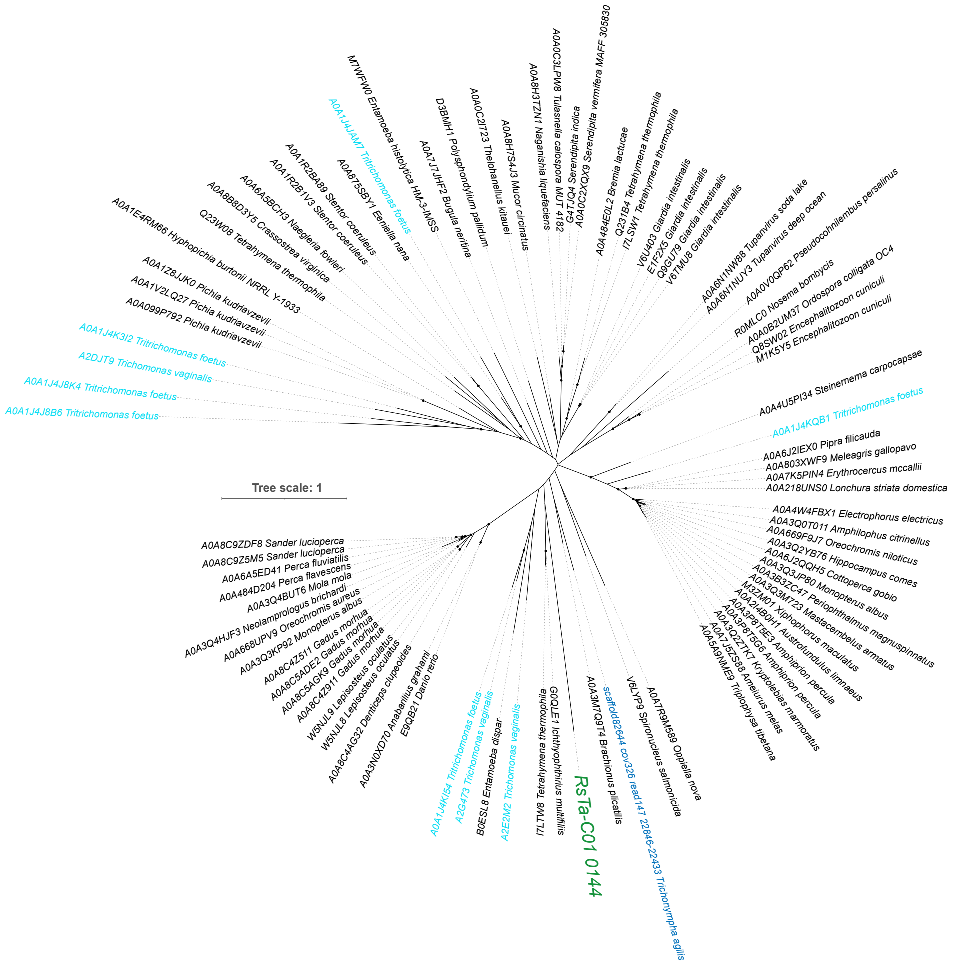
M7WFW0 Entamoeba histolytica HM-3-IMSS
A0A1J4JAM7 Tritrichomonas foetus
A0A875SBY1 Eeniella nana
A0A7J7JHF2 Bugula neritina
D3BMH1 Polysphondylium pallidum
A0A0C2I7Z3 Thelohanellus kitauei
A0A8H7S4J3 Mucor circinatus
A0A8H3TZN1 Naganishia liquefaciens
A0A0C3LPW8 Tulasnella calospora MUT 4182
A0A0C2XQX9 Serendipita vermifera MAFF 305830
A0A1R2BA89 Stentor coeruleus
A0A1R2B1V3 Stentor coeruleus
A0A6A5BCH3 Naegleria fowleri
A0A8B8D3Y5 Crassostrea virginica
Q23W08 Tetrahymena thermophila
A0A1E4RM66 Hyphopichia burtonii NRRL Y-1933
A0A1Z8JJK0 Pichia kudriavzevii
A0A1V2LQ27 Pichia kudriavzevii
A0A099P792 Pichia kudriavzevii
A0A1J4K3I2 Tritrichomonas foetus
A2DJT9 Trichomonas vaginalis
A0A1J4J8K4 Tritrichomonas foetus
A0A1J4J8B6 Tritrichomonas foetus
A0A484E0L2 Bremia lactucae
Q231B4 Tetrahymena thermophila
I7LSW1 Tetrahymena thermophila
V6U403 Giardia intestinalis
E1F2X5 Giardia intestinalis
Q9GU79 Giardia intestinalis
V6TMU8 Giardia intestinalis
A0A6N1NW88 Tupanvirus soda lake
A0A6N1NUY3 Tupanvirus deep ocean
A0A0V0QP62 Pseudocohnilembus persalinus
R0MLC0 Nosema bombycis
A0A0B2UM37 Ordospora colligata OC4
Q8SW02 Encephalitozoon cuniculi
M1K5Y5 Encephalitozoon cuniculi
A0A4U5PI34 Steinernema carpocapsae
A0A1J4KQB1 Tritrichomonas foetus
A0A6J2IEX0 Pipra filicauda
A0A803XWF9 Meleagris gallopavo
A0A7K5PIN4 Erythrocercus mccallii
A0A218UNS0 Lonchura striata domestica
A0A4W4FBX1 Electrophorus electricus
A0A3Q0T011 Amphilophus citrinellus
A0A669F9J7 Oreochromis niloticus
A0A3Q2YB76 Hippocampus comes
A0A6J2QQH5 Cottoperca gobio
A0A3Q3JP80 Monopterus albus
A0A3B3ZC47 Periophthalmus magnuspinnatus
A0A3Q3M723 Mastacembelus armatus
M3ZM01 Xiphophorus maculatus
A0A3Q4B0H1 Austrofundulus limnaeus
A0A3P8TSE3 Amphiprion percula
A0A3P8T9G6 Amphiprion percula
A0A3Q2ZTK7 Kryptolebias marmoratus
A0A7J5Z588 Ameiurus melas
A0A5A9NME9 Triplophysa tibetana
Tree scale: 1
A0A8C9ZDF8 Sander lucioperca
A0A8C9Z5M5 Sander lucioperca
A0A6A5ED41 Perca fluviatilis
A0A484D204 Perca flavescens
A0A3Q4BUT6 Mola mola
A0A3Q4HJF3 Neolamprologus brichardi
A0A668UPV9 Oreochromis aureus
A0A3Q3KP92 Monopterus albus
A0A8C4Z511 Gadus morhua
A0A8C5ADE2 Gadus morhua
A0A8C5AGK9 Gadus morhua
A0A8C4Z911 Gadus morhua
W5NJL9 Lepisosteus oculatus
W5NJL8 Lepisosteus oculatus
A0A8C4AG32 Denticeps clupeoides
A0A3N0XD70 Anabarilius grahami
E9QB21 Danio rerio
A0A1J4KI54 Tritrichomonas foetus
A2G473 Trichomonas vaginalis
B0ESL8 Entamoeba dispar
A2E2M2 Trichomonas vaginalis
I7LTW8 Tetrahymena thermophila
G0QLE1 Ichthyophthirius multifiliis
RsTa-C01 0144
A0A3M7Q9T4 Brachionus plicatilis
scaffold82644 cov326 read147 22846-22433 Trichonympha agilis
V6LYP9 Spironucleus salmonicida
A0A7R9M589 Oppiella nova

Y

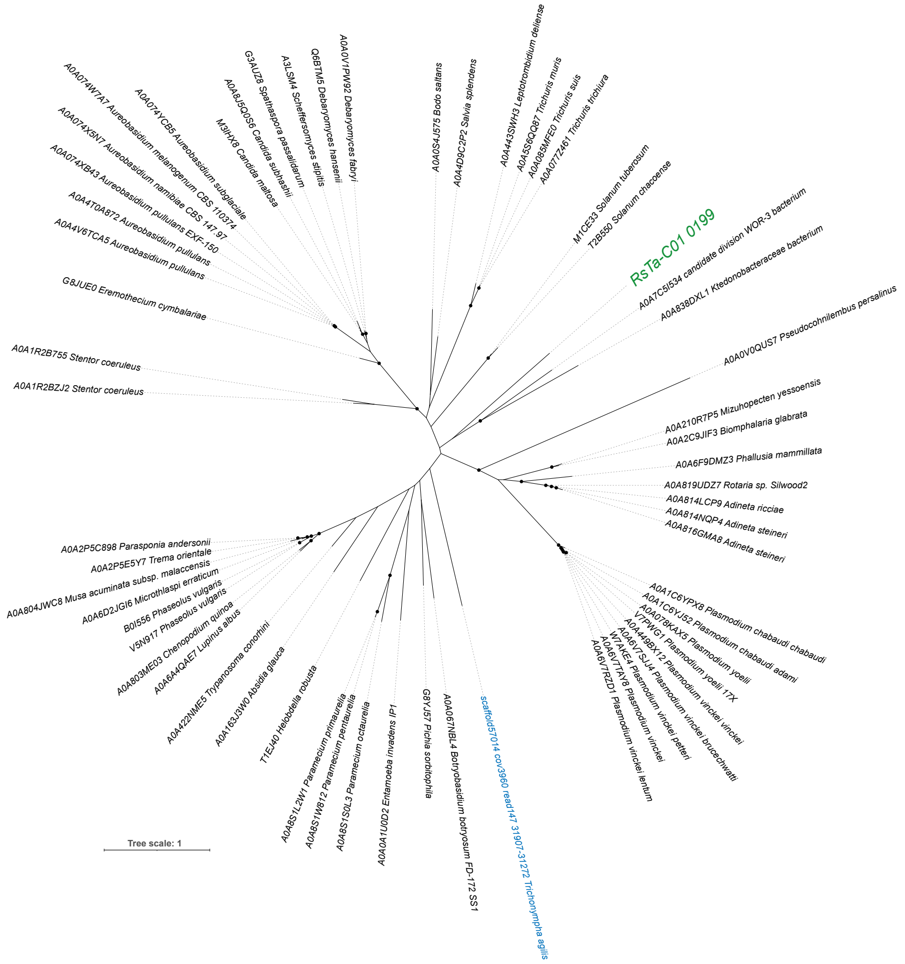
A0A074W7A7 Aureobasidium melanogenum CBS 110374
A0A074YCB5 Aureobasidium subglaciale
A0A074X5N7 Aureobasidium namibiae CBS 147.97
A0A074XB43 Aureobasidium pullulans EXF-150
A0A4T0A872 Aureobasidium pullulans
A0A4V6TCA5 Aureobasidium pullulans
M3IHX8 Candida maltosa
A0A8J5Q0S6 Candida subhashii
G3AUZ8 Spathaspora passalidarum
A3LSM4 Scheffersomyces stipitis
Q6BTM5 Debaryomyces hansenii
A0A0V1PW92 Debaryomyces fabryi
A0A0S4J575 Bodo saltans
A0A4D9C2P2 Salvia splendens
A0A443SWH3 Leptotrombidium deliense
A0A5S6QQ87 Trichuris muris
A0A085MFE0 Trichuris suis
A0A077Z461 Trichuris trichiura
M1CE33 Solanum tuberosum
T2B550 Solanum chacoense
RsTa-C01 0199
A0A7C5I534 candidate division WOR-3 bacterium
A0A838DXL1 Ktedonobacteraceae bacterium
A0A0V0QUS7 Pseudocohnilembus persalinus
G8JUE0 Eremothecium cymbalariae
A0A1R2B755 Stentor coeruleus
A0A1R2BZJ2 Stentor coeruleus
A0A210R7P5 Mizuhopecten yessoensis
A0A2C9JIF3 Biomphalaria glabrata
A0A6F9DMZ3 Phallusia mammillata
A0A819UDZ7 Rotaria sp. Silwood2
A0A814LCP9 Adineta ricciae
A0A814NQP4 Adineta steineri
A0A816GMA8 Adineta steineri
A0A2P5C898 Parasponia andersonii
A0A2P5E5Y7 Trema orientale
A0A804JWC8 Musa acuminata subsp. malaccensis
A0A6D2JGI6 Microthlaspi erraticum
B0I556 Phaseolus vulgaris
V5N917 Phaseolus vulgaris
A0A803ME03 Chenopodium quinoa
A0A6A4QAE7 Lupinus albus
A0A422NME5 Trypanosoma conorhini
A0A163J3W0 Absidia glauca
T1EJ40 Helobdella robusta
A0A8S1L2W1 Paramecium primaurelia
A0A8S1W812 Paramecium pentaurelia
A0A8S1S0L3 Paramecium octaurelia
A0A0A1U0D2 Entamoeba invadens IP1
G8YJ57 Pichia sorbitophila
A0A067NBL4 Botryobasidium botryosum FD-172 SS1
scaffold57014 cov3960 read147 31907-31272 Trichonympha agilis
A0A1C6YPX8 Plasmodium chabaudi chabaudi
A0A1C6YJ52 Plasmodium chabaudi adami
A0A078KAX5 Plasmodium yoelii
V7PWG1 Plasmodium yoelii 17X
A0A449BX12 Plasmodium vinckei vinckei
A0A6V7SJJ4 Plasmodium vinckei brucechwatti
W7AKE4 Plasmodium vinckei petteri
A0A6V7TAY8 Plasmodium vinckei
A0A6V7RZD1 Plasmodium vinckei lentum
Tree scale: 1

Z

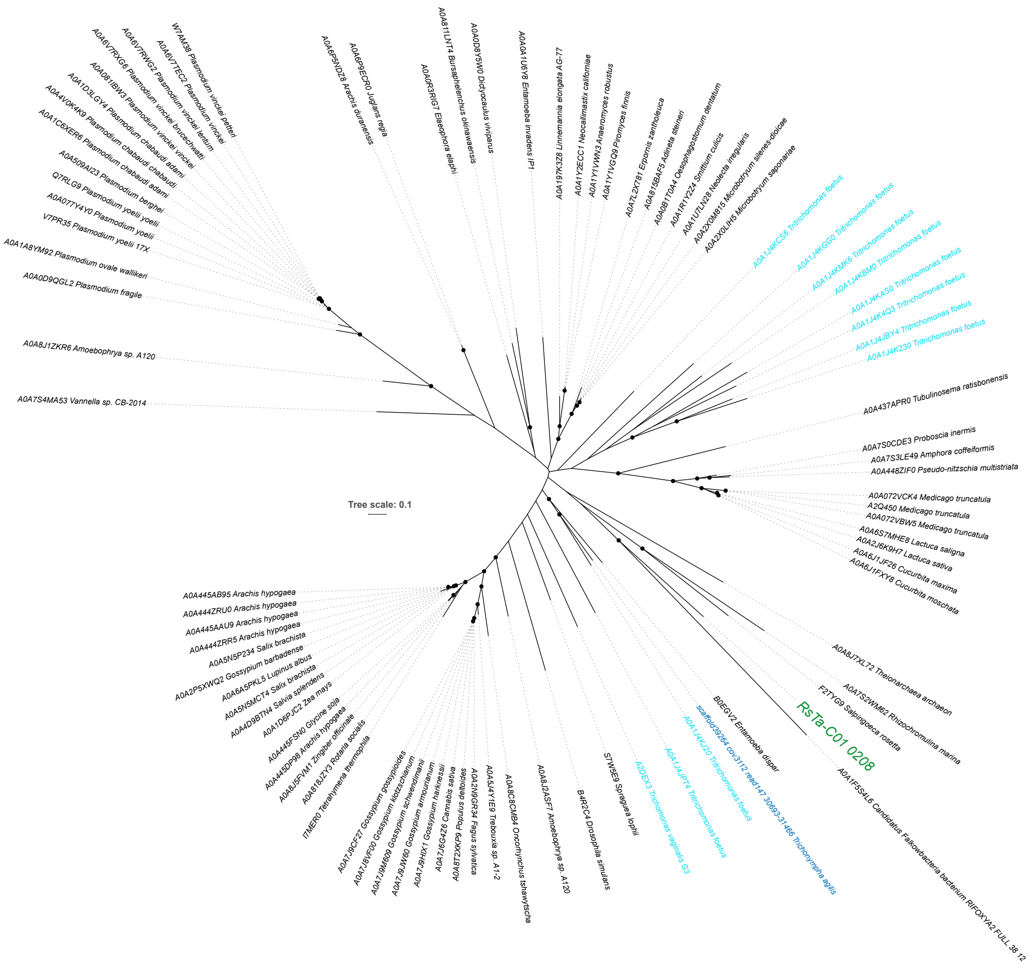
W7AM38 Plasmodium vinckei petteri
A0A6V7TEC2 Plasmodium vinckei
A0A6V7RWG2 Plasmodium vinckei lentum
A0A6V7RXG6 Plasmodium vinckei brucechwatti
A0A081IBW3 Plasmodium vinckei vinckei
A0A1D3LGY4 Plasmodium chabaudi adami
A0A4V0K4K9 Plasmodium chabaudi chabaudi
A0A1C6XER6 Plasmodium chabaudi adami
A0A509AI23 Plasmodium berghei
Q7RLG9 Plasmodium yoelii yoelii
A0A077Y4Y0 Plasmodium yoelii
V7PR35 Plasmodium yoelii 17X
A0A1A8YM92 Plasmodium ovale wallikeri
A0A0D9QGL2 Plasmodium fragile
A0A8J1ZKR6 Amoebophrya sp. A120
A0A7S4MA53 Vannella sp. CB-2014
A0A6P5NDZ8 Arachis duranensis
A0A6P9ECR0 Juglans regia
A0A0R3RIG7 Elaeophora elaphi
A0A811LNT4 Bursaphelenchus okinawaensis
A0A0D8Y5W0 Dictyocaulus viviparus
A0A0A1U6Y8 Entamoeba invadens IP1
A0A197K3Z8 Linnemannia elongata AG-77
A0A1Y2ECC1 Neocallimastix californiae
A0A1Y1VWN3 Anaeromyces robustus
A0A1Y1VGQ9 Piromyces finnis
A0A7L2X781 Erpornis zantholeuca
A0A815BAF5 Adineta steineri
A0A0B1T0A4 Oesophagostomum dentatum
A0A1R1Y2Z4 Smittium culicis
A0A1U7LN28 Neolecta irregularis
A0A2X0M815 Microbotryum silenes-dioicae
A0A2X0LIH5 Microbotryum saponariae
A0A1J4KCS6 Tritrichomonas foetus
A0A1J4KGG0 Tritrichomonas foetus
A0A1J4KMK6 Tritrichomonas foetus
A0A1J4KBM0 Tritrichomonas foetus
A0A1J4KAS0 Tritrichomonas foetus
A0A1J4K4Q3 Tritrichomonas foetus
A0A1J4JBY4 Tritrichomonas foetus
A0A1J4K230 Tritrichomonas foetus
A0A437APR0 Tubulinosema ratisbonensis
A0A7S0CDE3 Proboscia inermis
A0A7S3LE49 Amphora coffeiformis
A0A448ZIF0 Pseudo-nitzschia multistriata
A0A072VCK4 Medicago truncatula
A2Q450 Medicago truncatula
A0A072VBW5 Medicago truncatula
A0A6S7MHE8 Lactuca saligna
A0A2J6K9H7 Lactuca sativa
A0A6J1JF26 Cucurbita maxima
A0A6J1FXY8 Cucurbita moschata
Tree scale: 0.1
A0A445AB95 Arachis hypogaea
A0A444ZRU0 Arachis hypogaea
A0A445AAU9 Arachis hypogaea
A0A444ZRR5 Arachis hypogaea
A0A5N5P234 Salix brachista
A0A2P5XWQ2 Gossypium barbadense
A0A6A5PKL5 Lupinus albus
A0A5N5MCT4 Salix brachista
A0A4D9BTN4 Salvia splendens
A0A1D6PJC2 Zea mays
A0A445FSN0 Glycine soja
A0A445DP98 Arachis hypogaea
A0A8J5FVM1 Zingiber officinale
A0A818JZY3 Rotaria socialis
I7MER0 Tetrahymena thermophila
A0A7J9CF27 Gossypium gossypioides
A0A7J8VF00 Gossypium klotzschianum
A0A7J9M609 Gossypium schwendimanii
A0A7J9JW60 Gossypium armourianum
A0A7J9HIX1 Gossypium harknessii
A0A7J6G4Z6 Cannabis sativa
A0A8T2XKP9 Populus deltoides
A0A2N9GR34 Fagus sylvatica
A0A5J4Y1E9 Trebouxia sp. A1-2
A0A8C8CMB4 Oncorhynchus tshawytscha
A0A8J2ASF7 Amoebophrya sp. A120
B4R2C4 Drosophila simulans
S7W5E9 Spraguea lophii
A2DEX3 Trichomonas vaginalis G3
A0A1J4JPT4 Tritrichomonas foetus
A0A1J4KJ20 Tritrichomonas foetus
scaffold39264 cov3112 read147 30693-31466 Trichonympha agilis
B0EGV2 Entamoeba dispar
RsTa-C01 0208
F2TYG9 Salpingoeca rosetta
A0A7S2WM62 Rhizochromulina marina
A0A8J7XL72 Theionarchaea archaeon
A0A1F5S4L6 Candidatus Falkowbacteria bacterium RIFOXYA2 FULL 38 12

AA

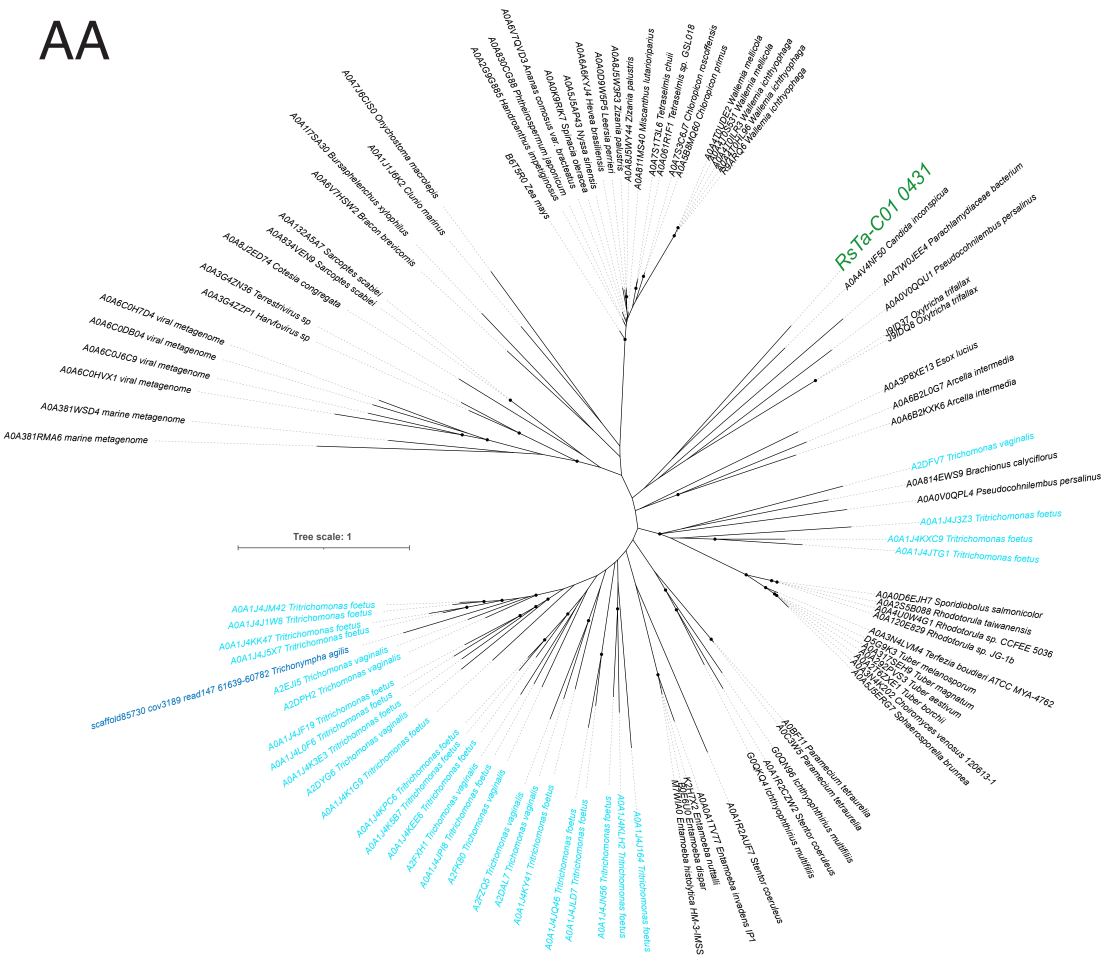


AB

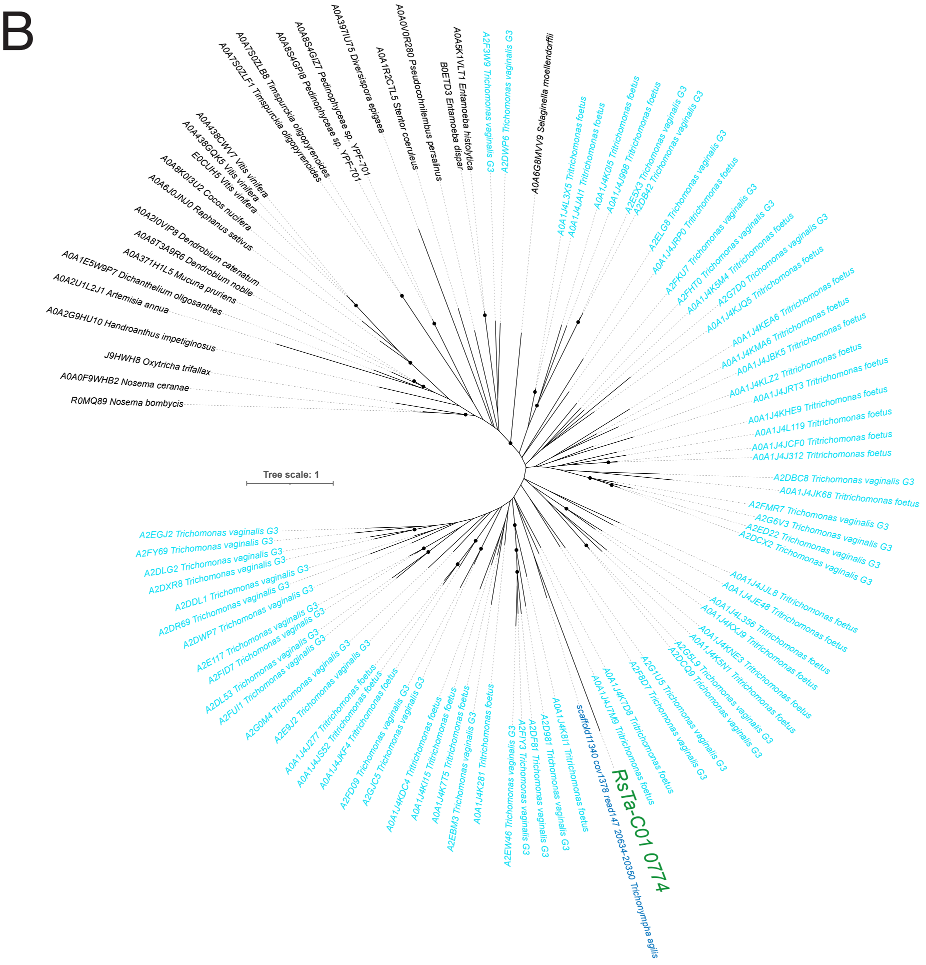

AC

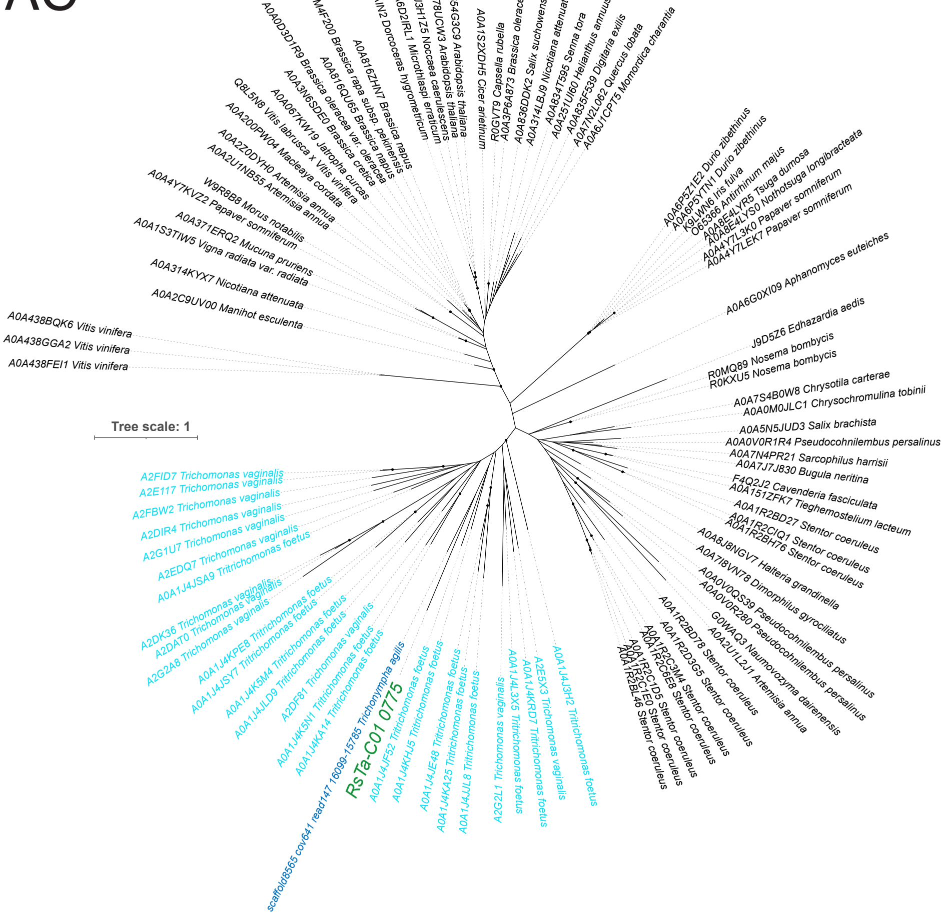
Tree scale: 1
RsTa-C01 0775
scaffold8565 cov641 read147 16099-15785 Trichonympha agilis

AD

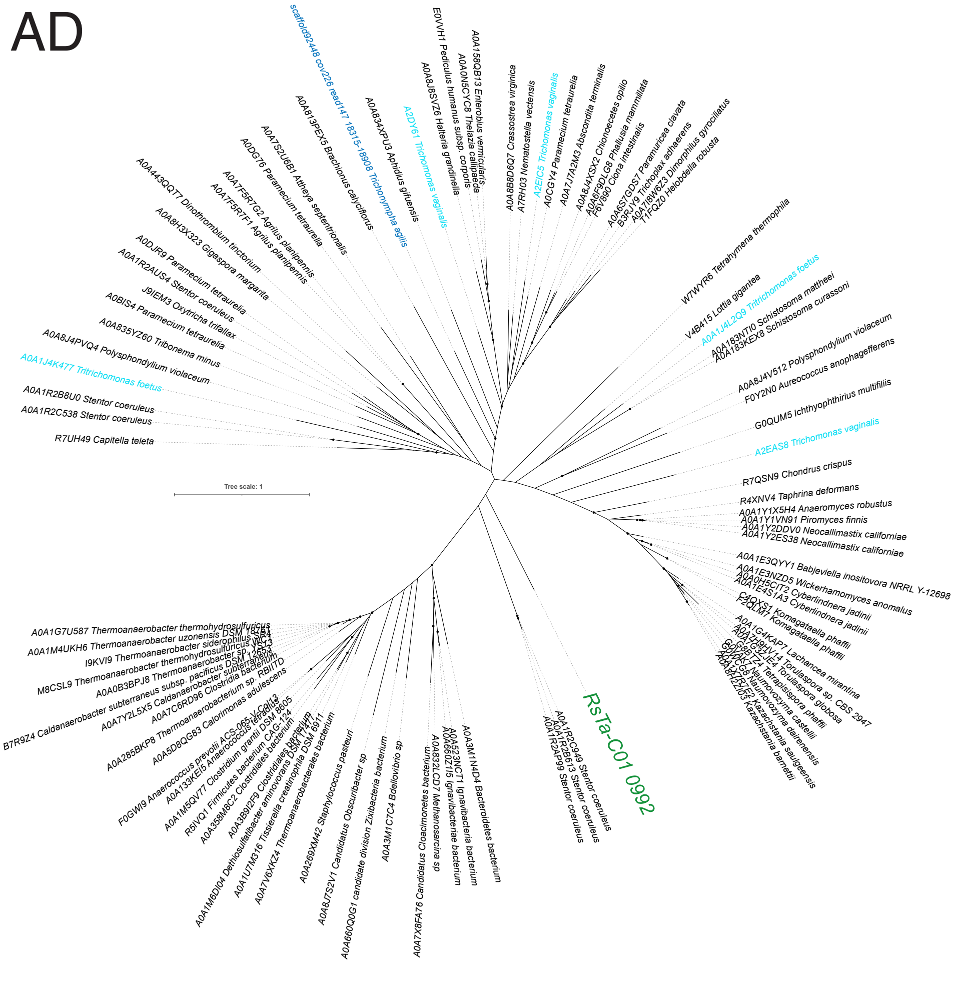
Tree scale: 1
RsTa-C01 0992
scaffold92448 cov226 read147 18315-18908 Trichonympha agilis

**Figure S9.** Phylogenetic positions of eukaryotic-like genes contained in the genomes of CfP3-15, NkDv07, and RsTa-C01. The following genes were used as queries to identify homologues as reference sequences. (**A**) CfP3-15_0049. (**B**) CfP3-15_0165. (**C**) CfP3-15_0235. (**D**) CfP3-15_0282. (**E**) CfP3-15_0445. (**F**) CfP3-15_0456. (**G**) CfP3-15_0548. (**H**) CfP3-15_0552. (**I**) CfP3-15_0565. (**J**) CfP3-15_0581. (**K**) CfP3-15_0591. (**L**) CfP3-15_0627. (**M**) CfP3-15_0653. (**N**) CfP3-15_0780. (**O**) CfP3-15_0790. (**P**) CfP3-15_0852. (**Q**) CfP3-15_0856. (**R**) CfP3-15_0934. (**S**) NkDv07_0114. (**T**) NkDv07_0121. (**U**) NkDv07_0512. (**V**) NkDv07_0536. (**W**) NkDv07_0615. (**X**) RsTa-C01_0144. (**Y**) RsTa-C01_0199. (**Z**) RsTa-C01_0208. (**AA**) RsTa-C01_0431. (**AB**) RsTa-C01_0774. (**AC**) RsTa-C01_0775. (**AD**) RsTa-C01_0992. Highly supported nodes (SH-aLRT value ≥ 80% and ultrafast bootstrap support value ≥ 95%) are highlighted with closed circles. The sequence IDs highlighted by green, blue, and light blue correspond to the proteins of the endosymbiotic *Clostridia* used to find homologs, proteins of the protist hosts, and proteins from parabasalid protists, respectively.

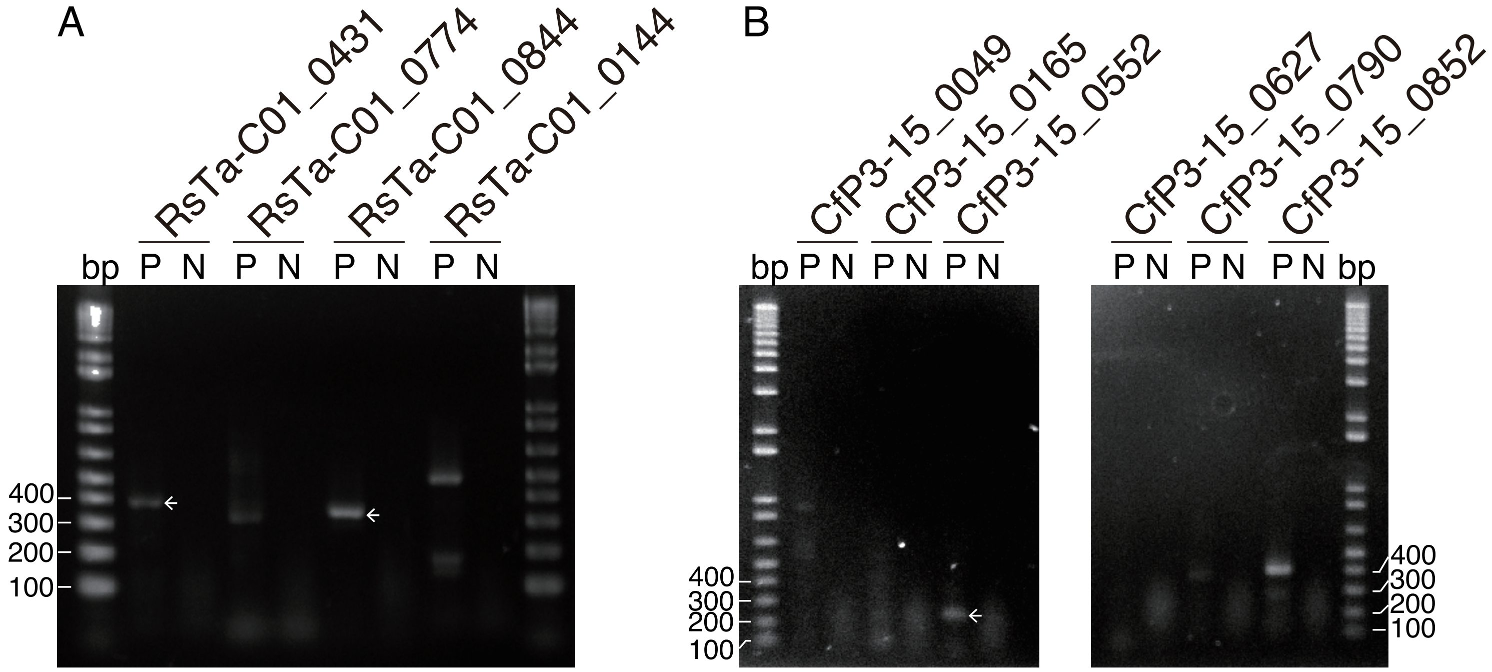


**Figure S10.** RT-PCR analysis of eukaryotic-like protein-coding genes. Agarose gel electrophoresis of RT-PCR products from total RNA of *R. speratus* guts (**A**) and *C. formosanus* guts (**B**). N, negative control (RNA samples without reverse transcription); P, products of RT-PCR using primer sets specific to the respective eukaryotic-like protein-coding genes (see Table S2). The amplification products of eukaryotic-like protein-coding genes are indicated by arrows, which were confirmed by Sanger sequencing.

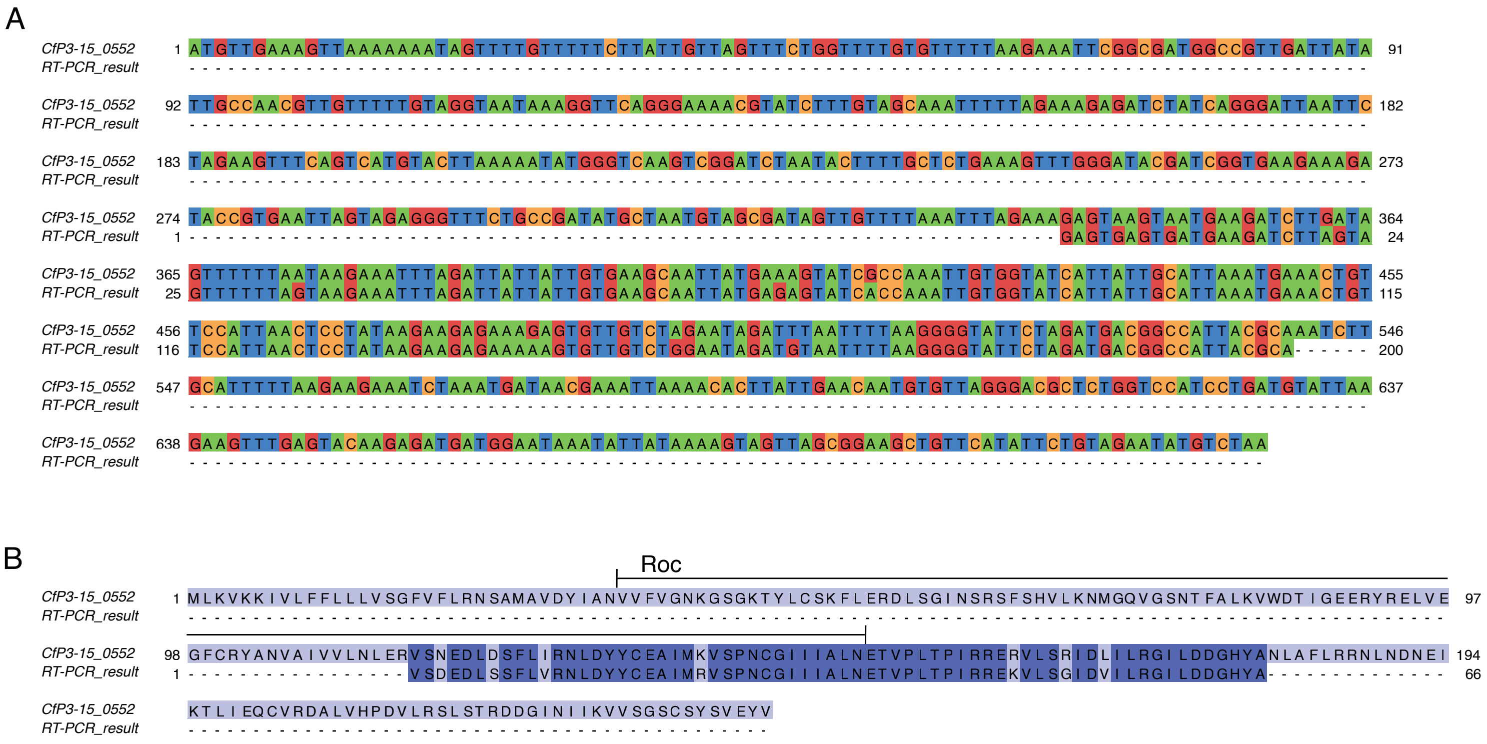


**Figure S11.** Comparison of a eukaryotic-like gene, CfP3-15_0552, in the CfP3-15 genome and that obtained by RT-PCR from *C. formosanus* guts. (**A**) Nucleotide sequences. (**B**) Amino acid sequences of **A**.

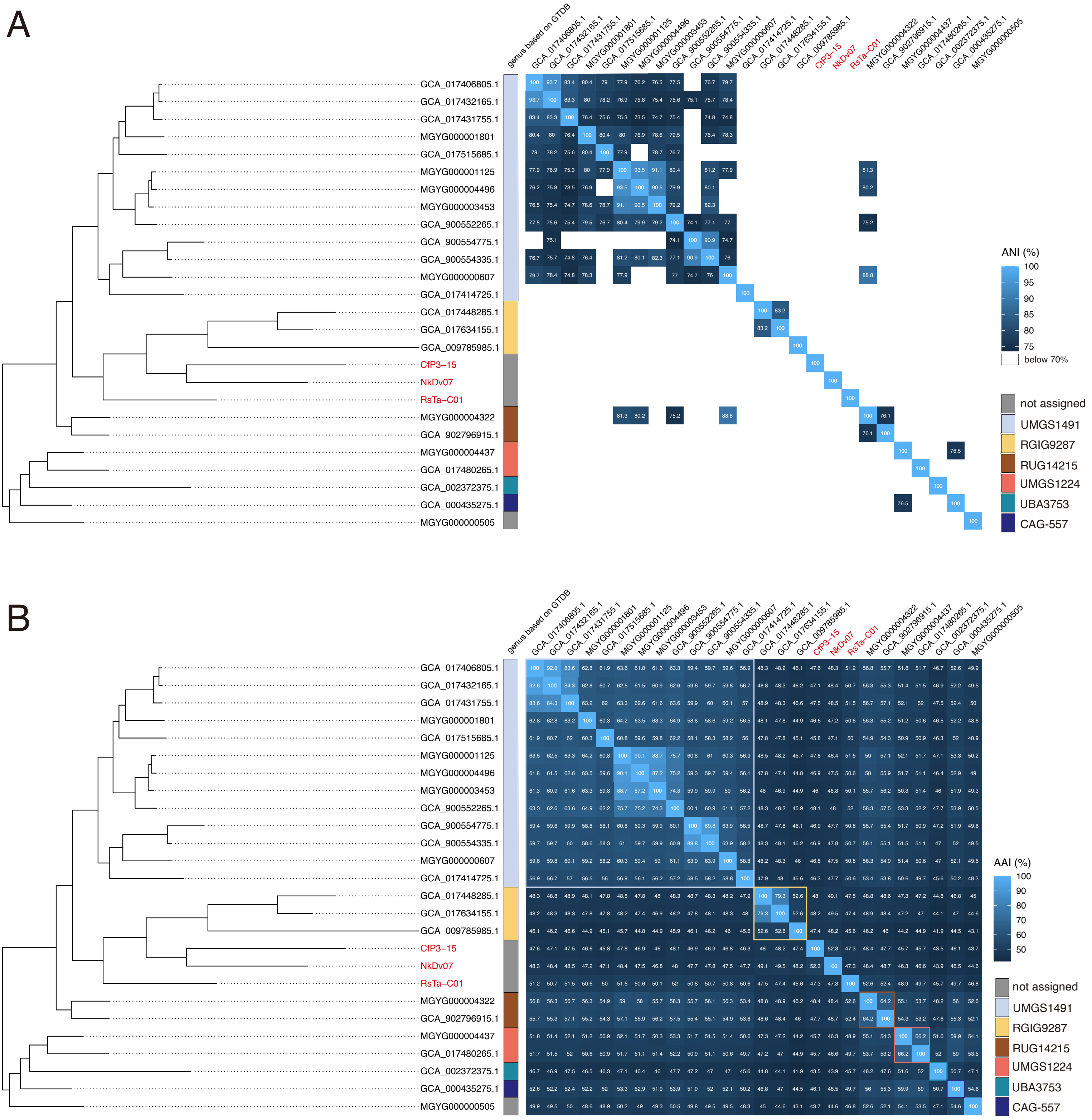


**Figure S12.** Pairwise ANI (**A**) and AAI (**B**) heatmap among the genomes in the small genome clade. The phylogenetic tree of the small genome clade is shown. In **B**, the genus-level clade based on GTDB r207 are indicated.

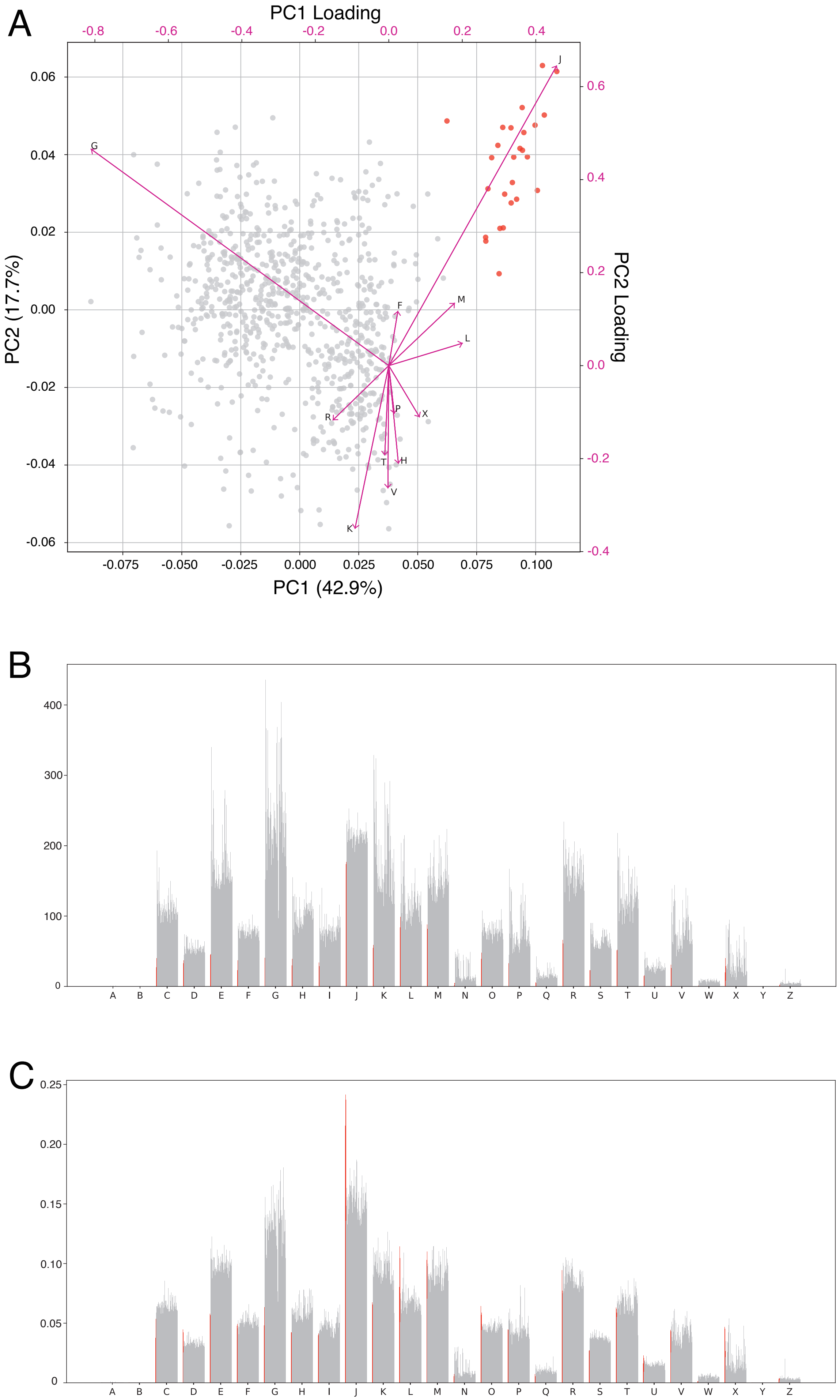


**Figure S13.** Principal component analysis of genomes of "Acutalibacteraceae" (Table S3) based on the relative abundance of the COG functional categories (**A**) and comparison of the number (**B**) and ratio (**C**) of the COG functional categories. Genomes belonging to the small genome clade are highlighted with red. Arrows indicate the loadings of each functional category. Several arrows are omitted to improve visibility. See also the legend to Fig. S5.

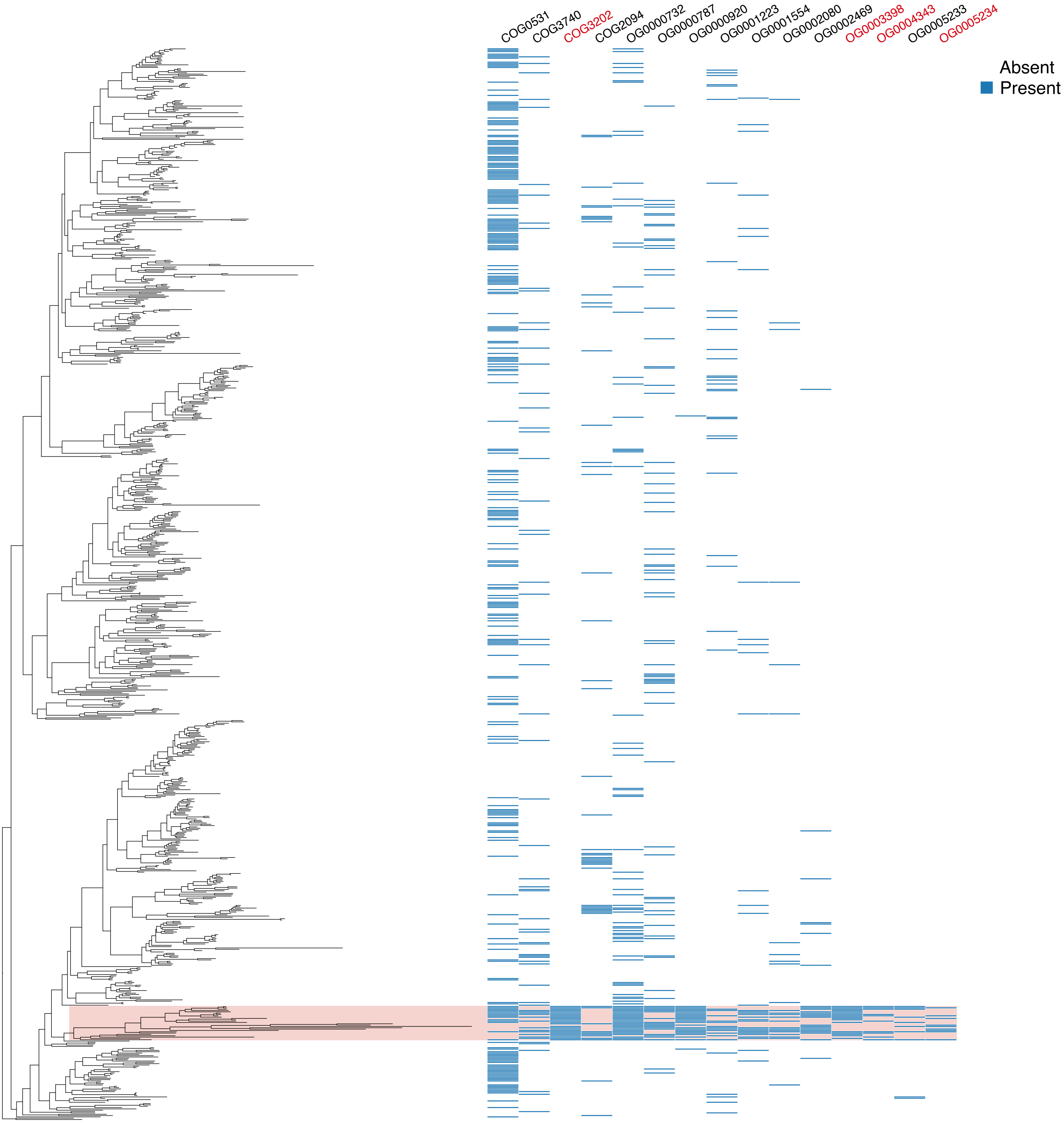


**Figure S14.** Distribution of genes acquired by the last common ancestor of the small genome clade, predicted in the gene flux analysis. The phylogenetic tree corresponds to that shown in Fig. 5A with the small genome clade being shaded in red.

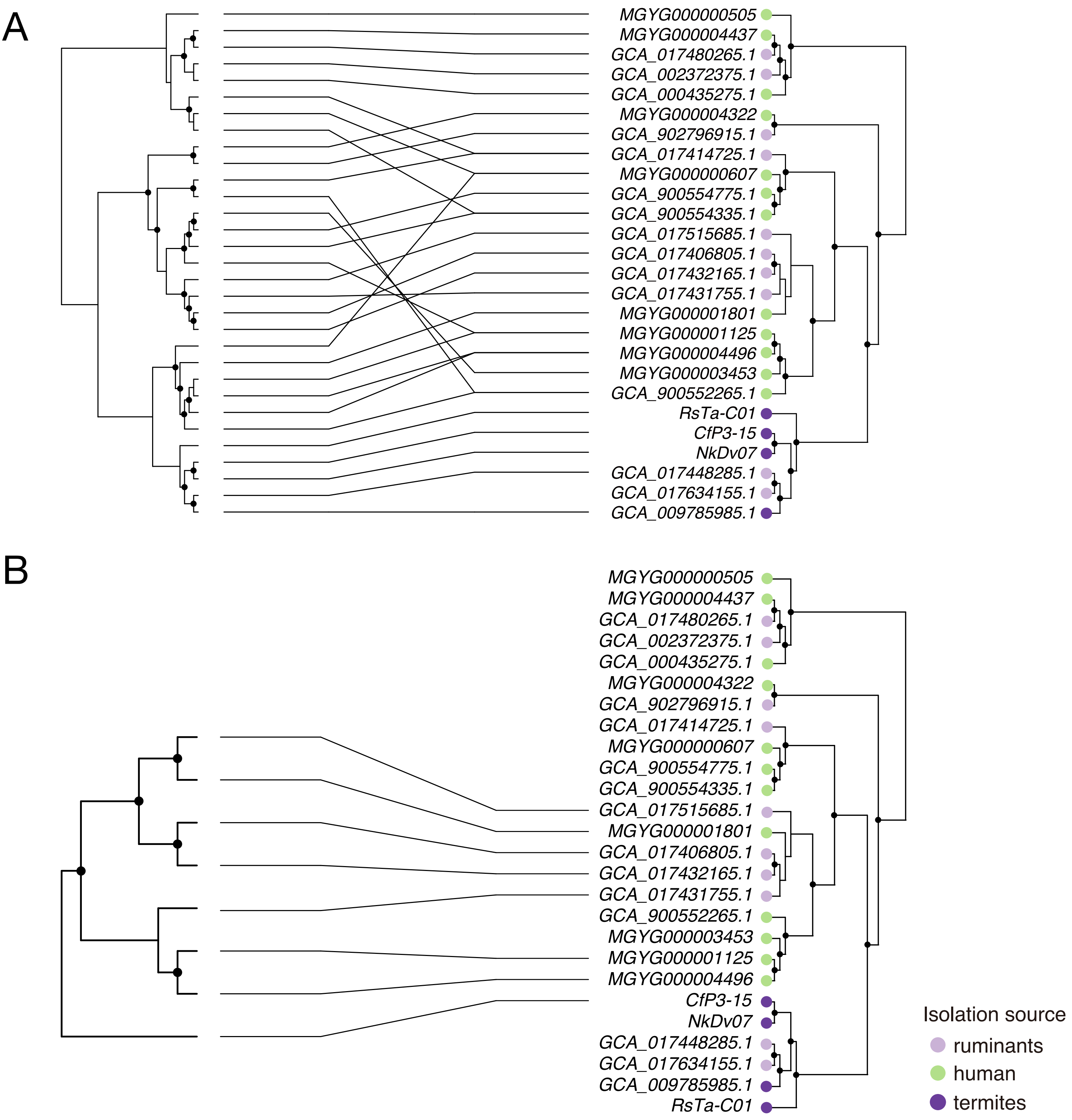


**Figure S15.** Comparison between the phylogenomic tree of the small genome clade and the gene tree of their ATP/ADP translocase. (**A**) Comparison of the *Clostridia* ATP/ADP translocase I (left) with the phylogenomic tree (right). (**B**) Comparison of the *Clostridia* ATP/ADP translocase II (left) and the phylogenomic tree (right). Highly supported nodes (SH-aLRT value ≥ 80% and ultrafast bootstrap support value ≥ 95%) are indicated with a closed circle.

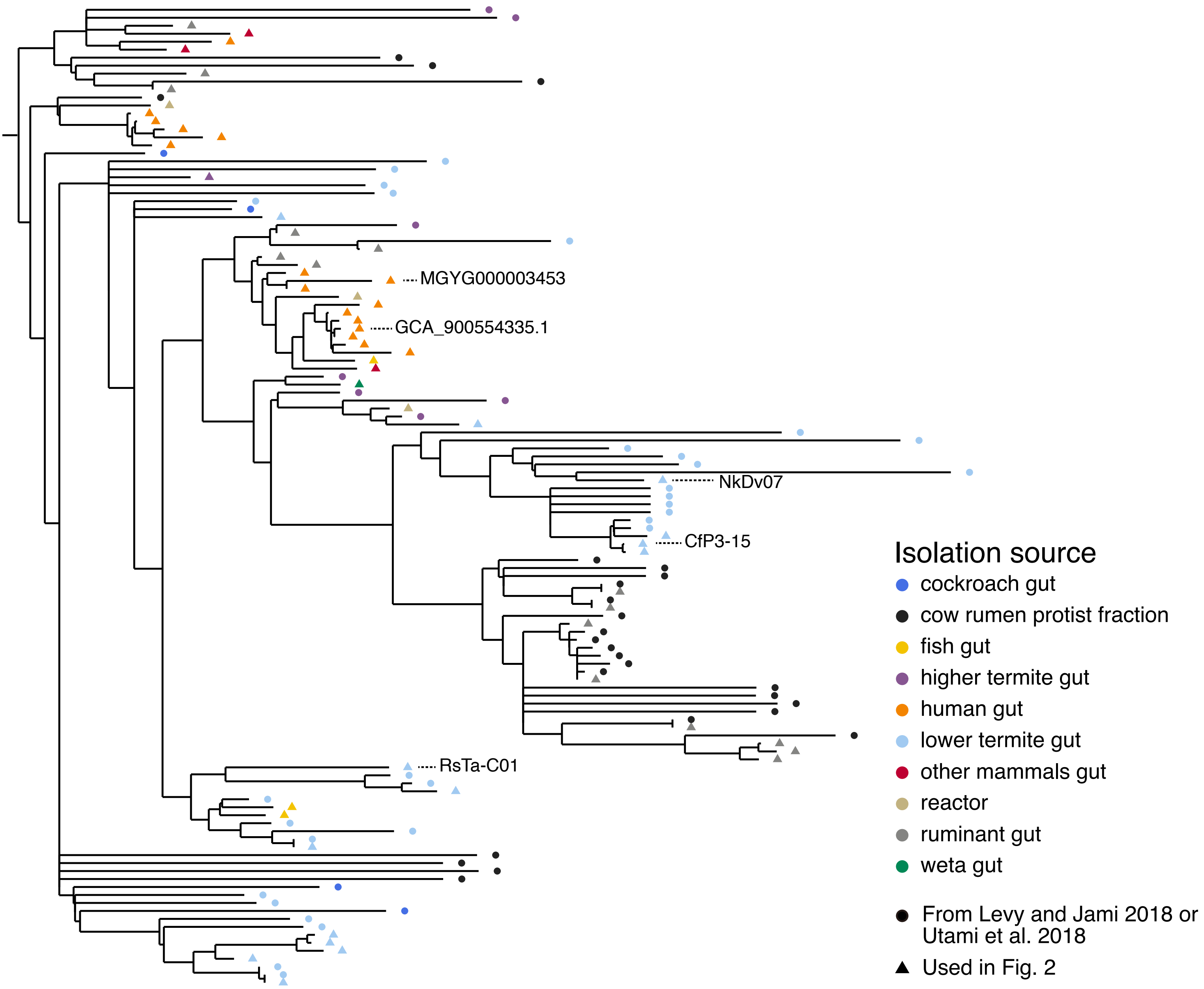


**Figure S16.** Phylogenetic positions of 16S rRNA gene amplicon sequences derived from the protistan fraction of a cattle rumen [31] and the entire guts of termites and cockroaches [32].
